## Supplementary material for "Genomic Analysis of Two Phlebotomine Sand Fly Vectors of *Leishmania* from the New and Old World": Manuscript File

### **SUPPLEMENTAL METHODS**

#### Manual Annotation Methods

*Digestion Genes*

*Peptidases, GHF13 and GHF18*

The Peptidase and Glycoside Hydrolase Families (GHF) (defined as described in the MEROPS and CAZy databases, respectively) in sand flies genomes were characterized using FAT software [1], which integrates HMMER (http://hmmer.janelia.org/) and BLAST+ tools [2] to filter the initial dataset and perform automatic annotation. The filter step used the HMG-box conserved domain (Pfam code varied between families) to identify and extract only proteins containing such a domain in both datasets.

*N-acetylhexosaminidases*

VectorBase [3-5] was screened for sand fly n-acetylhexosamisnidases through tBlastn searches against *Tribolium castaneum* homologs [6]. Genbank Accession numbers: *Drosophila melanogaster* (DmNAG1: NP_523924; DmNAG2: NP_525081; and DmFDL: NP_725178), *Anopheles gambiae* (AgNAG1: XP_315391; AgNAG2: XM_307483; AgFDL: XP_308677; and AgHEX: XM_319210), *Aedes aegypti* (AaNAG1: EAT43909; AaNAG2: EAT40440; AaFDL: EAT36388; and AaHEX: EAT43655), and *T. castaneum* (TcNAG1: ABQ95982; TcNAG2: ABQ95983; TcNAG3: ABQ95984; TcFDL: ABQ95985; TcHEX1: XM_970563; TcHEX2: XM_970567; and TcHEX3: XM_970565). Conserved domain architecture was identified through CDD, Prosite, and Pfam database searches.

*Chitin deacetylases*

VectorBase was screened for sand fly chitin deacetylase (CDA) through tBlastn searches against *T. castaneum* homologs [7]. Genbank Accession numbers: *T. castaneum* (TcCDA1: ABU25223; TcCDA2: ABU25224; TcCDA3: ABW74145; TcCDA4: ABW74146; TcCDA5: ABW74147; TcCDA6: ABW74149; TcCDA7: ABW74150; TcCDA8: ABW74151; and TcCDA9: ABW74152), *An. gambiae* (AgCDA1: EAA00275; AgCDA2: EAA43313; AgCDA3: EAA12484; AgCDA4: EAA06323; and AgCDA5: EAA12207); *Ae. aegypti* (AeCDA1: EAT45316; AeCDA2: EAT45315; AeCDA3: EAT42807; and AeCDA4: EAT42983), and *D. melanogaster* (DmCDA1: AGB94755; DmCDA2: AAF49121; DmCDA3: NP_609806; DmCDA4: AAF50937; DmCDA5: ABV53594; and DmCDA9: AHN56315). Conserved domain architecture was identified through CDD, Prosite, and Pfam database searches.

*Peritrophin-like proteins*

VectorBase [3-5] was screened for *Lutzomyia longipalpis* peritrophins by carrying out tBlastn searches against peritrophin-like proteins from many insects, for instance: *Lucilia cuprina* (AAC37261, AAC70784, AAK01058, AAB38414, and AAB70878); *An. Gambiae* (AAV31069); *Chrysomya bezziana* (AAK01057, AAB86623, and AAD25103); *D. melanogaster* (NP_524982 and NP_524983); *Trichoplusia ni* (AAC47557 and AAC47556); *Armigeres* subalbatus (AY439786); *Ae. Aegyti* (AAL05409, AAM94157, AAM94156, AAM94155, and AAM94154); and the sand fly peritrophins identified in cDNA libraries [8, 9]. This initial screening gave rise to 10 unique *Lu. longipalpis* peritrophins. Such sequences were translated, and their protein sequences were used to further search Bfor novel sand fly peritrophins. Genbank Accession numbers for proteins displayed in the phylogenetic tree: *T. castaneum* peritrophin access numbers were retrieved from [10] whereas available sand fly peritrophin access numbers were retrieved from [8, 9]. Conserved domain architecture was identified through CDD, Prosite, and Pfam database searches.

<u>*Immunity Genes*</u>

*JAK/STAT*

OrthoMCL was used to search for orthologous genes between the 2 sand fly species and 6 other insect species (*D. melanogaster, Anophles albimanus, An. Gambiae, Ae. Aegypti, Culex quinquefasciatus*, *Rhodnius prolixus*). The genome sequences for these organisms were downloaded from VectorBase [3-5], with the exception of *D. melanogaster*, which was downloaded from FlyBase [11]. Following the OrthoMCL comparisons, a list of known immunity genes and ID's were downloaded from FlyBase for *D. melanogaster*. There were a total of 5 JAK/STAT genes found using this method; Hopscotch, Domeless, STAT92E, Upd3, SOCS36E. The ID's were used to search for the OrthoMCL clusters for the 5 JAK/STAT genes. From the OrthoMCL gene cluster ID, it was possible to find the corresponding ID's for the 2 sand fly species. 5 VectorBase ID's were found in each sand fly species for the 5 JAK/STAT genes. VectorBase searches were performed to retrieve the sand-fly ID's, each gene was downloaded to assess the quality of the gene model on VectorBase. In order to assess the quality of the gene model, Artemis [12] was used. The gene models were checked to make sure that that the coding sequences started and ended with the correct codons. In addition, the coding sequences were checked to make sure that there were no stop codons in the coding sequences themselves. All of the gene models were of good quality and everything was in the right place. Once the gene quality had been assured, the gene ID's could then be confirmed.

To find more JAK/STAT genes, the Insect Immunity Database (Bordenstein Lab, NSF DEB-1046149) was used. This website contains a list of immunity genes from different insects, the JAK/STAT gene list has been mainly compiled using *D. melanogaster*. 2 genes were found from this list that were also present in our OrthoMCL gene clusters, these were; p38b, and mekk1. The genes were again found using VectorBase and the gene models were confirmed using Artemis.

*Prophenoloxidase genes*

The same OrthoMCL gene clusters were used as in the JAK/STAT gene annotations. Using the FlyBase immunity gene list, we found 3 Prophenoloxidase genes. These 3 genes turned out to all belong to the same gene cluster. The gene models were downloaded for all of the 3 prophenoloxidase genes and they were assessed using Artemis. All were confirmed to be good quality. Another phylogenetic tree is being created using these genes, more phenoloxidase genes are being searched for currently.

*Toll Signaling Genes*

The annotation was focused on the core signaling molecules in the Toll pathway and also on the upstream pathogen recognition receptors including PGRPs and GNBPs. This manual annotation started by using blastp against the two sand fly genomes from Vectorbase with the *Drosophila* protein sequences of the core Toll signaling genes. The *Drosophila* genes and protein sequences were obtained from FlyBase (Toll: FBgn0262473; PGRP-SA: FBgn0030310; GNBP1: FBgn0040323; Tube: FBgn0003882; Pelle: FBgn0010441; Myd88: FBgn0033402; Cactus: FBgn0000250; Dorsal: FBgn0260632). Gene models of the top hits from the blast were checked by Artemis to ensure the correct annotation of coding region. Subsequently the protein (transcribed) sequences of the hits were used to align with *Drosophila* counterpart by ClustalW [13]. A score above 20 was set as the arbitrary threshold to confirm it was a 'true' homologue. Furthermore, SMART [14] was used to confirm that the characteristic protein domains essential for the gene function are present in the sand fly homologues.

*IMD/ROS genes*

For manual annotation, blast programs were used with IMD/ROS core genes from *Drosophila* or in-house sequences from *Lu. longipalpis* as query. The *Drosophila* genes and protein sequences were obtained from Flybase. Gene models from the blast highest hits were checked by Artemis to ensure the correct annotation of coding region.

*Galactose-binding proteins*

VectorBase was screened for sand fly galectins through tBlastn searches against fly, mosquito, and sand fly homologs retrieved from Genbank. Genbank accession numbers: *Crassostrea virginica* (CVgalec: ABG75998.1), *Ae. Aegypti* (AeGal1: XP_001656933; AeGal2: XP_001655870; AeGal3: EAT44461; AeGal5: XP_001657305; AeGal6: EAT43258; AeGal8: XP_001662146; AeGal11: EAT44861; AeGal12: EAT38239; AeGal13: EAT38241; AeGal14: EAT38240; and AeGal_ XP_001650704: XP_001650704), *An. Gambiae* (AgGal1: EAA14815;AgGAl2: EAA06231; AgGAl3: EAA10348; AgGal4: EAL41552; AgGal5: EAA05052; AgGAl6: EAA13122; AgGal7: EAA13132; AgGal8: EAA09838; AgGal9: EGK97239; and AgGal10: EAL38639), *D. melanogaster* (DmGalA: AAF51564; DmGalB: ACZ94129; DmGalD: AFH03490; DmGalE: ACZ94130; DmGalF: AGB92325; DmGal_CG5335: AAF57667; DmGal_CG11374: AAF51563; DmGal_peroxin23: AAF49038; DmGal_CG13950: AAF51443; and DmGal_CG14879: AHN57362), P. papatasi (PpGalecA: AY538600), *Lu. longipalpis* (LuloGalecR: ABV60341), and *Biomphalaria glabrata* (BgGal_ABQ09359: ABQ09359). Conserved domain architecture was identified through CDD, Prosite, and Pfam database searches.

*MAP Kinases*

A bidirectional BLAST method was applied to identify new orthologous genes belonging to the ERK, JNK and p38 MAPKs pathways. All RefSeq proteins of *Homo sapiens*, *An. gambiae*, *D. melanogaster* and *Caenorhabditis elegans* were downloaded from Genbank and a set of 79

MAPKs was made by selecting representative amino acid sequences from these species. Set of homologue sequences used as query: NP_001036065.1, NP_003945.2, XP_310236.3, NP_109587.1, NP_005912.1, NP_003609.2, XP_307879.4, XP_309868.4, XP_316357.4, XP_316502.4, XP_318144.4, XP_322064.4, NP_572458.3, NP_477163.1, NP_477353.1, NP_477361.1, NP_491683.1, NP_491087.1, NP_492620.1, NP_501365.1, NP_504721.1, NP_509682.1, NP_524080.1, NP_511098.1, NP_002742.3, NP_620602.2, NP_620448.1, NP_620581.1, NP_620637.1, NP_620407.1, NP_002743.3, NP_660143.1, NP_660186.1, NP_002437.2, NP_663304.1, NP_005195.2, NP_004570.2, NP_723541.1, NP_727335.1, NP_610817.1, NP_611399.2, NP_741431.1, NP_741537.1, NP_477089.3, NP_001001671.3, NP_649137.3, NP_006292.3, NP_872069.1, NP_004324.2, NP_001193733.1, NP_001229243.1, NP_001229488.1, XP_310813.5, NP_976226.1, NP_650750.4, NP_001645.1, NP_002410.1, NP_002745.1, NP_002739.1, NP_002871.1, NP_003001.1, NP_995972.1, NP_996277.1, NP_004663.3, NP_002960.2, NP_149132.2, XP_005248576.1, XP_005250103.1, XP_005258356.1, NP_002746.1, XP_006715545.1, XP_006723285.1, NP_009112.1, NP_001021270.1, NP_001022584.1, NP_741430.3, NP_001024971.1, NP_002737.2, NP_620590.2.

This query dataset was blasted against the *P. papatasi* and *Lu. longipalpis* genome. The genomic regions with highest similarity to MAPKs were extracted and blasted against all downloaded RefSeq proteins. Only the genomic regions which matched to MAPKs in the second blast round were considered as a MAPK gene locus. The exons were identified by the comparison of the genes found to the RNA-seq data assembled using different methodologies. The RNA-seq assemblies were also screened by a similar bidirectional blast approach.

*TGF-beta*

Manual annotation was performed by identifying *D. melanogaster* ortholog genes related to TGF-beta molecules from FlyBase (http://flybase.org; dcapentaplegic: FBgn0000490; maverick: FBgn0039914 ; mother against decapentaplegic: FBgn0011648; smad on X: FBgn0025800; punt: FBgn0003169 ; saxophone: FBgn0003317; baboon: FBgn0011300; thickveins: FBgn0003716; tolloid: FBgn0003719; fusel: FBgn0039932; inwardly rectifying potassium channel 2: FBgn0039081; supernumerary limbs: FBgn0267841; spichthyin: FBgn0032451; short gastrulation: FBgn0003463). These sequences were used as a query using BLAST search tools against the two sand fly genomes from VectorBase. Gene models identified in the sand fly genomes were checked by Artemis or Web Apollo and metadata was added.

*Circadian Rhythm Genes*

Using a bioinformatic approach, we identified several genes involved in the control of circadian rhythms, behaviour and sensory ecology in other insect species. Orthologous sequences, mostly from *D. melanogaster*, *An. gambie*, *Ae. aegypti*, and *Cx. quinquefasciatus* were used as a query to perform HMMER, BLASTP, and TBLASTN searches on the *Lu. longipalpis* and *P. papatasi* genomes and on their predicted protein sets. Subsequently, some gene models were improved manually or using gene prediction programs such as Augustus [15] and Fgenesh+ [16]. Final gene models have been created using Artemis [17]. The clock and behaviour related genes identified and their main characteristics are detailed in Tables S1 and S2.for *Lu. longipalpis* and *P. papatasi*, respectively.

*cycle* (*cyc*), *Clock* (*Clk*), and *vrille* (vri) genes were experimentally obtained. We first designed degenerate primers in conserved regions of vertebrate and invertebrate ortholog genes (*Mus musculus*, *Gallus gallus*, *Danio rerio*, and *D. melanogaster*; GenBank accession numbers AAC53200, AAF26365, AAD27749, and AAF50516). Based on these first sequences, new specific primers were made to obtain further sequences. The 5' and 3' ends were obtained using RACE techniques [18]. cDNA sequences were then used as query to search and edit the *Lu. longipalpis* and *P. papatasi* gene models automatically annotated.

*Lu. longipalpis* and *P. papatasi* genes that belong to serine/threonine phosphatase (Pp1-alpha, Pp1-beta, Pp2-alpha, Pp2-beta, Pp4, and PpV-6, and Pp7) and photolyase superfamilies (cry-1, cry-2, phr, (6-4) phr) were differentiated by performing phylogenetic analyses using vertebrate and invertebrate ortholog genes. To do so, alignments of vertebrate and invertebrate ortholog sequences were performed in ClustaL X [19], manually edited in Jalview v2.6.1 [20] and subsequently aligned using G-INS-I strategy in MAFFT [21]. For the phylogenetic reconstruction, different evolutionary models (JTT, LG, DCMut, MtREV, MtMam, MtArt, Dayhoff, WAG, RtREV, CpREV, Blosum62, and VT) were tested using the ProtTest v2.4 [22]. Finally, maximum likelihood trees with 1,000 bootstrap pseudo-replicates of the original data were built in MEGA6 [23].

In the case of the pickpocket (PPK) and transient receptor potential (TRP) cation channel protein families, interative searches were performed with each new obtained protein sequence as query until no new genes were identified in each major subfamily or lineage. Once gene modelling process was finished, the number of predicted transmembrane domains in sand fly TRP and PPK sequences was established using TOPCONS [24]. Besides, functional domains were identified using Pfam v27.0 database [25]. Finally, a phylogenetic analysis with sequences from these protein families and those from *D. melanogaster* was performed. TRP sequences from *Drosophila* melanogaster were obtained from Peng et al. [26]. In the case of PPKs, fruit fly sequences were obtained from Zelle et al. [27]. To build the TRP and PPK phylogenetic trees, we aligned and edited the sequences the same way as described above.

<u>*Chemosensory Genes*</u>

The three major chemoreceptor gene families (odorant, gustatory, and ionotropic receptors) were manually annotated using *Anopheles gambiae* and *Drosophila melanogaster* gene models as queries for tBLASTn analysis of the *P. papatasi* and *Lu. longipalpis* genome assemblies in VectorBase [3, 5, 28]. Exon-intron boundaries, where ambiguous, were assessed using the splice site prediction tool [29] on the Berkeley Drosophila Genome Project web site (http://www.fruitfly.org/seq_tools/splice.html). Gene models were evaluated by multiple alignment of each receptor family with the *An. gambiae* and *D. melanogaster* peptide sequences using ClustalX [19, 30]. When necessary, gaps were filled and indels evaluated by BLASTn analysis of the sequence read archives (SRAs) in NCBI (*Lu. longipalpis*: SRX016812, SRX016813, SRX1537403; *P. papatasi*: SRX027115, SRX027117, SRX1009382). Partial gene models encoding less than 300 amino acids were omitted from the analysis. MEGAHIT [31] used to produce *de novo* assemblies for the field isolates Jacobina 14 and Jacobina 23 to search for several genes suspected to be missing from the *Lu. longipalpis* reference assembly. See [32] for additional details.

*RNA Genes and MicroRNAs*

Identification and characterization of RNAi genes - *D. melanogaster* RNA interference core proteins involved in siRNA, miRNA and piRNA pathways were used to search for similarity using BLASTP against all predicted polypeptide genes in *Lu. longipalpis* and *P. papatasi* genomes. Identification of sand fly predicted polypeptides was confirmed by similarity searches against annotated proteins in the *Drosophila* genome using BLASTP. RNAi proteins identified by this strategy were analyzed to determine size and domain organization with InterProScan and gene were refined using SoftBerry and manual curation. Identification of microRNAs and piRNAs - Small RNA libraries were constructed from adult female *Lu. longipalpis* and deep sequenced using the Illumina platform. Small RNAs were mapped against the *Lu. longipalpis* genome using Bowtie. miRNAs were identified using miRDeep2 [33]. Candidate precursors were manually curated using 4 different criteria: (1) Size distribution of mapped reads on precursors, (2) Homogeneity of 5' end of mapped reads, (3) detection in more than one library and (4) conservation in other insects. To identify piRNA clusters, *Luztomya* scaffolds were separated in 2000 nt bins using in-home perl scripts that were used as reference for mapping reads libraries using Bowtie [34]. 2000 nt bins with a profile of mapped reads showing size distribution between 24 and 30 nt were selected. These were analyzed for ping-pong signature [35], nucleotide enrichment (weblogo) and offset between reads mapped in positive and negative strands.

*G-Protein Coupled Receptors*

A novel classifier pipeline was utilized to identify the GPCRs from the *P. papatasi* and *Lu. longipalpis* genome assemblies at VectorBase using the set of known GPCR peptides from *Ae. aegypti, An. gambiae, Apis mellifera, D. melanogaster, H. sapiens,* and *Pediculus humanus* [36]. The identified sand fly GPCR sequences were aligned using BLAST against the non-redundant protein sequences (nr) in NCBI. Putative functions were assigned based on the best and the most informative BLAST hits, in most of the cases the later was from *D. melanogaster*. The sand fly sequences were aligned with homologous genes, mostly from insects used in the classifier pipeline, using MultAlin [37] and trees from Clustal Omega [38]. The sand fly gene models were manually corrected using the annotation tool Apollo. Identified and manually annotated GPCRs were used to search (tBLASTn) the sand fly genome scaffolds for additional GPCR genes. When available, the gene models were confirmed using transcript evidence from Expressed Sequence Tags (ESTs) and RNA sequencing (RNAseq).

*Cytochrome P450s*

To identify all potential members of the cytochrome P450 superfamily we aligned the predicted gene sets of *Lu. longipalpis* (LlonJ1.6) and *P. papatasi* (PpapI1.6) to a reference collection of 2,942 curated CYPs from a wide variety of arthropods [39] using BLAST v2.10.0+ [2]. Since it is known that automated gene prediction methods quite frequently contain mistakes we proceeded to manually curate the automatically predicted CYP genes. To this end, we mapped publicly available

*Lu. longipalpis* (SRR535765) and *P. papatasi* (ERR3714261, ERR3714267, ERR3714268) publicly available RNAseq data on the reference genomes using Hisat2 v2.1.0 [40] with parameters "--dta-cufflinks". Subsequently, Bam2wig v3.0.1 was used with default parameters to generate BigWig files, in order to obtain transcription levels along the genome [41]. The BigWig files were then loaded as separate tracks in a locally deployed instance of the Apollo genome browser (v2.6.0) [42] that contained the sand fly genome sequences as well as the automatic gene predictions. We also identified CYP genes that were missed by automatic gene prediction using tblastn searches of the reference CYP set against each genome assembly. The Apollo genome browser was used for manual curation of the P450 genes.

A phylogenetic analysis of the *Lu. longipalpis* and *P. papatasi* CYPs was conducted using the curated CYPome of the major malaria vector *An. gambiae [39]* as reference. More specifically, we first aligned the amino acid sequence of the CYPomes of the three species using MAFFT v7.490 [43] with the "auto" parameter. Next, we trimmed the alignment using Trimal v1.2rev59 [44] with parameters "-gt 0.50". IQ-TREE 1.6.12 [45] was used with parameters "*-alrt 5000 -bb 5000 -m MFP*" to reconstruct a maximum likelihood phylogeny with 5,000 bootstraps using the ultrafast bootstrap algorithm UFBoot2 [46]. Amino-acid model selection was performed using ModelFinder [47]. The human CYP51A1 was used as an outgroup. Finally, tree visualization and post-processing was performed using Evolview v3 [48].

*Salivary Protein Genes*

Forty-nine *P. papatasi* and 35 *Lu. longipalpis* putative salivary genes deposited at NCBI [49] were mapped to the sand fly assemblies using BLAST.

*Heat Shock Protein Genes*

Comparison of *D. melanogaster* proteins with GO terms relating to heat shock and response to hypoxia were compared by BLAST to the scaffolds and predicted genes in *Lu. longipalpis* and *P. papatasi*. Genes that were identified as possible orthologs were then compared by BLAST to the arthropod-specific BLAST database using CLC Genomics Workbench 7.

*Cuticular Protein Genes*

Sequence motifs characteristic of various families of cuticle proteins [50] were blasted (tBLASTn) [51] against the official gene set for *Lu. longipalpis* (LlonJ1.4) and *P. papatasi* (Ppal1.4), which were obtained from VectorBase (http://vectorbase.org). Potential cuticle proteins were further analyzed with CutProtFam-Pred, a tool for predicting cuticular protein families described by Ioannidou et al. [52], to assign genes to gene families. To find the closest putative homolog to cuticle protein genes from *Lu. longipalpis* and *P. papatasi*, genes were searched against the official gene sets for *Ae. aegypti* and *An. gambiae* downloaded from VectorBase as well as *D. melanogaster* genes obtained from FlyBase (http://flybase.org). The gene with the lowest e-value was considered the closest putative homolog.

*Vitamin Metabolism Genes*

Known protein sequences associated with vitamin metabolism from *D. melanogaster* were acquired through batch download and compared using BLASTagainst peptides from *P. papatasi* and *Lu. longipalpis* to identify the highest e-value and bit score matches through the use of CLC Genomics Workbench 7. These peptides from *P. papatasi* and *Lu. longipalpis* were subsequently compared against databases for *An. gambiae* and *Ae. aegypti* to identify specific orthologs and to confirm their identity.

*Hormonal Signaling Genes*

*D.melanogaster* proteins with GO terms relating to insulin signaling processes were compared to the hypothetical scaffolds of the two sand fly species *Lu. longipalpis* and *P. papatasi* using BLAST. Predicted genes that were identified as possible orthologs were then compared by BLAST to the arthropod-specific BLAST database using CLC Genomics Workbench 7 to validate the identity of each gene.

*Antioxidant Genes*

Comparison of *D. melanogaster* proteins with GO terms relating to antioxidant activity and response to oxidative stress were compared using BLAST to predicted genes of the two sand fly species *Lu. longipalpis* and *P. papatasi* allowed for identification of orthologs in each species. Predicted gene that were identified as orthologs were then compared by BLAST to an arthropod-specific BLAST database using CLC Genomics Workbench 7 for gene identity validation.

*Aquaporin Genes*

Aquaporins were identified based upon BLAST comparison to those from mosquitoes and other higher flies [53]. Putative aquaporin genes were subsequently compared to an arthropod-specific BLAST database using CLC Genomics Workbench 7 to validate each as an AQP.

*Novel Viruses*

Two strategies were used to investigate the possible presence of novel viruses in the sand fly genomes. First, reads were aligned onto known viral sequences. The Taxonomy Browser of NCBI was used to select 1795534 viral sequences from GenBank. 1791477 of these proved suitable for use. The GS Reference Mapper 3.0 software (454 Life Sciences) was used to align the *Lutzomyia* and *Phlebotomus* *.sff files onto the downloaded viral GenBank fraction. Reads of less than 75 bases in length were discarded and the hit match threshold was set at 85% identity. *Lutzomyia* Illumina *.fq files were first groomed using read cleaner (Gatherer, unpublished) and then aligned onto the same downloaded viral GenBank fraction using BWA [54] and Bowtie [34] implemented in valet (Gatherer, unpublished). Output was viewed in Tablet [55].

Secondly, the NCBI Taxonomy Browser was used to search for a reference genome from each family of viruses as defined by the International Union for Taxonomy of Viruses (ICTV), retrieving 101 reference genomes. BLASTN was then used to search the *Lutzomyia* and *Phlebotomus* assembled transcript sets for homologous sequences. A list was compiled of hits at expectation threshold of $<10^{-3}$ and match longer than 100 nucleotides, produced a total of 11 non-redundant hits for *Lutzomyia* and 6 for *Phlebotomus*. Each candidate hit was then used to search the whole of GenPept by BLASTX to determine its nearest protein relative. In addition, BLASTN was used with an expectation threshold of <$10^{-3}$ to match individual genome contigs to the viral GenBank fraction described above. Genome contigs were also BLASTed against bracoviral and retroviral fractions of GenBank.

### SUPPLEMENTAL RESULTS: MANUAL ANNOTATION

#### Transposable Elements

Transposable elements (TEs) are coding and non-coding sequences which have a mechanism allowing them to jump within a genome. They are undoubtedly important to analyze and categorize, because they usually make up a significant part of a genome and because of their mobility they thought to be one of the driving forces of the evolution. Genome sizes differ greatly among insect genomes, sometimes even in closely related species. The genome size of *Phlebotomus papatasi* (347 Mb) is approximately twice the size of *Lutzomyia longipalpis* (154 Mb). We found that *P. papatasi* genome is composed of 5.65% of TE derived sequences while the *Lu. longipalpis* genome contains only 0.57%. As might be expected due to differences in the genome size, all TE classes and orders were expanded in the *P. papatasi* genome comparted to *Lu. longipalpis*. The amount of TE sequence is likely underestimated, due to the nature of the short-read assembly process producing many contigs, sometimes shorter than the length of full-length TEs. Analysis of scaffolds did not improve TE yield, most likely due to the gaps in the assembly. Sequenced BACs were also screened for the presence of TEs and were useful in identification of several full-length retrotransposons.

##### *Long-terminal repeat retrotransposons (LTR)*

We have identified canonical elements belonging to two of the main superfamilies of LTR retrotransposons in both the *P. papatasi* and *Lu. longipalpis* genomes: gypsy, and pao/bel; and elements corresponding to the Copia and DIRS superfamilies only in *P. papatasi*.

The gypsy group is the most diverse, containing elements belonging to five of the previously characterized lineages (gypsy, Mag, CsRn1, Osvaldo-like, Mdg1 and Mdg3) and a phylogenetically new lineage represented by sequences obtained from both genomes. These sequences grouped together with 100% bootstrap value and they constitute full-length elements, containing all the functional domains (RT, RH, RVE). The CsRn1 lineage was not identified in either sand fly genome.

The Bel-Pao superfamily has been previously classified into seven lineages (Pao, Sinbad, Bel, Tas, Suzu, Flow and Dan) which tend to cluster with the host species phylogeny [1, 2]. Most of the sequences belonging to this superfamily in both sand fly genomes correspond to the Bel lineage while a minority (eight sequences from the *P. papatasi* and six from the *Lu. longipalpis*) correspond to the Pao lineage. None of the other lineages within this superfamily are present in these genomes. One sequence from the *P. papatasi* genome clustered with the *copia* reference sequences and five with the DIRS references. Neither *copia* nor DIRS elements were identified in the *Lu. longipalpis* genome.

The Gypsy and Bell groups are represented mostly by the same families in both genomes. For instance, the same superfamilies of gypsy are present or absent in both genomes and the “novel” lineage also has representative sequences of both. For the pao/bel group the situation is similar, only elements belonging to the Pao and Bel families are present (the Pao constituting a clade with 88% bootstrap value).

*Non-LTR*

Non-LTR retrotransposon *in-silico* screening identified that the *P. papatasi* genome possess 0.92% non-LTRs, whereas the *Lu. longipalpis* genome only contains 0.22%, which is considerably lower than that of *D. melanogaster* (approximately 6%) [3]. In sand flies, the clades which contributed most to the genome size are L2, RTE, Jockey, I and CR1. *P. papatasi* also has more variation of non-LTRs than *Lu. longipalpis*. The I clade is the only clade more abundant in *Lu. longipalpis* (0.05%) than in *P. papatasi* (0.04%). Members of clades R2 and R4, as well as L1 are not detectable in these genomes. Overall, due to the large size of a full-length non-LTR (5-9 Kb), and comparatively short length of assembled contigs the amount of non-LTRs are likely underrepresented in this study.

*Class II (DNA) TEs*

Annotated DNA transposon content in both sand fly species is lower than that of *Drosophila melanogaster* (*P. papatasi* 1.1% of the sequenced genome, *Lu. longipalpis* 0.05%, and *D. melanogaster* 2.0% [3]). It is unclear how much of these differences can be attributed to the quality of the genome assemblies, particularly for elements with such low overall abundances. Interestingly, P-elements, which are the most abundant DNA transposons in *D. melanogaster* appear to be missing from both sequenced sand fly genomes. Given that P-elements are a recent invasion in *D. melanogaster*, but have been found in other Diptera, it is not clear whether these sand flies have some defense against P-elements, or the lack of invasion is due to random chance. However, the lack of P-elements in sand flies opens up their possible use for future control strategies. Other DNA transposons, such as the Tc1/mariner elements are found both in *Drosophila* and the sand fly genomes. Information about each Transposable Element's distribution can be found in Table 1.

*Miniature Inverted-repeat Transposable Elements (MITEs)*

MITEs are short, non-autonomous and do not code for any protein. They have the terminal inverted repeats (TIRs) and target site duplication (TSD) on each flank. These features are the signature used for *in silico* searches of MITEs. Biologically, ITRs are used for the excision of the MITEs from the genome by the corresponding autonomous element's active transposase, and insertion in a new location. There are 39 and seven annotated MITEs *P. papatasi* and *Lu. longipalpis*, respectively. Annotated MITEs also appear to occupy a significantly higher percentage of the genome in *P. papatasi* than *Lu. longipalpis*. MITEs that are flanked by TA target site duplications appear to be the most abundant and occupies >2% of the *P. papatasi*. It is not clear how these comparisons may be influenced by the quality of genome assembly.

<u>Immunity</u>

In spite of the potential importance of the sand fly immune system for influencing *Leishmania* development and survival in the sand fly gut the immune response is poorly studied. There have been a number of studies on immune peptides, although to date, this work has been restricted to defensins [4, 5]. It was shown that manipulation of the immune system could lead to inhibition of

*Leishmania* development in the gut. Gene depletion via RNAi of negative regulator of IMD pathway caspar [6] led to a reduction in *Leishmania* population in the gut of *Lu. longipalpis*.

*Toll Signaling*

The *Drosophila* Toll signaling pathway is essential in the defense response against Gram-positive bacteria and fungi [7]. Sensing of these pathogens is achieved at the cell surface by recognition of conserved pathogen associated molecules in Gram positive bacteria and fungi. Two pattern recognition receptor (PRR) gene families play a role in this, namely, Peptidoglycan Recognition Proteins (PGRPs) and Glucan Binding Proteins (GNBPs). Recognition results in the activation of proteolytic cascades followed by activation of the Toll signaling pathway. Here we focused on the core signaling molecules in the Toll pathway and on the upstream pathogen recognition receptors including PGRPs and GNBPs.

In both sand fly species' genomes, Toll signaling pathway is highly conserved as homologues of all the core genes can be manually annotated confidently (Table S5). For the upstream pattern recognition receptors, PGRPs, *Lu. longipalpis* has four homologues and *P. papatasi* has two homologues. All the sand fly homologues share a conserved peptidoglycan binding domain as in the *Drosophila* PGRPs. In *Lu. longipalpis*, two homologues LLOJ002444 and LLOJ005642 have the conserved residues for the enzymatic activity detected in *Drosophila PGRP-SA* and *–SD* [8]. These two homologues also show higher identity with *PGRP-SA* compared to *PGRP-SD*. In *P. papatasi*, the residues essential for the PGRP-SA and –SD catalytic function is conserved in PPAI010204 only but not in PPAI007689. PPAI010204 also shares higher homology with *PGRP-SA*.

For GNBPs, *Lu. longipalpis* has one GNBP homologue with higher homologue to *Drosophila* GNBP3 and *P. papatasi* has four GNBP homologues. Out of the four GNBP homologues in *P. papatasi*, two are highly likely to be paralogues. In *P. papatasi*, PPAI000880 shows the highest homology with *Drosophila GNBP1*, while PPAI010440 exhibits highest homology with *Drosophila GNBP3*. Both *Lu. longipalpis* and *P. papatasi* have five Toll homologues. All the sand fly Toll receptors have the TIR domain present.

Only one homologue of each of the core signaling molecule downstream of the PRRs, *MyD88*, *tube* and *Pelle*, can be found in both *Lu. longipalpis* and *P. papatasi*, with a death domain present in all homologues.

Two homologues of Cactus are present in the *Lu. longipalpis* genome with *P. papatasi* only having one. Similarly, *Lu. longipalpis* has two *Dorsal/Dif* paralogues and *P. papatasi* has one homologue. The characteristic Rel domain of NF-κB protein family is also found in the sand fly homologues. For the NF-κB inhibitor *Cactus*, the Ankyrin repeat is present.

*IMD/ROS genes*

The Insect Immune Deficiency Pathway (IMD) has an essential role in insect defense against gram-negative bacteria in *Drosophila*. [9-11]. The activation of the IMD pathway is based upon recognition of molecules derived from the bacterial wall through specific receptors (termed peptidoglycan recognition proteins - PGRPs) located on the surface of immunocompetent cells. After receptor activation, a signaling cascade is triggered in the cell cytoplasm culminating with the activation of NF-κB transcription factors such as relish, resulting in expression of antimicrobial peptides. The IMD pathway seems to be highly conserved among dipterans, including *Drosophila*, *Aedes*, *Anopheles* and both sand fly species analysed in this study (Table S6). Homologues of several genes performing different roles in this pathway were identified for *Lu. longipalpis* and *P. papatasi* (Table S6). Negative regulators Caspar, Caudal and Pirk were found for both sand fly species and Caspar was identified for *P. papatasi* for the first time. Interestingly, Caspar knockdown down-regulates *Leishmania* growth within the gut of *Lu. longipalpis* infected with *L. mexicana* and *L. infantum* [6]. Core signalling genes such as IMD, IRD5, IAP2, Effete, Bendless, CYLD; the transcription factor Relish and the Relish activator Dredd are present for both species. TAB1 that forms a complex with TAK1 during the IMD cascade was only found in *Lu. longipalpis*. The IMD pathway is also connected with other immune-related pathways such as the one responsible for the formation of reactive oxygen species [12]. The production of reactive oxygen species (ROS) such as hydrogen peroxide ($H_2O_2$) and hypochlorous acid occur through the activation of the enzyme Dual oxidase (DUO ) [12, 13]. Calcium ions activate DUOX molecules to produce $H_2O_2$. In the presence of chloride ions, DUOX transforms hydrogen peroxide in hypochlorite which is extremely active against microorganisms. The ROS pathway is a conserved mechanism to control the insect gut flora and protect against pathogenic microorganisms. DUOX is present in the genome of *Lu. longipalpis* and *P. papatasi*, DUOX is upregulated through p38-mediated activation of ATF2. p38 is also present in both sand fly genomes (Table S6). Down-regulation of this pathway involves sequential induction of calcineurin B (CanB) and MKP3, inhibiting p38 activation and reducing *DUOX* transcription [14].

##### *Galactose-binding proteins*

Galactose-binding proteins (galectins) are a diverse family of proteins playing roles in development and immunity [15]. Comparing the sand flies' galectin protein sequences with other Diptera, clusters encompassing protein encoded by shared as well as independent orthologs were noticed (Fig. S3; Table S8). For instance, the *Le. major* receptor in the *P. papatasi* midgut PpGalec [16] (here named *PpGalecA*) shares similarities only with its *Lu. longipalpis* ortholog-encoded protein, though the latter bears an extra galactose-binding domain (Fig. S3; Table S8). Similarly, other sand fly specific galectins were identified, such as the *Lu. longipalpis LuloGalec* [17] (here named *LuloGalecR*), its putative ortholog *LuloGalecR PpGalecR, and LuloGalecII*. On the other hand, the sand fly *GalecB, GalecC, GalecD*, and *Galec_peroxin23* galectins exhibited orthologs in mosquitoes and/or flies (Fig. S3; Table S8). Even within a shared protein cluster, a recent gene duplication event was noticed by the presence of two PpGalecB genes in different scaffolds of the *P. papatasi* genome (Fig. S3; Table S8). Such galectin diversity may be an adaptation of the sand fly immune system to cope with specific symbiont and pathogenic microorganisms.

##### *TGF-beta*

Transforming growth factor beta (TGF-beta) belongs to a family of multifunctional cytokines found in organisms that go from arthropods and mammals, which regulates functions as diverse as cell differentiation and growth, adhesion, migration, and immune responses [18]. Nevertheless, the TGF-beta pathway components are quite conserved across evolution [19]. TGF-beta is synthesized as a precursor molecule containing a hydrophobic signal peptide, a poorly conserved N-terminal pro-peptide and a highly conserved and active C-terminal domain. The cytokine is secreted as a mature homo or heterodimer [18]. This cytokine superfamily comprises more than forty members, grouped into the following subfamilies according to structure or function: TGF-beta stritu sensu, BMPs (“bone morphogenic proteins”), MIF (“Mullerian inhibitory factor”) and activin/inhibin [20]. Activin/inibin is a polymeric peptide involved in many processes in *Drosophila* as growth, development and neural functions [21]. The TGF-beta family is highly conserved among arthropods and is directly involved in immunity. In the malaria vector *An. stephensi*, a TGF-beta homolog named (As60A) was implicated in the insect immune response to *Plasmodium* [22]. It was also determined that the mammalian TGF-beta 1 ingested in the blood regulates nitric oxide production, modulating the *An. stephensi* immune response against the parasite [23]. The up-regulation of a TGF-beta gene (activin/inhibin subfamily) has been observed in *Lu. longipalpis* upon infection with *Leishmania iinfantum chagasi* [24].

We identified fourteen genes in *Lu. longipalpis* genome and fourteen genes in *P. papatasi* genome related to TGF-beta or TGF-beta pathways (Table S9). We found one copy of each gene in *Lu. longipalpis* or *P. papatasi* genomes. Additionally, we searched for ortholog genes in *Glossina morsitans*, *An. gambiae* and *Ae. aegypti* genomes and we also found one copy of each gene for these species with exception of *maverick* that was not found in *Ae. aegypti* and *smad on X* that was not found in *An. gambiae*. Among these genes we found two molecules from the TGF-beta superfamily that contain TGF-beta conserved domains, two TGF-beta transcription factors that contain a SMAD (or MAD, for mothers against decapentaplegic) domains, four TGF-beta receptors that contain TGF-beta receptor conserved domains, 1 Dpp (or BMP) ligand that contains a metallopeptidase, a CUB (for complement C1r/C1s, Uegf, Bmp1) and a EGF (for epidermal growth factor) domain, 2 Dpp negative regulators that contain SMAD domains, another negative regulator that contains a WD-40 (for motifs that have approximately forty amino acids often terminating in a Trp-Asp dipeptide) and a F-box (for a motif firstly described in cyclin-F) domain, 1 Dpp inhibitor that contains a magnesium transporter domain, and a Dpp antagonist that contains a CHDN (for chordin) domain and a VWF (for von Willebrand factor) domain.

*MAP Kinases*

Mitogen-activated protein kinases (MAPK) are key components of a series of vital signal transduction pathways that regulate processes such as growth, metabolism, apoptosis, and innate immune responses. The MAPK pathways are a cascade of four regulatory serine-threonine protein kinases which results in the activation of a MAPK that that can regulate effector proteins and transcription factor. MAPKs are phosphorylated and activated by the MAP2Ks (MAPK kinases), also known as MEKs. The MAP2Ks are phosphorylated and activated by MAP3Ks (MAPK kinase kinases), also known as MEKKs [25]. The whole pathway activation starts by the phosphorylation of a MAP4K which is controlled by different cell stimulus. The main MAPKs pathways are the ERK pathway (extracellular signal-regulated kinase), JNK/SAPK pathway (c-Jun NH2-terminal kinase/stress-activated protein kinase) and p38 pathway [26].

In mosquitoes infected with the malaria parasite there is an increase in the production of nitric oxide, which inhibits the proliferation of *Plasmodium* inside the insect. This reaction is caused by the activation of a MAPK signaling pathway [27]. It is known that the activation of the MAPK pathway leading to the NOS production in mosquitoes infected with *Plasmodium* is regulated by the human TGF-beta ingested with the blood [28].

Sixteen different MAPK gene loci were identified in the genome of *Lu. longipalpis* and 15 in the *P. papatasi* genome (Table S10). A *Lu. longipalpis* homologue of *MAP2K3* (ABR28347.1) was found in the transcriptome, but its codifying gene was not in the genome. The number of *Lu. longipalpis* MAPKs is equal to *An. gambiae* [29]. The difference between the two species is that the mosquito has two JNK paralogues and one ERK copy while the sand fly has one JNK and two ERKs. Sand flies also have NLK (Nemo-like kinase) but we only focused on the main MAPKs pathways.

*Other immune-related genes*

Phenoloxidases are ubiquitous type 3 copper-containing enzymes also involved in insect cellular and humoral immune defences. The activated enzyme oxidises phenolic compounds to produce melanin and encapsulate invading pathogens [30]. Due to its high importance and high degree of conservancy among organisms, two prophenoloxidase homologs were found in *Lu. longipalpis* and *P. papatasi* genomes (Table S11).

A TEP-1-like molecule was also found in the genome of *Lu. longipalpis* and *P. papatasi* (Table S5). Tep-1 is a complement-like peptide previously described in *An. gambiae* that interacts and is cleaved by the LRIM-APLC1 complex, targeting *Plasmodium* for destruction [31]. The presence of orthologs of a complement-like system in the sand fly haemolymph raises the possibility that such machinery might be part of sand fly innate immune defences against pathogens.

Eicosanoids act in several lines of defence in insects, inducing phagocytosis, aggregation and haemocyte migration towards an invading pathogen [32]. Cyclooxygenases (COX) are not found in insects, being involved in one of the pathways leading to eicosanoid biosynthesis. Instead, a COX-like molecule was identified in the *Drosophila* genome, possibly exerting a similar role [33]. A COX-like ortholog is present in both sand fly genomes (Table S5), possibly being involved in pathogen killing through eicosanoid production.

Salivary Protein Genes

Proteomics and transcriptomics have identified the most abundant sand fly salivary proteins from *P. papatasi* and *Lu. longipalpis* sand flies. There are 49 *P. papatasi* and 35 *Lu. longipalpis* putative salivary genes deposited at the NCBI [34]. We mapped these sequences to the sand fly assemblies (Table S12).

During blood feeding, female sand flies inoculate saliva into the host dermis where salivary components disrupt hemostasis and the immune system of the host to maintain blood flow. Salivary proteins counteract the coagulation system by blocking clotting and inducing local vasodilation. Additionally, distinct sand fly salivary proteins have immunomodulatory properties reducing inflammation and modulating cellular recruitment to the bite site [34]. Counter-intuitively, repeated exposure to sand fly bites or inoculation of distinct salivary proteins induces an adaptive immune response that can be measured as a delayed-type hypersensitivity response and a specific anti-sand fly saliva antibody response. The antagonistic interaction of some of the sand fly salivary proteins with the host coagulation and immune systems exerts an evolutionary pressure. We can appreciate this phenomenon in the frequency of gene duplication events and the expansion of several sand fly salivary families resulting in the presence of several homologues for some genes. These patterns are highlighted by the presence of motifs assigning conserved families of sand fly salivary proteins that otherwise share a low degree of identity between members when comparing *P. papatasi* and *Lu. longipalpis*. This observation suggests a fast evolution pace for salivary genes for example when compared to digestive or olfactory molecules. Below we summarize the function and features of the salivary proteins that were found in the search of the *P. papatasi* and *Lu. longipalpis* databases.

*Yellow related proteins*

Yellow related proteins are abundant molecules in both species. Despite their size, around 40-kDa, they bind and trap biogenic amines such as serotonin, histamine and others in a central pocket. Removal of biogenic amines from the environment inhibits the activation of the coagulation system or reduces the availability of inflammatory mediators. LJM111, a yellow protein from *Lu. longipalpis*, has anti-inflammatory properties inhibiting IL-17, TNF-α, IFN-γ and neutrophil migration in an ovalbumin-induced neutrophil migration mouse model [35]. LJM11 and LJM17, other yellow proteins from *Lu. longipalpis* are immunogenic and immunization with LJM11 protects mice from *Leishmania* infection [36]. Conversely, immunization with PpSP44, the most abundant yellow protein from *P. papatasi*, led to exacerbation of parasite infection [37]. No function has been assigned to yellow proteins from the *P. papatasi* sand fly.

*D7 related Proteins*

PPTSP30 and LJL13 D7 belong to the D7 related proteins and have been found in New and Old World species of sand flies. They are classified as part of the odorant binding family of proteins [38]. They are also present in several other blood feeding insects such as mosquitoes and black flies. The function of these proteins remains unknown in sand flies, but it is possible that they also work by binding biogenic amines and eicosanoids similarly to mosquito D7 proteins [39].

*Silk related/collagen binding/32kDa family of proteins*

These salivary proteins are commonly found in saliva of most studied sand fly species. PpSP32 from *P. papatasi* has been characterized as the main marker of exposure to sand fly bites in an endemic area of cutaneous leishmaniasis in Tunisia [40]. There have been no functional studies undertaken on the *Lu. longipalpis* salivary protein LJL13.

*Antigen 5 family of proteins*

PPTSP29 and LJL34 belong to the Antigen 5 family of proteins. These proteins are commonly present in many bloodsucking insects and are associated to the CAP family of proteins composed of cysteine-rich secretory proteins, Antigen 5, and pathogenesis-related 1 proteins [41].

*Apyrases*

Apyrases are enzymes that hydrolyze ATP and ADP preventing platelet aggregation and facilitating blood feeding. Apyrases are present in saliva of all sand fly species studied; LJL23 and PPTSP36 represent the Apyrases for *Lu. longipalpis* and *P. papatasi* sand flies, respectively [41].

*Maxadilan*

One of the best-characterized molecules in *Lu. longipalpis* saliva, maxadilan, it is a powerful vasodilator and an anti-inflammatory peptide of 6.8 kDa [42]. Vaccination with DNA encoding maxadilan protected mice against *Leishmania* infection [43].

*Anti-coagulant*

LJL143 from *Lu. longipalpis*, named Lufaxin, is an inhibitor of Factor Xa, disrupting both the intrinsic and extrinsic coagulation cascades. Moreover, by its interaction with the PAR receptors it can modulate inflammation [44].

*Endonucleases*

LJL138 from *Lu. longipalpis*, named Lundep, is an endonuclease that cleaves DNA and, at a lower efficiency, RNA. Lundep has a role in blood feeding through inhibition of the intrinsic pathway of coagulation. Additionally, Lundep cleaves neutrophil extracellular traps enhancing the survival of *Leishmania* parasites upon deposition into the host tissue [45].

*10-kDa family of proteins*

LJS192, LJS169 and LJM19 are proteins of unknown function that are abundant in the saliva of *Lu. longipalpis*. Proteins of similar molecular weight but with low identity have been identified from the *Le. intermedia* salivary transcriptome [46]. Vaccination with LJM19 protects hamsters from cutaneous [47] and visceral [48] leishmaniasis.

*C-type lectin*

LJL91, LJL15, LJL18, LJM10 and LJS142 from *Lu. longipalpis* are members of the C-type lectin family of proteins that seems to be unique to New World sand flies. No function has been assigned to this family of proteins.

PpSP56.6

Previously identified by transcriptomic analysis, the function of PpSP56.6 from *P. papatasi* is unknown. There is one other member of this family identified from *P. sergenti* saliva [49].

Amylase PPTAMY, a putative alpha-amylase from *P. papatasi*, hydrolyzes dietary starch to maltose, which is then cleaved to glucose by alpha glucosidase. Starch if one of the major components in the natural diet of *P. papatasi*.

<u>Digestion</u>

Digestive processes in Phlebotomine sand flies are of key interest in terms of the relevance to the gut dwelling *Leishmania* parasite. Successful development and emergence from the digesting blood meal is an essential feature for the development of a transmissible parasite population in the gut. A number of studies suggest that expression of digestive proteases may be influenced by the presence of *Leishmania* [50]. Inhibition of trypsin like protease via gene depletion enhanced *Leishmania* survival in *Lu. longipalpis* [51].

*Peptidases*

Peptidases (E.C. 3.4) are enzymes responsible for the hydrolysis of peptide bonds. They are classified according to the site of cleavage in the substrate, in exopeptidases (terminal bonds) or endopeptidases (internal bonds). Besides that, peptidases are classified by their catalytic mechanism in aspartic, cysteine, glutamic, metallo, asparagine, serine, threonine and mixed type peptidases. A more recent approach is the classification of peptidases in families based on the similarity of amino acid sequences, which reflect the overall structure and the catalytic mechanism of homologous enzymes [52].

We conducted a systematic search in the genomes of *Lu. longipalpis* and *P. papatasi* for genes belonging to all known peptidase families, using HMMER/blast and the results are presented in Table S13. Both genomes contain at least 376 protease genes belonging to several families (64 *Lu. longipalpis* and 62 in *P. papatasi*), with a predominance of serine proteinases (48.4 and 46.3% of total protease genes, respectively). This distribution is similar to the observed in other dipterans known genomes, where serine proteases account for 45-64 % of all protease genes. This is consistent with the involvement of serine proteases in larval and adult sand fly digestion. Because larval and adult initial protein digestion rely mainly on the same type of proteinases from family S1, tissue specific gene expression studies are necessary to elucidate if some gene expansion in this family occurred during sand fly adaptation to hematophagy.

Interestingly, two proteinase families were found to be unique to sand flies among dipterans. These are families C95 and M87. Families C95 and M87 are proteinases which are present in other animals [53, 54] and its occurrence in *Acyrtosiphon pisum* suggests that these are ancient sequences which might be present in the hexapodan ancestor. However, its occurrence is sparse throughout the animal kingdom, and more detailed studies are necessary to elucidate their evolutionary origin in sandflies. Notably, family M87 correspond to chloride channel associated proteins, and as the chloride channel is the target of several insecticides, its occurrence in sand flies can be the basis for insecticide selectivity between these vectors and other organisms.

Nevertheless, the number of protease genes in the sand fly genomes seem to be lower than the observed in other dipteran species. This could be a result from under representation arising from assembly or search strategy issues, or just a reflection of different protease gene expansions in culicidae and brachycera. These expansions might have occurred mainly in family S1, which harbor the majority of serine proteinase sequences. However, the number of serine proteinase genes found in sand fly genomes (177 and 174 for *Lu. longipalpis* and *P. papatasi*) is similar to the observed in other insect genomes (like *Acyrthosiphon pisum*, 197 genes, or *Apis mellifera*, 108 genes).

*Carbohydrases*

Carbohydrases, also known as glycoside hydrolases, are essential enzymes involved in sugar metabolism. They hydrolyse glycosidic bonds and are mainly classified by the substrate and glycosidic bond they recognize. Recently, they were categorized by amino acid sequence similarities in families (Glycoside Hydrolase Families, or GHFs), which reflect the overall enzyme structure and catalytic mechanism [55]. In sand fly genomes, the number of GH genes we found using a general approach (HMMER/Blast) is quite low (±30) when compared with other insect genomes (~100). This is probably a result of gene fragmentation and masking of low-quality sequences. Because of that, we focused in two GH families of special interest, GHF13 and GHF18.

*Glycoside Hydrolase Family 13 – Starch, Glycogen and Sucrose metabolism*

In general, members of GHF13 are enzymes active on alpha-glycosidic bonds which are typical of starch or glycogen. Most members of GHF13 are typical amylases and alpha-glucosidases, but GHF13 also contains enzymes with pullulanase, cyclomaltodextrinase or transferase activities, among others. Both *Lu. longipalpis* and *P. papatasi* genomes contain at least 15 GHF13 genes, summarized in Table S14. These numbers were updated in a more recent and depth analysis of GH13 proteins in *Lu. longipalpis* that was published elsewhere [56], with an account of at least 21 proteins. These high copy numbers reflect not only the multi enzymatic nature of this glycosidase family, but also the importance of these enzymes in larval and adult metabolism. These enzymes are involved mainly in starch/glycogen digestion in larvae and sucrose digestion in adults of both sexes. Accordingly, amylase and alpha-glucosidase were already detected and partly characterized in *Lu. longipalpis* [57] and some *Phlebotomus* species [58, 59] .

Sucrose digestion in insects is routinely performed by alpha-glucosidases (E.C. 3.2.1.20), as reports for beta-fructosidase (E.C. 3.2.1.26) are very scarce [60]. As expected, sand fly genomes do not contain genes belonging to GHFs 32, 68 and 100, which are the gene families for which this activity was already reported [55]. Comparison with other insect genomes suggest the presence of genes orthologous to sand fly GHF13 genes in other dipterans genomes, as described in Table S14. In many cases, several sand fly sequences are more similar to the same ortholog in *D. melanogaster*, *An. gambiae*, *Ae. aegypti* or *Culex quinquefasciatus*, which suggest that GHF13 diversification might have happened after the divergence of dipteran families.

The initially discovered number of GHF13 genes in sand fly genomes (15) is similar to the observed in *D. melanogaster* (16) and *An. gambiae* (17), higher than the number in other vectors as *Rhodinus prolixus* (6) and *G. morsitans* (8), but lower than the numbers found in *Ae. aegypti* (27) and *C. quinquefasciatus* (33). This expansion in GH13 gene family in some insect genomes could be related to the fact that alpha glucosidase is a major membrane protein in the several insect midgut epithelia (especially in the order Diptera), being the target receptor of several bacterial endotoxins [61]. This might be the motor of evolutionary co-adaptations between insects and bacterial pathogens, which could involve gene duplications, mutations and presence of isoforms conferring resistance to the insect against entomopathogenic microorganisms. Interestingly, the infection of *Lu. longipalpis* adult females with *Le. mexicana* changes the expression of alpha-glucosidase genes in the gut, but it is still not clear if this is a direct mechanism involving regulation by the pathogen, or of it is a indirect consequence of the general effects of the parasite in the digestion of sugar and blood [56].

*Chitinase and Chitinase-like proteins*

Insect glycoside hydrolases from family 18 (GH18) comprise chitinases and chitinase-like proteins. Insect genomes contain several copies of GH18 genes, as a reflection of the pivotal role of chitin structures in insect anatomy, development and physiology. Chitinases and chitinase-like proteins are categorized into eight groups based on their catalytic activity, domain structure (presence of catalytic domains, linker regions and chitin-binding domains) and amino acid sequence similarities. These groups have distinct physiological roles which were assigned using expression pattern and RNAi silencing studies in *D. melanogaster*, *Tribolium castaneum*, *An. gambiae* and *Ae. aegypti* [62, 63]. They are involved, for example, in molting, digestion, peritrophic matrix turnover, developmental morphogenesis and cell signaling. In the sand fly genomes we have found 11 (*Lu. longipalpis*) and 12 (*P. papatasi*) GH18 genes, representing all known chitinase groups and including 1 Imaginal Disc Growth Factors for each specie. Original VectorBase [64-66] gene sequences were completed and corrected using de novo assembly of transcriptome data. Characteristics of curated chitinase gene sequences are summarized in Table S15. In the case of *Lu. longipalpis* chitinase genes, ten are present in the original assembly and have VectorBase Accession numbers, and 1 were found only in de novo assembly of RNA-Seq data. GH18 gene copy numbers in sand fly genomes are more similar to the observed in *Drosophila* (16 genes, with 6 IDGF copies) than the genomes of *An. gambiae* (20 genes, 2 IDGFs), *Ae. aegypti* (16, 2 IDGFs) or *T. castaneum* (22, 2 IDGFs). GH18 proteins in sand fly genomes contain a variable number of catalytic domains (1-4) and chitin binding domains (0-4), presence or absence of signal peptide or transmembrane regions and highly glycosylated linker regions. The conservation observed in the chitinase gene number and gene structure among the dipteran genomes is probably related to the conservation of chitin role and metabolism in the insect orders. As observed in other insects, the multigenic nature of GH18 family points their members as interesting models for gene expression and physiological studies as well as targets for insecticide molecules.

*N-acetylhexosaminidases*

N-acetylhexosaminidases (HEXs) belong to the glycosyl-hydrolase family 20 [67], and participate in N-glycan processing (fused lobe proteins - FDLs; [68] as well as digestion of carbohydrate oligomers to monomers [69]. The HEX family member exo-splitting β-N-acetylglucosaminidases (NAGs) have been implicated in cuticular chitin turnover during insect molting as well as in peritrophic matrix degradation [69-71]. In sand flies, four orthologs belonging to n-acetylhexosaminidases groups I-IV were identified in each sand fly species (Table S16; Fig S4). Interestingly, a gene duplication event has taken place only in the sand flies NAG1 gene, giving rise to an extra NAG gene (NAG3), which is shared by both sand fly species (Table S16; Fig S4), therefore, sand flies bear the necessary machinery to recycle chitin along with an extra gene copy that may perform a more specialized function.

*Chitin deacetylases*

Chitin deacetylase proteins (CDA) belong to the carbohydrate esterase family CE4 and converts chitin into chitosan via N-deacetylation. This reaction might contribute to the binding of some proteins to chitosan in chitinous structures in the cuticle and peritrophic matrix [72]. In the red flour beetle (*T castaneum*), five CDA groups have been described, displaying specific expression in either cuticle or peritrophic matrix [73]. In sand flies, CDAs belonging to the five groups presented in insects were identified (Table S17; Fig. S5), suggesting chitin deacetylation may also be an important biological process in the sand fly's cuticle and PM.

*Peritrophin-like proteins*

Peritrophins are abundant component of insect PM and cuticle, in which they create the structural scaffold by cross-linking chitin fibrils [74]. These proteins bear similar chitin-binding domains (CBDs) to chitinases and can also display mucin-like motifs, which are heavily glycosylated [75]. In sand flies, 30 peritrophin genes were identified in *P. papatasi*, whereas 22 homologs were described in *Lu. longipalpis* (Table S18). As far as domain number and diversity, up to nine CBD domains were found in a single peritrophin (PpPer9), and multiple sand fly peritrophins were predicted to be glycosylated by exhibiting mucin-like motifs (Table S18). In a comparative analysis between sand flies and *T. castaneum* [76] CBD domains, multiple sand fly and RFB domains belonging to the CPAP subgroup (cuticular proteins analogous to peritrophins) clustered together (Fig. S6). On the other hand, only one sand fly CBD domain clustered together with the *T. castaneum* CBD domains of the PMP subgroup (peritrophic matrix proteins) associated with PM scaffolding (Fig. S6). These differences highlight the functional specialization of peritrophin-like proteins playing a role in peritrophic matrix formation as well as a more conserved role for the peritrophins associated with the insect exoskeleton.

Aquaporins

Aquaporins (AQPs) are required for the transportation of water and other small solutes across cell membranes. We have identified six aquaporin genes from both species of sand flies (Table S19; Fig. S7). This is similar to the number present in mosquitoes (N = 6), but two and four less than *Drosophila* and *Glossina*, respectively [77]. Members of each AQP group previously identified from insects are present in the sand fly genomes, which includes those involved in the water and the recently assigned to be involved in glycerol transport [78].

Circadian Rhythm

As far as we know, all organisms present an endogenous mechanism that allows the responsiveness to different environmental stimuli such as light and temperature, the biological clocks. Among them, the circadian clocks (which last about 24 hours), regulate the rhythms most closely related to daily behaviours such as locomotion, oviposition and larval eclosion.

In the model species, *D. melanogaster*, the molecular basis of the circadian clock has been well studied for decades. Genes that are known to influence several behaviours or encode for receptors involved in the detection of heat, water or mechanical stimuli are controlled by clock genes and have been characterized, mainly in *D. melanogaster* and *Ap. mellifera* [79-85].

The major components of the circadian clock form three regulator feedback loops. In the main loop, the *Clock (Clk)* and *cycle (cyc)* genes encode two activators, CLK and CYC, that form a heterodimer and bind in regulatory sequences called E-boxes (CACGTG) in the promoters of *period (per)* and *timeless* (*tim*) genes, activating their transcription. After a series of post-translational changes, PER and TIM proteins form a dimer, enter in the nucleus and inhibit CLK/CYC function, in a cyclic manner. In the second loop, the genes *vrille* (*vri*) and *PAR domain protein 1* (*Pdp1*), which are also are cyclically activated by the CLK/CYC heterodimer, encode a repressor (VRI) and an activator (PDP1), respectively, which in turn compete for the same site in the *Clk* promoter, regulating its transcription. The last described loop involves the regulation of CLK/CYC targets by *clockwork-orange* (*cwo*) gene [82].

Insects that are vectors of many pathogens, such as virus, protozoans and worms, for example, have a precise influence of the circadian clock to a successful vectorial capacity [86]. Although hematophagy and host seeking are controlled by the clock, the molecular regulation is poorly understood in sand flies. In *Lu. longipalpis*, the expression pattern of the core loop was described and present striking differences when compared to the *Drosophila* pattern, mainly in *cyc* expression. While *Drosophila* presents a constitutive *cyc* expression, *Lu. longipalpis* has a cyclic expression of this gene, with great amplitude [87]. Until now, no description of the molecular clock of *P. papatasi* has been available.

All core clocks genes were found in *Lu. longipalpis* (Table S20) and *P. papatasi* (Table S21) genomes (*period, timeless, cycle, Clock*, photolyases, *clockwork orange*, *vrille*, *Pdp1*), as well as post-translational modifiers (kinases and phosphatases). Other output circadian genes were successfully confirmed in both genomes (*timeout*, *single minded*, *slimb, tango, nemo, cacophony, paralytic, narrow abdomen, slowpoke, nocte, ataxin-2, circadian trip* and *takeout-like*). The amount of takeout/Juvenile Hormone Binding Protein copies found in *Lu. longipalpis* and *P. papatasi* was far below than it is found in other dipterans (5-6 in sand flies against 24-30 copies in the other genomes). Surprisingly, it was found two copies of the *casein-kinase-2* gene in *Lu. longipalpis* genome, whereas in *P. papatasi* and other dipterans there is only one copy of this gene. Another difference between the sand fly genomes is that serine/threonine phosphatase 2-beta (Pp2-b) is not found in *Lu. longipalpis*, while Pp1-beta and Pp4 are absent in *P. papatasi*. The most striking finding is the absence of a cryptochrome-1 in *Lu. longipalpis* genome and transcriptome, and its presence in the species *P. papatasi* (Fig. S8).

Nine and ten members of the transient receptor potential (TRP) cation channel family have been found in *Lu. longipalpis* and *P. papatasi* genomes, respectively. The presence of six predicted transmembrane domains and the PF00520 (ion transport protein family) and PF12796 (ankyrin repeats) domains, characteristic features of this protein family, was confirmed in the sequences of all candidates. *D. melanogaster* orthologous sequences of *pyrexia*, *TRPA5* and *TRPP* seem to be absent in both sand fly genomes. No differences in TRPM, TRPML, TRPC and TRPA subfamilies were detected between both sand fly species, except for water which gene (belonging to TRPA subfamily) with five and three paralogs in *Lu. longipalpis* and *P. papatasi*, respectively. Interestingly, NompC (belonging to TRPN subfamily) has been identified in *Lu. longipalpis* genome; however, it is absent in *P. papatasi*. Regarding TRPV subfamily, *inactive* and *nanchung* ortholog were detected only in *P. papatasi*. The TRP phylogenetic tree showed a separation of the different TRP subfamilies [88] and high bootstrap values supporting the different clades (Fig. S9).

In the case of pickpocket (PPK) family, fourteen and thirteen members have been identified in *Lu. longipalpis* and *P. papatasi* genomes, respectively. All candidates presented the two predicted transmembrane domains and the PF00858 domain corresponding to amiloride-sensitive sodium channel superfamily. The phylogenetic tree demonstrated a division of the six different PPK subfamilies, as proposed by Zelle et al. [89] (Fig. S10). None of the genes represented in *D. melanogaster* PPK subfamily III was present in the sand fly genomes. The orthologue of *ppk28*, which is related to water perception in *D. melanogaster* [90, 91], has been identified in both sand fly species (Fig. S10). Other *D. melanogaster ppk* orthologues (*ppk16*, *ppk31* and *ppk3*) were identified in *P. papatasi* and *Lu. longipalpis* (Fig. S10). In the case of sand fly *ppk100-105*, it was impossible to identify clear fruit fly orthologous sequences. Interestingly, sand fly *ppk100*-103 and *ppk-like* from *Lu. longipalpis* grouped to the conserved subgroup *ppk*, *rpk* and *ppk26* (Fig. S10). It is important to mention that *Lu. longipalpis ppk101*, *ppk102* and *ppk*-like are clustered in the same region of the genome (Table S20). An identical situation was observed for *P. papatasi ppk100a* and *ppk100b* and *ppk102* and *ppk103* (Table S21).

Other behavioural genes identified include: 1) *foraging* (*for*), which has been related to different patterns of locomotor activity in *Drosophila* [85, 92], locusts [93] and honeybees [79, 94]; 2) *malvolio* (*mlv*), which has been associated to labour division in honey bee [95] and normal taste behaviour in *Drosophila* [96]; 3) *stripe* (*sr*), involved in fly orientation in *Ap. mellifera* [97]; and 4) *piezo*, which mediates noxious mechanosensory stimuli for *Drosophila* [80].

Interestingly, both sand fly species presented a number of receptors related to the detection of moist air (Llonwtwr1-5 and Ppapwtrw1-3) [98] greater than that observed in other insects [99]. Besides the Dmelppk28 orthologue, related to water reception [90], has been identified in both sand fly genomes. The presence of these receptors could be related to the susceptibility to dehydration of these insects due to its small size. In fact, a high constant humidity is required for rearing *Lu. longipalpis* and *P. papatasi* in laboratory [100]. Regarding TRP superfamily, the main difference in those members (*nanchung*, *inactive* and *NompC*) is related to hearing [101]. Functional genetics studies would be necessary to assess the meaning of these differences between both species. The number of PPKs identified in both sand fly species (13 and 14) is lower than those reported from *An. gambiae* and *D. melanogaster*, 31 and 18 respectively [89]. Both sand fly genomes encode five different *ppks*, which clustered with the conserved group *ppk1*, *ppk2* and *ppk26* of subfamily IV. A similar gene expansion has been observed in *An. gambiae*, specifically with *ppk26* and four related subunits [89], however, in case of sand flies its relation to the IV subfamily members was not well resolved in our phylogenetic tree. Besides, at least one *ppk* from one sand fly genome has been identified in all PPK subfamilies, except for subfamily III, which is not shared by all dipteran species [89]. The differences found between *Lu. longipalpis* and *P. papatasi ppk* members, e.g. Llonppk9, Llonppk13 or Ppapppk23; need further investigation. Regarding other genes identified, e.g. *foraging* or *stripe*, behavioural and functional genetic studies would be necessary to confirm the conservation of these gene functions in sand flies [102].

<u>Cytochrome P450s</u>

CYP6AK and CYP6AG have been considerably expanded in *Lu. longipalpis* and at a lesser extent in *P. papatasi*, while CYP9J/9L has been almost equally expanded in both species (Fig. S11). Further examination suggests that these expansions have been probably caused by tandem gene duplications. In particular, expansions of CYP9J/9L and CYP6AK have each formed two large gene clusters in the *Lu. longipalpis* genome, which consist of 15 consecutive CYP9J/9L-like and 10 consecutive CYP6AK-like genes, respectively. The CYP6AG expansion also formed a gene cluster that is conserved in both sand flies and it is composed of seven and five tandem genes in *Lu. longipalpis* and *P. papatasi*, respectively.

The CYP3 clan is generally associated with xenobiotic detoxification [103]. Indeed, two of the three expanded subfamilies have been previously implicated in xenobiotic metabolism and insecticide resistance in mosquito vector species. In particular, CYP9J enzymes are involved in xenobiotic detoxification and pyrethroid resistance in the dengue vector, *Ae. aegypti* [104]. *CYP6AK1* has been associated with *A. gambiae* permethrin resistance. Therefore, expansions of these subfamilies in *Lu. longipalpis* and *P. papatasi* could possibly reflect sand fly-specific adaptations to environmental challenges posed by xenobiotics.

Sand fly P450 diversity is much more limited in the other three CYP clans. The Mito clan is complete in *Lu. longipalpis* and *P. papatasi*, since all the conserved genes are present in both sand flies (Fig. S11). Importantly, the Mito clan genes implicated in ecdysteroid metabolism are present in both sand flies, even though they are all fragmented in *P. papatasi*. Namely, CYP315A1, CYP302A1, and CYP314A1 are present and cluster confidently with their *A. gambiae* orthologs (Fig. S11).

The CYP2 clan is also complete, with a sand fly ortholog in each of the major clades of this clan (Fig. S11). There is a total of 13 sand fly CYPs, three of which are full-length. These full-length genes are the orthologs of CYP303A1, CYP304B1, and CYP305A1. More specifically, *Lu. longipalpis* and *P. papatasi* have three and four CYP304B1-like genes respectively, most probably indicating species-specific duplications. The ecdysteroid metabolism genes of this clan, CYP306A1 and CYP307A2 are also present. Finally, orthologs of the mosquito CYP15B1, which is implicated in the juvenile hormone biosynthesis, is also detected in both sand fly genomes.

*Lu. longipalpis* and *P. papatasi* have contracted CYP4 clans, mostly caused by an *An. gambiae*-specific expansion of the CYP4H subfamily. Orthologs for each member of the conserved CYP4G subfamily which is involved in cuticular hydrocarbon biosynthesis, also exist in both sand fly genomes.

It is worth noting that there are some conserved P450s that appear to be duplicated in *P. papatasi*, such as CYP314A1, CYP307A2, CYP15B1 and CYP4G17. However, all these genes are known single-copy genes in insects and they are also single-copy in *Lu. longipalpis*. As a result, the apparent *P. papatasi*-specific duplication is most likely due to the fragmentation of the *P. papatasi* genome assembly.

RNA genes and MicroRNAs

RNA interference (RNAi) refers to pathways that utilize small non-coding RNAs (ncRNAs) associated with Argonaute proteins to regulate gene expression. Most animals, including insects, have at least three separate classes of small ncRNAs known as microRNAs (miRNAs), small interfering RNAs (siRNAs) and piwi-interacting RNAs (piRNAs). These classes of small RNA differ in their mechanism of biogenesis and action. Specifically, each class of small RNA is associated with different argonaute proteins and, in the case of miRNAs and siRNAs, also ribonuclease III proteins such as Dicer and Drosha and small dsRNA binding protein partners. The miRNA pathway regulates the expression endogenous gene thus being required for development and most biological responses. The siRNA pathway controls the expression of transposable elements in somatic tissues and also mediates a powerful antiviral defense. The piRNA pathway regulates the activity of transposable elements in the animal germline.

The mechanism of RNAi has been extensively characterized utilizing the fruit fly *D. melanogaster* as an animal model. Thus, we have utilized *D. melanogaster* RNAi genes as a reference to help identify and characterize core RNAi genes and small ncRNAs in *Lu. longipalpis*. Utilizing this approach, we have identified genes encoding core genes for all three RNAi pathways in the *Lutzomyia* genome: *Drosha, Pasha, loqs, Dcr-1 and AGO1* (two separate copies) for the miRNA pathway; *R2D2*, *Dcr-2* and *AGO2* for the siRNA pathway; *AGO3* and *AGO4* Piwi genes for the piRNA pathway (Table S24). We have also found orthologs for most RNAi genes in the *P. papatasi* genome although we could not find genes corresponding to R2D2 and AGO2 and only found 2 copies of piwi genes. These genes are not likely to be absent in the *P. papatasi* genome but rather just not included in the current assembly.

In addition, we have deep sequenced and analyzed small RNA libraries from female adults of *Lu. longipalpis* in order to annotated potential ncRNA genes corresponding to miRNA genes and piRNA clusters. We annotated 88 unique and 113 total miRNA precursor genes, including 58 unique genes that are conserved in other insects and 30 unique miRNA genes that seem to be *Lu. longipalpis* specific. Total number of miRNA genes is comparable to what has been observed in *Ae. aegypti* that has 101 annotated miRNA precursors. However, it is likely that we have missed a few miRNA genes as *D. melanogaster* has 256 miRNA precursor genes annotated. We also identified 29 genomic clusters that generate piRNAs in the *Lu. longipalpis* genome based on a pattern search approach first applied to *D. melanogaster*. Although piRNA clusters tend to be repeat rich regions and show low sequence conservation comparing different insects, we have clearly identified conserved characteristics in *Lutzomyia* piRNAs compared to *Drosophila* such as size and a clear ping-pong signature associated with a piRNA-specific amplification mechanism.

In summary, we have found RNAi genes and non-coding RNAs corresponding to all three major RNAi pathways in *Lu. longipalpis* and *P. papatasi*. These RNAi pathways show different degrees of conservation compared to *D. melanogaster* and other Dipteran insects, with miRNAs being the most conserved and piRNAs the most divergent.

#### Heat Shock Protein Genes

Comparison of *D. melanogaster* proteins with GO terms relating to heat shock and response to hypoxia to the hypothetical scaffolds of the two sand fly species *Lu. longipalpis* and *P. papatasi* allowed for identification of 83 and 78 orthologs, respectively, in each species (Table S25). These comparisons were accomplished by BLAST comparison of the known *D. melanogaster* heat shock and hypoxia proteins to the scaffolds and predicted genes in *Lu. longipalpis* and *P. papatasi*. Genes that were identified as possible orthologs were then compared by BLAST to the arthropod-specific BLAST database using CLC Genomics Workbench 7. Due to a high degree of similarity, 53 of the *Lu. longipalpis* proteins require no revisions while 14 require minor revisions and 16 require large-scale revisions. For *P. papatasi* gene predictions, 44 require no revisions while 13 require minor revisions and 21 require major revisions. Overall, there appears to be no interesting expansions or retractions in gene families associated with heat shock or hypoxia for either sand fly species.

#### Cuticular Protein Genes

Cuticular protein sequences containing the R&R consensus chitin-binding domain, hence belonging to the Cuticle Protein R&R (CPR) family, were identified from *Lu. longipalpis* and manually annotated using CLC Genomics Workbench 7. Relatively all of the predicted CPR family genes need substantial revision (these are marked with a Y in cuticular (Table S26). *Lu. longipalpis*, 74, and *P. papatasi*, 84, have far fewer genes encoding CPR proteins than *D. melanogaster, An. gambiae*, and *Ae. aegypti* (101, 158 and 240, respectively, [105]). An additional 43 sequences belonging to other cuticular protein families were identified for *Lu. longipalpis* and 41 additional sequences for *P. papatasi*. All of these putative cuticular proteins were defined based upon sequence similarity through BLAST to proteins that were either isolated directly from cuticle or appeared in genomic analysis of *Drosophila* as cuticle proteins. Although we note a major reduction in the number of cuticle proteins, we are hesitant to confirm this is the case due to the fact that many of the CPR gene predictions were of low quality and need revision, suggesting that a few to many genes might not have been predicted or are predicted incorrectly.

#### Hormonal Signaling

##### *Juvenile hormone signaling*

Juvenile hormone signaling gene were identified by BLAST comparison of the known *D. melanogaster* heat shock and hypoxia proteins to the scaffolds and predicted genes in *Lu. longipalpis* and *P. papatasi*. Genes that were identified as possible orthologs were then compared by BLAST to the arthropod-specific BLAST database using CLC Genomics Workbench 7. Genes identified as components of the juvenile hormone signaling pathway are highly conserved between *D. melanogaster*, *An. gambiae*, *Ae. aegypti* and the two sand fly species, *Lu. longipalpis* and *P. papatasi* (Table S27). This is fairly unsurprising given the deep evolutionary conservation for the fundamental aspects of juvenile hormone signaling [106]. The methoprene tolerant (met) gene is recognized as the best candidate juvenile hormone receptor. In Brachycera, two paralogous JH receptor candidates Met and gce are thought to transduce the juvenile hormone signaling pathway. The *gce* gene is homologous to “Met” in other Diptera, such as mosquitoes. In the two sand flies, the *gce* gene is more homologous to given gene model rather than the met gene, which is similar to that in *An. gambiae* and *Ae. aegypti*.

##### *Insulin signaling*

Comparison of *D. melanogaster* proteins with GO terms relating to insulin signaling processes to the scaffolds of the two sand fly species *Lu. longipalpis* and *P. papatasi* using BLAST methods described in the insulin signaling section allowed for identification of 31 and 30 orthologs, respectively, in each species associated with insulin signaling (Table S28). Due to a high degree of similarity, 14 of the *Lu. longipalpis* proteins require no revisions while 5 require minor revisions and 12 require large-scale revisions. 13 of the *P. papatasi* gene predictions also require no revisions while 7 require minor revisions and 10 require major revisions. While not all insulin signaling proteins were identified this does not indicate a lack of missing proteins in the genome. It is likely that further assessment of the genomes would identify the potential missing genes.

Antioxidants

Comparison of *D. melanogaster* proteins with GO terms relating to antioxidant activity and response to oxidative stress to predicted genes of the two sand fly species *Lu. longipalpis* and *P. papatasi* allowed for identification of orthologs in each species. Due to a high degree of similarity, 38 of the 65 identified *Lu. longipalpis* orthologs require no revisions while 13 require minor revisions and 14 require large-scale revisions. 31 of the 61 identified *P. papatasi* predicted models also require no revisions while 19 require minor revisions and 11 require major revisions (Table S29). This analysis revealed a likely expansion of proteins with peroxidase and peroxiredoxin functions in both sand fly species analyzed, which was also noted in mosquito species [107-109].

Vitamin Metabolism

A comparison between *D. melanogaster* and the hypothetical scaffolds of *P. papatasi* and *Lu. longipalpis* revealed high levels of similarity between the three species genes known to be involved in the in vitamin metabolic processes. In *P. papatasi* and *Lu. longipalpis,* a total of 59 and 58 genes were found, respectively, to be potentially associated with vitamin metabolism. High similarity was noted between *D. melanogaster, An. gambiae* and *Ae.aegypti* and the two sand fly species. Only a small percentage of the hypothetical transcripts required revision (marked Y in revision required column in Table S30).

Novel Viruses

The *P. papatasi* and *Lu. longipalpis* genomes are essentially similar in terms of their integrated virus complement. For *Lu. longipalpis* there appear to be traces of ancient bracoviral insertions in to the *Lutozmyia* genome, and perhaps some retroviral sequences. HF586479: Cotesia congregata bracovirus, pos. 140201-140316 is annotated as a “proviral locus”. It is found in a variety of genomes of insects *Dendroctonus ponderosae* (mountain pine beetle), and *Drosophila* but also in *Schistosoma*. EU001284: *Glyptapanteles flavicoxis* (parasitic wasp) bracovirus segment 29, pos. 11878-11995, is also found in various insect genomes. The region of assembly of the *Lutzomyia* reads is in a long non-coding 3’ stretch. AJ289710: human endogenous retrovirus (hERV) H, pos. 3187-3452. BXU82084: Bacteriophage X transposon IS2, pos. 872-1204. Phage DNA is frequently inserted into deep sequencing experiments as a quantitative control.

To further investigate the bracoviral insertions, RNAseq *.fq reads were aligned onto the bracovirus fraction alone of GenBank (1303 polydnaviridae family GenBank sequences). Like the *.sff reads, the RNASeq reads aligning to the bracovirus fraction are mostly, with the exception of the hits to EU001284.

Unlike *Lutzomyia*, there are several hits to *Wolbachia* phages, indicating probably *Wolbachia* infection of flies used for sequencing, not found in *Lutzomyia*. AB161975: W0CauB1 appears to be present in its entirety and AB036666: WO is nearly complete. HQ906663: wVitA has its entire 5'-half present and HQ906662: wVitA has patchier coverage.

BLAST of transcripts with viral reference genomes: In both *P. papatasi* and *Lu. longipalpis*, all matches of longer than 100 nucleotides can be accounted for as cellular genes. In particular, ribonucleoside diphosphate reductase is found in both viral and cellular genomes. Heat shock proteins, thymidylate kinases, signal recognition particles and ubiquitins account for the rest. BLAST of transcripts and genome contigs onto viral GenBank fraction: In *Lu. longipalpis*, 19 transcripts have BLAST hits to retroviral sequences, and 79 to bracoviral sequences. In *P. papatasi*, the corresponding figures are 18 and 63 respectively.

#### SUPPLEMENTAL FIGURES

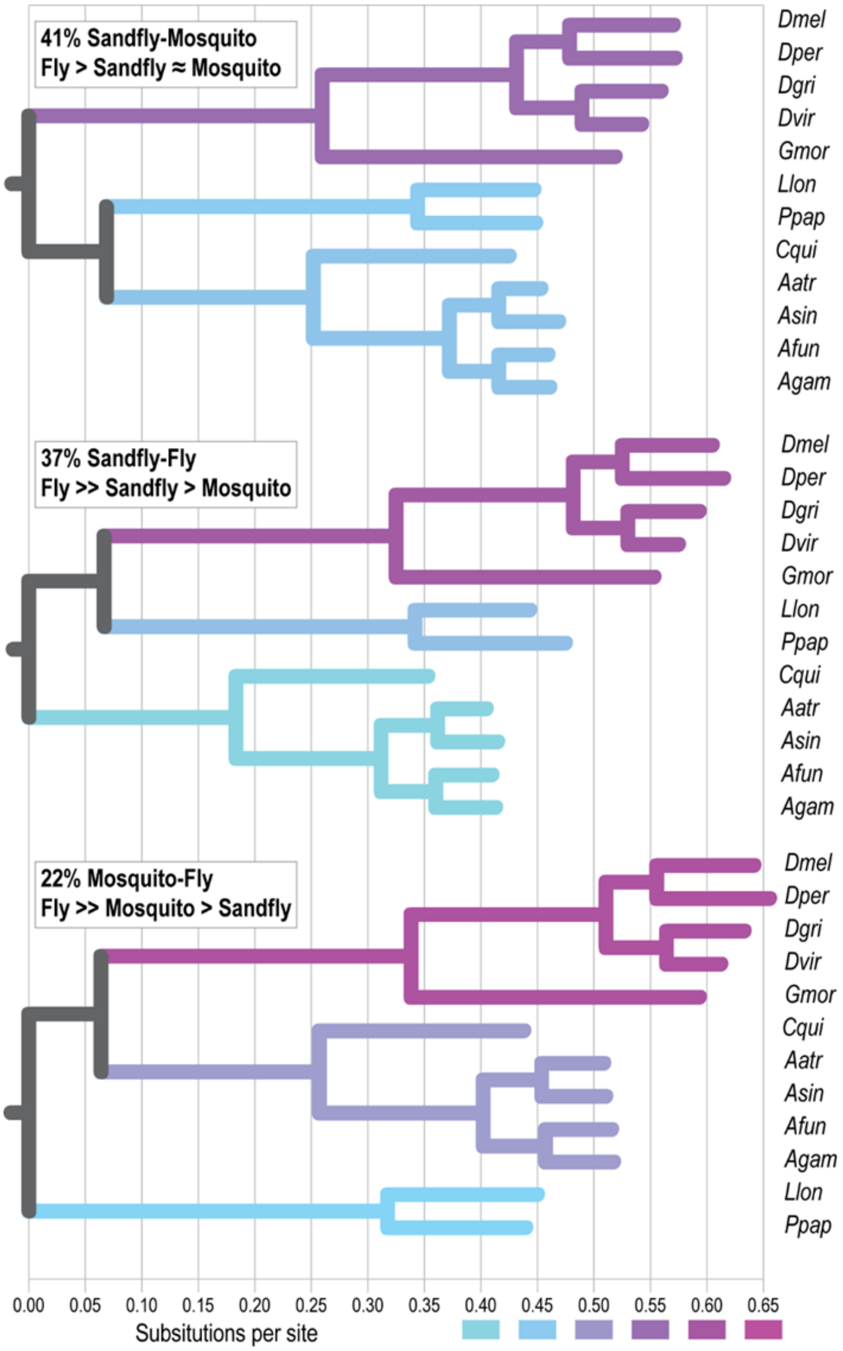

**S1 Figure. Conflicting phylogenetic signals.** Analysis of the gene phylogenies of individual orthologous groups identified three major topologies with sand fly-mosquito (41%), sand fly-fly (37%), or mosquito-fly (22%) sister clades. Comparisons of average branch lengths for each topology suggest that, although substitution rates in flies are always higher, orthologs that support the sand fly-mosquito topology show the lowest substitution rates in flies and the smallest differences in substitution rates among the fly, sand fly, and mosquito clades. In contrast, the sand fly-fly and mosquito-fly topologies show much higher substitution rates in flies and much greater differences in substitution rates among the three clades.

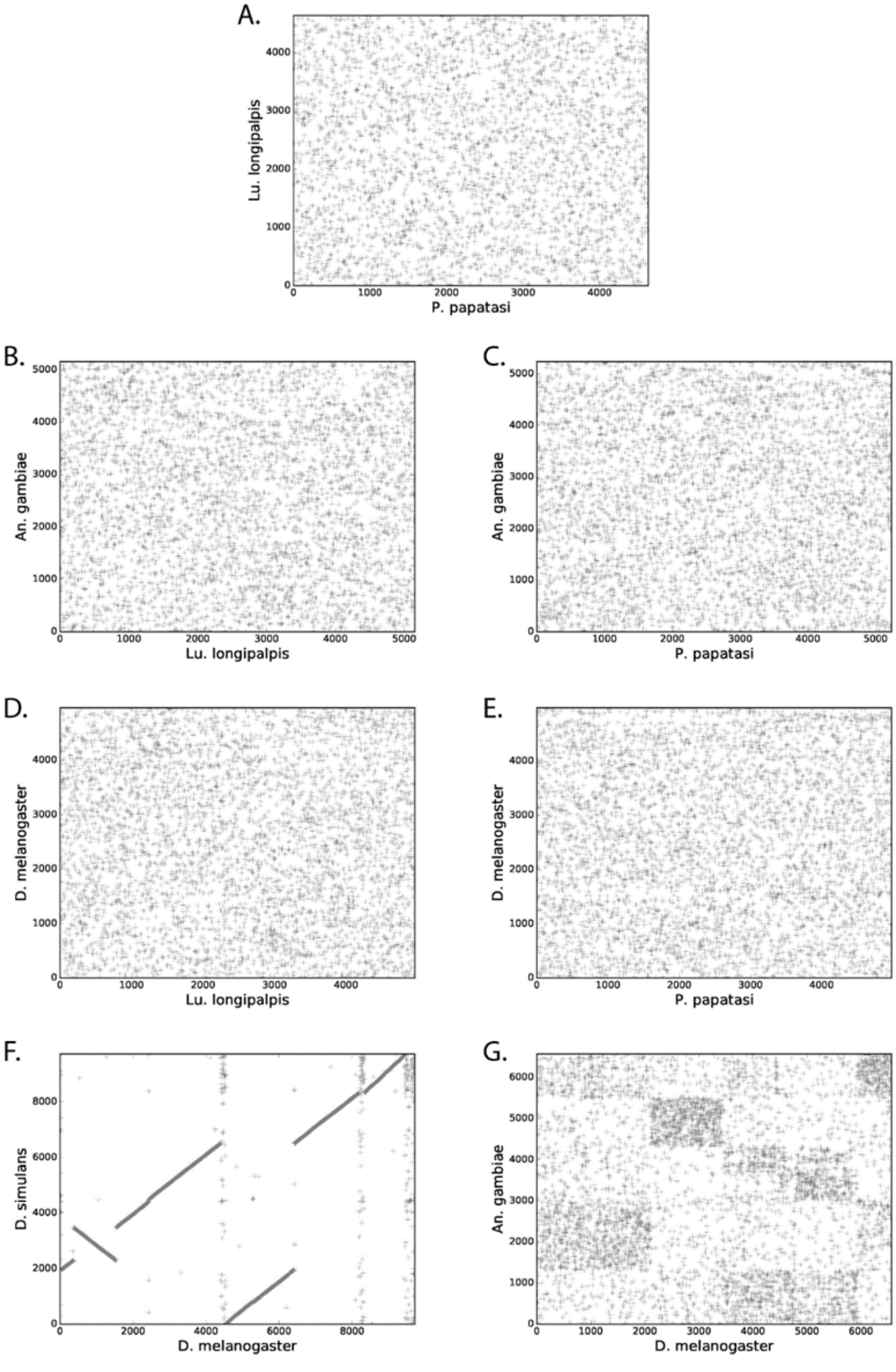

**S2 Figure. Scatter plots of ortholog locations in each pair of genomes.** 1- to -1 ortholog groups were associated utilizing OrthoDB and scaffolds with only one gene were removed prior to generating plots. (**A**) *P. papatasi* vs. *Lu. longipalpis*. (**B**) *Lu. longipalpis* vs. *An. gambiae*. (**C**) *P. papatasi* vs. *An. gambiae*. (**D**) *Lu. longipalpis* vs. *D. melanogaster*. (**E**) *P. papatasi* vs. *D. melanogaster*. (**F**) *D. melanogaster* vs. *D. simulans*. (**G**) *D. melanogaster* vs. *An. gambiae*.

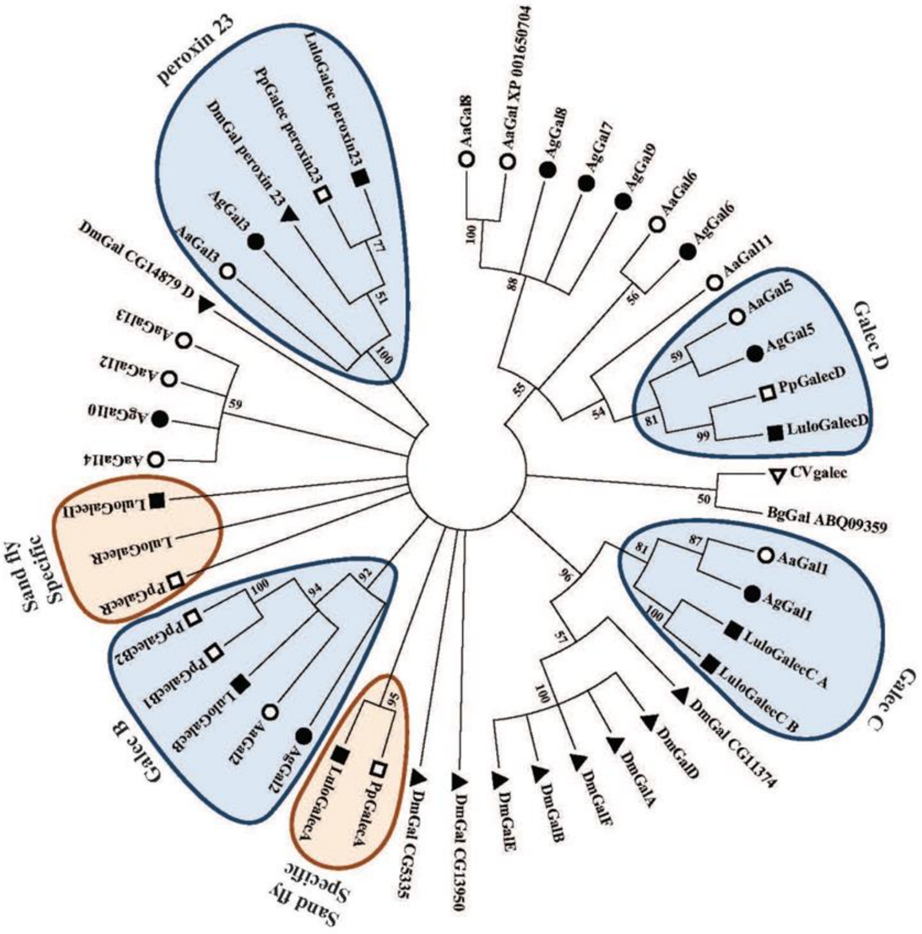

**S3 Figure. Clustering of sand fly galectin protein sequences.** Condensed Neighbor-Joining tree depicting clustering among galectin protein sequences of sand flies (*P. papatasi* and *Lu. longipalpis*; open and filled squares, respectively), mosquitoes (*Ae. aegypti* and *An. gambiae*; open and filled circles, respectively), fly (*D. melanogaster;* filled triangle), eastern oyster (*C. virginica*; upside-down open triangle), and freshwater snail (*B. glabrata*; upside-down filled triangle). Branches encompassing shared orthologs are highlighted by blue shades. Sand fly specific clusters and genes are highlighted by orange shades. The evolutionary distances were computed using the p-distance method and are in the units of the number of amino acid differences per site. One thousand bootstrap replicates were performed, and only branches displaying at least 50% confidence are shown.

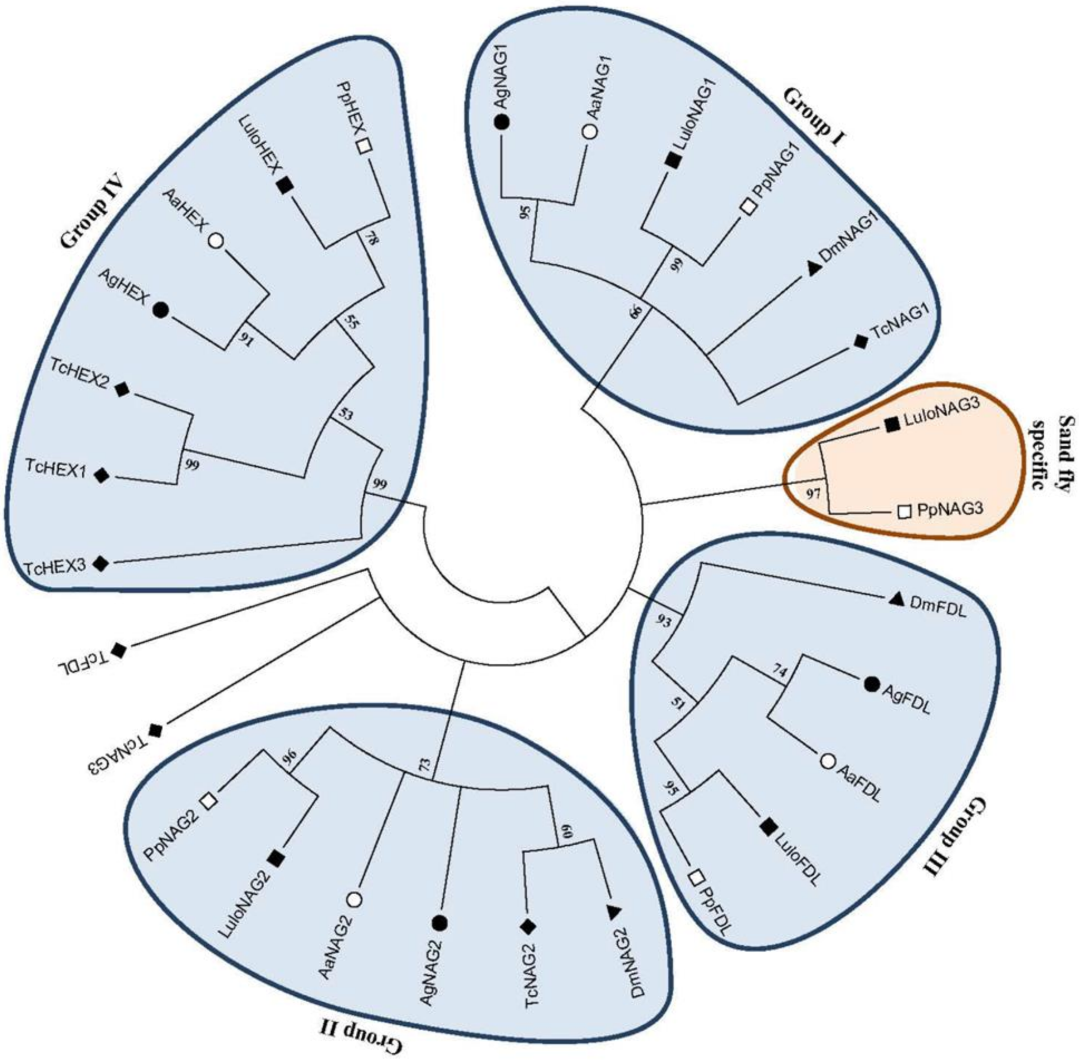

**S4 Figure. Condensed Neighbor-Joining tree depicting clustering among n-acetylhexosaminidase** protein sequences of sand flies (*P. papatasi* and *Lu. longipalpis*; open and filled squares, respectively), mosquitoes (*Ae. aegypti* and *An. gambiae*; open and filled circles, respectively), fly (*D. melanogaster*; filled triangle), and beetle (*T. castaneum*; filled diamond). Branches encompassing sequences belonging to group I-IV n-acetylhexosaminidases are highlighted by a blue shade. The sand fly specific cluster is highlighted by an orange shade. The evolutionary distances were computed using the p-distance method and are in the units of the number of amino acid differences per site. One thousand bootstrap replicates were performed, and only branches displaying at least 50% confidence are shown.

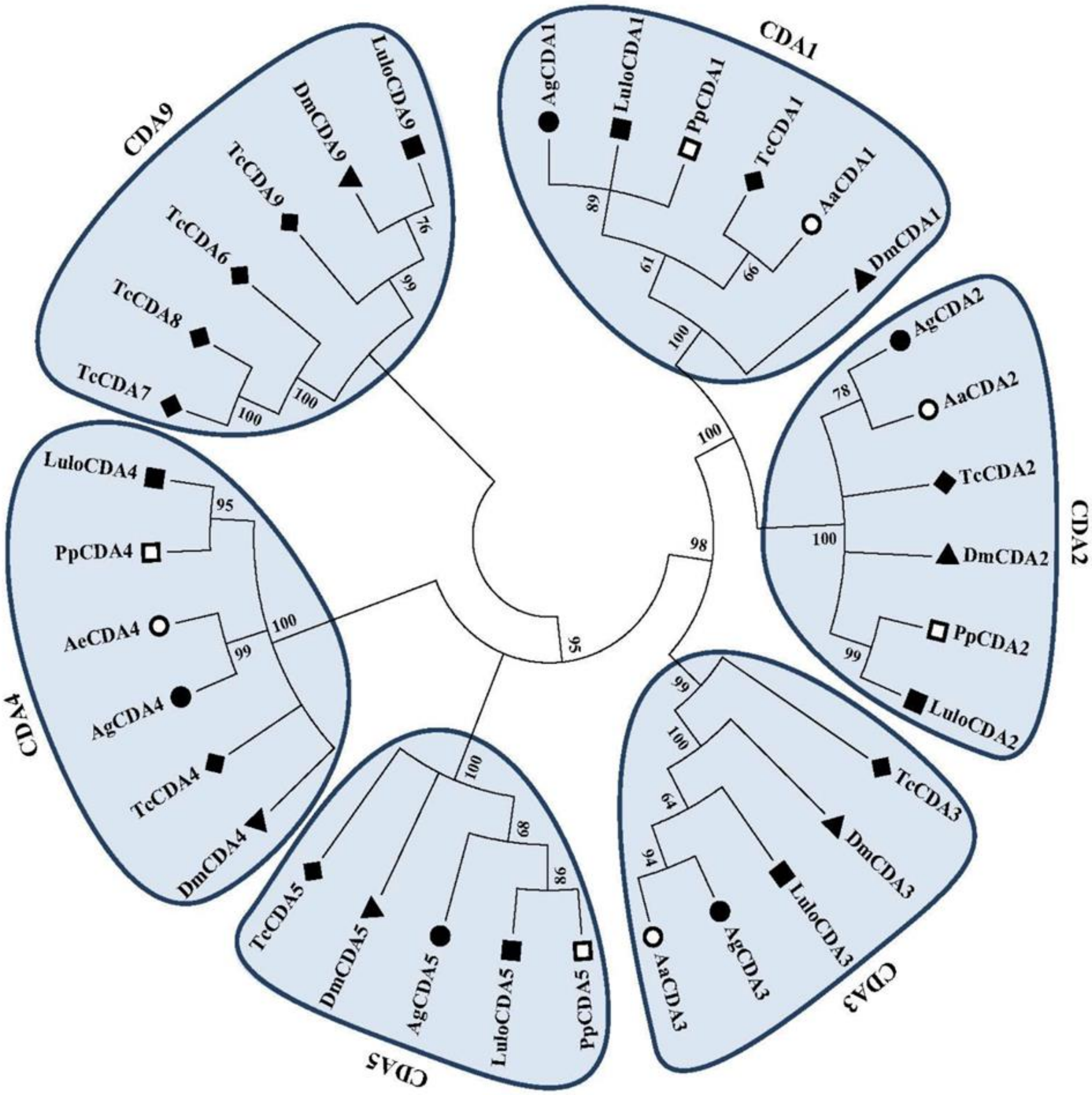

**S5 Figure. Condensed Neighbor-Joining tree depicting clustering among chitin deacetylase catalytic domain sequences** of sand flies (*P. papatasi* and *Lu. longipalpis*; open and filled squares, respectively), mosquitoes (*Ae. aegypti* and *An. gambiae*; open and filled circles, respectively), fly (*D. melanogaster*; filled triangle), and beetle (*T. castaneum*; filled diamond). Branches encompassing sequences belonging to group 1-5 and 9 CDA are highlighted by blue shades. The evolutionary distances were computed using the p-distance method and are in the units of the number of amino acid differences per site. One thousand bootstrap replicates were performed, and only branches displaying at least 50% confidence are shown.

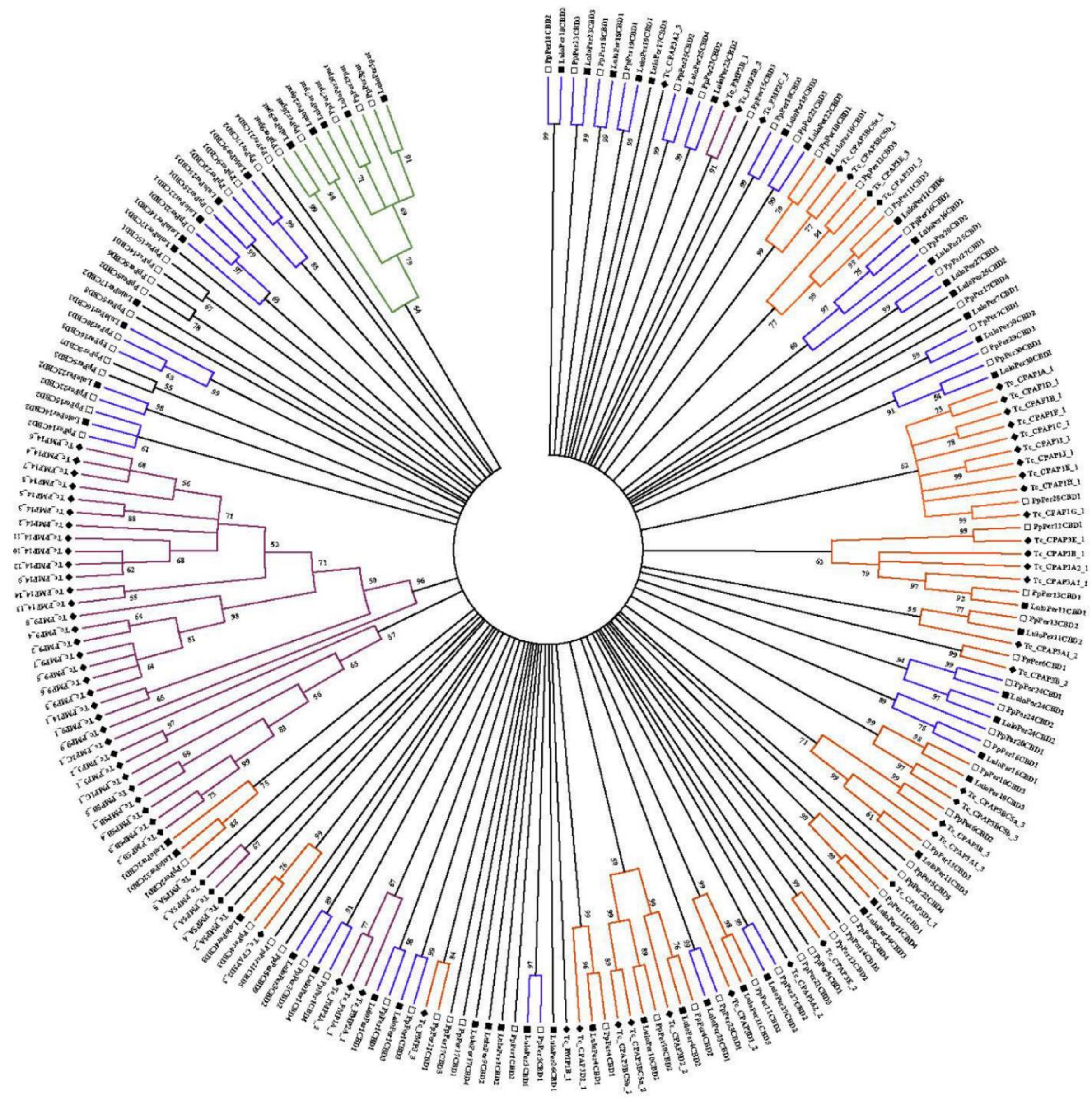

**S6 Figure. Condensed Maximum likelihood tree depicting peritrophin CBD domain similarities** among the sand flies *P. papatasi* and *Lu. longipalpis* and the red flour beetle *T. castaneum*. Open squares, filled squares, and filled diamonds represent *P. papatasi*, *Lu. longipalpis*, and *T. castaneum* domains, respectively. Branches exclusive to *T. castaneum* were color-coded in magenta; those specific to sand flies were highlighted in blue. The branch encompassing the CBD-like domain "CBDput" is highlighted in green. The branches shared by sand flies and RFB CBD domains are color-coded in orange. Maximum likelihood tree was constructed using the Whelan and Goldman (WAG) model with Gamma distributed among Invariant sites (G+I), as suggested by the Model test function of the Mega6 software. One thousand bootstrap replicates were performed, and only branches displaying at least 50% confidence are shown.

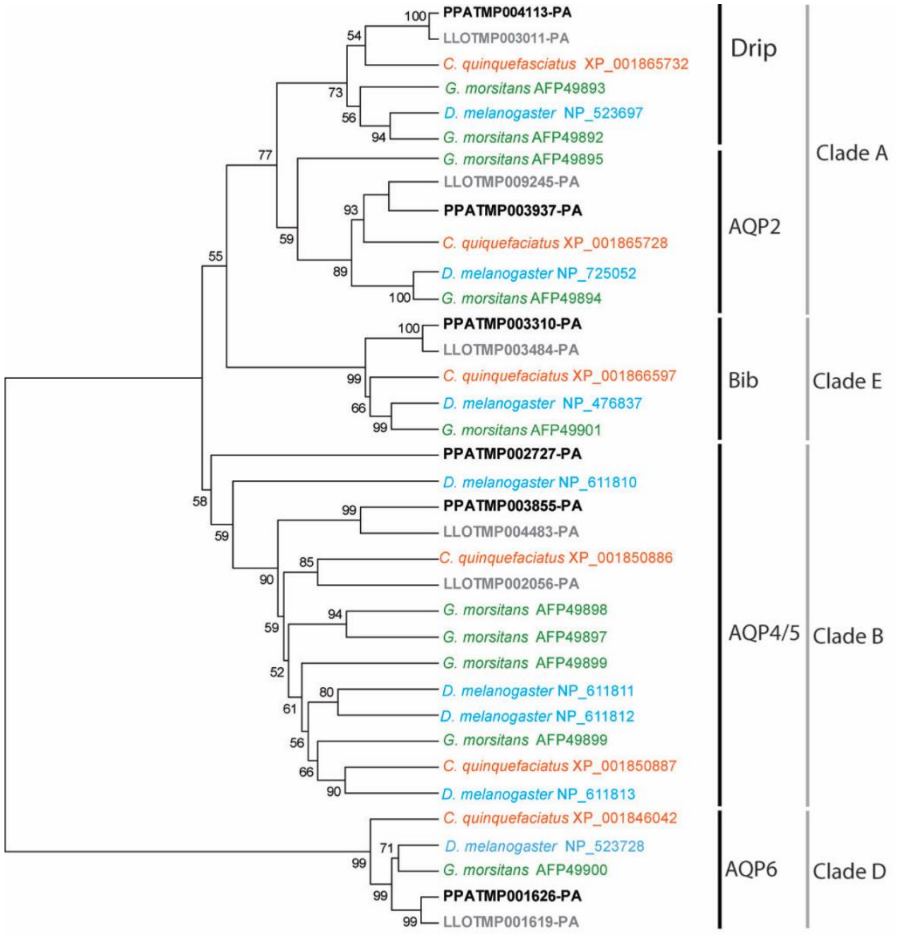

**S7 Figure. Comparison of predicted aquaporins from other flies.** Neighbor-joining tree was produced using MEGA6 using Dayhoff Model and pairwise matching; branch values indicate support following 3000 bootstraps; values below 50% are omitted.

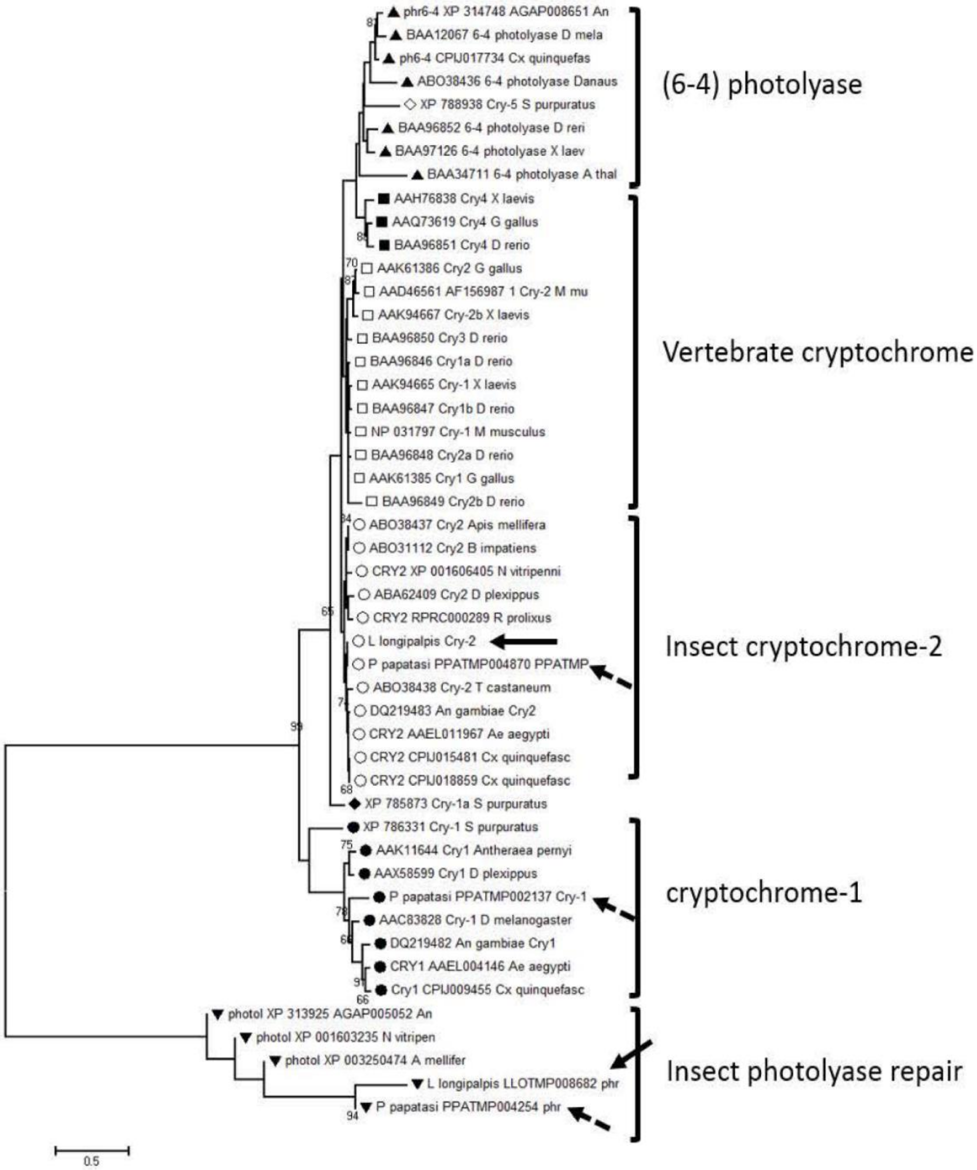

**S8 Figure. Molecular phylogenetic analysis of vertebrate and invertebrate photolyases containing *Lu. longipalpis* and *P. papatasi* gene models.** The different photoyases are displayed on the right. The evolutionary history was inferred by using the Maximum Likelihood method based on the Jones-Taylor-Thorton + four gamma categories with 1000 bootstrap replicates (showing only above 65). Sequences with squares are vertebrate cryptochromes (black – cry-4, white – cry-1, cry-2, and cry-3); sequences with black triangles represent (6-4) insect photolyases; sequences with inverted black triangles are reprenting all insect photolyase repair proteins; and sequences with a dot symbol show insect cryptochromes (black –cry-1, white –cry-2). Dashed arrows point to *P. papatasi* photolyase sequences and straight arrows to *Lu. longipalpois* photolyase sequences.

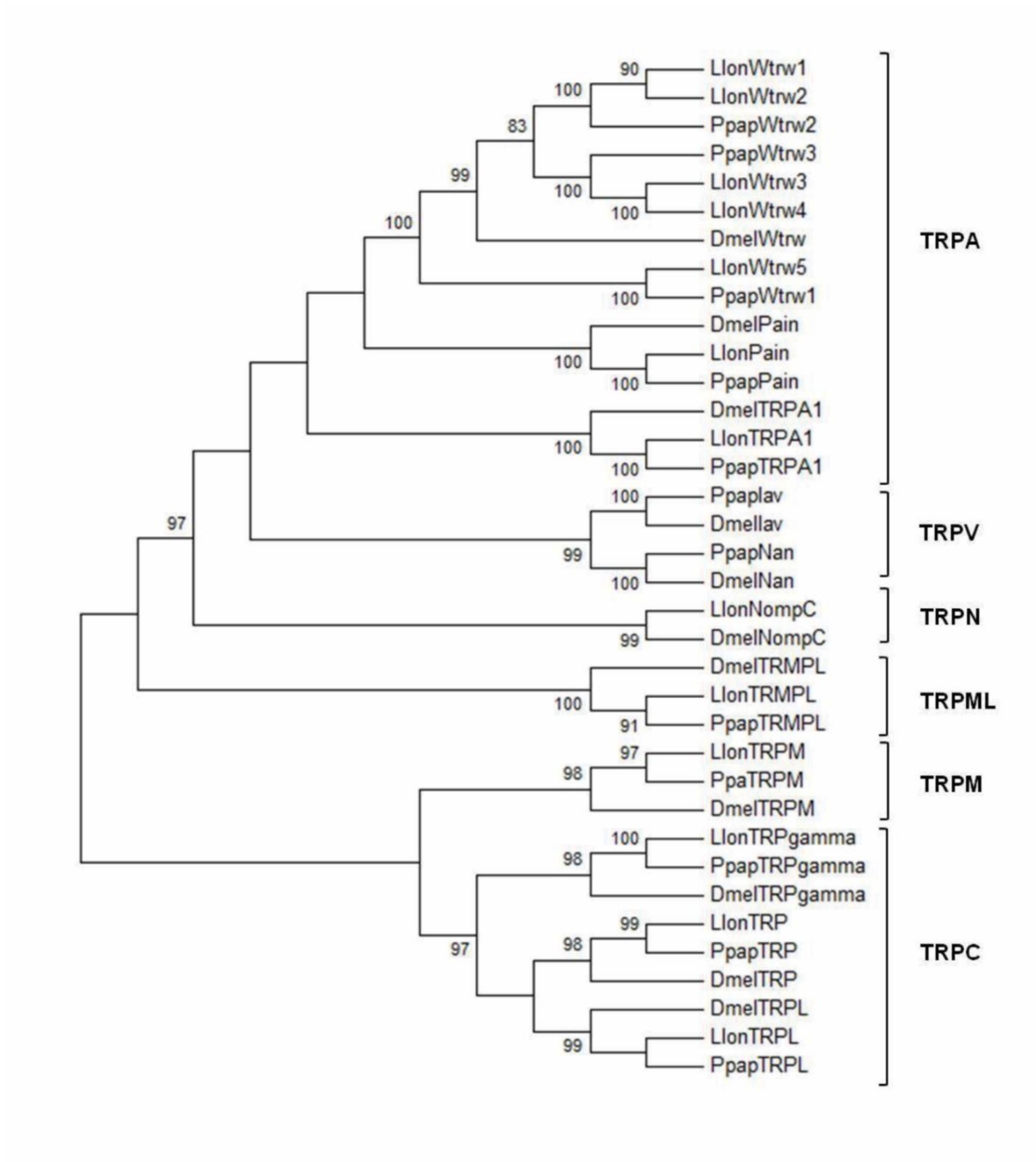

**S9 Figure. Molecular phylogenetic analysis of *Lu. longipalpis, P. papatasi* and *D. melanogaster* TRP channel sequences.** The different TRP subfamilies are displayed on the right. The evolutionary history was inferred by using the Maximum Likelihood method based on the Whelan and Goldman +Freq. model with 1000 bootstrap replicates.

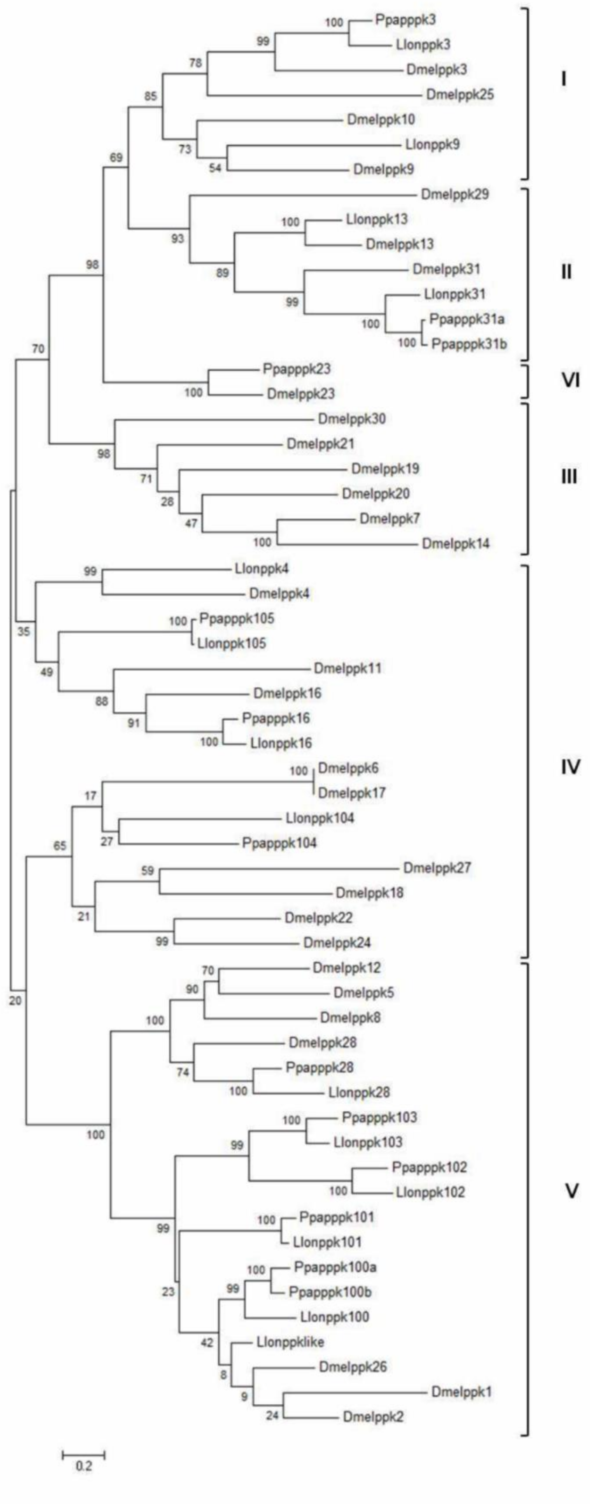

**S10 Figure. Molecular phylogenetic analysis of *Lu. longipalpis, P. papatasi* and *D. melanogaster* PPK sequences.** The different PPK subfamilies are displayed on the right. The evolutionary history was inferred by using the Maximum Likelihood method based on the Whelan and Goldman +Freq. model with 1000 bootstrap replicates.

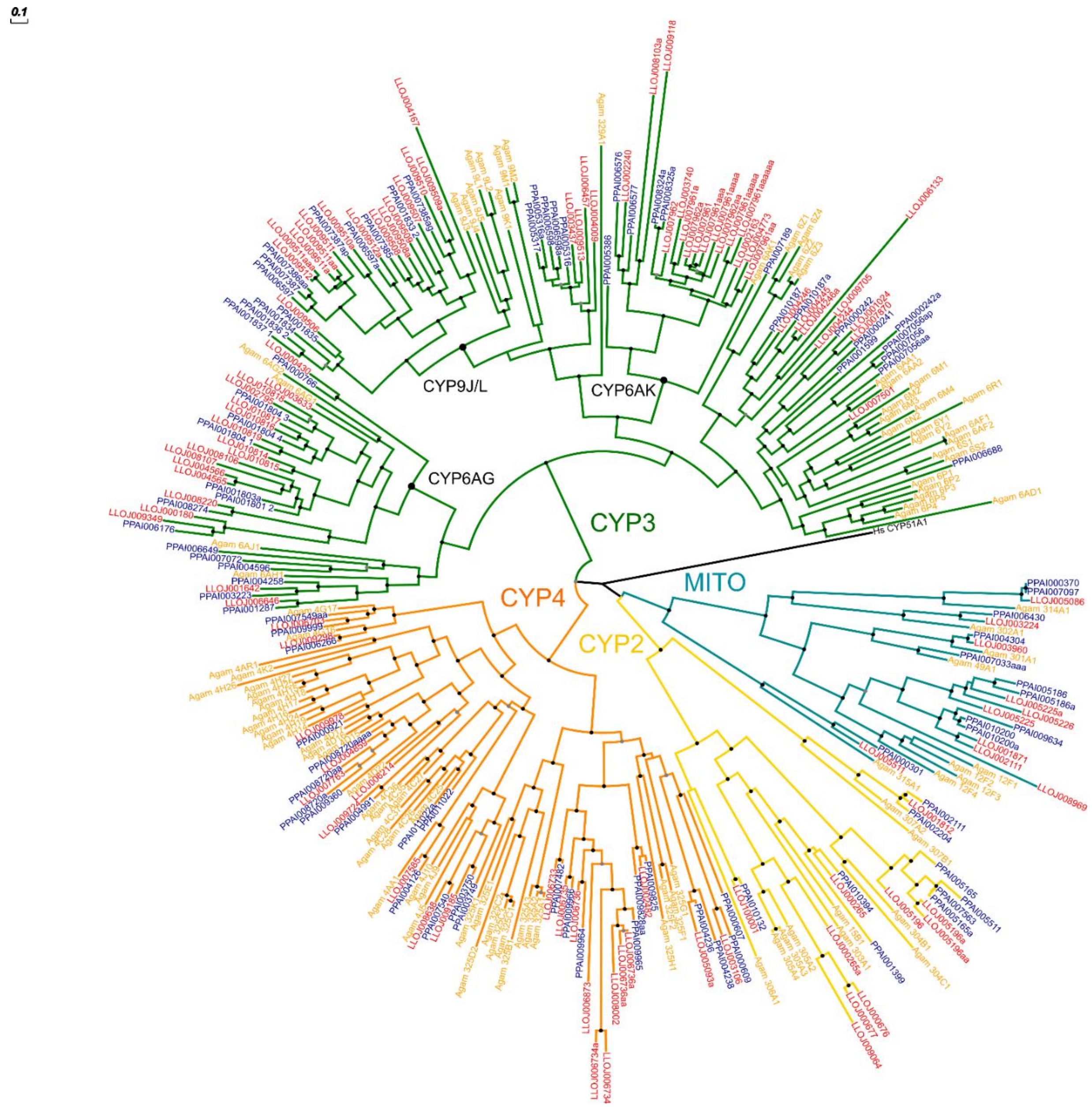

**S11 Figure. Comparison of predicted aquaporins from other flies.** Neighbor-joining tree as produced using MEGA6 using Dayhoff Model and pairwise matching; branch values indicate support following 3000 bootstraps; values below 50% are omitted.

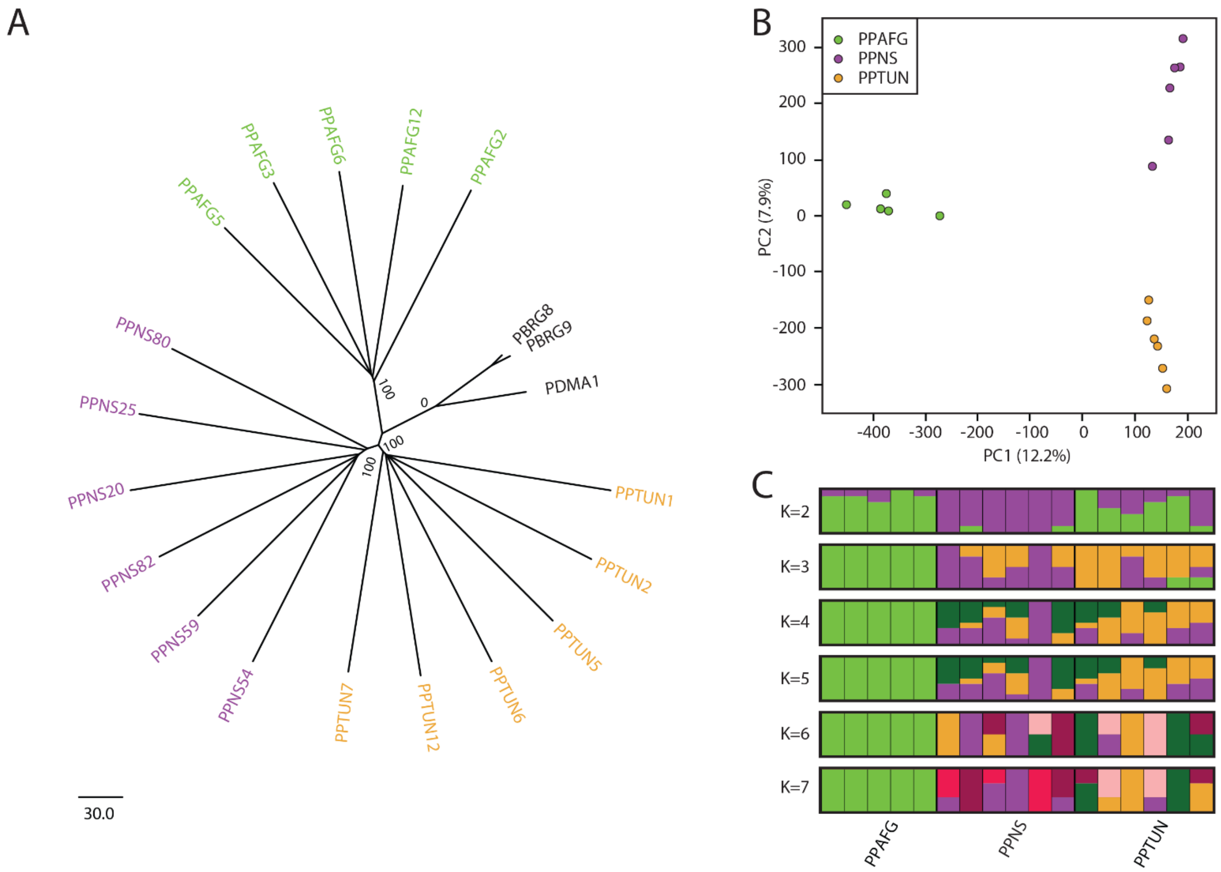

**S12 Figure. *Phlebotomus papatasi* population structure.** Inferred population structure of *P. papatasi* individuals collected from Afghanistan (PPAFG; green), North Sinai - Egypt (PPNS; purple), and Tunisia (PPTUN; orange). (**A**) Phylogenetic Analysis. Rooted neighbor joining (NJ) radial tree generated with the Adegenet and ape packages of R. We included both *P. bergeroti* (PBRG; black) and *P. duboscqi* (PDMA; gray), and used *P. duboscqi* to root the trees. Bootstrap values represent the percentage of 1,000 replicates. (**B**) Principle component analysis (PCA). Individuals are plotted according to their coordinates on the first two principal components (PC1 and PC2). (**C**) Admixture analysis. Ancestry proportions for Admixture models from *K* = 2 to *K* = 7 ancestral populations. Each individual is represented by a thin vertical line, partitioned into *K* coloured segments representing the individual's estimated membership fractions to the *K* clusters. These data are the average of the major *q*-matrix clusters derived by CLUMPAK analysis.

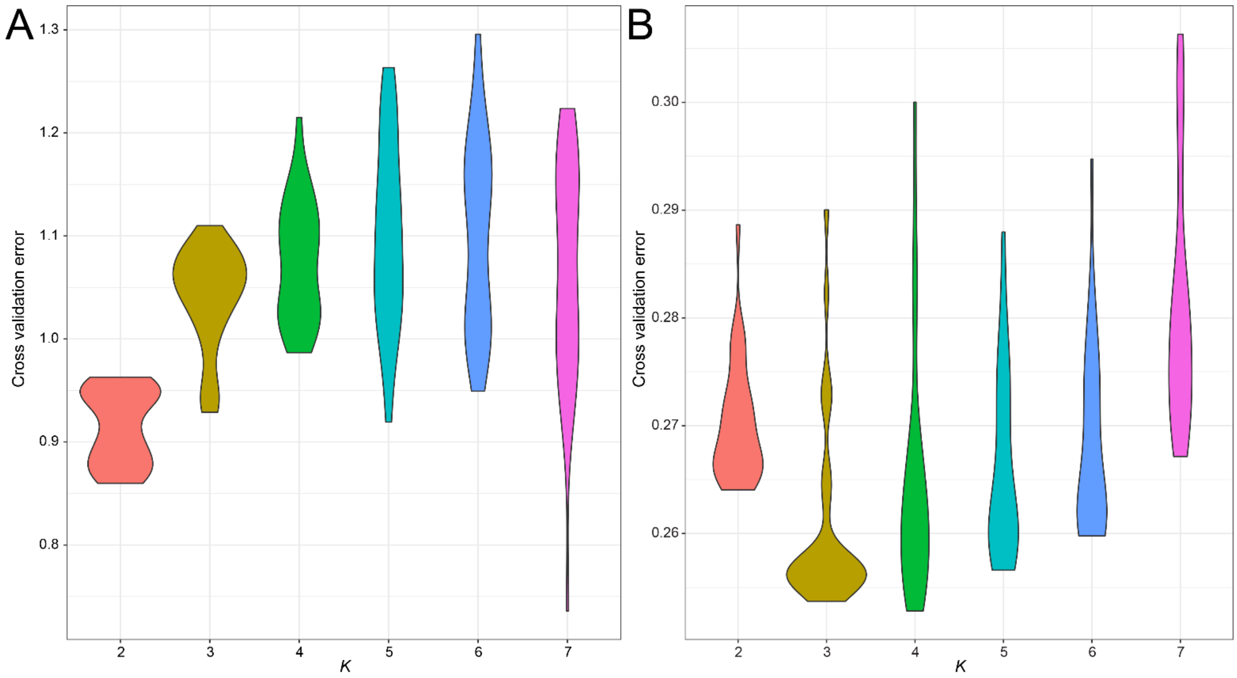

**S13 Figure. Admixture cross validation error**. Violin plot of the cross-validation error for each of 30 replicates for each *K* value. (**A**) *Phlebotomus papatasi* populations. (**B**) *Lutzomyia longipalpis* populations.

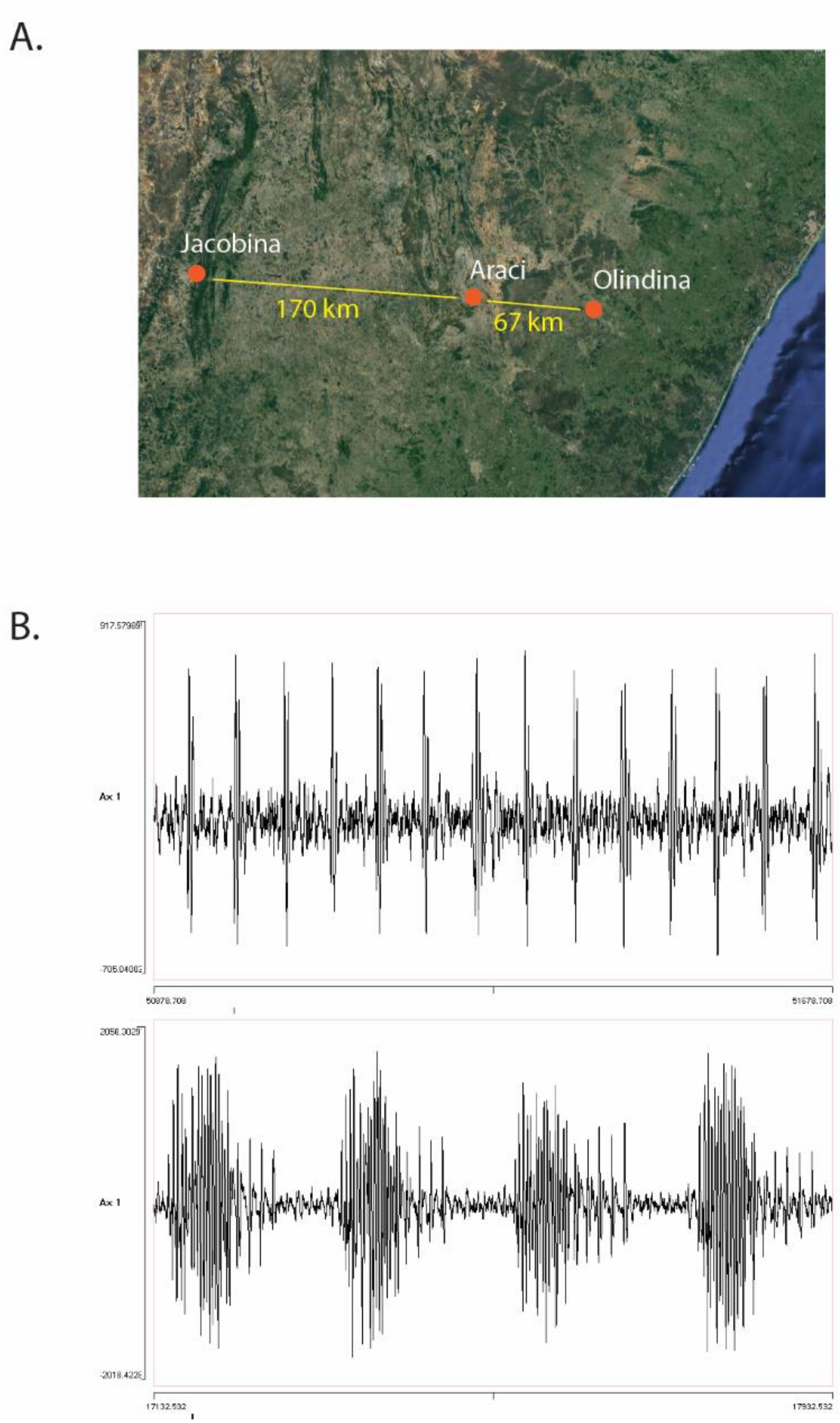

**S14 Figure. Male copulatory courtship songs from Araci and Olinda.** (**A**) Approximate distance of Araci and Olinda from Jacobina. (**B**) Male copulatory courtship song tracings of *Lutzomyia longialpis* males collected from Araci and Olindina. The figure shows ~1 s of song in each case. Map Source: Google Imagery TerraMetrics.

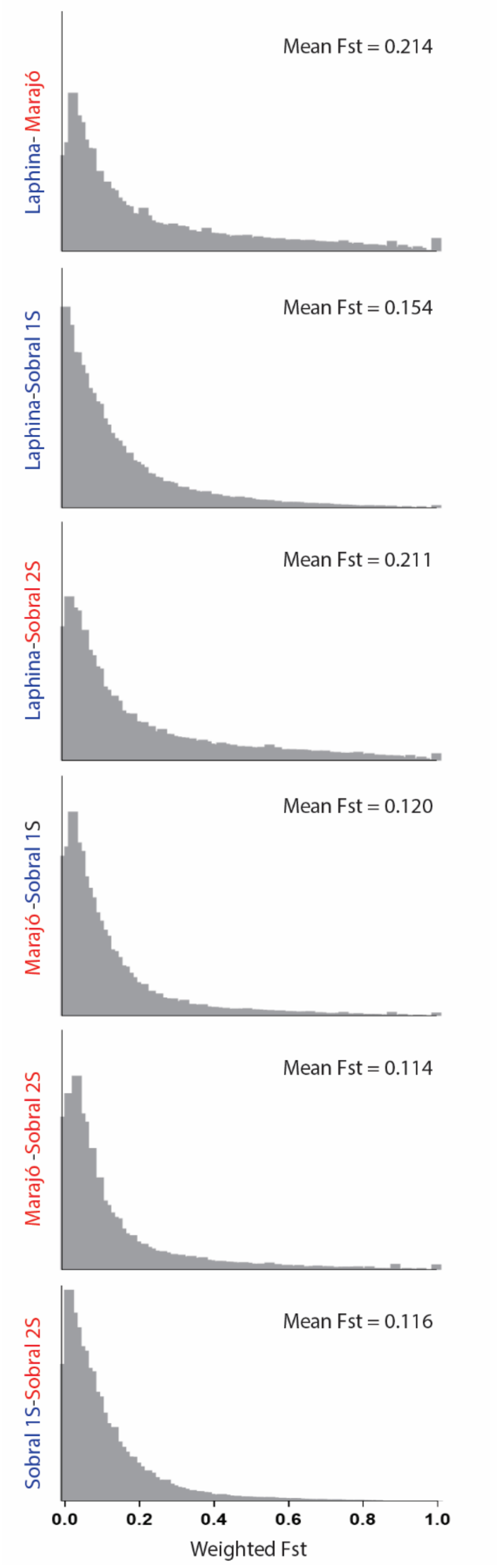

**S15 Figure. Distribution plots of the pairwise $F_{ST}$ between the different populations of *Lutzomyia longipalpis*.** Weighted $F_{ST}$ values for 1kb non-overlapping windows were calculated across the genome for each population comparison.

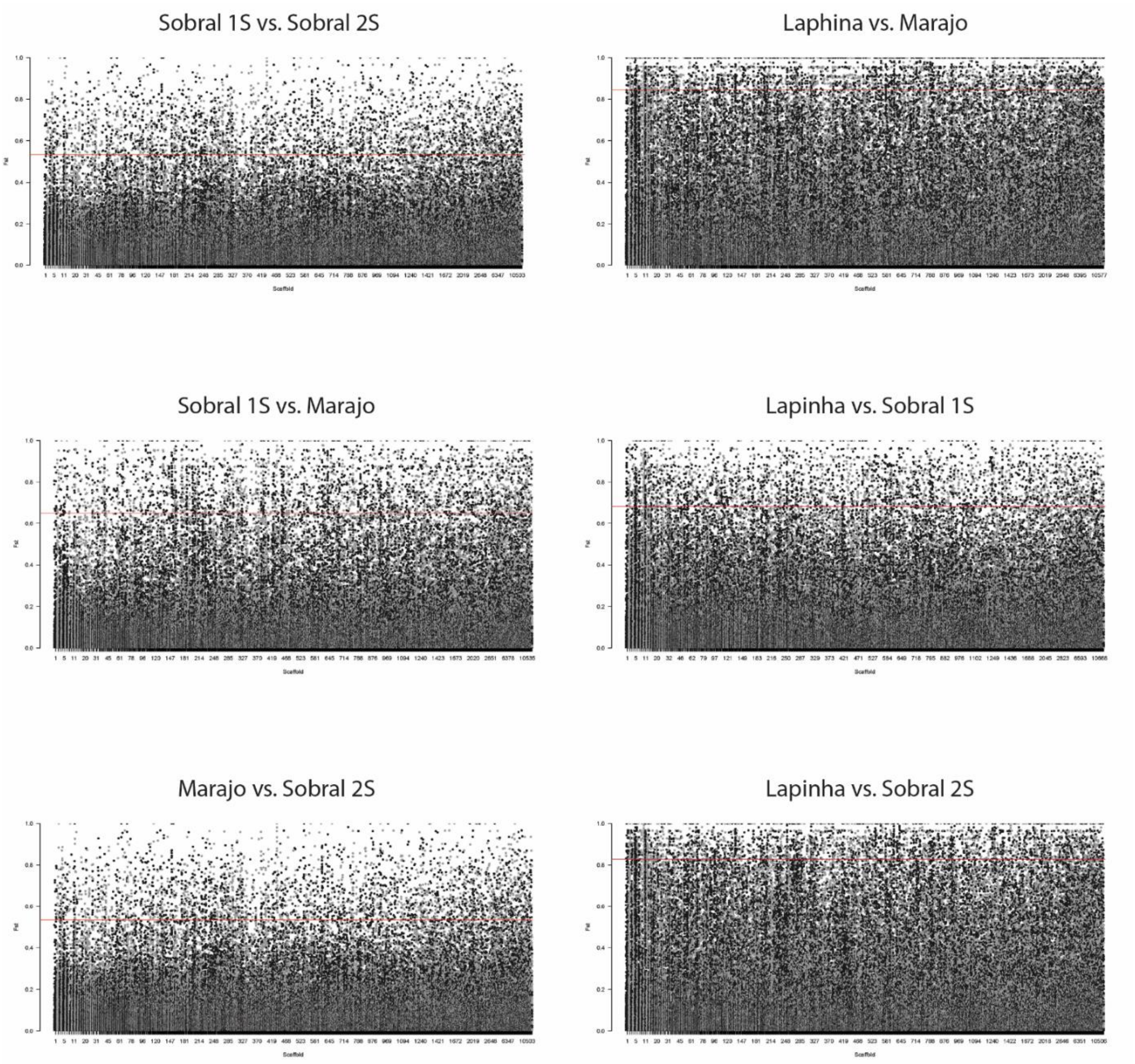

**S16 Figure. Manhattan plots of the pairwise $F_{ST}$ between the different populations of *Lutzomyia longipalpis*.** The red horizontal lines indicate the upper 0.05% of $F_{ST}$ distribution over the entire genome.

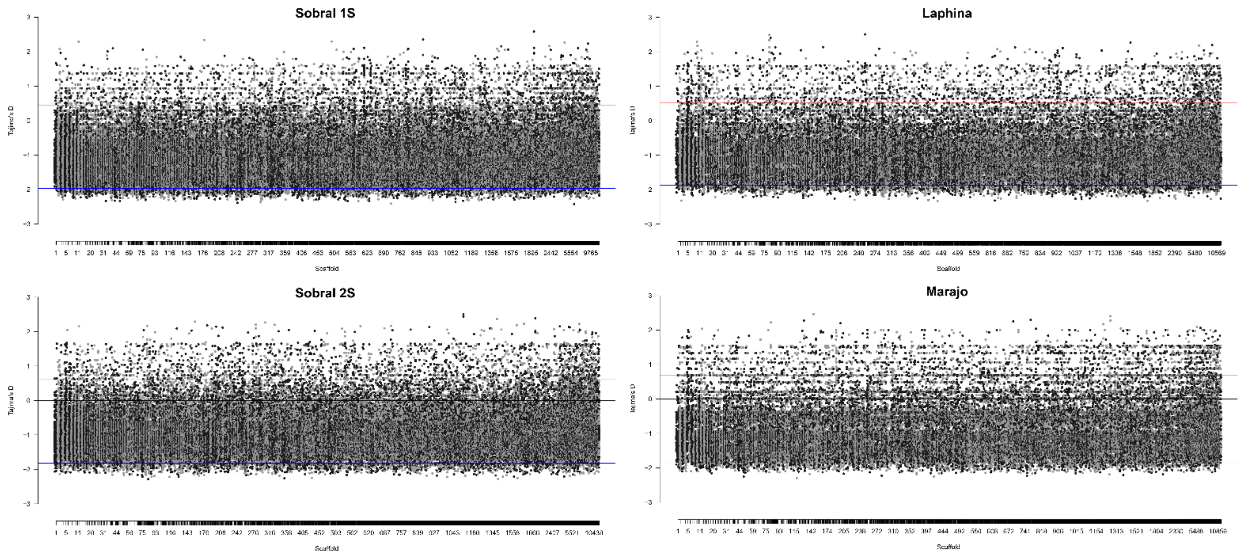

**S17 Figure. Manhattan plot of Tajimas'D for each population of *Lutzomyia longipalpis*.** The red and blue horizontal lines indicate the upper and lower 0.05% of Tajima's D distribution, respectively.

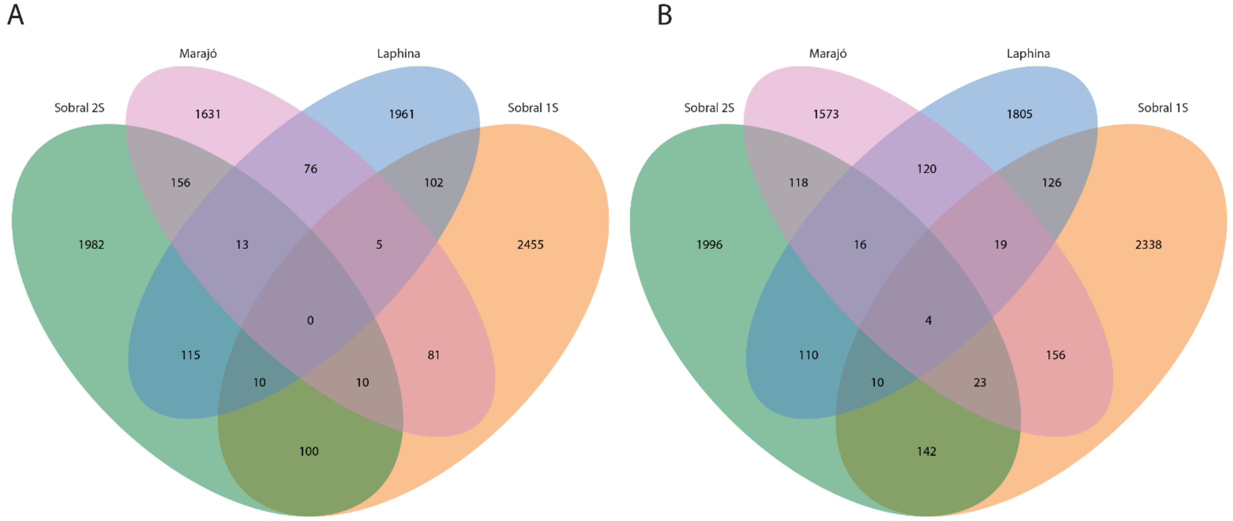

**S18 Figure. Genomic regions with high (outlier) Tajimas'D for different populations of *Lutzomyia longipalpis*.** (**A**) The Venn diagram summarizes the numbers of 1 kb genomic windows with Tajimas'D values in the upper 2.5% of the different populations. (**B**) The Venn diagram summarizes the numbers of 1 kb genomic windows with Tajimas'D values in the lower 2.5% of the different populations.

**S1 Data File:** *Phlebotomus papatasi* CYPome Fasta File.
**S2 Data File:** *Lutzomyia longipalpis* CYPome Fasta File.

### SUPPLEMENTAL TABLES

**Table S1.** Assembly statistics of *Phlebotomus papatasi* and *Lutzomyia longipalpis* draft genome assemblies.

| Feature | *P. papatasi* | *Lu. longipalpis* |
|---|---|---|
| Assembly Version | Ppap_1.0 | Llon_1.0 |
| GenBank Assembly | GCA_000262795.1 | GCA_000265325.1 |
| Assembly Size | 363.768 Mb | 154.229 Mb |
| Gap Length | 18,378,439 | 11,456,874 |
| Spanned Gaps | 32,373 | 24,164 |
| Sequencing coverage | 345.39 Mb | 147.77 Mb |
| Total number of scaffolds | 106,826 | 11,532 |
| Scaffold N50 | 28 kb | 85.1 kb |
| Scaffold L50 | 2,066 | 491 |
| Total number of contigs | 139,199 | 35,696 |
| Contig N50 | 5,795 | 7,481 |
| Contig L50 | 16,042 | 5,523 |
| GC Content | 34.3 % | 35.9 % |
| Gene Set | Ppapl1.4 | LlonJ1.4 |
| # Predicted protein-coding | 11,216 | 10,311 |
| % of Genome | 0.003 % | 0.007 % |
| Non-coding genes | 444 | 339 |
| Number of RNAseq | 11,664 | 10,699 |
| Protein-coding transcripts | 11,220 | 10,330 |

**Table S2.** BUSCO assessment.

| BUSCOs | *P. papatasi* | *Lu. longipalpis* |
|---|---|---|
| Total | 262 | 261 |
| Complete and single copy | 249 | 238 |
| Complete and duplicate B | 13 | 23 |
| Fragmented | 32 | 9 |
| Missing | 9 | 33 |
| Total searched | 303 | 303 |
| **Completeness** | 86.5% | 86.1% |

**Table S3.** RNAseq Samples *Lu. Longipalpis.*

| Species | BioSample | Run | Secondary Sample Accession | MBases | Sample name | feeding pattern | life stage |
|---|---|---|---|---|---|---|---|
| *Lu. Longipalpis* | SAMN01096292 | SRR535760 | SRX174785 | 4,042 | LLON_S0H-1 | sugar fed from same batch of bloodfed/infected insects | adult female |
| | - | SRR535761 | SRX174785 | 3,114 | | | |
| | SAMN01096293 | SRR535765 | SRX174788 | 2,476 | LLON_S0H-2 | sugar fed from same batch of bloodfed/infected insects | adult |
| | SAMN01096290 | SRR535770 | SRX174792 | 3,852 | LLON_I6H-1 | 6h after blood feeding a meal containing 2x10 6 /ml Leishmania infantum | adult |
| | - | SRR535771 | SRX174792 | 2,956 | | | |
| | SAMN01096291 | SRR535774 | SRX174794 | 2,287 | LLON_I6H-2 | 6h after blood feeding a meal containing 2x10 6 /ml Leishmania infantum | adult |
| | SAMN01096295 | SRR535758 | SRX174784 | 3,755 | LLON_L4-2 | | fourth instar larvae |
| | - | SRR535759 | SRX174784 | 2,845 | | | |
| | SAMN01096406 | SRR535779 | SRX174797 | 2,998 | LLON_L4-4 | | fourth instar larvae |
| | SAMN01096284 | SRR535777 | SRX174796 | 3,075 | LLON_B6H-1 | 6h after blood feeding | adult |
| | - | SRR535778 | SRX174796 | 2,345 | | | |
| | SAMN01096285 | SRR535757 | SRX174783 | 1,156 | LLON_B6H-2 | 6h after blood feeding | adult |
| | SAMN01096288 | SRR535766 | SRX174789 | 4,672 | LLON_I24H-1 | 24h after blood feeding a meal containing 2x10 6 /ml Leishmania infantum | adult |
| | | SRR535767 | SRX174789 | 3,543 | | | |
| | SAMN01096289 | SRR535768 | SRX174790 | 1,891 | LLON_I24H-2 | 24h after blood feeding a meal containing 2x10 6 /ml Leishmania infantum | adult |
| | SAMN01096282 | SRR535772 | SRX174793 | 3,556 | LLON_B24H-1 | 24h after blood feeding | adult |
| | | SRR535773 | SRX174793 | 2,757 | | | |

| | | | | | | | |
|---|---|---|---|---|---|---|---|
| | SAMN01096283 | SRR535756 | SRX174782 | 1,793 | LLON_B24H-2 | 24h after blood feeding | adult |
| | SAMN01096286 | SRR535775 | SRX174795 | 3,041 | LLON_I144H-1 | 144h after blood feeding a meal containing 2x10 6 /ml Leishmania infantum | adult |
| | | SRR535776 | SRX174795 | 2,312 | | | |
| | SAMN01096287 | SRR535769 | SRX174791 | 1,786 | LLON_I144H-2 | 144h after blood feeding a meal containing 2x10 6 /ml Leishmania infantum | adult |
| | SAMN01096280 | SRR535762 | SRX174786 | 4,302 | LLON_B144H-1 | 144h after blood feeding | adult |
| | | SRR535763 | SRX174786 | 3,272 | | | |
| | SAMN01096281 | SRR535764 | SRX174787 | 2,735 | LLON_B144H-2 | 144h after blood feeding | adult |
| *P. papatasi* | SAMN05936232 | SRR4446962 | SRX2265643 | 5,891 | 730019 | Sugar fed | adult female |
| | | SRR4446804 | SRX2265636 | 337 | | | |
| | | SRR4446797 | SRX2265626 | 364 | | | |
| | SAMN05936231 | SRR4446967 | SRX2265644 | 6,940 | 730015 | Sugar fed | adult male |
| | | SRR4446806 | SRX2265638 | 357 | | | |
| | | SRR4446798 | SRX2265625 | 384 | | | |
| | SAMN05936235 | SRX2265647 | SRR4446963 | 6,357 | 730031 | | larvae and pupae |
| | | SRR4446802 | SRX2265633 | 367 | | | |
| | SAMN05936234 | SRR4446965 | SRX2265641 | 5,375 | 730027 | 6 hours post blood meal | adult female |
| | | SRR4446800 | SRX2265632 | 336 | | | |
| | | SRR4446796 | SRX2265628 | 366 | | | |
| | SAMN05936230 | SRR4446966 | SRX2265646 | 5,871 | 730011 | 6 hours post *L. major*-infected blood meal | adult female |

| | | | | | | |
|---|---|---|---|---|---|---|
| | SRR4446807 | SRX2265640 | 319 | | | |
| | SRR4446794 | SRX2265623 | 347 | | | |
| SAMN05936238 | SRR4446803 | SRX2265634 | 335 | 730049 | 24 hours post blood meal | adult female |
| | SRR4446964 | SRX2265649 | 5,653 | | | |
| | SRR4446793 | SRX2265629 | 396 | | | |
| SAMN05936233 | SRR4446809 | SRX2265645 | 5,346 | 730023 | 24 hours post *L. major*-infected blood meal | adult female |
| | SRR4446808 | SRX2265639 | 352 | | | |
| | SRR4446792 | SRX2265627 | 386 | | | |
| SAMN05936237 | SRR4446968 | SRX2265642 | 3,392 | 730039 | 96 hours post blood meal | adult female |
| | SRR4446805 | SRX2265637 | 214 | | | |
| | SRR4446799 | SRX2265631 | 222 | | | |
| SAMN05936236 | SRR4446969 | SRX2265630 | 7,886 | 730035 | 96 hours post *L. major*-infected blood meal | adult female |
| | SRR4446801 | SRX2265635 | 405 | | | |
| | SRR4446795 | SRX2265630 | 440 | | | |

**Table S4.** Microsynteny Block Statistics.

| Species | Count | Average Size | Median Size | Maximum Size |
|---|---|---|---|---|
| *Lu. longipalpis* vs.*P. papatasi* | 499 | 4.2 | 3 | 24 |
| *Lu. longipalpis* vs. *An. gambiae* | 504 | 4.3 | 3 | 39 |
| *P. papatasi* vs. *An. gambiae* | 307 | 4.3 | 3 | 21 |
| *D. melanogaster* vs. *D. simulans* | 243 | 41.6 | 5 | 700 |
| *D. melanogaster* vs. *An. gambiae* | 1037 | 2.5 | 2 | 10 |

**S5 Table.** Toll pathway annotation.

| Species | Gene | ID | Scaffold | Base pair range on supercontig |
|---|---|---|---|---|
| ***Lu. Longipalpis*** | LuloToll1 | LLOJ007208 | Scaffold544 | 33,079-36,578 |
| | LuloToll2 | LLOJ002222 | Scaffold1536 | 10,062-12,521 |
| | LuloToll3 | LLOJ001885 | Scaffold142 | 134,843-137,926 |
| | LuloToll4 | LLOJ002166 | Scaffold1514 | 1,810-5,706 |
| | LuloToll5 | LLOJ004777 | Scaffold286 | 98,546-100,447 |
| | LuloPGRP-SA1 | LLOJ005642 | Scaffold361 | 60,049-61,864 |
| | LuloPGRP-SA2 | LLOJ002444 | Scaffold1627 | 4,150-12,137 |
| | LuloPGRP-S3 | LLOJ003999 | Scaffold2373 | 8,198-8,820 |
| | LuloPGRP-S4 | LLOJ004539 | Scaffold268 | 40,944-41,418 |
| | LuloGNBP3 | LLOJ002383 | Scaffold16 | 276,750-281,261 |
| | LuloTube | LLOJ005198 | Scaffold327 | 58,512-60,136 |
| | LuloPelle | LLOJ001101 | Scaffold12 | 330,669-334,979 |
| | LuloMyd88 | LLOJ009628 | Scaffold9 | 296,479-306,442 |
| | LuloCac1 | LLOJ004612 | Scaffold272 | 113,310-137,735 |
| | LuloCac2 | LLOJ008968 | Scaffold786 | 67,382-82,912 |
| | LuloDifA | LLOJ002098 | Scaffold15 | 93,362-96,422 |
| | LuloDifB | LLOJ002097 | Scaffold15 | 91,191-93,034 |
| ***P. papatasi*** | PpToll1 | PPAI010510 | Scaffold8 | 1,152,463-1,156,686 |
| | PpToll2 | PPAI010503 | Scaffold8 | 455,955-459,845 |
| | PpToll3 | PPAI006632 | Scaffold42729 | 8,717-12,634 |
| | PpToll4 | PPAI001388 | Scaffold1394 | 15,847-33,278 |
| | PpToll5 | PPAI009420 | Scaffold5805 | 1,406-2,515 |
| | PpPGRP-SA | PPAI010204 | Scaffold729 | 6,726-11,936 |
| | PpPGRP-S2 | PPAI007689 | Scaffold45674 | 827-1,827 |
| | PpGNBP1 | PPAI000880 | Scaffold1225 | 7,143-16,719 |
| | PpGNBP2A | PPAI002587 | Scaffold189 | 14,063-14,970 |
| | PpGNBP2B | PPAI002588 | Scaffold189 | 18,505-19,068 |
| | PpGNBP3 | PPAI010440 | Scaffold78 | 366,823-372,348 |
| | PpTube | PPAI003151 | Scaffold2113 | 26,151-27,620 |
| | PpPelle | PPAI007366 | Scaffold44566 | 625-1,344 |
| | PpMyd88 | PPAI006982 | Scaffold43446 | 2,116-3,039 |
| | PpCac | PPAI010893 | Scaffold90 | 131,533-133,250 |
| | PpDif1 | PPAI001149 | Scaffold13 | 871,814-875,824 |

| Gene | *P. papatasi* | *Lu. longipalpis* | *A. aegypti* | *A. gambiae* | *D. melanogaster* |
|---|---|---|---|---|---|
| **Toll** | 5 | 5 | 2 | 10 | 9 |
| **PGRPs** | 2 | 4 | 4 | 3 | 7 |
| **GNBP** | 4 | 1 | 7 | 7 | 3 |
| **Tube** | 1 | 1 | 1 | 1 | 1 |
| **Pelle** | 1 | 1 | 1 | 1 | 1 |
| **Myd88** | 1 | 1 | 1 | 1 | 1 |
| **Cactus** | 1 | 1 | 1 | 1 | 1 |
| **Dorsal/Dif** | 1 | 1 | 2 | 1 | 1 |

**Table S6.** Insect immune deficiency pathway annotation.

| Species | Gene | ID | Scaffold | Base pair range on supercontig | Observations |
|---|---|---|---|---|---|
| ***Lu. longipalpis*** | PGRP-LC | | | | |
| | PGRP-LB | | | | |
| | PGRP-SC1 | | | | |
| | PGRP-SC2 | | | | |
| | PGRP-LCa | | | | |
| | PGRP-LCx | | | | |
| | PGRP-LE | | | | |
| | IMD | LLOJ002454-RA | Scaffold163 | 71031-73281 | Base error on 5' M |
| | IRD5 | LLOJ003785-RA | Scaffold2255 | 245-421 | Missing 30 AA 5' portion |
| | JRA | | | | |
| | KAY | | | | |
| | KEY | | | | |
| | FADD | | Scaffold2131 | | Missing 5' sequence poor quality (multiple N) |
| | Dredd | LLOJ001647-RA | Scaffold135 | 131418-132026 | |
| | Caspar | LLOJ002950-RA | Scaffold1853 | 10833-14280 | |
| | CaspL1 | | | | |
| | Rel | LLOJ002098-RA | Scaffold15 | 93407-963231 | Missing 5' end |
| | IAP2 | LLOJ000523-RA | Scaffold1082 | 30325-42500 | |
| | Tab2 - TAK1-associated binding protein 2 | LLOJ009749-RA | Scaffold920 | 31630-35530 | Missing 3' sequence poor quality (multiple N) |
| | Tak1 | | | | |
| | Pirk | LLOJ004926-RA | Scaffold30 | 90777-91250 | |
| | UBI-P63E | | | | |
| | UEV1A | | | | |
| | AP1 | | | | |
| | DSP1 | | | | |
| | Caudal | LLOJ004702-RA | Scaffold280 | 6964-7506 | Base error on 5' M |
| | RPD3 | | | | |
| | PUC | | | | |
| | Attacin-C | LLOJ005408-RA | Scaffold340 | 100805-101365 | |
| | Cecropin | | Scaffold1655 | | Poor quality at Scaffold. Present at good quality in contigs. I have the full annotated sequence in house. |

| | | | | |
|---|---|---|---|---|
| POSH | | | | |
| dUSP36 | | | | |
| SKPA/SLMB/DCUL1 | | | | |
| Bendless | LLOJ004109-RA | Scaffold244 | 65316-65005 | Complete CDS |
| CYLD | LLOJ007012-RA | Scaffold52 | 162005-177559 | Complete CDS |
| EFFETE | LLOJ002835-RA | Scaffold1784 | 294-2239 | Complete CDS |
| DNR-1 | LLOJ004935-RA | Scaffold30 | 160039-183921 | Complete CDS. There are 17 X in the middle of the sequence that should be removed. |
| CalcineurinB | LLOJ006044-RA | Scaffold401 | 112545-117231 | ROS Pathway - Complete CDS |
| MKP3 | LLOJ005232-RA | Scaffold33 | 67772-94556 | ROS Pathway - Complete CDS |
| p38b | LLOJ010513-RA | Scaffold6 | 119076-124326 | ROS Pathway - Complete CDS |
| COX | LLOJ003300-RA | Scaffold2 | 591-621594 | ROS Pathway - Complete CDS |
| DuOX | | | | ROS Pathway; coding sequence split into 3 parts: First part is LLOTMP010038-RA (Scaffold986:9698-23477:1) coding for the animal peroxidase domain. THIS NEEDS TO BE CONFIRMED; Second part is LLOTMP002199-RA (Scaffold1528:18696-19934:-1) coding for EF hand 1 and Ferric reductase like transmembrane component; Third part is LLOTMP002198-RA (Scaffold1528:17305-18670:-1) coding for FAD-binding domain 8 and Ferric reductase NAD binding domain 6. |
| DuOX partial seq - 5' end | LLOJ010503-RA | Scaffold986 | 20640-23477 | ROS Pathway - Partial sequence. This is the 5' end |

| | | | | | |
|---|---|---|---|---|---|
| | | | | | of the DUOX gene. Missing 3' end. The 3' end is on scaffold 1528. |
| | DuOX partial seq - 3' end | LLOJ010494-RA | Scaffold1528 | 18080-19983 | ROS Pathway - Partial sequence. This is the 3' end of the DUOX gene. Missing 5' end. The 5' end is on scaffold 986. |
| | TEP1 | LLOJ007923-RA | Scaffold634 | 37315-55768 | Complement-like protein - Complete CDS |
| | Fosfolipase C beta | LLOJ001939-RA | Scaffold1440 | 37196-45904 | Complete CDS |
| | Prophenoloxidase | LLOJ001742-RA | Scaffold1380 | 6504-13245 | Complete CDS |
| *P. papatasi* | IMD | PPAI001062-RA | Scaffold129 | 4371-21382 | |
| | IRD5 | | | | |
| | Dredd | | | | |
| | Caspar | | | | |
| | Relish | PPAI012820-RA | Scaffold10 | 266434-284397 | IMD Pathway. Missing 5' end. |
| | IAP2 | | | | |
| | Tab2 - TAK1-associated binding protein 2 | | | | |
| | Pirk | | | | |
| | Caudal | | | | |
| | Attacin-C | | | | |
| | Bendless | PPAI004722-RA | Scaffold289 | 3738-4393 | IMD Pathway. Complete CDS |
| | CYLD | PPAI005913-RA | Scaffold373 | 689-4946 | IMD Pathway. Complete CDS |
| | EFFETE | PPAI001122-RA | Scaffold13 | 491487-495614 | IMD Pathway. Complete CDS |
| | DNR-1 | PPAI003240-RA | Scaffold217 | 119829-126407 | IMD Pathway. Complete CDS |
| | CalcineurinB | PPAI009875-RA | Scaffold66 | 353071-358750 | ROS Pathway - Complete CDS |
| | MKP3 | PPAI004376-RA | Scaffold268 | 90864-94448 | ROS Pathway - Missing 5' end |
| | p38b | | | | |
| | COX | PPAI012814-RA | Scaffold126 | 40550-86015 | Complete CDS |
| | DuOX | PPAI003114-RA | Scaffold21 | 558486-576040 | ROS Pathway - Complete CDS |
| | TEP1 | PPAI005309-RA | Scaffold3235 | 5995-8652 | Complement-like protein. Missing 5' end |
| | Fosfolipase C epsilon | PPAI010847-RA | Scaffold893 | 35958-47337 | ROS Pathway - Complete CDS |
| | Prophenoloxidase | PPAI010450-RA | Scaffold783 | 24285-27358 | Complete CDS |

**Table S7.** Jak/Stat pathway annotation.

| Species | Gene | Transcript | Scaffold # | Base pair range on supercontig |
|---|---|---|---|---|
| *Lu. longipalpis* | Dome | LLOJ002449-PA | Scaffold163 | 14907-41091 |
| | Hopscotch | LLOJ005862-PA | Scaffold389 | 19572-22451 |
| | Mekk1 | LLOJ007004-PA | Scaffold518 | 34530-52570 |
| | P38b | LLOJ010513-PA | Scaffold6 | 113170-124326 |
| | Socs36e | LLOJ009450-PA | Scaffold872 | 49378-71024 |
| | Stat92e | LLOJ007427-PA | Scaffold57 | 82155-112275 |
| | Upd3 | LLOJ007923-PA | Scaffold634 | 37315-55768 |
| | | LLOJ007922-PA | Scaffold634 | 31843-37409 |
| *P. papatasi* | Dome | PPAI001065-PA | Scaffold129 | 44265-62863 |
| | Hopscotch | PPAI002271-PA | Scaffold173 | 157296-159579 |
| | Mekk1 | PPAI009824-PA | Scaffold651 | 37748-50232 |
| | P38b | na | na | na |
| | Socs36e | na | na | na |
| | Stat92e | PPAI002926-PA | Scaffold2012 | 4799-9580 |
| | Upd3 | PPAI003567-PA | Scaffold2333 | 2089-8960 |
| | | PPAI005310-PA | Scaffold3235 | 8990-14592 |
| | | PPAI005309-PA | Scaffold3235 | 5995-8652 |

| **JAK/STAT gene name** | ***P. papatasi*** | ***Lu. Longipalpis*** | ***D. Melanogaster*** | ***An. Albimanus*** | ***An. Gambiae*** | ***Ae. Aegypti*** | ***Cu.quinqu*** | ***Rh. Prolixus*** |
|---|---|---|---|---|---|---|---|---|
| Hopscotch | 1 | 1 | 1 | 1 | 1 | 1 | 1 | 1 |
| Domeless | 1 | 1 | 1 | 1 | 1 | 1 | 1 | 1 |
| STAT92E | 1 | 1 | 1 | 0 | 1 | 1 | 1 | 1 |
| SOCS36E | 0 | 1 | 1 | 1 | 1 | 1 | 1 | 1 |
| p38b | 0 | 1 | 1 | 1 | 1 | 3 | 2 | 1 |
| Upd3 | 3 | 2 | 11 | 5 | 11 | 8 | 18 | 1 |
| Mekk1 | 1 | 1 | 2 | 1 | 1 | 1 | 1 | 1 |
| **Total** | 7 | 8 | 18 | 10 | 17 | 16 | 25 | 7 |

**S8 Table.** Galectin family annotation.

Detailed description of galectins identified in sand flies is shown. Gene names, scaffold whereby genes are mapped, base pair range in the scaffold, GenBank accession number, and number of GLEC domains are described.

| Species | Gene | Transcript | Scaffold # | Base pair range on supercontig | GenBank (V1.1) accession # | GLEC domain |
|---|---|---|---|---|---|---|
| *Lu. longipalpis* | **LuloGalecA** | LLOJ010541-RA | Scaffold269 | 31641-32834 | | 3 |
| | **LuloGalecB** | LLOJ003096-RA | Scaffold193 | 93715-95078 | | 2 |
| | **LuloGalecC** | LLOJ007344-RA | Scaffold561 | 15695-16176 | | 4 |
| | **LuloGalecD** | LLOJ000277-RA | Scaffold1032 | 11171-13201 | | 1 |
| | **LuloGalecR** | LLOJ009549-RA | Scaffold887 | 26206-31999 | ABV60341 | 1 |
| | **LuloGalec_Peroxin23** | LLOJ004425-RA | Scaffold2617 | 22399-25050 | | 1 |
| | **LuloGalecII** | LLOJ009548-RA | Scaffold887 | 21503-24893 | | 1 |
| *P. papatasi* | **PpGalecA** | PPAI007147-RA | Scaffold43916 | 1632-2853 | AY538600 | 2 |
| | **PpGalecBa** | PPAI007331-RA | Scaffold44416 | 367-1564 | | 2 |
| | **PpGalecBb** | PPAI002605-RA | Scaffold1890 | 16200-17339 | | 2 |
| | **PpGalecC** | PPAI008432-RA | Scaffold57589 | 3109-3571 | | 1 |
| | **PpGalecD** | PPAI000755-RA | Scaffold1198 | 41229-42179 | | 1 |
| | **PpGalecR** | PPAI007711-RA | Scaffold45770 | 626-2887 | | 1 |

Number of galectin genes.

| Galectins | *P. papatasi* | *Lu. longipalpis* | *A. aegypti* | *A. gambiae* | *D. melanogaster* |
|---|---|---|---|---|---|
| **Total** | **7** | **8** | **12** | **10** | **6** |

**Table S9.** Transforming growth factor-beta family annotation.

| Species | Gene | Transcript | Scaffold # | Base pair range on supercontig | TGF-beta domain | SMAD domain | TGF-beta receptor domain | Metallopeptidase domain | CUB domain | EGF domain | WD-40 domain | F-box domain | Magnesium transporter domain | CHRD domain | VWF dom |
|---|---|---|---|---|---|---|---|---|---|---|---|---|---|---|---|
| *Lu. longipalpis* | Lldpp | LLOJ006419-RA | Scaffold451 | 22413-30051 | 1 | | | | | | | | | | |
| | Llmav | LLOJ005524-RA | Scaffold351 | 125246-134929 | 1 | | | | | | | | | | |
| | Llmad | LLOJ009024-RA | Scaffold796 | 36871-56646 | | 2 | | | | | | | | | |
| | Llsmox | LLOJ009024-RA | Scaffold796 | 36871-56646 | | 2 | | | | | | | | | |
| | Llpunt | LLOJ000177-RA | Scaffold1012 | 4730-9275 | | | 1 | | | | | | | | |
| | Llsax | LLOJ006339-RA | Scaffold439 | 10108-12433 | | | 2 | | | | | | | | |
| | Llbabo | LLOJ003539-RA | Scaffold21 | 230942-233465 | | | 1 | | | | | | | | |
| | Lltkv | LLOJ008051-RA | Scaffold655 | 31338-42397 | | | 1 | | | | | | | | |
| | Lltld | LLOJ000403-RA | Scaffold106 | 54120-68892 | | | | 1 | 1 | 1 | | | | | |
| | Llfus | LLOJ005418-RA | Scaffold341 | 8631-10441 | | 2 | | | | | | | | | |
| | LlIrk2 | LLOJ008806-RA | Scaffold763 | 30635-38904 | | 2 | | | | | | | | | |
| | Llslmb | LLOJ006764-RA | Scaffold5 | 129585-140234 | | | | | | | 1 | 1 | | | |
| | Llspict | LLOJ005267-RA | Scaffold332 | 66570-68090 | | | | | | | | | 1 | | |
| | Llsog | LLOJ000799-RA | Scaffold1131 | 54895-73573 | | | | | | | | | | 1 | 1 |
| *P. papatasi* | Ppdpp | PPAI007784-RA | Scaffold46 | 552344-557726 | 1 | | | | | | | | | | |
| | Ppmav | PPAI008670-RA | Scaffold51 | 176758-179366 | 1 | | | | | | | | | | |
| | Ppmad | PPAI007902-RA | Scaffold4654 | 1349-8308 | | 2 | | | | | | | | | |
| | Ppsmox | PPAI007902-RA | Scaffold4654 | 1349-8308 | | 2 | | | | | | | | | |

| | | | | | | | | | | | | | | |
|---|---|---|---|---|---|---|---|---|---|---|---|---|---|---|
| Pppunt | PPAI006514-RA | Scaffold420 | 50724-55103 | | | 1 | | | | | | | | |
| Ppsax | PPAI010817-RA | Scaffold889 | 41494-42901 | | | 2 | | | | | | | | |
| Ppbabo | PPAI008219-RA | Scaffold4821 | 5284-7860 | | | 1 | | | | | | | | |
| Pptkv | PPAI008219-RA | Scaffold4821 | 5284-7860 | | | 1 | | | | | | | | |
| Pptld | PPAI008935-RA | Scaffold530 | 54753-60174 | | | | 1 | 1 | 1 | | | | | |
| Ppfus | PPAI010627-RA | Scaffold842 | 2486-4923 | | 2 | | | | | | | | | |
| PpIrk2 | PPAI002168-RA | Scaffold1689 | 25974-31932 | | 2 | | | | | | | | | |
| Ppslmb | PPAI004879-RA | Scaffold299 | 2427-7576 | | | | | | | 1 | 1 | | | |
| Ppspict | PPAI010360-RA | Scaffold762 | 29884-36922 | | | | | | | | | 1 | | |
| Ppsog | PPAI001672-RA | Scaffold15 | 606454-613653 | | | | | | | | | | 1 | 1 |

TGF-beta domain: Conserved TGF-beta domain that includes two of the conserved cysteines.
SMAD domain: SMAD or MAD domain found in MAD related proteins that bind to DNA.
TGF-beta receptor domain: Serine/threonine kinase receptor domain that include a single transmembrane domain and a specific hydrophilic Cys-rich ligand-binding domain.
Metallopeptidase domain: Catalytic domain of metallopeptidases composed by a three-layer alpha-beta-alpha structure.
CUB domain: Structural motif found in extracellular and plasma membrane-associated proteins.
EGF domain: Epidermal growth factor domain that includes six cysteine residues involved in disulphide bonds.
WD-40 domain: Repeated WD40 motifs function as a site for protein-protein interactions involved in signal transduction and transcription regulation.
F-box domain: Identified first in cyclin-F as a protein-protein interaction motif that is necessary to link protein complexes such as Skp1-cullin-F-box protein ligase complexes.
Mg transporter domain: Catalysis of the transfer of magnesium ions through a membrane.
CHRD domain: The CHRD domain is a novel domain identified in chordin, an inhibitor of bone morphogenetic proteins.
VWF domain: The von Willebrand factor domain is present in intracellular proteins involved in transcription, DNA repair, ribosomal and membrane transport .

**S10 Table.** Mitogen activated protein kinase family annotation.

| Species | Gene | Transcript | Scaffold/contig | Base pair range | GenBank |
|---|---|---|---|---|---|
| *Lu. Longipalpis* | **MAPK1 (ERK2)** | LLOJ008154-RA | Scaffold670 | 28939-29878 | |
| | **MAPK8 (JNK1)** | LLOJ005677-RA | Scaffold366 | 138910-157006 | |
| | **MAPK11 (P38b)** | LLOJ010513-RA | Scaffold6 | 119076-124326 | |
| | **MAPK15 (ERK7)** | LLOJ004347-RA | Scaffold257 | 15915- 20039 | |
| | **MAP2K1 (MEK1/DSOR1)** | LLOJ010496-RA | Scaffold188 | 166292-169419 | |
| | **MAP2K3** | | orphan | | ABR28347.1 |
| | **MAP2K4** | LLOJ002043-RA | Scaffold148 | 2404-7521 | |
| | **MAP2K7** | LLOJ001452-RA | Scaffold1298 | 37784-39128 | |
| | **MAP3K4 (MEKK1)** | LLOJ007004-RA | Scaffold518 | 34530-52701 | |
| | **MAP3K5 (MEKK5/6)** | | Contig35012 | 451- 3738 | |
| | **MAP3K7 (TAK)** | LLOJ010544-RA | Scaffold694 | 43814-49966 | |
| | **MAP3K10 (MLK2/3)** | LLOJ002365-RA | Scaffold1595 | 15830-28107 | |
| | **MAP3K12/13** | LLOJ009180-RA | Scaffold817 | 25883-37021 | |
| | **B-RAF** | LLOJ006454-RA | Scaffold459 | 15641-18266 | |
| | **MOS** | LLOJ006057-RA | Scaffold405 | 39152-40413 | |
| | **MAP4K1** | LLOJ002088-RA | Scaffold1498 | 18047-32210 | |
| | **MAP4K4** | LLOJ009619-RA | Scaffold9 | 56864-99818 | |
| *P. papatasi* | **MAPK1 (ERK2)** | PPAI005249-RA | Scaffold319 | 80689-88419 | |
| | | PPAI008443-RA | Scaffold49775 | 1948-2751 | |
| | **MAPK8 (JNK1)** | PPAI008376-RA | Scaffold493 | 233-770 | |
| | | PPAI008377-RA | Scaffold493 | 1385-2127 | |
| | **MAPK11 (P38b)** | PPAI003699-RA | Scaffold23631 | 767-5980 | |
| | **MAPK15 (ERK7)** | PPAI009597-RA | Scaffold605 | 52694-55083 | |
| | | PPAI009598-RA | Scaffold605 | 55180-55764 | |
| | **MAP2K1 (MEK1/DSOR1)** | PPAI008154-RA | Scaffold479 | 28213-35690 | |
| | **MAP2K3** | PPAI006033-RA | Scaffold383 | 64914-71075 | |
| | **MAP2K4** | PPAI009824-RA | Scaffold651 | 37748-50232 | |
| | **MAP2K7** | PPAI000315-RA | Scaffold1069 | 18678-22947 | |
| | | PPAI000316-RA | Scaffold1069 | 26311-27225 | |
| | | PPAI000317-RA | Scaffold1069 | 27774-28720 | |
| | **MAP3K4 (MEKK1)** | PPAI005224-RA | Scaffold3171 | 2786-5853 | |
| | **MAP3K5 (MEKK5/6)** | PPAI001629-RA | Scaffold149 | 29277-30044 | |
| | | PPAI001630-RA | Scaffold149 | 29059-71054 | |
| | **MAP3K7 (TAK)** | PPAI009080-RA | Scaffold54377 | 1843-2321 | |
| | | PPAI009094-RA | Scaffold4525 | 4084-4926 | |
| | **MAP3K10 (MLK2/3)** | PPAI000857-RA | Scaffold122 | 118782-122138 | |
| | **MAP3K12/13** | PPAI003648-RA | Scaffold23552 | 110168-11004 | |
| | **B-RAF** | PPAI002155-RA | Scaffold1680 | 24036-28926 | |
| | **MOS** | PPAI003648-RA | Scaffold23552 | 10168-11004 | |
| | **MAP4K1** | PPAI002155-RA | Scaffold1680 | 24036-28926 | |
| | **MAP4K4** | PPAI003333-RA | Scaffold2209 | 8155-23951 | |
| | | PPAI003751-RA | Scaffold2380 | 3199-8715 | |
| | | PPAI005831-RA | Scaffold3648 | 2344-7192 | |

**S11 Table.** Prophenoloxidase family annotation.

| Species | Gene | Transcript | Scaffold # | Base pair range |
|---|---|---|---|---|
| *Lu. longipalpis* | | LLOJ009210-RA | Scaffold823 | 38321-41617 |
| | | LLOJ001742-RA | Scaffold1380 | 6504-13245 |
| *P. papatasi* | | PPAI010450-RA | Scaffold783 | 24285-27358 |
| | | PPAI012836-RA | Scaffold46630 | 3170-3867 |
| | | PPAI002201-RA | Scaffold17 | 423823-427052 |

**S12 Table.** Salivary protein annotation.

| Salivary Family of proteins | Sand fly species | Function | Protein name | Transcript | Exons | supercontig # | Base pair range on supercontig | GenBank (V1.1) accession # from vector base | GenBank (V1.1) accession # from gene bank | Job ID | comments |
|---|---|---|---|---|---|---|---|---|---|---|---|
| Yellow related proteins | *Lutzomyia longipalpis* | Biogenic amine binding proteins | LJM11 | LLOJ001468-RA | 173412-17357<br>173601-174065<br>175015-175943<br>176002-178309 | Scaffold 13 incomplete sequence | 173412-178309 | AY445935 | AY445935 | 23895 | |
| Yellow related proteins | *Lutzomyia longipalpis* | Biogenic amine binding proteins | LJM17 | LLOJ001469-RA | 182575-183683<br>182745-182884<br>182967-183680<br>183742-184039 | Scaffold 13 incomplete sequence | 182575-184039 | AF132518 | AF132518 | 23898 | |
| Yellow related proteins | *Lutzomyia longipalpis* | Anti-inflammatory /anti-arthritis | LJM111 | LLOJ001468-RA | 174811-174945<br>175011-175943<br>176001-176221 | Scaffold 13 | 174811-176221 | DQ192488 | DQ192488 | 23899 | |
| D7 related protein | *Lutzomyia longipalpis* | unknown | LJL13 | LLOJ009780-RA | 4450-4375<br>4309-3914<br>3846-3378 | Scaffold 926 | 4450-3378 | AF420274 | AF420274 | 23900 | |
| Silk related/collagen binding/32kDa protein | *Lutzomyia longipalpis* | unknown | LJL04 | LLOJ004915-RA/RB | 11256-11193<br>10979-10941<br>10849 -10651<br>10587-10256 | Scaffold 30 | 11256-9756 | AY455906 | AY455906 | 23901 | |

| | | | | | | | | | | | |
|---|---|---|---|---|---|---|---|---|---|---|---|
| | | | | | 10087-9756 | | | | | | |
| Antigen-5 protein | *Lutzomyia longipalpis* | unknown | LJL34 | LLOJ002578-RA | 847-667 601-148 | Scaffold 1694 | 847-148 | AF132511 | AF132511 | 24004 | incomplete sequence |
| Apyrase | *Lutzomyia longipalpis* | Ecto ADPase, inhibitor of platelet aggregation | LJL23 | LLOJ003550-RA | 72742-72670 71469-71200 69595-68745 67006-66137 | Scaffold 210 | 72742-66137 | AF131933 | AF131933 | 24025 | |
| Maxadilan | *Lutzomyia longipalpis* | Vasodilator | | LLOJ003615-RA | 31852-33568 34570-34853 35764-36409 36476-37178 37718-37935 39465-39748 39810-40041 40693-41301 41368-42084 42604-42857 43094-44245 | Scaffold 215 | 31852-44245 | | | 23903 | |
| Anti-coagulant | *Lutzomyia longipalpis* | Anti-coagulant, inhibitor of factor Xa | LJL143 | LLOJ001514-RA | 15179-15334 15396-16256151 38-15334 15399-16281 | Scaffold 1307 | 15179-16281 | AY445936 | AY445936 | 23906 | |

| | | | | | | | | | | | |
|---|---|---|---|---|---|---|---|---|---|---|---|
| Endonuclease | *Lutzomyia longipalpis* | DNAse activity | LJL138 | LLOJ005361-RA | 36116-36121 36388-36886 37738-38454 40419-40156 40418-40546 42763-43122 | Scaffold 34 | 36116-43122 | AY455916 | AY455916 | 23998 | |
| 10kDa protein | *Lutzomyia longipalpis* | unknown | LJM19 | LLOJ006680-RA | 44033-43980 43907-43785 42783-42761 42667-42596 42502-42389 39744-39616 38317-38204 38111-37873 | Scaffold 486 | 44033-37873 | AY438271 | AY438271 | 23999 | missing the 3' part |
| 10kDa protein | *Lutzomyia longipalpis* | unknown | LJS192 | LLOJ006680-RA | 44033-43980 43907-43785 42783-42761 42667-42596 42502-42389 39744-39616 38317-38204 38111-37873 | Scaffold 486 | 44033-37873 | AY438270 | AY438270 | 24000 | |

| | | | | | | | | | | |
|---|---|---|---|---|---|---|---|---|---|---|
| 10kDa protein | *Lutzomyia longipalpis* | unknown | LJS169 | LLOJ005853-RA | 6728-6677 6614-6492 6411-6235 5493-5446 5036-4854 | Scaffold 388 | 6728-4854 | AY455912 | AY455912 | 24001 |
| C-type lectin | *Lutzomyia longipalpis* | unknown | LJL91 | LLOJ004397-RA | 356369-356164 356092-355751 353035-352898 352822-352358 | Scaffold 26 | 356369-352358 | AY445934 | AY445934 | 24005 |
| C-type lectin | *Lutzomyia longipalpis* | unknown | LJL15 | LLOJ004397-RA | 356369-356164 356092-355751 353035-352898 352822-352358 | Scaffold 26 | 356369-352358 | DQ190946 | DQ190946 | 24009 |
| C-type lectin | *Lutzomyia longipalpis* | unknown | LJL18 | LLOJ004397-RA | 356369-356164 356092-355751 353035-352898 352822-352358 | Scaffold 26 | 356369-352358 | DQ190947 | DQ190947 | 24012 |
| C-type lectin | *Lutzomyia longipalpis* | unknown | LJM10 | LLOJ004397-RA | 356369-356164 356092-355751 353035-352898 352822-352358 | Scaffold 26 | 356369-352358 | DQ192486 | DQ192486 | 24013 |

| | | | | | | | | | | |
|---|---|---|---|---|---|---|---|---|---|---|
| C-type lectin | *Lutzomyia longipalpis* | unknown | LJS142 | LLOJ004397-RA | 356369-356164<br>356092-355751<br>353035-352898<br>352822-352358 | Scaffold 26 | 356369-352358 | DQ192487 | DQ192487 | 24022 |
| | *Lutzomyia longipalpis* | | LJM06 | LLOJ004398-RA | 36650-363814<br>363139-363561 | Scaffold 26 | 36650-363561 | AY453401 | AY453401 | |
| | *Lutzomyia longipalpis* | | LJL17 | LLOJ007904-RA | 49522-50177<br>50373-50634<br>51134-51490 | Scaffold629 | 49522-51490 | AY452695 | AY452695 | |
| | *Lutzomyia longipalpis* | | 10.7 kDa | LLOJ006680-RA | 44033-43980<br>43907-42785<br>42783-42761<br>42677-42596<br>42502-42389<br>39744-39616<br>38317-38204<br>38111-37873 | Scaffold486 | 44033-37873 | AY438271 | AY438271 | |
| | *Lutzomyia longipalpis* | | 2.5 kDa | LLOJ008391-RA | 36315-36279<br>35981-35780<br>34562-34326<br>32109-31893<br>31792-30773 | Scaffold70 | 36315-30773 | AY438269 | AY438269 | |

| | | | | | | | |
|---|---|---|---|---|---|---|---|
| *Lutzomyia longipalpis* | 16.1 kDa | LLOJ006038-RA | 84373-84706 84773-85424 | Scaffold401 | 84373-85424 | AY455917 | AY455917 |
| *Lutzomyia longipalpis* | 49 kDa | LLOJ008570-RA | 135-576 696-880 954-1439 | Scaffold7245 | 135-1439 | AY455913 | AY455913 |
| *Lutzomyia longipalpis* | 6 kDa | LLOJ008400-RA | 163486-163-269 163193-162902 | Scaffold70 | 163486-162902 | AY455915 | AY455915 |
| *Lutzomyia longipalpis* | 15 kDa | LLOJ003729-RA | 84852-84946 85023-85644 | Scaffold220 | 84852-85644 | AY455914 | AY455914 |
| *Lutzomyia longipalpis* | AMY | LLOJ005909-RA | 3590-4227 4361-5055 5377-5528 5823-6340 | Scaffold244 | 3590-6340 | AF132512 | AY455908 |
| *Lutzomyia longipalpis* | ADA | LLOJ004125-RA | 174920-174958 `75539-175613 `75688-`76494 177893-178240 178336-181149 | Scaffold396 | 174920-181149 | AF234182 | AF234182 |
| *Lutzomyia longipalpis* | HYAL | LLOJ000009-RA | 128970-129862 131827-132288 134370-135216 135303-135747 | Scaffold128 | 128970-136484 | AF132515 | AF132515 |

| | | | | | | | |
|---|---|---|---|---|---|---|---|
| | | | 135943-136484 | | | | |
| *Lutzomyia longipalpis* | NTFE-2 | LLOJ009119-RA | 95030-94786<br>94716-94038<br>93961-93545<br>87557-87292<br>87088-86545<br>86469-85798<br>84884-84538 | Scaffold805 | 95030-84538 | | AF132510 |
| *Phlebotomus papatasi* | PPSP2.5 | | 19574-19764<br>18976-19373<br>18283-18571 | Scaffold2250 incomplete sequence | 19574-18571 | | JQ988875 |
| *Phlebotomus papatasi* | PPSP12 | | 46-107/2198-2341<br>1821-2130 | Scaffold55921/Scaffold62574 | Scaffold55921:46-107/Scaffold62574:1821-2341 | AF335485 | JQ988874 |
| *Phlebotomus papatasi* | PPSP14.2b | | 572-628<br>699-842<br>902-1200 | Scaffold75296 | 572-1200 | | |
| *Phlebotomus papatasi* | PPSP14.3 | | 6633-6690<br>2751-2897<br>2379-2463<br>2000-2111/14293-14562 | Scaffold48699/Scaffold1426 | Scaffold48699:6633-2111/Scaffold1426:14293-14562 | | |

| | | | | | | | | | | |
|---|---|---|---|---|---|---|---|---|---|---|
| | *Phlebotomus papatasi* | | PPSP15 | | 1589-1645/198-642 | Scaffold5990/Scaffold27678 | Scaffold5990:1589-1645/Scaffold27678:198-642 | | JQ988879 | |
| | *Phlebotomus papatasi* | | PPSP28 | PPAI003080-RA | 100284-100217 100154-99762 99695-99288 | Scaffold209 | 100284-99288 | AF335488 | AF335488 | |
| Yellow related proteins | *Phlebotomus papatasi* | Biogenic amine binding proteins | PPTSP42 | PPAI000161-RA | 38501-39937 | Scaffold1012 | 38501-39937 | JQ988885 | JQ988885.1 | 27064 |
| Yellow related proteins | *Phlebotomus papatasi* | Biogenic amine binding proteins | PPTSP44 | PPAI002956-RA | 35513-40323 | Scaffold203 | 35513-40323 | JQ988886 | JQ988886.1 | 22029 |
| D7 related protein | *Phlebotomus papatasi* | unknown | PPTSP30 | PPAI003079-RA | 91520-92491 | Scaffold209 | 91520-92491 | JQ988884 | JQ988884.1 | 22030 |
| Silk related/collagen binding/32kDa protein | *Phlebotomus papatasi* | unknown | PPTSP32 | PPAI002597-RA | 111838-113971 | Scaffold189 | 111838-113971 | JQ988888 | JQ988888 | 27065 |
| Antigen-5 protein | *Phlebotomus papatasi* | unknown | PPTSP29 | PPAI008255-RA | 1140-4960 | Scaffold4853 | 1140-4960 | JQ988887 | JQ988887.1 | 22031 |
| APYRASE | *Phlebotomus papatasi* | Ecto ADPase, inhibitor of platelet aggregation | PPTSP36 | PPAI002442-RA | 15173-17739 | Scaffold1814 | 15173-17739 | JQ988892 | JQ988892 | 2070 |
| | *Phlebotomus papatasi* | | PPTSP34 | | 727-891 949-1889 | Scaffold47164 | 727-1889 | JQ988889 | JQ988889 | |
| SP56.6 | *Phlebotomus papatasi* | unknown | PPTSP56 | PPAI005803-RA | 43568-60885 | Scaffold362 | 43568-60885 | JQ988890 | JQ988890 | 27068 |
| APHA-AMYLASE | *Phlebotomus papatasi* | putatice amylase | PPTAMY | PPAI010151-RA | 134306-142939 | Scaffold72 | 134306-142939 | JQ988891 | JQ988891 | 27069 |

1996

**S13 Table.** Peptidase annotation.

| Peptidase group | Fam. | PFAM | Short Description | Lu. lon | Ph. pap | Aed. aeg. | Anp. gam | Cul. qui | Dro mel | Dro pse | Glo mor |
|---|---|---|---|---|---|---|---|---|---|---|---|
| | A01 | PF00026 | Eukaryotic aspartyl protease | 4 | 2 | 1 | 1 | 1 | 13 | 13 | 3 |
| | A22 | PF01080 | Presenilin | 1 | 1 | 3 | 3 | 4 | 3 | 3 | 3 |
| Aspartic | A25 | PF03418 | Germination protease | 1 | 0 | 0 | 0 | 0 | 0 | 0 | 0 |
| | A28 | PF09668 | Aspartate protease | 0 | 1 | 1 | 1 | 0 | 1 | 1 | 0 |
| | A33 | - | skin SASPase | 0 | 1 | 0 | 0 | 0 | 0 | 0 | 0 |
| | C01 | PF00112 | Papain family cysteine protease | 5 | 8 | 15 | 13 | 19 | 15 | 12 | 10 |
| | C02 | PF00648 | Calpain family cysteine protease | 5 | 4 | 9 | 20 | 7 | 4 | 3 | 3 |
| | C12 | PF01088 | Ubiquitin carboxyl-terminal hydrolase, family 1 | 2 | 3 | 3 | 3 | 3 | 4 | 4 | 4 |
| | C13 | PF01650 | Peptidase C13 family | 1 | 1 | 1 | 1 | 1 | 1 | 1 | 1 |
| | C14 | PF00656 | Caspase domain | 2 | 5 | 9 | 16 | 13 | 7 | 7 | 6 |
| | C15 | PF01470 | Pyroglutamyl peptidase | 0 | 1 | 1 | 1 | 1 | 1 | 1 | 1 |
| | C19 | PF00443 | Ubiquitin carboxyl-terminal hydrolase | 18 | 17 | 21 | 39 | 29 | 23 | 18 | 24 |
| | C26 | PF07722 | Peptidase C26 | 2 | 1 | 4 | 5 | 4 | 8 | 6 | 7 |
| | C44 | PF00310 | Glutamine amidotransferases class-II | 3 | 1 | 5 | 4 | 5 | 7 | 6 | 7 |
| Cysteine | C46 | PF01079 | Hint module | 1 | 0 | 1 | 1 | 1 | 1 | 1 | 1 |
| | C48 | PF02902 | Ulp1 protease family, C-terminal catalytic domain | 3 | 3 | 3 | 4 | 3 | 7 | 5 | 9 |
| | C54 | PF03416 | Peptidase family C54 | 2 | 3 | 2 | 2 | 2 | 2 | 2 | 2 |
| | C56 | PF01965 | DJ-1/PfpI family | 2 | 1 | 1 | 2 | 1 | 2 | 1 | 5 |
| | C64/C85 | PF02338 | OTU-like cysteine protease | 5 | 3 | 4 | 5 | 4 | 7 | 6 | 4 |
| | C65 | PF10275 | Peptidase C65 Otubain | 1 | 0 | 1 | 1 | 1 | 2 | 1 | 1 |
| | C67 | - | CylD peptidase" | 1 | 1 | 1 | 1 | 1 | 0 | 1 | 1 |
| | C78 | PF07910 | Peptidase family C78 | 3 | 2 | 2 | 2 | 2 | 2 | 2 | 2 |
| | C86 | PF02099 | Josephin | 1 | 1 | 0 | 1 | 1 | 1 | 2 | 1 |
| | C95 | - | lysosomal 66.3 kDa protein | 1 | 1 | 0 | 0 | 0 | 0 | 0 | 0 |
| | C97 | PF05903 | PPPDE putative peptidase domain | 2 | 2 | 2 | 1 | 2 | 2 | 1 | 1 |
| | M01 | PF01433 | Peptidase family M1 | 17 | 22 | 27 | 39 | 23 | 23 | 24 | 14 |
| | M02 | PF01401 | Angiotensin-converting enzyme | 5 | 5 | 8 | 11 | 9 | 6 | 7 | 7 |
| | M03 | PF01432 | Peptidase family M3 | 2 | 2 | 2 | 2 | 2 | 2 | 2 | 2 |
| | M08 | PF01457 | Leishmanolysin | 1 | 1 | 1 | 1 | 1 | 1 | 1 | 1 |
| | M10 | PF00413 | Matrixin | 3 | 3 | 8 | 4 | 9 | 3 | 2 | 3 |
| | M12B | PF01421 | Reprolysin (M12B) family zinc metalloprotease | 5 | 4 | 7 | 13 | 11 | 14 | 7 | 11 |
| | M13 | PF05649 | Peptidase family M13 | 6 | 5 | 9 | 9 | 9 | 30 | 27 | 25 |
| | M14 | PF00246 | Zinc carboxypeptidase | 20 | 26 | 24 | 38 | 30 | 32 | 28 | 28 |
| | M16 | PF00675 | Insulinase (Peptidase family M16) | 5 | 4 | 10 | 13 | 12 | 12 | 16 | 15 |
| | M17 | PF00883 | Cytosol aminopeptidase family, catalytic domain | 6 | 6 | 5 | 5 | 6 | 9 | 13 | 11 |
| Metallo | M19 | PF01244 | Membrane dipeptidase (Peptidase family M19) | 5 | 6 | 7 | 6 | 4 | 6 | 4 | 4 |
| | M20 | PF01546 | Peptidase family M20/M25/M40 | 6 | 3 | 1 | 2 | 0 | 7 | 0 | 4 |
| | M22 | PF00814 | Glycoprotease family | 2 | 1 | 2 | 2 | 2 | 2 | 4 | 4 |
| | M23 | PF01551 | Peptidase family M23 | 0 | 0 | 0 | 1 | 0 | 0 | 0 | 0 |
| | M24 | PF00557 | Metallopeptidase family M24 | 8 | 7 | 11 | 16 | 8 | 11 | 6 | 12 |
| | M28 | PF04389 | Peptidase family M28 | 5 | 6 | 3 | 13 | 5 | 15 | 9 | 9 |
| | M38 | PF01979 | Amidohydrolase family | 4 | 4 | 0 | 1 | 1 | 4 | 1 | 6 |
| | M41 | PF01434 | Peptidase family M41 | 3 | 1 | 3 | 3 | 3 | 4 | 3 | 4 |
| | M48 | PF01435 | Peptidase family M48 | 1 | 1 | 1 | 1 | 1 | 4 | 4 | 1 |
| | M49 | PF03571 | Peptidase family M49 | 1 | 0 | 0 | 1 | 1 | 1 | 1 | 1 |
| | M67 | PF01398 | JAB1/Mov34/MPN/PAD-1 ubiquitin protease | 5 | 5 | 5 | 7 | 4 | 8 | 4 | 10 |

| | | | | | | | | | | |
|---|---|---|---|---|---|---|---|---|---|---|
| | M76 | PF09768 | Peptidase M76 family | 1 | 1 | 1 | 1 | 1 | 1 | 1 | 1 |
| | M79 | PF02517 | CAAX protease self-immunity | 1 | 1 | 1 | 1 | 1 | 1 | 1 | 1 |
| | M87 | PF08434 | Chloride channel accessory protein | 1 | 1 | 0 | 0 | 0 | 0 | 0 | 0 |
| Serine | S01 | PF00089 | Trypsin | 112 | 126 | 370 | 560 | 428 | 306 | 215 | 221 |
| | S08 | PF00082 | Subtilase family | 6 | 4 | 6 | 5 | 5 | 6 | 5 | 5 |
| | S09 | PF00326 | Prolyl oligopeptidase family | 7 | 6 | 10 | 18 | 13 | 45 | 11 | 52 |
| | S10 | PF00450 | Serine carboxypeptidase | 5 | 3 | 5 | 4 | 5 | 5 | 6 | 4 |
| | S14 | PF00574 | Clp protease | 1 | 1 | 1 | 2 | 1 | 1 | 1 | 3 |
| | S16 | PF05362 | Lon protease (S16) C-terminal proteolytic domain | 1 | 0 | 1 | 1 | 1 | 1 | 1 | 3 |
| | S24 | PF00717 | Peptidase S24 | 2 | 1 | 0 | 0 | 0 | 0 | 0 | 0 |
| | S28 | PF05577 | Serine carboxypeptidase S28 | 7 | 5 | 8 | 10 | 9 | 6 | 7 | 2 |
| | S33 | PF12697 | Alpha/beta hydrolase family | 24 | 17 | 11 | 24 | 11 | 25 | 9 | 13 |
| | S54 | PF01694 | Rhomboid family | 4 | 4 | 3 | 5 | 3 | 9 | 6 | 7 |
| | S59 | PF04096 | Nucleoporin autopeptidase | 1 | 1 | 1 | 1 | 1 | 1 | 1 | 1 |
| | S60 | PF00405 | Transferrin | 4 | 3 | 6 | 7 | 9 | 5 | 6 | 5 |
| | S63 | - | EGF-like module containing mucin-like hormone receptor-like 2 | 1 | 1 | 1 | 1 | 1 | 4 | 1 | 1 |
| | S72 | PF05454 | Dystroglycan (Dystrophin-associated glycoprotein 1) | 1 | 1 | 2 | 2 | 2 | 2 | 2 | 2 |
| | S81 | PF05497 | Destabilase | 1 | 1 | 2 | 2 | 2 | 7 | 3 | 1 |
| Threonine | T01 | PF00227 | Proteasome subunit | 12 | 13 | 15 | 21 | 18 | 29 | 37 | 21 |
| | T02 | PF01112 | Asparaginase | 3 | 4 | 2 | 3 | 5 | 5 | 4 | 6 |
| | T03 | PF01019 | Gamma-glutamyltranspeptidase | 2 | 3 | 6 | 3 | 5 | 4 | 4 | 6 |
| | T06 | - | polycystin-1 | 1 | 0 | 0 | 0 | 0 | 0 | 0 | 0 |
| | Total[a] | | | 376 | 376 | 703 | 983 | 800 | 808 | 586 | 646 |

**S14 Table.** Glycosidase Hydrolase family 13 annotation.

| Species | Gene | Transcri pt | Scaffol d # | Base pair | Description | Gene status | Justification | | Obs | Signal Peptide |
|---|---|---|---|---|---|---|---|---|---|---|
| *Lu. longipal* | LLONA MY1 | LLOJ000 566-RA | Scaffol d1090 | 51770- 53700 | Alpha-amylase | *Truncated 5' | Similarity with XP_001985943.1 and | Rev erse | | No |
| | LLONA MY2 | LLOJ004 841-RA | Scaffol d296 | 13595- 15335 | Alpha-amylase | Complete | Similarity with AAO17923.2 and AAA92243.1 | Fow ard | | Yes |
| | LLONA MY3 | LLOJ002 257-RA | Scaffol d1546 | 27920- 39494 | Alpha-amylase | Complete | Similarity with XP_001660907.1 and | Fow ard | | Yes |
| | LLONA MY4 | LLOJ004 838-RA | Scaffol d296 | 160- 11278 | Alpha-amylase | Complete with 3' domain duplicated | Similarity with XP_005185446.1 and | Rev erse | Equal to LLOTMP0048 | Yes |
| | LLONA MY5 | LLOJ004 839-RA | Scaffol d296 | 2784- 6233 | Alpha-amylase | Complete | Similarity with XP_004529971.1 and | Fow ard | Equal to LLOTMP0048 | Yes |
| | LLONA MY6 | LLOJ004 880-RA | Scaffol d3 | 46114- 56430 | Alpha-amylase | 2 genes ( 1 complete plus 1 truncated 3') | Similarity with EFN63697.1 and AAA92237.1 | Rev erse | | Yes both |
| | LLONA MY7 | LLOJ004 881-RA | Scaffol d3 | 52460- 63531 | Alpha-amylase | 2 genes (1 gene truncado 3' e 1 completo) | Similarity with XP_005185446.1 and | Fow ard | | Yes both |
| | LLONA MY8 | LLOJ004 882-RA | Scaffol d3 | 63145- 64814 | Alpha-amylase | Complete | Similarity with AAO13293.1 and BAB32521.1 | Rev erse | | Yes |
| | LLONA MY9 | LLOJ005 909-RA | Scaffol d396 | 3590- 7348 | Alpha-amylase | Complete with 5' domain duplicated | Similarity with AAD32192.1 and AAA92235.1 | Fow ard | | Yes |
| | LLONC D98 | LLOJ006 803-RA | Scaffol d5 | 479539- 487315 | CD98hc | Truncated 5' | Similarity with XP_001655437.1 and | Fow ard | | No |
| | LLONA MY10 | LLOJ008 156-RA | Scaffol d6719 | 3215- 5732 | Alpha-amylase | Truncated 3' | Similarity with XP_002063229.1 and | Rev erse | | Yes |
| | LLONA MY11 | LLOJ008 629-RA | Scaffol d732 | 27159- 32069 | Alpha-amylase?? | Truncated 5' | Similarity with XP_001659278.1 and | Fow ard | Bad Sequence | No |
| | LLONB RA1 | LLOJ005 533-RA | Scaffol d352 | 48422- 60071 | 1,4-Alpha -glucan branching enzyme | Truncated 5' | Similarity with XP_002006790.1 and | Fow ard | | No |
| *P. papatasi* | PPATA MY1 | PPAI0107 60-RA | Scaffol d875 | 29555- 40817 | Alpha-amylase | 2 genes duplicated at beginning – First with signal peptide | Similarity with XP_001649787.1 and | Rev erse | | |
| | PPATA MY2 | PPAI0102 58-RA | Scaffol d74 | 175129- 183344 | Alpha-amylase | Complete with Signal peptide | Similarity with XP_001660906.1 and | Rev erse | | |
| | PPATA MY3 | PPAI0102 57-RA | Scaffol d74 | 163348- 174909 | Alpha-amylase | Complete with Signal peptide | Similarity with XP_001660909.1 and | Rev erse | | |
| | PPATA MY4 | PPAI0102 56-RA | Scaffol d74 | 153174- 163056 | Alpha-amylase | Complete with Signal peptide | Similarity with XP_001660907.1 and | Rev erse | | |
| | PPATA MY5 | PPAI0102 55-RA | Scaffol d74 | 146430- 148321 | Alpha-amylase | Complete with Signal peptide | Similarity with XP_001660908.1 and | Rev erse | | |
| | PPATA MY6 | PPAI0107 59-RA | Scaffol d875 | 17428- 22094 | Alpha-amylase – trealose | Complete with Signal peptide | Similarity with XP_005186326.1 and | Rev erse | | |
| | PPATA MY7 | PPAI0101 50-RA | Scaffol d72 | 118446- 126822 | Alpha-amylase | 2 genes | Similarity with EFN63697.1 and AAA92235.1 | Rev erse | | |
| | PPATA MY8 | PPAI0101 51-RA | Scaffol d72 | 134306- 142939 | Alpha-amylase | 2 genes - 1 with signal peptide and 1 complete | Similarity with EFN63697.1 and BAB32543.1 | Rev erse | | |
| | PPATA MY9 | PPAI0101 48-RA | Scaffol d72 | 104696- 116160 | Alpha-amylase | Complete | Similarity with AAO14612.1 and AAA92237.1 | For ward | | |
| | | PPAI0017 89-RA | Scaffol d1540 | 25221- 30557 | Alpha-amylase | Truncated 5' and 3' | Similarity with XP_001659278.1 | Rev erse | | |
| | PPATB RA1 | PPAI0089 63-RA | Scaffol d5325 | 3184- 6883 | 1,4-Alpha -glucan branching enzyme | Truncated 5' | Similarity with XP_001959638.1 and | Rev erse | | |
| | | PPAI0048 54-RA | Scaffol d2979 | 7708- 9147 | CD98hc | Truncated 5' and 3' | Similarity with XP_001655437.1 and | For ward | | |

**S15 Table.** Chitinase family annotation.

| Species | Name/description | Transcript | Scaffold | Base Pair Range | signal P | catalitic domain | CBD | length Aa |
|---|---|---|---|---|---|---|---|---|
| *Lu. Longipalpis* | Llcht7 | LLOJ009389-RA | Scaffold864 | 5025-6367 | yes | 2 | 1 | 1006 |
| | cht12 like protein | LLOJ003713-RA | Scaffold22 | 171881-176204 | yes | 1 | 0 | 389 |
| | LlIDGF4 | LLOJ008789-RA | Scaffold76 | 69632-72255 | yes | 1 | 0 | 441 |
| | | LLOJ004095-RA | Scaffold2438 | 22-1333 | | | | |
| | Llcht1/midgut cht | LLOJ000006-RA | Scaffold1 | 63082-67552 | yes | 1 | 1 | 473 |
| | Llcht5 | LLOJ008303-RA | Scaffold69 | 56110-62401 | yes | 1 | 1 | 576 |
| | chitinase like protein | LLOJ009205-RA | Scaffold822 | 59819-62113 | yes | 1 | 0 | 392 |
| | Llcht8 | LLOJ003119-RA | Scaffold1934 | 2910-4887 | yes | 1 | 0 | 410 |
| | chitinase like protein | LLOJ006041-RA | Scaffold401 | 99424-101989 | yes | 1 | 1 | 491 |
| | Llcht10 | LLOJ006362-RA | Scaffold44 | 131615-140847 | yes | 3 | 3 | 1829/partial |
| | Llcht2 | LLOJ008646-RA | Scafold736 | 26000-33995 | yes | 1 | 0 | 487 |
| | Llcht6 | | orphan | | yes | 1 | 1 | 747/partial |
| *P. papatasi* | PpCht6 | PPAI000832-RA | Scaffold121 | 188380-199213 | yes | 1 | 1 | 2324 |
| | chitinase like protein | PPAI000950-RA | Scaffold1249 | 21440-30369 | No | 1 | 0 | 452 |
| | PpCht7 | PPAI001579-RA | Scaffold1465 | 2791-6550 | No | 2 | 1 | 994 |
| | PpCht5 | PPAI002483-RA | Scaffold1836 | 6030-14822 | No | 1 | 1 | 449/partial |
| | PpCht10 | PPAI003548-RA | Scaffold2324 | 14268-22615 | No | 4 | 4 | 1987 |
| | chitinase like protein | PPAI004649-RA | Scaffold2841 | 743-4696 | yes | 1 | 0 | 485 |
| | | PPAI004650-RA | Scaffold2841 | 12580-13845 | | | | |
| | chitinase like protein | PPAI007799-RA | Scaffold4609 | 973-1971 | No | 1 | 0 | 307/partial |
| | chitinase like protein | PPAI008028-RA | Scaffold47091 | 1646-6883 | No | 1 | 0 | 470/partial |
| | PpCht4 | PPAI009441-RA | Scaffold5844 | 565-1174 | No | 1 | 0 | 443/partial |
| | | PPAI002059-RA | Scaffold163 | 158880-15969 | | | | |
| | PpIDGF | PPAI009732-RA | Scaffold63 | 203447-206693 | yes | 1 | 0 | 443 |
| | PpCht1/midgut cht | PPAI009937-RA | Scaffold67 | 133926-135374 | yes | 1 | 0 | 470 |
| | | PPAI009938-RA | Scaffold67 | 138396-144637 | | | | |
| | cht10 like protein (?) | PPAI010017-RA | Scaffold687 | 52518-60360 | No | 3 | 1 | 1118/partial |

**S16 Table.** Hexosaminidase family annotation.

Detailed description of n-acetylhexosaminidases identified in sand flies is shown. Gene names, scaffold whereby genes are mapped, base pair range in the scaffold, number of glycohydro 20 domains, and number of glycohydro 20b2 domains are described.

| Species | Gene | Transcript | Scaffold # | Base pair range | Glycohydro 20 | Glycohydro 20b2 |
|---|---|---|---|---|---|---|
| *Lu. longipalpis* | LuloNAG1 | LLOJ001716-RA | Scaffold137 | 132443-137441 | 1 | 1 |
| | LuloNGA2 | LLOJ002196-RA | Scaffold1527 | 14092-18453 | 1 | 1 |
| | LuloNGA3 | LLOJ002001-RA | Scaffold146 | 190122-196971 | 1 | 1 |
| | LuloFDL | LLOJ004141-RA | Scaffold245 | 41024-50912 | 1 | 1 |
| | LuloHEX | LLOJ003464-RA | Scaffold207 | 36386-43078 | 1 | 1 |
| *P. papatasi* | PpNAG1 | PPAI008055-RA | Scaffold4727 | 385-1581 | 1 | |
| | PpNGA2 | PPAI007424-RA | Scaffold44824 | 2533 8568 | 1 | 1 |
| | PpNGA3 | PPAI003424-RA | Scaffold2268 | 9498-20213 | 1 | 1 |
| | PpFDL | PPAI004287-RA | Scaffold2632 | 2797-9660 | 1 | |
| | PpHEX | PPAI000447-RA | Scaffold110 | 43948 49643 | 1 | 1 |

Number of n-acetylhexosaminidases genes.

| N-acetyl hexosaminidases | Gene | *P. papatasi* | *L. longipalpis* | *A. gambiae* | *A. aegypti* | *D. melanogaster* | *T. castaneum* |
|---|---|---|---|---|---|---|---|
| | NAG | 2 | 2 | 2 | 2 | 2 | 2 |
| | FDL | 1 | 1 | 1 | 1 | 1 | 1 |
| | HEX | 1 | 1 | 1 | 1 | 0 | 3 |
| **Total** | | **4** | **4** | **4** | **4** | **3** | **6** |

**S17 Table.** Chitinase deacetylase family annotation.

Detailed description of chitin deacetylases identified in sand flies is shown. Gene names, scaffold whereby genes are mapped, base pair range in the scaffold, and number of CBDs, LDLa, and CDA domains are described.

| Species | Gene | Transcript | Scaffold # | Base pair range | CBD domain | LDLa domain | CDA domain |
|---|---|---|---|---|---|---|---|
| *Lu. longipalpis* | LuloCDA1 | LLOJ007910-RA | Scaffold63 | 118979-131052 | 1 | 1 | 1 |
| | LuloCDA2 | LLOJ007911-RA | Scaffold63 | 138775-152294 | 1 | 1 | 1 |
| | LuloCDA3 | LLOJ007595-RA | Scaffold593 | 64642-69829 | 1 | 1 | 1 |
| | LuloCDA4 | LLOJ006479-RA | Scaffold461 | 47503-59468 | 2 | | 1 |
| | LuloCDA5 | LLOJ005142-RA | Scaffold322 | 26433-27614 | | | 1 |
| | LuloCDA9 | LLOJ005424-RA | Scaffold34 | 24191-26951 | | | 1 |
| *P. papatasi* | PpCDA1 | PPAI005534-RA | Scaffold34 | 611270-612917 | | | 1 |
| | PpCDA2 | PPAI005535-RA | Scaffold34 | 630278-635209 | 1 | | 1 |
| | PpCDA4 | PPAI009437-RA | Scaffold583 | 16931-26189 | 1 | | 1 |
| | PpCDA5 | PPAI008909-RA | Scaffold53 | 149562-153239 | | | 1 |

Number of chitin deacetylase genes.

| Chitin Deacetylases | Gene | *P. papatasi* | *L. longipalpis* | *A. gambiae* | *D. melanogaster* | *T. castaneum* |
|---|---|---|---|---|---|---|
| Total | CDA | 4 | 6 | 6 | 6 | 9 |

**S18 Table.** Peritrophin family annotation.

Detailed description of peritrophins identified in sand flies are shown. Gene names, scaffold whereby genes are mapped, base pair range in the scaffold, GenBank access number, number and type of CBD domains, number of mucin-like motif, such as S-T tandem repeats or T-P rich regions are described.

| Species | Gene | Transcript | Scaffold # | Base pair range | GenBank (V1.1) accession # | CBD domain | CBD-like domain[1] | Mucin-like domain (S-T tandem repeats) | T-P-rich region |
|---|---|---|---|---|---|---|---|---|---|
| *Lu. longipalpis* | **LuloPer1** | LLOJ006497-RA | Scaffold464 | 51712-53102 | EU124588 | 4 (Type 1) | | | |
| | **LuloPer2** | LLOJ003676-RA | Scaffold217 | 68692-69367 | EU124602 | 1 (Type 1) | | | |
| | **LuloPer3a** | LLOJ002647-RA | Scaffold1711 | 4339-6324 | | 2 (Type 1) | 1 | 1 | |
| | **LuloPer3b** | LLOJ004906-RA | Scaffold3 | 473357-474138 | | | | | |
| | **LuloPer4** | LLOJ003410-RA | Scaffold2040 | 19086-27302 | | 3 (Type 1) | | | |
| | **LuloPer7** | LLOJ002081-RA | Scaffold1493 | 15517-16211 | | 1 (Type 1) | 1 | | |
| | **LuloPer9** | LLOJ006576-RA | Scaffold474 | 3871-6910 | | 2 (Type 1) | 1 | | |
| | **LuloPer10** | LLOJ007615-RA | Scaffold5963 | 2565-6708 | | 3 (Type 1) | | | |
| | **LuloPer11** | LLOJ002676-RA | Scaffold172 | 80007-90140 | | 6 (Type 1) | | | |
| | **LuloPer14** | LLOJ006496-RA | Scaffold464 | 49900-50692 | | 3 (Type 1) | | 1 | |
| | **LuloPer16** | LLOJ000527-RA | Scaffold1082 | 52971-53915 | | 3 (Type 1) | | | 1 |
| | **LuloPer17** | LLOJ006500-RA | Scaffold464 | 60212-64414 | | 4 (Type 1) | | | |
| | **LuloPer18** | LLOJ006499-RA | Scaffold464 | 59214-59897 | | 3 (Type 1) | | | |
| | **LuloPer19** | LLOJ000526-RA | Scaffold1082 | 50302-53130 | | 1 (Type 1) | | | |
| | **LuloPer22** | LLOJ006495-RA | Scaffold872 | 17904-18738 | | 3 (Type 1) | | | 1 |
| | **LuloPer23** | LLOJ009445-RA | Scaffold1082 | 48309-50748 | | 3 (Type 1) | | | |
| | **LuloPer24** | LLOJ000525-RA | Scaffold3 | 362554-365807 | | 2 (Type 1) | | | |
| | **LuloPer25** | LLOJ004900-RA | Scaffold68 | 81157-82300 | | 4 (Type 1) | | | |
| | **LuloPer26** | LLOJ008206-RA | Scaffold924 | 31951-34753 | | 1 (Type 1) | 1 | | |
| | **LuloPer27** | LLOJ008206-RA | Scaffold1260 | 61537-65771 | | 2 (Type 1) | | 1 | |
| | **LuloPer30** | LLOJ001522-RA | Scaffold131 | 39846-45296 | | 2 (Type 1) | 1 | | |
| | **LuloPer32** | LLOJ004516-RA | Scaffold266 | 61643-62145 | EU124607 | 1 (Type 1) | | | |
| *P. papatasi* | **PpPer1** | PPAI009353-RA | Scaffold573 | 55119-56306 | EU031912 | 4 (Type 1) | | | |

| | | | | | | | | | | |
|---|---|---|---|---|---|---|---|---|---|---|
| **PpPer2** | PPAI009723-RA | Scaffold629 | 59171-60434 | EU047543 | 1 (Type 1) | | | | | |
| **PpPer3** | PPAI006556-RA | Scaffold424 | 3891-5068 | EU045354 | 2 (Type 1) | 1 | | 1 | | |
| **PpPer4** | PPAI006974-RA | Scaffold4344 | 8019-8947 | | 3 (Type 1) | | | | | |
| **PpPer5a** | PPAI005563-RA | Scaffold3427 | 793-1617 | | 9 (Type 1) | | | | | |
| **PpPer5b1** | PPAI007266-RA | Scaffold3427 | 2115-3480 | | | | | | | |
| **PpPer5b2** | PPAI007267-RA | Scaffold44207 | 1314-4014 | | | | | | | |
| **PpPer6** | PPAI001604-RA | Scaffold1475 | 37185-37974 | | 2 (Type 1) | | | | | 1 |
| **PpPer7** | PPAI002253-RA | Scaffold1722 | 29083-30398 | | 1 (Type 1) | 1 | | | | |
| **PpPer9** | PPAI007659-RA | Scaffold45559 | 8856-9604 | | 1 (Type 1) | 1 | | | | |
| **PpPer10** | PPAI004716-RA | Scaffold2886 | 9426-15872 | | 3 (Type 1) | | | | | |
| **PpPer11** | PPAI004749-RA | Scaffold29 | 534135-536847 | | 3 (Type 1) | | | | | |
| **PpPer12** | PPAI001263-RA | Scaffold134 | 131298-133319 | | 3 (Type 1) | | | | | 1 |
| **PpPer13** | PPAI004750-RA | Scaffold29 | 547260-548379 | | 3 (Type 1) | | | | | |
| **PpPer14** | PPAI009350-RA | Scaffold573 | 45836-46651 | | 3 (Type 1) | | | | | |
| **PpPer15** | PPAI009351-RA | Scaffold573 | 48583-49534 | | 3 (Type 1) | | | | | 1 |
| **PpPer16** | PPAI003170-RA | Scaffold2120 | 16177-17202 | | 3 (Type 1) | | | | | 1 |
| **PpPer17** | PPAI009356-RA | Scaffold573 | 60025-61162 | | 4 (Type 1) | | | | | |
| **PpPer18** | PPAI009355-RA | Scaffold573 | 58812-59940 | | 3 (Type 1) | | | 1 | | |
| **PpPer19** | PPAI003168-RA | Scaffold2120 | 9508-10552 | | 1 (Type 1) | | | | | |
| **PpPer20** | PPAI003169-RA | Scaffold2120 | 14138-14970 | | 3 (Type 1) | | | | | |
| **PpPer21** | PPAI000790-RA | Scaffold12 | 960694-984053 | | 5 (Type 1) | | | | | |
| **PpPer22** | PPAI009352-RA | Scaffold573 | 51069-52817 | | 4 (Type 1) | | | 1 | | |
| **PpPer23** | PPAI003166-RA | Scaffold2120 | 2504-3621 | | 3 (Type 1) | | | | | |
| **PpPer24** | PPAI003167-RA | Scaffold2120 | 3989-5043 | | 2 (Type 1) | | | | | |
| **PpPer25** | PPAI010501-RA | Scaffold8 | 109637-110161 | | 2 (Type 1) | | | | | 1 |
| **PpPer26** | PPAI004431-RA | Scaffold270 | 94310-95337 | | | 1 | | | | |
| **PpPer27** | PPAI008214-RA | Scaffold482 | 18787-23903 | | 2 (Type 1) | | | 1 | | |
| **PpPer28** | PPAI001796-RA | Scaffold1547 | 1805-3212 | | 1 (Type 1) | | | | | |
| **PpPer29** | PPAI002251-RA | Scaffold1722 | 19175-20095 | | 1 (Type 1) | 1 | | | | |

| | | | | | |
|---|---|---|---|---|---|
| **PpPer30** | PPAI002252-RA | Scaffold1722 | 23947-24787 | 1 (Type 1) | 1 |

Number of peritrophin genes.

| **Peritrophins** | **Gene** | ***P. papatasi*** | ***L. longipalpis*** |
|---|---|---|---|
| | Per1 | 1 | 1 |
| | Per2 | 1 | 1 |
| | Per3 | 1 | 2 |
| | Per4 | 1 | 1 |
| | Per5 | 2 | 0 |
| | Per6 | 1 | 0 |
| | Per7 | 1 | 1 |
| | Per9 | 1 | 1 |
| | Per10 | 1 | 1 |
| | Per11 | 1 | 1 |
| | Per12 | 1 | 0 |
| | Per13 | 1 | 0 |
| | Per14 | 1 | 1 |
| | Per15 | 1 | 0 |
| | Per16 | 1 | 1 |
| | Per17 | 1 | 1 |
| | Per18 | 1 | 1 |
| | Per19 | 1 | 1 |
| | Per20 | 1 | 0 |
| | Per21 | 1 | 0 |
| | Per22 | 1 | 1 |
| | Per23 | 1 | 1 |
| | Per24 | 1 | 1 |
| | Per25 | 1 | 1 |
| | Per26 | 1 | 1 |
| | Per27 | 1 | 1 |
| | Per28 | 1 | 0 |
| | Per29 | 1 | 0 |
| | Per30 | 1 | 1 |
| | Per32 | 0 | 1 |
| **Total** | | **30** | **22** |

**S19 Table.** Aquoporin family annotation.

| Species | Gene ID | Symbol | *D. melanogaster* | *Tribolium* | *P. humanus* | *Glossina* | *Aedes* | Sand fly |
|---|---|---|---|---|---|---|---|---|
| *P. papatasi* | PPAI004113 | Drip | FBpp0303806 | XP_972862 | PHUM195520 | GMOY003126 | AAEL003512 | LLOJ003011 |
| | PPAI003937 | AQP2 | FBpp0087236 | XP_008190404 | PHUM195520 | GMOY012008 | AAEL003550 | LLOJ009246 |
| | PPAI003310 | Bib | FBpp0079519 | XP_968782 | PHUM617300 | GMOY012056 | AAEL004741 | LLOJ003484 |
| | PPAI002727 | AQP4/AQP5 | FBpp0072016 | XP_008193675 | PHUM474700 | GMOY009435 | AAEL005008 | LLOJ004483 |
| | PPAI003855 | AQP4/AQP5 | FBpp0072016 | XP_008193675 | PHUM369010 | GMOY009435 | AAEL005008 | LLOJ004483 |
| | PPAI001626 | AQP6 | FBpp0289514 | XP_008196202 | PHUM127940 | GMOY006916 | AAEL014255 | LLOJ001619 |
| *Lu. longipalpis* | LLOJ003011 | Drip | FBpp0303806 | XP_972862 | PHUM195520 | GMOY003126 | AAEL003512 | PPAI004113 |
| | LLOJ009245 | AQP2 | FBpp0087236 | XP_008190404 | PHUM434530 | GMOY012008 | AAEL003550 | PPAI004113 |
| | LLOJ003484 | Bib | FBpp0079519 | XP_968782 | PHUM617300 | GMOY012056 | AAEL004741 | PPAI003310 |
| | LLOJ004483 | AQP4/AQP5/Eglp | FBpp0072013 | XP_008193675 | PHUM474700 | GMOY009435 | AAEL005008 | PPAI003855 |
| | LLOJ002056 | AQP4/AQP5/Eglp | FBpp0297476 | XP_008200635 | PHUM474700 | GMOY009435 | AAEL005001 | PPAI003855 |
| | LLOJ001619 | AQP6 | FBpp0289514 | XP_008196202 | PHUM127940 | GMOY009439 | AAEL014255 | PPAI001626 |

**S20 Table. Details of *Lutzomyia longipalpis* circadian and behavior genes and proteins.** Columns: Gene – the assigned gene and protein name (NTE – N-terminus missing, CTE – C-terminus missing, INT –problems in the assembly, FUS – two gene models located in the same scaffold were fused; JOI – gene model spans scaffolds); OGS – the official gene number in the 10,429 genes in LlonJ1.1, prefix is LLOJ; Scaffold (Sc) – the LlonJ1.1 genome assembly supercontig ID and Contig (Ct) – the LlongJ1 genome assembly contig ID, prefix is Scaffold; Coordinates – the nucleotide range from the first position of the start codon to the last position of the stop codon in the scaffold; Strand + is forward and - is reverse; Introns – number of introns; AAs – number of encoded amino acids in the protein; Comments – comments on the OGS gene model and repairs to be done in the genome assembly.

| Gene | OGS | Scaffold/Contig | Coordinates | Strand | Introns | AAs | Comments |
|---|---|---|---|---|---|---|---|
| ***tim*-JOI** | - | Sc166 and Sc329 | 65630-74870 and 106250-107029 | - ; + | 12 | 1065 | New gene model. LLOTMP002513 (Sc166) and LLOTMP005222 (Sc329) were fused and edited |
| ***tim2*** | 2988 | Sc188 | 69219-79438 | + | 9 | 1031 | Fine as it is |
| ***per*-JOI** | - | Ct49844 and Sc1003 | 963-1046 and 7944-14194 | - ; + | 7 | 1169 | New gene model. First exon of the model was localized on Contig49844 (LlonJ1 assembly). The rest of the model was in LLOTMP000134 (Sc1003). |
| **cwo** | 7025 | Sc520 | 47143-50439 | - | 5 | 923 | Fine as it is |
| ***Clk*-JOI** | - | Sc48 and Sc108 | 204225-210403 and 105635-106138 | + | 7 | 696 | New gene model. LLOTMP006614 (Sc 48) and LLOTMP000502 (Sc108) were fused and edited |
| **cyc** | 264 | Sc103 | 104825-112093 | + | 7 | 622 | Exons 1 to 4 were removed and initial methionine was properly fixed |
| ***sim*-NTE** | ***798*** | Sc1131 | 44553-56676 | + | 7 | 784 | NTE region is missed |
| **tgo** | 3291 | Sc2 | 436819-448928 | + | 8 | 621 | Final part of the model was edited |
| ***sgg*-CTE** | ***6077*** | Sc408 | 69292-72843 | + | 1 | 122 | CTE region is missed |
| **dbt** | 4904 | Sc3 | 424178-439399 | + | 6 | 359 | Fine as it is |
| ***CKII alpha*** | ***4026*** | 2394 | 1353-2369 | + | 0 | 338 | Fine as it is |
| **CKII beta-JOI** | - | Sc2377 and Sc68 | 11469-13761 and 23168-25415 | + | 5 | 277 | New gene model. LLOTMP004003 (Sc2377) and LLOTMP008200 (Sc68) were fused |
| ***Pp1a*** | - | - | - | - | - | 162 | Present only in the larval L4 transcriptome assembly |
| **Pp1beta** | 6762 | Sc5 | 106939-123172 | - | 5 | 332 | Fine as it is |
| ***Pp2a*** | ***8900*** | Sc778 | 60175-68895 | - | 4 | 309 | First exon was extend and four last exons were excluded |
| **Pp4** | 3815 | Sc227 | 34843-37174 | - | 3 | 307 | Fine as it is |

| | | | | | | | |
|---|---|---|---|---|---|---|---|
| ***PpV-6*** | ***10016*** | Sc979 | 48828-51945 | - | 2 | 302 | Maintained only the first three exons of the original model. Third exon extended |
| **Pp7-FUS** | - | Sc874 | 8757-21833 | - | 9 | 695 | New gene model. LLOTMP009458 and LLOTMP009459 were fused and edited |
| ***nmo*-NTE** | ***3956*** | Sc235 | 3-10491 | + | 3 | 138 | NTE region is missed |
| ***cry2*-JOI** | - | - | - | - ; + ; - | 5 | 742 | New gene model. The initial part of the model was present in Contig06585 (3-1564), followed by Contig12558 (2-202) and Contig06584 (144-974) |

| Gene | OGS | Scaffold/Contig | Coordinates | Strand | Introns | AAs | Comments |
|---|---|---|---|---|---|---|---|
| ***Phr*-JOI** | - | Ct83811 and Sc741 | 1486-2245 and 34561-36367 | + ; + | 3 | 333 | New gene model. The first 760 bp of the gene model were in Contig83811. The rest of the gene model was located in LLOTMP008682 (Sc741) |
| ***norpA*** | 1939 | Sc1440 | 37196- 45904 | - | 8 | 1081 | Fine as it is |
| ***vri*** | 7416 | Sc568 | 62113-76422 | + | 2 | 469 | Fine as it is |
| ***Pdp1*** | 6612 | Sc48 | 18671-106058 | + | 4 | 267 | Initial methionine was fixed including a new exon |
| ***slmb*** | 6764 | Sc5 | 129351-138574 | + | 10 | 700 | Fine as it is |
| ***cac*-INT** | 6699 | Sc489 | 105764-130064 | - | 20 | 1185 | Multiple changes |
| ***Na*-FUS** | - | Sc137 | 64371-83060 | - | 10 | 3074 | New gene model. LLOTMP001706 and LLOTMP001707 were fused and edited |
| ***para*-NTE** | 7479 | Sc576 | 62487-85952 | - | 27 | 1600 | Multiple changes |
| ***slo*-NTE** | 1559 | Sc132 | 16087-51527 | + | 19 | 1018 | NTE region is missed |
| ***Slob*** | 8864 | Sc770 | 8281-9598 | + | 2 | 158 | First, second and last exons were eliminated |
| ***nocte*-CTE** | 4544 | Sc268 | 88697-92269 | + | 3 | 574 | CTE region is missed |
| ***Atax-2*** | 4561 | Sc269 | 109694-126353 | - | 9 | 935 | Fine as it is |
| ***ctrip*** | 138 | Sc1003 | 20136-40553 | - | 11 | 2100 | Exons 7th and 10th were edited |
| ***to1*** | 6030 | Sc401 | 20817-21728 | - | 2 | 189 | LLOTMP006030 was divided in two gene models (*to1* and *to2*). Last exon extended |
| ***to2*** | - | Sc401 | 18743-19697 | - | 3 | 252 | New gene model. First exon added and 2nd exon extended |
| ***to3*** | 8362 | Sc7 | 47865-49626 | - | 3 | 244 | LLOTMP008362 was divided in two different models (*to3* and *to4*); last exon extended |
| ***to4*-NTE** | - | Sc7 | 40986-41804 | - | 1 | 192 | New gene model |

| | | | | | | | |
|---|---|---|---|---|---|---|---|
| ***to5*** | 6987 | Sc514 | 25417-28591 | - | 2 | 248 | Fine as it is |
| ***to6*** | 8792 | Sc76 | 114913-117338 | - | 1 | 219 | Fine as it is |
| ***Rh3*** | 5188 | Sc326 | 40469 - 47369 | + | 2 | 380 | The 3rd exon has been eliminated |
| ***Rh7*** | 9159 | Sc813 | 59510 60663 | + | 2 | 325 | Fine as it is |
| ***Piezo*-NTE** | 5459 | Sc346 | 1205-5153 | + | 3 | 682 | NTE region is missed |
| ***nompC*** | 3746 | Sc223 | 5365-38664 | + | 17 | 1928 | Fine as it is |
| ***pain-INT*** | 6694 | Sc489 | 4579-1227 | + | 5 | 763 | Multiple changes |
| ***TrpA1*** | 4961 | Sc302 | 27240- 43231 | + | 13 | 1273 | Fine as it is |
| ***wtwr*** | 9849 | Sc941 | 53070-57180 | + | 5 | 971 | Fine as it is |
| ***wtwr*** | 7806 | Sc1655 | 3447-6841 | + | 1 | 997 | Fine as it is |
| ***wtwr*** | 7983 | Sc6425 | 8892 - 11961 | - | 3 | 977 | Fine as it is |
| ***wtwr*** | 6605 | Sc4796 | 21-545 | + | 2 | 988 | Fine as it is |
| ***wtwr*** | 9848 | Sc941 | 44585- 47985 | + | 2 | 979 | Fine as it is |
| ***trp*-JOI/CTE** | - | Sc1365 and Sc1033 | 46030-48856; 18801-23064 | + ; + | 17 | 902 | New gene model. LLOTMP001696 (Sc1365) and LLOTMP000286 (Sc1033) were fused. |
| ***trpgamma*-NTE** | 6929 | Sc508 | 80435-84475 | - | 4 | 591 | The initial methionine is missed |
| ***Trpl*-NTE** | - | Sc118 | 176287-179149 | + | 2 | 331 | New gene model. The initial methionine is missed. |
| ***Trpm*-NTE/INT** | 7263 | Sc552 | 32964-65802 | - | 24 | 1654 | Initial methionine is missed and internal problems were detected |
| ***Trpml*** | 170 | Sc655 | 161373-163489 | + | 1 | 655 | Fine as it is |
| ***ppk3*** | 3220 | Sc198 | 22341-23961 | - | 2 | 487 | Fine as it is |

| Gene | OGS | Scaffold/Contig | Coordinates | Strand | Introns | AAs | Comments |
|---|---|---|---|---|---|---|---|
| ***pkk9*** | - | Sc88 | 17945 - 176385 | + | 1 | 458 | New gene model |
| ***ppk4*** | 7095 | Sc530 | 14760-16306 | - | 3 | 451 | Fine as it is |

| | | | | | | | |
|---|---|---|---|---|---|---|---|
| ***ppk13*-FUS** | - | Sc2154 | 16046 -34392 | + | 9 | 448 | LLOTMP003631 and LLOTMP003631were fused and edited |
| ***ppk16*** | 4198 | Sc2475 | 17735-20905 | - | 2 | 530 | Fine as it is |
| ***ppk16-like*** | 5025 | Sc31 | 42322-47258 | + | 6 | 915 | Fine as it is |
| ***ppk26*** | 9835 | Sc94 | 68591-78879 | + | 6 | 787 | Fine as it is |
| ***ppk26-like*** | 1025 | Sc1188 | 22343-30678 | - | 4 | 575 | Fine as it is |
| ***ppk28*** | 1020 | Sc1187 | 19039-28307 | + | 4 | 504 | Fine as it is |
| ***ppk31*-NTE/CTE** | 7955 | Sc639 | 19353-20752 | - | 3 | 339 | Partial model. CTE and NTE regions were missed |
| ***ppk100*** | 5117 | Sc32 | 206361-210069 | + | 4 | 532 | Fine as it is |
| ***ppk101*** | 9834 | Sc94 | 56855-64178 | - | 5 | 533 | Fine as it is |
| ***ppk102*** | - | Sc94 | 60671-64403 | - | 2 | 547 | New gene model |
| ***ppk*-NTE** | - | Sc299 | 61443-63745 | - | 3 | 404 | New gene model.NTE region was missed |
| ***mlv*** | 4951 | Sc301 | 25631-32046 | - | 11 | 565 | Fine as it is |
| ***Pkg2*-1D** | 1613 | Sc134 | 123869-138596 | - | 7 | 1022 | Fine as it is |
| ***for*-NTE** | 7793 | Sc61 | 40100- 48763 | - | 7 | 515 | NTE region is missed. The last seven exons were eliminated |
| ***sr*-CTE** | 3205 | Sc970 | 5116-8025 | + | 0 | 184 | CTE region was extended |

Official full names:, tim: timeless, per: period, cwo: clockwork orange, Clk: clock, cyc: cycle, sim: Single minded, tgo: tango, sgg: shaggy, dbt: doubletime, CKII alpha: Casein kinase II alpha, CKII beta: Casein kinase II beta, Pp1a: serine/threonine-protein phosphatase 1 alpha, Pp2a: serine/threonine-protein phosphatase 2 alpha, Pp4: serine/threonine-protein phosphatase 4, PpV-6: serine/threonine-protein phosphatase V-6, Pp7: serine/threonine-protein phosphatase 7, nmo: nemo, cry2: Cryptochorme2, phr: DNA photolyase, photorepair (phr), norpA: Phosphoinositide phospholipase C, vri: Vrille, Pdp1: Par-domain protein1, slmb: Supernumerary limbs, cac: cacophony, na: narrowabdomen, para: paralytic, slo: Calcium-activated potassium channel slowpoke , slob: Slowpoke binding protein, nocte: No circadian temperature entrainment, Atax-2: Ataxin-2, ctrip: Circadin trip, to: JHBP/takeout, Rh3: Ultraviolet sensitive opsin, LWO: long-wavelength opsin, Rh7: Rhodopsin, nompC: No mechanoreceptor potential C , pain: painless, TrpA1: transient receptor potential A , wtrw: water witch , trp: transient receptor potential protein, trp gamma: transient receptor potential gamma, trpL: transient receptor potential L , trpm: transient receptor potential cation channel, melastatin subfamily, trpml: transient receptor potential cation channel, mucolipin subfamily, ppk: pickpocket, mlv: malvolio, Pkg2-1D: cGMP-dependent protein kinase, isozyme 1, for: foraging or cGMP-dependent protein kinase, isozyme 2, sr: stripe.

**S21 Table. Detailed information about *Phlebotomus papatasi* circadian and behavior genes and proteins.** Columns: *Gene* – assigned gene and protein (NTE – N-terminus missing, CTE – C-terminus missing, INT – problems in the assembly, FUS – two gene models located in the same scaffold were fused; JOI – gene model spans scaffolds); OGS – the official gene number in the 12,678 genes in Ppapl1.1, prefix is PPAI; Scaffold (Sc) – the Ppapl1.1 genome assembly supercontig ID and Contig (Ct) – the Ppapl1 genome assembly contig ID, prefix is Scaffold; Coordinates – nucleotide range from the first position of the start codon to the last position of the stop codon in the scaffold; Strand + is forward and - is reverse; Introns – number of introns; AAs – number of encoded amino acids in the protein found in the gene model; Comments – comments about the OGS gene model, and repairs that were done in the genome assembly.

| Gene | OGS | Scaffold/Contig | Coordinates | Strand | Introns | AAs | Comments |
|---|---|---|---|---|---|---|---|
| ***tim*-FUS** | 12818 | Sc47 | 137910-156773 | - | 13 | 1201 | PPATMP008008 and PPATMP008007 were fused |
| ***tim2*-JOI** | - | Sc597 | 3338-4648 | - | 9 | 969 | New gene model. It was distributed in Sc597, Scaffold44752 and Scaffold52134 |
| ***per*-JOI** | 6484 | Ct101627.1 and Sc006484 | 4747-9741 and 350-435 | + ; - | 7 | 1158 | 1st exon was located in Contig101627.1 and the rest of the model was in Scaffold4185 (PPATMP006484) |
| ***cwo*-INT** | 12819 | Sc430 | 2201-11463 | - | 2 | 619 | Manual edition |
| ***Clk*** | 2844 | Sc2 | 440906-446226 | + | 6 | 706 | Fine as it is |
| ***cyc*** | 10392 | Sc77 | 344459-357605 | + | 7 | 634 | 1st exon was eliminated and the initial methionine fixed |
| ***sim*-NTE/CTE** | 1673 | Sc15 | 618709-626223 | - | 3 | 240 | The NTE and CTE regions were missed |
| ***tgo*-NTE/CTE** | 9623 | Sc61 | 60289-66121 | - | 6 | 556 | The NTE and CTE regions were missed |
| ***sgg*-NTE** | 10158 | Sc72 | 200643-211489 | + | 4 | 331 | The NTE region was missed |
| ***dbt*- NTE** | 7687 | Sc45652 | 3543-4142 | + | 0 | 199 | The NTE region was missed |
| ***CKII alpha*** | 516 | Sc112 | 71465-72481 | - | 0 | 338 | Fine as it is |
| ***CKII beta*-FUS** | 11154 | Sc996 | 9684-20145 | + | 5 | 227 | New gene model. PPATMP011154 and PPATMP011154 were fused and edited |
| ***Pp1a*-NTE** | 5749 | Sc3582 | 603-11379 | + | 2 | 163 | The NTE region was missed |
| ***Pp2a*-NTE/CTE** | 6622 | Sc42725 | 17875-18809 | + | 1 | 216 | The NTE and CTE regions were missed |
| ***Pp2b*-FUS/CTE** | 000839/?000840 | Sc1211 | 15663-39415 | + | 6 | 401 | New gene model. PPATMP000839 and PPATMP000840 were fused. Note there are still and PPAI000839 & PPAI000840 |

| | | | | | | | |
|---|---|---|---|---|---|---|---|
| ***PpV-6*-NTE** | 3603 | Sc23515 | 3045-4044 | + | 1 | 247 | The NTE region is missed |
| ***Pp7*-JOI/NTE** | 009968/?006145 | Sc672 and Sc3936 | 26965-35214 and 7053-7637 | + | 3 | 376 | PPATMP009968 (Sc672) and PPATMP006145 (Sc3936) were fused and edited. NTE region was missed Note there is still PPAI009968 & PPAI006145 |
| ***nmo*-FUS/NTE** | 12817 | Sc594 | 27097-44269 | + ; - | 6 | 360 | New gene model. PPATMP009517 and PPATMP009518 were fused |
| ***cry1*-NTE** | 2137 | Sc1671 | 26175-26932 | - | 0 | 253 | Model is partial and only contains the last exon |
| ***cry2*-JOI** | 12817 | Sc46912 and Sc2983 | 2643-4050, 301-1506 | + ; - | 4 | 743 | New gene model. PPATMP009517 (Sc46912) and PPATMP009518 (Sc2983) were fused. |
| ***phr*** | 4254 | Sc261 | 63214-64919 | + | 1 | 515 | Fine as it is |
| ***norpA*-JOI** | 007015/?007016 | Sc4354 and Sc64711 | - | - | 5 | 801 | New gene model IN Sc4354 two predicted protein (PPATMP007015 and PPATMP007016) were fused.<br>In Sc 64711 a new predicted protein was created. Note that PPAI007015 and PPAI007016 are both still there. |

| Gene | OGS | Scaffold/Contig | Coordinates | Strand | Introns | AAs | Comments |
|---|---|---|---|---|---|---|---|
| ***vri*-JOI** | ?????? | Sc26106 and Sc64592 | 222-977 and 189-764 | - | 0 | 344 | New gene model |
| ***Pdp1*-NTE** | 2842 | Sc2 | <273251-281247 | + | 2 | 190 | NTE region was missed |
| ***slmb*-INT** | 4879 | Sc299 | 7110-7561 | + | 0 | 150 | Assembly problem |
| ***cac*-NTE** | 12826 | Sc809 | 835-1791 | - | 2 | 267 | Model is partial and only contains three exons; first four exons of the original model were deleted. |
| ***na*-CTE** | 8229 | Sc48310 | 2841->8148 | + | 2 | 1122 | CTE region is missed |
| ***para*-JOI** | 003387/?394&95 | Sc2248 and Sc1094 | - | - , - ,+, + | 16 | 980 | New gene model. Four gene models were fused: PPATMP003387 (coordinates: 9174-14426) and PPATMP003386 (coordinates: 1151-7841) located in Sc2248; PPATMP000394 (coordinates: 7644-14047) and PPATMP000395 (coordinates: 26395-28039) located in Sc1094. Multiple assembly problems. Note all three IDs still there, please check. |
| ***slo*-JOI/CTE** | - | Multiple | - | +, - , - ,+ ,+ | 10 | 626 | New gene model. Five gene models were fused and edited. PPATMP007684 (Sc45640, coordinates 5741-6062); PPATMP009501 (Sc5903, coordinates: 3467-4002); PPATMP003816 (Sc2407, coordinates: 2902-24213); PPATMP005694 (Sc3539, coordinates: 5161-8967) and PPATMP006643 (Sc42743, coordinates: 3497-9448) All still there |
| ***nocte*-NTE** | 2962 | Sc2032 | <3916-11712 | + | 3 | 1660 | NTE region was missed |
| ***Atax-2*** | 4801 | Sc2931 | 2462-12528 | - | 3 | 401 | CTE region was extended |
| ***ctrip*-FUS** | 12822 | Sc816 | 50562-51523 | + | 4 | 1684 | New gene model. Two gene models (PPATMP010549 and PPATMP010550) were fused and edited |

| | | | | | | | |
|---|---|---|---|---|---|---|---|
| ***to1*-NTE** | 7441 | Sc44847 | <7218-7718 | + | 0 | 166 | NTE region is missed |
| ***to2*** | 12824 | Sc1162 | <11478-11916 | - | 0 | 145 | PPATMP000647 was divided in two models (*to2* and *to3*). Initial methionine was missed |
| ***to3*-NTE** | ?????? | Sc1162 | <6974-7842 | - | 2 | 225 | New gene model. Initial methionine was missed. |
| ***to4*-NTE** | 648 | Sc1162 | <20371-21129 | - | 1 | 225 | PPATMP000648 was splitted in two models (*to4* and *to5*); Initial methionine was missed. |
| ***to5*-NTE** | ?????? | Sc1162 | <17873-18785 | - | 2 | 226 | New gene model. Initial methionine was missed |
| ***Rh3*-FUS** | 012839/?0 12816 | Sc2 | 1226541-1240886 | - | 2 | 379 | Two gene models (PPATMP002883-RA and PPATMP002881) were fused and edited Note: 2 different IDs map |
| ***LWO*** | 4207 | Sc26 | 547523-549089 | + | 1 | 381 | Fine as it is |
| ***Rh7*-INT** | 7252 | Sc44163 | ? | +; - | 1 | 136 | Gene model was edited. Assembly problems |
| ***Piezo*-INT** | 12821 | Sc38 | 253661-462668 | - | 19 | 1959 | Multiple changes |
| ***iav*** | 4901 | Sc3 | 279511-291919 | + | 5 | 1135 | Multiple changes |
| ***nan*-JOI** | 7810 | Sc4610 and Sc47892 | 4988-? | - ; + | 4 | 835 | New gene model. PPATMP007810 (Sc4610) and Contig8123.1 and Contig190787.1 (Sc47892) were fused. |
| ***pain*** | 3852 | Sc2424 | 160-2844 | + | 0 | 2685 | Fine at it is |
| ***TrpA1*-NTE** | 4036 | Sc2513 | <8698-20514 | - | 9 | 1119 | NTE region was missed |
| ***wtrw*** | 8786 | Sc52 | 76321-79438 | + | 1 | 1003 | Fine at it is |
| ***wtrw*** | 8787 | Sc52 | 99530-102726 | + | 1 | 979 | One extra exon was added at NTE region |
| ***wtrw*** | 8788 | Sc52 | 114914-117964 | + | 1 | 993 | Fine at it is |

| **Gene** | **OGS** | **Scaffold/Contig** | **Coordinates** | **Strand** | **Introns** | **AAs** | **Comments** |
|---|---|---|---|---|---|---|---|
| ***trp*-JOI** | 5961 | Sc377 | - | ? | ? | 707 | Manually created. Split in many different scaffolds. Partial model with CTE and NTE regions missed |
| ***trp gamma*-FUS/NTE** | 00775? | Sc12 | <491600-498054 | - | 3 | 581 | Two gene models (PPATMP00775 and PPATMP00776) were fused and edited. CONFUsing, please check. |
| ***trpL*** | 9654 | Sc615 | 2582-9933 | + | 5 | 1019 | Fine as it is |

| | | | | | | | |
|---|---|---|---|---|---|---|---|
| ***trpm*-NTE** | ????? | Sc46755, Sc5448, Sc50709 | - | ? | ? | 681 | Manually created. NTE region is missed |
| ***trpml*** | ????? | Sc43374, Sc87076, Sc27005 | - | ? | ? | 635 | Manually created |
| ***ppk3*-NTE** | ????? | Sc348 | <79190-80437 | - | 0 | 415 | NTE region was missed |
| ***ppk16*** | 6353 | Sc4085 | 2373-9139 | - | 4 | 526 | Four exons were added at CTE region |
| ***ppk16-like*** | 2706 | Sc191 | 136896-138806 | - | 2 | 575 | Fine as it is |
| ***ppk26-likeA*** | 9365 | Sc574 | 29919-35026 | + | 3 | 583 | Fine as it is |
| ***ppk26-likeB*-INT** | 9363 | Sc574 | 29919-35026 | + | 3 | 583 | Problems in the assembly of the scaffold were detected |
| ***ppk23*** | 1398 | Sc14 | 76160-80084 | - | 2 | 552 | Fine as it is |
| ***ppk28*** | 4084 | Sc2541 | 4922-12752 | - | 5 | 561 | Multiple changes |
| ***ppk31a*-NTE/CTE** | 9142 | Sc5514 | <321->4531 | - | 2 | 323 | 1st exon eliminated and the last exon extended |
| ***ppk31b*-NTE/CTE** | 3967 | 2495 | <21580->22580 | + | 1 | 266 | 1st exon eliminated and the last exon extended |
| ***ppk100*** | 1000 | 127 | 21689-25364 | + | 4 | 536 | Fine as it is |
| ***ppk101*** | 980 | 1261 | 6128-12698 | + | 2 | 516 | Fine as it is |
| ***ppk102*-CTE** | 978 | 1261 | 3674->4681 | + | 1 | 316 | The last exon eliminated |
| ***ppk-like*-NTE** | 12838 | 42794 | <5679-6785 | + | 1 | 344 | Two first exons were eliminated. NTE region was missed |
| ***mlv*-NTE/CTE** | ????? | 24350 | >641-<1568 | - | 2 | 251 | New gen model. Partial model |
| ***Pkg2-1D*** | 6090 | 39 | 103022-112773 | - | 6 | 812 | Fine as it is |
| ***sr*-NTE** | 5020 | 302 | >26270-69382 | - | 3 | 847 | The initial methionine was missed. Multiple changes |

Official full names:, tim: timeless, per: period, cwo: clockwork orange, Clk: clock, cyc: cycle, sim: Single minded, tgo: tango, sgg: shaggy, dbt: doubletime, CKII alpha: Casein kinase II alpha, CKII beta: Casein kinase II beta, Pp1a: serine/threonine-protein phosphatase 1 alpha, Pp2a: serine/threonine-protein phosphatase 2 alpha, Pp2a: serine/threonine-protein phosphatase 2 beta, PpV-6: serine/threonine-protein phosphatase V-6, Pp7: serine/threonine-protein phosphatase 7, nmo: nemo, cry1: Cryptochorme1, cry2: Cryptochorme2, phr: DNA photolyase, photorepair (phr), norpA: Phosphoinositide phospholipase C, vri: Vrille, Pdp1: Par-domain protein1, slmb: Supernumerary limbs, cac: cacophony, na: narrowabdomen, para: paralytic, slo: Calcium-activated potassium channel slowpoke , nocte: No circadian temperature entrainment, Atax-2: Ataxin-2, ctrip: Circadin trip, to: JHBP/takeout, Rh3: Ultraviolet sensitive opsin, LWO: long-wavelength opsin, Rh7: Rhodopsin, pain: painless, TrpA1: transient receptor potential A , wtrw: water witch , trp: transient receptor potential protein, trp gamma: transient receptor potential gamma, trpL: transient receptor potential L , trpm: transient receptor potential cation channel, melastatin subfamily, trpml: transient receptor potential cation channel, mucolipin subfamily, ppk: pickpocket, mlv: malvolio, Pkg2-1D: cGMP-dependent protein kinase, isozyme 1, sr: stripe.

**S22 Table.** Repertoire sizes of the three major chemoreceptor families in *Lu. longipalpis* and *P papatasi*. Splice variants were predicted for two GR genes in each species (Gr13 and Gr26).

| Species | ORs | GRs (genes) | IRs |
|---|---|---|---|
| *Lutzomyia longipalpis* | 140 | 91 (82) | 23 |
| *Phlebotomus papatasi* | 140 | 88 (77) | 28 |

**S23 Table.** G-protein coupled receptor family annotation.

Comparison of *Phelbotomus papatasi* and *Lutzomyia longipalpis* non-sensory and opsin G protein-coupled receptors (GPCRs) to other insects.

| GPCR class | GPCR Subclass/family* | *P. papatasi* | *Lu. longipalpis* | *D. melanogaster* | *A. aegypti* | *A. gambiae* | *P. h. humanus* |
|---|---|---|---|---|---|---|---|
| **Class A: Rhodopsin-like** | Biogenic amine | 15 | 15 | 17 | 26 | 18 | 21 |
| | Glycoprotein hormone | 4 | 4 | 4 | 3 | 3 | 5 |
| | Peptide | 24 | 23 | 30 | 33 | 25 | 21 |
| | Purine | 1 | 1 | 1 | 1 | 1 | 2 |
| | (Rhod)opsin | 5 | 5 | 7 | 10 | 11 | 3 |
| | Orphan | | | 32 | 16 | 22 | 17 |
| **Class B: Secretin-like** | Calcitonin-like | 3 | 3 | 3 | 3 | 3 | 3 |
| | Diuretic insect hormone | 2 | 2 | 3 | 2 | 2 | 1 |
| | Growth hormone releasing hormone | 4 | 4 | 3 | 4 | 4 | 2 |
| | HE6 like | 1 | 1 | 2 | 1 | 1 | 0 |
| | Latrophilin | 1 | 1 | 1 | 1 | 1 | 2 |
| | Methuselah-like | 7 | 7 | 15 | 9 | 7 | 8 |
| | Ocular Albinism | 1 | 1 | 0 | 1 | 1 | 1 |
| | Parathyroid hormone | 1 | 1 | 1 | 0 | 0 | 1 |
| | Orphan | 2 | 3 | 2 | 4 | 4 | 1 |
| **Class C: Metabotropic glutamate-like** | Metabotropic glutamate | 5 | 5 | 5 | 5 | 5 | 7 |
| | $GABA_B$ | 3 | 3 | 4 | 3 | 3 | 2 |
| | Orphan | 0 | 0 | 0 | 0 | 0 | 1 |
| **Class D: Atypical GPCRs** | Boss | 1 | 1 | 1 | 1 | 1 | 1 |
| | Frizzled/Smoothened | 4 | 3 | 6 | 11 | 8 | 5 |
| | Starry Night | 1 | 1 | 1 | 1 | 1 | 1 |
| | Orphan | 2 | 2 | 2 | 2 | 3 | 2 |
| | **Total** | 94 | 92 | **140** | **137** | **124** | **107** |

Comprehensive list of *Phelbotomus papatasi* non-sensory and opsin GPCRs.

| Species | GPCR class | GPCR subclass | GPCR family | *P. papatasi* GPCR | *P. papatasi* scaffold # | Base pair range on scaffold | Vectorbase (PpapI1.1) Gene ID |
|---|---|---|---|---|---|---|---|
| ***P. papatasi*** | **(1) Class A-Rhod(opsin) receptor family** | **Amine receptors** | **Muscarinic acetylcholine** | GPRmac1_1 | 3896 | 7489..9646 | PPAI006081 |
| | | | | GPRmac1_2§ | 3896 | 10322..10269 | NEW |
| | | | | GPRmac1_3 | 318 | 23923..31276 | PPAI005235 |
| | | | | GPRmac2 | 216 | 93760..100476 | PPAI003220 |
| | | | **Adrenergic** | GPRadr1 | | | |
| | | | | GPRadr2 | | | |
| | | | **Dopamine** | GPRdop1_1 | 296 | 78784..78580 | NEW |
| | | | | GPRdop1_2 | 139 | 13283..1091 | PPAI001377 |
| | | | | GPRdop1_3 | 1564 | 30508..29981 | PPAI001852 |
| | | | | GPRdop2 | 334 | 73913..65556 | PPAI005443 |
| | | | | GPRdop3_1 | 2510 | 20941..15326 | PPAI004032 |
| | | | | GPRdop3_2 | 44987 | 5425..4961 | PPAI007491 |
| | | | | GPRdop3_3 | 4989 | 3208..2416 | PPAI008468 |
| | | | | GPRdop3_4 | 49896 | 2611..2397 | NEW |
| | | | **Histamine** | GPRhis1_1 | 2900 | 15070..13982 | PPAI004760 |
| | | | | GPRhis1_2 | 2900 | 16322..16025 | PPAI004761 |
| | | | | GPRhis1_3 | 2900 | 16317..16784 | NEW |
| | | | **Melatonin** | GPRmtn1_1 | 51201 | 5739..4841 | PPAI008718 |
| | | | | GPRmtn1_2 | 51201 | 5536..5832 | PPAI008717 |
| | | | **Octopamine** | GPRoar1_1 | 63030 | 720..469 | NEW |
| | | | | GPRoar1_2 | 1953 | 18433..2120 | PPAI002761 |
| | | | | GPRoar1_3 | 2017 | 21568..22367 | PPAI002932 |
| | | | | GPRoar2_1 | 85 | 273392..273157 | PPAI010658 |
| | | | | GPRoar2_2 | 85 | 238998..259019 | PPAI010657 |
| | | | | GPRoar2_3 | 85 | 268521..268194 | PPAI010658 |
| | | | | GPRoar3 | 4 | 724535..738443 | PPAI006236 |
| | | | **Tyramine** | GPRtyr1 | 317 | 67847..66099 | PPAI00522 |
| | | | **Serotonin** | GPR5ht1a_1 | 2 | 1589842..1600089 | PPAI002892 |
| | | | | GPR5ht1a_2 | 2 | 1610028..1610851 | PPAI002893 |
| | | | | GPR5ht1b_1 | 2356 | 8615..8105 | NEW |
| | | | | GPR5ht1b_2 | 1439 | 25835..25591 | PPAI001528 |
| | | | | GPR5ht1b_3 | 1439 | 33631..34021 | PPAI001529 |
| | | | | GPR5ht1b_4 | 1910 | 4725..5327 | PPAI002707 |
| | | | | GPR5ht2a_1 | 75 | 309261..320192 | PPAI010317 |
| | | | | GPR5ht2a_2 | 64577 | 72..247 | NEW |
| | | | | GPR5ht2a_3 | 75 | 341430..341749 | NEW |
| | | | | GPR5ht7_1 | 102 | 43..468 | NEW |
| | | | | GPR5ht7_2 | 1545 | 29722..8896 | PPAI001792 |
| | | | | GPR5htorph2 | 75 | 352089..359167 | PPAI010318 |

| | | | | | |
|---|---|---|---|---|---|
| **Glycoprotein Hormone/Hormone** | | GPRbcn | 963 | 6543..12990 | PPAI005338 |
| | | GPRfsh | 5660 | 4211..13988 | PPAI005532 |
| | | GPRgph1_1 | 243 | 55332..55432 | NEW |
| | | GPRgph1_2 | 243 | 52706..52822 | NEW |
| | | GPRgph1_3 | 243 | 49132..50199 | PPAI003861 |
| | | GPRgph1_4 | 243 | 62395..64533 | PPAI003862 |
| | | GPRpyn_1 | 45935 | 785..4188 | PPAI007748 |
| | | GPRpyn_2 | 60331 | 879..1602 | NEW |
| **Peptide receptors** | **Adipokinetic hormone** | GPRakh | 689 | 43077..48721 | |
| | **CAPA** | GPRcap_1 | 1827 | 2776..2340 | |
| | | GPRcap_2 | 1827 | 1484..2073 | |
| | **CCAP** | GPRccp | 7643 | 13455..17854 | |
| | **Corazonin** | GPRczn | 435 | 99872..100076 | |
| | **Ecdysis triggering hormone receptor** | GPReth | 689 | 32..1178 | |
| | **FMRFamide** | GPRfmr | 154 | 111107..114206 | |
| | **Galanin/Allatostatin** | GPRals_1 | 57 | 107977..115249 | |
| | | GPRals_2 | 57 | 123959..132324 | |
| | **Gastrin/Bombesin** | GPRgrp | 11088 | 1300..1896 | |
| | **Leukokinin** | GPRllk1_1 | 3372 | 14087..14664 | |
| | | GPRlik1_2 | 613 | 54..894 | |
| | **Myosuppressin** | GPRmys1 | 4256 | 9055..11742 | |
| | **Neuropeptide** | GPRnpr1 | 37 | 86709..99295 | |
| | | GPRnpr2_1 | 175 | 135476..135735 | |
| | | GPRnpr2_2 | 45332 | 436..1369 | |
| | | GPRnpr3 | | | |
| | **SIFamide** | GPRsif | 42 | 943..5322 | |
| | **Sulfakinin** | GPRsfk1 | 312 | 95430..99665 | |
| | | GPRsfk2 | 4550 | 432..8774 | |
| | **Sex peptide receptor** | GPRspr1_1 | 43379 | 2946..2634 | |
| | | GPRspr1_2 | 57097 | 1590..1326 | |
| | | GPRspr1_3 | 69417 | 587..344 | |
| | | GPRspr1_4 | 2184 | 21897..21291 | |
| | **Somatostatin** | GPRsms1_1 | 54233 | 844..275 | |
| | | GPRsms1_2 | 2967 | 11943..11721 | |
| | | GPRsms1_3 | 44139 | 6932..3038 | |
| | **Tachykinin** | GPRtak1_1 | 54644 | 1740..2166 | |
| | | GPRtak1_2 | 46427 | 8525..8738 | |
| | | GRPtak1_3 | 3655 | 9688..18014 | |
| | | GPRtak2_1 | 24818 | 351..206 | |
| | | GPRtak2_2 | 25382 | 959..414 | |
| | | GPRtak2_3 | 364 | 18782..3670 | |

| | | | | | | |
|---|---|---|---|---|---|---|
| | | | GPRtak2_4 | 2647 | 13909..13744 | |
| | **Purine receptors** | **Opioid** | | | | |
| | | **Adenosine** | GPRads1_1 | 767 | 4314..9910 | |
| | | | GPRads1_2 | 767 | 10678..11512 | |
| | **(Rhod)opsin receptors** | Ultraviolent | GPRop1 | 26 | 99757..101623 | |
| | | Long | GPRop2 | 2 | 54032..76088 | |
| | | | GPRop3 | 44613 | 866..11902 | |
| | | | GPRop4 | 881 | 23..1004 | |
| | | | GPRop5 | 38 | 10087..19963 | |
| | **Orphan/Putative Class A GPCRs** | | GPRorp1 £ | | | |
| | | | GPRorp2 | | | |
| | | | GPRorp3 | | | |
| | | | GPRorp4 | | | |
| | | | GPRorp5 | | | |
| | | | GPRorp6 | | | |
| | | | GPRorp7 | | | |
| | | | GPRorp8 | | | |
| | | | GPRorp9 | | | |
| | | | GPRorp10 | | | |
| | | | GPRorp11 | | | |
| | | | GPRorp12 | | | |
| | | | GPRorp13 | | | |
| | | | GPRorp14 | | | |
| | | | GPRorp15 | | | |
| | | | GPRorp16 | | | |
| | | | GPRorp17 | | | |
| **(2) Class B – Secretin receptor family** | **Calcitonin-like/ PDF receptors** | | GPRcal1 | 22785 | 578..1329 | NEW |
| | | | GPRcal2 | 5446 | 9043..1965 | PPAI006646 |
| | | | GPRpdf | 782 | 457..1153 | NEW |
| | **Diuretic insect hormone receptors** | | GPRdih1_1 | 1911 | 11755..11536 | PPAI002708 |
| | | | GPRdih1_2 | 42935 | 3173..3723 | PPAI006749 |
| | | | GPRdih1_3 | 1911 | 18252..18677 | PPAI002709 |
| | | | GPRdih2_1 | 2125 | 568..684 | NEW |
| | | | GPRdih2_2 | 2125 | 20012..19626 | NEW |
| | | | GPRdih2_3 | 2125 | 23862..24132 | NEW |
| | | | GPRdih2_4 | 29177 | 595..1 | NEW |
| | | | GPRdih2_5 | 72295 | 1..78 | NEW |
| | **Growth hormone releasing hormone receptors** | | GPRghp1 | 6743 | 65432..78549 | PPAI005892 |
| | | | GPRghp2 | 298 | 1004..2954 | NEW |
| | **Latrophilin receptors** | | GPRcir1 | 351 | 599..1764 | NEW |
| | | | GPRcir2 | 32 | 14099..28674 | PPAI005889 |

| | | | | | |
|---|---|---|---|---|---|
| | **Methuselah-like receptors** | GPRmth1 | 785 | 1333..11458 | PPAI007663 |
| | | GPRmth2 | 8774 | 439..1390 | NEW |
| | | GPRmth3 | 4998 | 14679..10065 | PPAI007732 |
| | | GPRmth4 | 3902 | 55892..49078 | PPAI000278 |
| | | GPRmth5 | 235 | 14087..2289 | PPAI002775 |
| | | GPRmth6 | 9065 | 6543..1167 | PPAI009933 |
| | **Ocular albinism (Type I) receptors** | GPRoca1 | 997 | 1094..2865 | NEW |
| | **Parathyroid hormone receptors** | GPRpth | 4301 | 437..1126 | NEW |
| | **Orphan/ Putative Class B GPCRs** | GPRorp1 | | | |
| **(3) Class C – Metabotropic glutamate receptor family** | **Metabotropic glutamate receptors** | GPRmgl1_1 | 446 | 708..551 | NEW |
| | | GPRmgl1_2 | 45142 | 4170..3956 | NEW |
| | | GPRmgl1_3 | 42591 | 39..1 | NEW |
| | | GPRmgl1_4 | 369 | 686..587 | NEW |
| | | GPRmgl1_5 | 284 | 1182..1077 | NEW |
| | | GPRmgl1_6 | 336 | 77727..77571 | NEW |
| | | GPRmgl1_7 | 491 | 2136..1247 | NEW |
| | | GPRmgl2_1 | 42779 | 9957..9549 | PPAI006658 |
| | | GPRmgl2_1 | 25137 | 1267..435 | NEW |
| | | GPRmgl2_3 | 42751 | 2258..3225 | PPAI006647 |
| | | GPRmgl2_4 | 42751 | 5854..5173 | PPAI006648 |
| | | GPRmgl3 | 9664 | 467..1129 | NEW |
| | | GPRmgl4 | 344 | 9987..17044 | PPAI002265 |
| | | GPRrk1_1 | 1 | 111936..53290 | PPAI000007 |
| | **Rickets** | GPRrk1_2 | 1 | 49741..48491 | PPAI000006 |
| | | GPRrk1_3 | 1 | 43441..36693 | PPAI000005 |
| | | GPRgbb1_1 | 1315 | 2637..2419 | NEW |
| | | GPRgbb1_2 | 1315 | 12383..32024 | PPAI001220 |
| | | GPRgbb1_3 | 51752 | 460..318 | NEW |
| | | GPRgbb1_4 | 59998 | 1433..2258 | NEW |
| | | GPRgbb1_5 | 3635 | 5683..13269 | PPAI005823 |
| | | GPRgbb1_6 | 42772 | 2468..2728 | NEW |
| | | GPRgbb2_1 | 42974 | 4270..4007 | PPAI006769 |
| | **GABA(B) receptors** | GPRgbb2_2 | 57227 | 1259..1915 | NEW |
| | | GPRgbb2_3 | 62211 | 269..1024 | NEW |
| | | GRPgbb2_4 | 24716 | 1395..320 | NEW |
| | | GPRgbb2_5 | 2041 | 633..233 | PPAI002976 |
| | | GPRgbb2_6 | 3521 | 485..1152 | NEW |
| | | GPRgbb3_1 | 58 | 153830..178897 | PPAI009413 |
| | | GPRgbb3_2 | 24161 | 1568..1503 | NEW |
| | | GPRgbb3_3 | 58 | 197762..245260 | PPAI009413 |
| | **Orphan/Putative Class C GPCRs** | GPRorp1 | 9077 | 35099..44987 | PPAI004489 |
| | **Bride of Sevenless** | GPRbos | 57 | 263565..268317 | PPAI009316 |

| | | | | | |
|---|---|---|---|---|---|
| **(4) Class D- Atypical 7TM proteins** | **Frizzled** | GPRfr1_1 | 2824 | 21313..20321 | PPAI004613 |
| | | GPRfz1_2 | 2824 | 5199..4885 | PPAI004612 |
| | | GPRfz1_3 | 54316 | 2435..2660 | NEW |
| | | GPRfz1_4 | 1458 | 26346..26878 | PPAI001566 |
| | | GPRfz2 | 17 | 123366..121447 | PPAI002197 |
| | | GPRfz3 | 1081 | 37399..42670 | PPAI000362 |
| | | GPRfz4_1 | 1307 | 35838..35343 | NEW |
| | | GPRfz4_2 | 1307 | 34729..35156 | NEW |
| | | GPRfz4_3 | 1930 | 16993..16797 | PPAI002734 |
| | | GPRfz4_4 | 46463 | 3377..3773 | NEW |
| | | GPRfz4_5 | 45223 | 2250..2563 | NEW |
| | **Smoothened** | GPRsmo_1 | 47187 | 4599..5680 | PPAI008047 |
| | | GPRsmo_2 | 47187 | 4418..2376 | PPAI008046 |
| | **Starry night** | GPRstn | 854 | 32005..58994 | PPAI002355 |
| | **Orphan/Putative Class D GPCRs** | GPRorp1 | | | |
| | | GPRorp2 | | | |
| **(5) Short peptides/partial gene models** | | | | | |

Comprehensive list of *Lu. Longipalpis* non-sensory and opsin GPCRs.

| Species | GPCR class | GPCR subclass | GPCR family | *Lu. longipalpis* GPCR | *Lu. longipalpis* scaffold # or contig # | Base pair range on scaffold | Vectorbase (LlonJ1.1) Gene ID |
|---|---|---|---|---|---|---|---|
| ***Lu. longipalpis*** | **(1) Class A-Rhod(opsin) receptor family** | **Amine receptors** | **Muscarinic acetylcholine** | GPRmac1 | 3 | 559079..568406 | LLOJ004913 |
| | | | | GPRmac2 | 530 | 8673..11046 | LLOJ007094 |
| | | | **Adrenergic** | GPRadr1 | | | |
| | | | | GPRadr2 | | | |
| | | | **Dopamine** | GPRdop1 | 1881 | 489..2308 | NEW |
| | | | | GPRdop2 | 1883 | 233..1709 | NEW |
| | | | | GPRdop3 | 116 | 71091..83702 | LLOJ000916 |
| | | | **Histamine** | GPRhis | 193 | 159773..157938 | LLOJ003111 |
| | | | **Melatonin** | GPRmtn | 790 | 7868..9361 | LLOJ008997 |
| | | | **Octopamine** | GPRoar1 | 57 | 194217..221043 | LLOJ007432 |
| | | | | GPRoar2_1 | 968 | 26020..30479 | LLOJ009965 |
| | | | | GPRoar2_2§ | 968 | 40136..39970 | NEW |
| | | | | GPRoar2_3 | 968 | 32689..33013 | LLOJ009965 |
| | | | | GPRoar3 | 578 | 32099..17089 | LLOTMP010543 |

| | | | | | |
|---|---|---|---|---|---|
| | **Tyramine** | GPRtyr1 | 1573 | 8856..4931 | LLOJ002317 |
| | **Serotonin** | GPR5ht1a_1 | 752 | 197..506 | NEW |
| | | GPR5ht1a_2 | 2619 | 173..10196 | LLOJ004430 |
| | | GPR5ht1a_3 | 277 | 106943..101018 | LLOJ004661 |
| | | GPR5ht1b_1 | 523 | 65450..74495 | LLOJ007042 |
| | | GPR5ht1b_2 | 523 | 1036..425 | NEW |
| | | GPR5ht3_1 | 285 | 69600..70241 | LLOJ004763 |
| | | GPR5ht3_2 | 285 | 70243..70884 | LLOJ004763 |
| | | GPR5ht4 | 767 | 3209..7886 | LLOJ004409 |
| | | GPR5htorph2 | 1335 | 19571..14084 | LLOJ001604 |
| **Glycoprotein Hormone/Hormone** | | GPRbcn | 679 | 45087..62348 | LLOJ004322 |
| | | GPRfsh | 6643 | 1889..54 | NEW |
| | | GPRgph1_1 | 5227 | 869..769 | NEW |
| | | GPRgph1_2 | 99661 | 549..433 | NEW |
| | | GPRgph1_3 | 5227 | 416..57 | NEW |
| | | GPRgph1_4 | 341 | 528..1050 | NEW |
| | | GPRgph1_5 | 866 | 397..45 | NEW |
| | | GPRpyn | 244 | 54098..71077 | LLOJ009953 |
| **Peptide receptors** | **Adipokinetic hormone** | GPRakh | 659 | 54099..42138 | LLOJ008872 |
| | **CAPA** | GPRcap | 1004 | 67099..51032 | LLOJ006681 |
| | **CCAP** | GPRccp | 904 | 16098..33065 | LLOJ002554 |
| | **Corazonin** | GPRczn | 233 | 2145..8832 | LLOJ004497 |
| | **Ecdysis triggering hormone receptor** | GPReth | 991 | 6543..1022 | NEW |
| | **FMRFamide** | GPRfmr1_1 | 598 | 9045..9432 | LLOJ007619 |
| | | GPRfmr1_2 | 2517 | 2362..2600 | NEW |
| | | GPRfmr1_3 | 598 | 10688..11467 | LLOJ007619 |
| | **Galanin/Allatostatin** | GPRals | 326 | 101187..87054 | LLOJ001995 |
| | **Gastrin/Bombesin** | GPRgrp | 5448 | 64088..89043 | LLOJ003386 |
| | **Leukokinin** | GPRllk1_1 | 3589 | 55099..81076 | LLOJ004276 |
| | | GPRllk1_2 | 3589 | 2408..4806 | NEW |
| | **Myosuppressin** | GPRmys1 | 3228 | 58092..66087 | LLOJ006558 |

| | | | | | |
|---|---|---|---|---|---|
| | | GPRmys2 | 3228 | 43207..13066 | LLOJ006559 |
| | **Neuropeptide** | GPRnpr1 | 112 | 1990..45 | NEW |
| | | GPRnpr2 | 4338 | 2456..16098 | LLOJ005398 |
| | | GPRnpr3 | 342 | 55089..48097 | LLOJ004228 |
| | **SIFamide** | GPRsif | 2669 | 32088..51098 | LOJ007664 |
| | **Sulfakinin** | GPRsfk1 | 908 | 12..1309 | NEW |
| | | GPRsfk2 | 32 | 1..1508 | NEW |
| | **Sex peptide receptor** | GPRspr | 298 | 25086..32149 | LLOJ004850 |
| | **Somatostatin** | GPRsms1_1 | 2119 | 821..95 | LLOJ003560 |
| | | GPRsms1_2 | 677 | 2049..2271 | NEW |
| | | GPRsms1_3 | 99 | 822..1354 | NEW |
| | **Tachykinin** | GPRtak1_1 | 778 | 1016..1451 | NEW |
| | | GPRtak1_2 | 1834 | 4485..4698 | LLOJ002928 |
| | | GPRtak1_3 | 781 | 871..1005 | NEW |
| | | GPRtak1_4 | 807 | 211..969 | NEW |
| | | GPRtak2_1 | 1493 | 5242..10475 | LLOJ002079 |
| | | GPRtak2_2 | 251899 | 690..1710 | NEW |
| | | GPRtak2_3 | 1493 | 21464..24956 | LLOJ002080 |
| **Purine receptors** | **Adenosine** | GPRads1 | 908 | 6543..11256 | LLOJ004227 |
| **(Rhod)opsin receptors** | | GPRop1 | 736 | 3255..11298 | NEW |
| | | GPRop2 | 767 | 9887..436 | LLOJ005188 |
| | | GPRop3 | 4497 | 55897..33256 | LLOJ005189 |
| | | GPRop4 | 314 | 56998..88652 | LLOJ005884 |
| | | GPRop5 | 5668 | 76435..60113 | LLOJ008760 |
| **Orphan/Putative Class A GPCRs** | | GPRorp1 | | | |
| | | GPRorp2 | | | |
| | | GPRorp3 | | | |
| | | GPRorp4 | | | |
| | | GPRorp5 | | | |
| | | GPRorp6 | | | |
| | | GPRorp7 | | | |
| | | GPRorp8 | | | |

| | | | | | |
|---|---|---|---|---|---|
| | | GPRorp9 | | | |
| | | GPRorp10 | | | |
| | | GPRorp11 | | | |
| | | GPRorp12 | | | |
| | | GPRorp13 | | | |
| | | GPRorp14 | | | |
| | | GPRorp15 | | | |
| | | GPRorp16 | | | |
| | | GPRorp17 | | | |
| | **Calcitonin-like/ PDF receptors** | GPRcal1 | 87 | 9986..506 | NEW |
| | | GPRcal2 | 4338 | 42244..67708 | LLOJ002219 |
| | | GPRpdf | 47009 | 14389..22358 | LLOJ005442 |
| **(2) Class B – Secretin receptor family** | **Diuretic insect hormone receptors** | GPRdih1_1 | 224 | 837..606 | NEW |
| | | GPRdih1_2 | 226 | 108356..106621 | LLOJ003806 |
| | | GPRdih2 | 1273 | 34144..17341 | LLOJ001389 |
| | **Growth hormone releasing hormone receptors** | GPRghp1 | 5668 | 55987..64098 | LLOJ006544 |
| | | GPRghp2 | 325 | 77543..55099 | LLOJ004337 |
| | **Latrophilin receptors** | GPRcir1 | 996 | 433..10224 | NEW |
| | | GPRcir2 | 450 | 6588..13045 | NEW |
| | **Methuselah-like receptors** | GPRmth1 | 8709 | 54699..80886 | LLOJ008870 |
| | | GPRmth2 | 70086 | 3588..14087 | NEW |
| | | GPRmth3 | 548 | 23007..55901 | LLOJ004484 |
| | | GPRmth4 | 3289 | 4399..23067 | LLOJ007742 |
| | | GPRmth5 | 4990 | 45099..62349 | LLOJ008864 |
| | | GPRmth6 | 663 | 9987..548 | NEW |
| | **Ocular albinism (Type I) receptors** | GPRoca1 | 55990 | 65008..44589 | NEW |
| | **Parathyroid hormone receptors** | GPRpth | 5521 | 3509.435 | NEW |
| | **Orphan/ Putative Class B GPCRs** | GPRorp1 | 4498 | 540087..45532 | LLOJ009863 |
| **(3) Class C – Metabotropic glutamate receptor family** | **Metabotropic glutamate receptors** | GPRmgl1 | 209 | 115030..99753 | LLOJ003515 |
| | | GPRmgl2_1 | 1900 | 1129..739 | LLOJ003046 |
| | | GPRmgl2_2 | 469 | 449..251 | NEW |
| | | GPRmgl2_3 | 7312 | 436..130 | NEW |

| | | | | | |
|---|---|---|---|---|---|
| | | GPRmgl2_4 | 570 | 1093..98 | NEW |
| | | GPRmgl2_5 | 1251 | 39860..39180 | LLOJ001301 |
| | | GPRmgl3 | 876 | 43566..58097 | LLOJ009943 |
| | | GPRmgl4 | 325 | 2308..18954 | LLOJ008704 |
| | | GPRmgl5 | 5448 | 3299..24076 | LLOJ007663 |
| | **Rickets** | GPRrk | 1137 | 52088..18863 | LLOJ000822 |
| | **GABA(B) receptors** | GPRgbb1_1 | 854 | 1044..1262 | NEW |
| | | GPRgbb1_2 | 691 | 44983..52144 | LLOJ008323 |
| | | GPRgbb1_3 | 4840 | 4872..5021 | NEW |
| | | GPRgbb1_4 | 691 | 53857..63339 | LLOJ008323 |
| | | GPRgbb2_1 | 1066 | 7793..7527 | LLOJ000435 |
| | | GPRgbb2_2 | 776 | 3761..21370 | LLOJ008887 |
| | | GPRgbb3_1 | 190 | 42138..49130 | LLOJ003042 |
| | | GPRgbb3_2 | 931 | 1344..1279 | NEW |
| | | GPRgbb3_3 | 190 | 53213..64932 | LLOJ003042 |
| | **Orphan/Putative Class C GPCRs** | GPRorp1 | 44067 | 1306..25008 | LLOJ003399 |
| **(4) Class D- Atypical 7TM proteins** | **Bride of Sevenless** | GPRbos | 1 | 13099..88765 | LLOJ008818 |
| | **Frizzled** | GPRfz1_1 | 594 | 6150..21905 | LLOJ007599 |
| | | GPRfz1_2 | 3280 | 1080..1292 | NEW |
| | | GPRfz1_3 | 594 | 65687..65894 | NEW |
| | | GPRfz2 | 811 | 2303..393 | NEW |
| | | GPRfz4 | 199 | 106370..142852 | LLOJ003257 |
| | | GPRsmo1_1 | 1092 | 4984..11 | LLOJ000573 |
| | **Smoothened** | GPRsmo1_2 | 9442 | 1460..546 | NEW |
| | **Starry night** | GPRstn | 321 | 6789..11358 | LLOJ009854 |
| | **Orphan/Putative Class D GPCRs** | GPRorp1 | | | |
| | | GPRorp2 | | | |
| **(5) Short peptides/partial gene models** | | | | | |

**S24 Table.** MicroRNA annotation.

Number of genes in different RNA interference pathways from Dipteran insects.

| RNAi pathway | Gene name | *L.longipalpis* | *P.papatasi* | *D. melanogaster* | *A. aegypti* |
|---|---|---|---|---|---|
| siRNA | *Dcr-2* | 1 | 1 | 1 | 1 |
| | *r2d2* | 1 | NF | 1 | 1 |
| | *AGO2* | 1 | NF | 1 | 1 |
| | *loqs* | 1 | 1 | 1 | 1 |
| miRNA | *drosha* | 1 | 1 | 1 | 1 |
| | *pasha* | 1 | 1 | 1 | 1 |
| | *Dcr-1* | 1 | 1 | 1 | 1 |
| | *AGO1* | 2 | 1 | 1 | 2 |
| piRNA | *AGO3* | 1 | 1 | 1 | 1 |
| | *piwi/aub* | 4 | 2 | 2 | 7 |

NF-not found in current assembly.

Analysis of core RNAi genes in *Lu. longipalpis* compared to *D. melanogaster*.

| RNAi pathway | Gene name | *Drosophila* protein reference used for BLAST searches (Flybase reference) | *Drosophila* gene and protein length (Flybase) | *Lu. longipalpis* gene (Vectorbase ID) | *Lu. longipalpis* gene (Provisional gene name) | *Lutzomyia* gene/protein length (VectorBase) | Blast e-value | Identity | Scaffold number and position | Observations about genome assembly and gene models |
|---|---|---|---|---|---|---|---|---|---|---|
| siRNA | *Dcr-2* | Dcr-2-RA | 5684bp-1722aa | LLOJ006509 | *Dcr-2* | 4956bp / 1526aa | 0.0 | 32.7% | Scaffold466:6,387-22,877 | |
| | *r2d2* | r2d2-RA | 1786bp-311aa | LLOJ000693 | *r2d2* | 1129bp / 292aa | 2.00E-28 | 32.3% | Scaffold 1109:17,333-18,530 | Gene model requires revision (too many exons resulting in unexpected protein) |
| | *AGO2* | AGO2-RB | 4050bp-1214aa | LLOJ006148 | *AGO2* | 3383bp / 1086aa | 5.00E-155 | 40.5% | Scaffold 419:7,793-15,194 | |
| | *loqs* | loqs-RB/loqs-RD | 2017bp-465aa / 1334bp-359aa | LLOJ002667 | *loqs* | 7919bp / 389aa | 3E-132 / 2e-82 | 62.1% / 63.4% | Scaffold 172:2,714-17,598 | |
| miRNA | *drosha* | drosha-RA | 4290bp-1327aa | LLOJ001478 | *drosha* | 9867bp / 2438aa | 0.0 | 68.9% | Scaffold 13:248,698-262,799 | Possible duplication caused by an artifact in the assembly |

| | | | | | | | | | | |
|---|---|---|---|---|---|---|---|---|---|---|
| | *Dcr-1* | Dcr-1-RA | 6913bp-2249aa | LLOJ006865 | *Dcr-1* | 8054bp / 2243aa | 0.0 | 60.7% | Scaffold 50:215,363-224,716 | |
| | *AGO1** | AGO1-RC | 7528bp-984aa | LLOJ008700 | *AGO1-1* | 5026bp / 974aa | 0.0 | 94.2% | Scaffold 611:6,014-14,820 | |
| | | AGO1-RC | 7528bp-984aa | LLOJ007781 | *AGO1-2* | 2196bp / 731aa | 0.0 | 88.2% | Scaffold 611:15,891-18,957 | Possible duplication caused by an artifact in the assembly |
| | *pasha* | pasha-RA | 2847bp-642aa | LLOJ005101 | *pasha* | 2010bp / 669aa | 8.00E-157 | 59.1% | Scaffold 317:118,601-122,633 | |
| piRNA | *AGO3* | AGO3-RD | 2800bp-867aa | LLOJ007793 | *AGO3* | 5200bp / 1552aa | 0.0 | 42.4% | Scaffold 612:28,708-49,153 | Gene model requires revision (current gene too long) |
| | *piwi/aub*** | piwi-RA/aub-RA | 3073bp-843aa/2825bp/866aa | LLOJ008700 | *piwi-1* | 2843bp / 887aa | 0.0 | 45.7% / 45.7% | Scaffold 747:16,913-20,393 | |
| | | piwi-RA/aub-RA | 3073bp-843aa/2825bp/866aa | LLOJ000423 | *piwi-2* | 3078bp / 972aa | 0.0 | 43.1% / 43.1% | Scaffold 1063:19,681-25,005 | |
| | | piwi-RA/aub-RA | 3073bp-843aa/2825bp/866aa | LLOJ005362 | *piwi-3* | 4385bp / 1414aa | 0.0 | 43.2% / 43.2% | Scaffold 34:40,427-54,140 | Gene model requires revision (current gene too long) |
| | | piwi-RA/aub-RA | 3073bp-843aa/2825bp/866aa | LLOJ006685 | *piwi-4* | 2614pb / 865aa | 0.0 | 41.9% / 41.9% | Scaffold 487:44,683-48,080 | |

* 2 separate genes in the genome of *Lu. longipalpis* showed homology to Drosophila AGO1
**4 separate genes in the genome of *Lu. longipalpis* showed homology to Drosophila piwi/aub genes

Analysis of core RNAi genes in *P. paptasi* compared to *Lu. longipalpis*

| RNAi pathway | Gene name | *Lutzomyia* protein reference used for BLAST searches (Vectorbase) | *Lutzomyia* gene and protein length (Vectorbase) | *P. papatasi* gene (Vectorbase ID) | *P. papatasi* gene (Provisional gene name) | *P. papatasi* gene/protein length (VectorBase) | Blast e-value | Identity | Scaffold number and position | Observations about genome assembly and gene models |
|---|---|---|---|---|---|---|---|---|---|---|
| siRNA | *Dcr-2* | LLOJ006509 | 4956bp-1526aa | PPAI008140 | *Dcr-2* | 1644bp / 547 aa | 0.0 | 64.7% | Scaffold47738: 4,153-5,796 | Gene model requires revision (current gene too small) |
| | *r2d2* | LLOJ000693 | 1129bp-292aa | NF | | | | | | Gene missing from current genome assembly |
| | *AGO2* | LLOJ006148 | 3383bp-1086aa | NF | | | | | | Gene missing from current genome assembly |
| miRNA | *loqs* | LLOJ002667 | 7919bp-389aa | PPAI006255 | *loqs* | 1040bp / 218aa | 9.00E-151 | 87.7% | Scaffold4: 1,392,711-1,397,363 | |
| | *drosha* | LLOJ001478 | 9867bp-2438aa | PPAI000853 | *drosha* | 3249bp / 1082aa | 0.0 | 79.7% | Scaffold1219: 28,667-33,606 | |
| | *Dcr-1* | LLOJ006865 | 8054bp-2243aa | PPAI005440 | *Dcr-1* | 5588bp / 1839aa | 0.0 | 82.7% | Scaffold334: 13,905-25,891 | |
| | *AGO1** | LLOJ007780 / LLOJ007781 | 5026bp-974aa / 2196bp-731aa | PPAI005304 | *AGO1* | 1538bp / 477aa | 0.0 / 2E-173 | 94.3% / 100% | Scaffold3232: 1,621-10,587 | Gene model requires revision (current gene too small) |
| | *pasha* | LLOJ005101 | 2010bp-669aa | PPAI009756 | *pasha* | 1269bp / 422aa | 8.00E-154 | 79% | Scaffold63318: 2,547-3,815 | Gene model requires revision (current |

| | | | | | | | | | | |
|---|---|---|---|---|---|---|---|---|---|---|
| | | | | | | | | | | gene too small) |
| piRNA | *AGO3* | LLOJ007793 | 5200bp-1552aa | PPAI005870 | *AGO3* | 2391bp / 649aa | 0.0 | 74.2% | Scaffold3695:4,737-14,529 | Gene model requires revision (current gene too small) |
| | *piwi* ** | LLOJ008700 / LLOJ000423 / LLOJ005362 / LLOJ006685 | 2843bp-887aa / 3078bp-972aa / 4385bp-1414aa / 2614pb- 865aa | PPAI001776 | *piwi-1* | 2928bp / 975aa | 0.0 | 65.3% - 77.2% *** | Scaffold1532: 22,048-30,014 | |
| | | LLOJ008700 / LLOJ000423 / LLOJ005362 / LLOJ006685 | 2843bp-887aa / 3078bp-972aa / 4385bp-1414aa / 2614pb- 865aa | PPAI005745 | *piwi-2* | 1194bp / 393aa | 2e-173 - 0.0 *** | 60.6% - 76.8% *** | Scaffold3576: 753-1,946 | Gene model requires revision (current gene too small) |

NF-not found in current assembly
* The two putative AGO1 genes from *Lu. longipalpis* showed homology to the same gene in the *P. papatasi* genome
** The four putative piwi genes from *Lu. longipalpis* showed similarity to two genes in the *P. papatasi* genome
*** Range of values for all 4 piwi genes

**S25 Table.** Heat shock and hypoxia gene family annotation.

| Species | Gene Name | Gene Symbol | Lutzomyia Vectorbase ID | D. melanogaster | A. aegypti | A. gambiae | Revisions required |
|---|---|---|---|---|---|---|---|
| *Lu. Longipalpis* | Adenosine deaminase acting on RNA | Adar | LLOJ008903 | FBgn0026086 | AAEL002522 | AGAP000185 | N |
| | Checkpoint suppressor homolog | Ches-1 | LLOJ006903 | FBgn0029504 | AAEL001536 | AGAP010023 | Major |
| | Cyclic nucleotide gated ion channel | Cng | LLOJ008345 | FBgn0261612 | AAEL007345 | AGAP009050 | Major |
| | Cyclin dependent kinase 9 | Cdk9 | LLOJ005668 | FBgn0019949 | AAEL013002 | AGAP008541 | N |
| | Cyclin T-like | CycT | LLOJ003789 | FBgn0025455 | AAEL004839 | AGAP006678 | Minor |
| | DNA-directed RNA polymerase II subunit | RPII33 | LLOJ003173 | FBgn0026373 | AAEL008755 | AGAP010321 | N |
| | DNAJ-like 1 | DNAJ1 | LLOJ002180 | FBgn0031322 | AAEL006899 | AGAP010239 | N |
| | DNAJ-like 2 | DNAJ2 | LLOJ001204 | FBgn0038145 | AAEL005165 | AGAP005981, AGAP000008 | N |
| | DNAJ-like 3 | DNAJ3 | LLOJ008501 | FBgn0031256 | AAEL009946, AAEL013020 | AGAP008327 | N |
| | DNAJ-like 4 | DNAJ4 | LLOJ005042 | FBgn0263106 | AAEL003588 | AGAP007107 | N |
| | DNAJ-like 5 | DNAJ5 | LLOJ007103 | FBgn0038145 | AAEL005165 | AGAP005981, AGAP000008 | Minor |
| | Dynamin | dyn | LLOJ010016 | FBgn0003392 | AAEL007288 | AGAP003018 | Major |
| | eas-ethanolamine kinase | eas | LLOJ000742 | FBgn0000536 | AAEL009765 | AGAP000010 | Major |
| | Ecdysone induced protein | Eip | LLOJ003014 | FBgn0264490 | AAEL004578 | AGAP004494 | Major |
| | eIF2B-beta | eIF2B-beta | LLOJ003864 | FBgn0024996 | AAEL005825 | AGAP007097 | N |
| | eIF2B-gamma | eIF2B-gamma | LLOJ005927 | FBgn0034029 | AAEL011442 | AGAP005210 | N |
| | enoyl-CoA-hydratase | ech | LLOJ002903 | FBgn0038049 | AAEL003202 | AGAP001715 | N |
| | Exportin-1 | XPO-1 | LLOJ001426 | FBgn0020497 | AAEL001484 | AGAP009929 | Major |
| | Glutathione s transferase | GstD | LLOJ004462 | FBgn0001149 | AAEL001078, AAEL001061 | AGAP004164 | N |
| | Guanylyl cyclase 89D | Gyc89D | LLOJ006750 | FBgn0038435, FBgn0038436 | AAEL013026, AAEL014569 | AGAP004564 | N |
| | Guanylyl Cyclase at 88E | Gyc88E | LLOJ000691 | FBgn0038295 | AAEL013328 | AGAP001985 | N |
| | hairy | h | LLOJ000067 | FBgn0001168 | AAEL005480, AAEL011943 | AGAP006699 | N |
| | Hangover | hang | LLOJ010026 | FBgn0026575 | AAEL011847 | AGAP010021 | Major |
| | Heat shock factor | Hsf | LLOJ009750 | FBgn0001222 | AAEL010319 | AGAP011082 | Minor |
| | Heat shock protein 22 | Hsp22 | LLOJ005427 | FBgn0001223 | AAEL013350 | AGAP005548 | Minor |
| | Heat shock protein 23 | Hsp23 | LLOJ007367 | FBgn0001224 | AAEL013350 | AGAP005548 | N |
| | Heat shock protein 26 | Hsp26 | LLOJ006489 | FBgn0001225 | AAEL013350 | AGAP005548 | N |
| | Heat shock protein 27 | Hsp27 | LLOJ007364 | FBgn0001226 | AAEL013350, AAEL013345, AAEL013339 | AGAP005547, AGAP005548, AGAP007158, AGAP007159 | N |
| | Heat shock protein 70A | Hsp70A | LLOJ006488 | FBgn0013275 | AAEL801273 | AGAP004944 | N |
| | Heat shock protein 70Ba | Hsp70Ba | LLOJ005733 | FBgn0042854 | AAEL801273 | AGAP004944 | N |

| | | | | | | |
|---|---|---|---|---|---|---|
| Heat shock protein 70Bb | Hsp70Bb | LLOJ005732 | FBgn0013278 | AAEL801273 | AGAP004944 | N |
| Heat shock protein 70C | Hsp70C | LLOJ007339 | FBgn0026418 | AAEL017315 | AGAP010331 | N |
| Heat shock protein 83 | Hsp83 | LLOJ004224 | FBgn0001233 | AAEL011708, AAEL014843, AAEL014845, AAEL011704 | AGAP006958 | Major |
| Heat shock protein cognate 70-1 | Hsc70-1 | LLOJ008388 | FBgn0001216 | AAEL018061 | AGAP004944 | N |
| Heat shock protein cognate 70-2 | Hsc70-2 | LLOJ001213 | FBgn0001217 | AAEL018062 | AGAP004944 | N |
| Heat shock protein cognate 70-3 | Hsc70-3 | LLOJ002147 | FBgn0001218 | AAEL017349 | AGAP004192 | N |
| Heat shock protein cognate 70-4 | Hsc70-4 | LLOJ006821 | FBgn0001219 | AAEL018061 | AGAP002076 | N |
| Heat Shock Protein Cognate 70-5 | Hsc70-5 | LLOJ003101 | FBgn0001220 | | AGAP010876 | N |
| Heterogeneous nuclear ribonucleoprotein | Hrb87F | LLOJ005415 | FBgn0004237 | AAEL010467 | AGAP002374 | Minor |
| Hormone receptor 4 | Hr4 | LLOJ003352 | FBgn0023546 | AAEL005850 | AGAP004693 | Major |
| Hypoxia inducible factor-prolyl hydroxylase | hph | LLOJ007742 | FBgn0264785 | AAEL002798 | AGAP003523 | N |
| Jun-N-Terminal Kinase | Jnk | LLOJ005677 | FBgn0000229 | AAEL008634 | AGAP009461 | N |
| Locomotion defects | loco | LLOJ001915 | FBgn0020278 | AAEL007358 | AGAP002411 | Major |
| Lon Protease | Lon | LLOJ008442 | FBgn0036892 | AAEL006474 | AGAP010451 | N |
| Menin | Mnn | LLOJ002837 | FBgn0031885 | AAEL005686, AAEL005553 | AGAP008130 | N |
| Methusela | mth | LLOJ002123 | FBgn0023000 | AAEL010982 | AGAP006215, AGAP006216 | N |
| Methyltransferase 2 | Mt2 | LLOJ001788 | FBgn0028707 | AAEL006166 | AGAP012836, AGAP004101 | Major |
| Mitochondrial import receptor subunit Tom40 | Tom40 | LLOJ006852 | FBgn0016041 | AAEL007001 | AGAP007871 | Minor |
| Mitochondrial inner membrane import translocase | Tim9 | LLOJ005065 | FBgn0027358 | AAEL002486 | AGAP007167 | Minor |
| Mitogen activated protein kinase kinase | Mekk | LLOJ007004 | FBgn0024329 | AAEL010466 | AGAP002371 | Major |
| Mixed lineage kinase | Mlk | LLOJ002365 | FBgn0030018 | AAEL012999 | AGAP002710 | N |
| Multi drug resistance protein 4 | Mrp4 | LLOJ000639 | FBgn0263316 | AAEL013854, AAEL013567 | AGAP006427 | N |
| Multidrug resistance protein 49 | Mdr49 | LLOJ000437 | FBgn0004512 | AAEL010379 | AGAP005639 | Minor |
| p38 map kinase | p38 | LLOJ010513 | FBgn0024846 | AAEL008379 | AGAP012148 | Major |
| Painless transient receptor potential channel protein | pain | LLOJ006694 | FBgn0060296 | AAEL006835 | AGAP012995, AGAP013366, AGAP013463 | Major |
| Para-like sodium channel | para | LLOJ007479 | FBgn0264255 | AAEL008297 | AGAP004707 | Major |
| Phosphoglycerate mutase family member | Pgam | LLOJ005892 | FBgn0023517 | AAEL006915 | AGAP002535 | N |
| Phosphomannomutase | PMM | LLOJ005377 | FBgn0036300 | AAEL009497 | AGAP011353 | N |
| Poly(ADP-ribose) glycohydrolase | Parg | LLOJ001887 | FBgn0023216 | AAEL001470 | AGAP000589 | N |
| Prohibitin | Phb | LLOJ002103 | FBgn0002031 | AAEL009345 | AGAP009323 | Minor |

| | | | | | | | |
|---|---|---|---|---|---|---|---|
| | Protein tumerous imaginal disc | Ptid | LLOJ001601 | FBgn0002174 | AAEL011055, AAEL007728 | AGAP007565 | Minor |
| | Protein tyrosine phosphatase 61F | Ptp61F | LLOJ008161 | FBgn0267487 | AAEL001919 | AGAP005352 | N |
| | Ras which interacts with calmodulin | Ric | LLOJ002204 | FBgn0265605 | AAEL009180, AAEL015232 | AGAP005159 | N |
| | Rutebaga adenylate cyclase | rut | LLOJ004622 | FBgn0003301 | AAEL017420, AAEL009022 | AGAP000727 | Major |
| | Sentrin specific protease | Ulp1 | LLOJ005857 | FBgn0027603 | AAEL008952 | n/a | Minor |
| | Similar-like Hypoxia inducible factor | sima | LLOJ000925 | FBgn0266411 | AAEL015383,AAEL001097 | AGAP002942 | N |
| | Sodium dependent phosphate cotransporter | SDPT | LLOJ004400 | FBgn0010651 | AAEL001209 | AGAP008931 | N |
| | Sorting nexin 33 isoform X1 | SH3PX1 | LLOJ000862 | FBgn0040475 | AAEL003416 | AGAP011751 | N |
| | Synapsin | syn | LLOJ004480 | FBgn0004575 | AAEL004549 | AGAP003318 | N |
| | Tango protein | tgo | LLOJ003291 | FBgn0264075 | AAEL010343 | AGAP009748 | Major |
| | Technical knockout | tko | LLOJ009320 | FBgn0003714 | AAEL008169 | AGAP013102 | N |
| | Temperature induced paralytic E sodium channel | tipE | LLOJ009582 | FBgn0003710 | AAEL002127 | AGAP002820 | N |
| | Thiodoxin reductase 1 | Trxr-1 | LLOJ007938 | FBgn0020653 | AAEL002886 | AGAP000565 | N |
| | THO-complex subunit-1 | THO1 | LLOJ002135 | FBgn0037382 | AAEL002563 | AGAP004070 | N |
| | THO complex subunit 2 | tho2 | LLOJ006531 | FBgn0031390 | AAEL001716 | AGAP008236 | N |
| | Transient receptor potential A1 | TrpA1 | LLOJ004961 | FBgn0035934 | AAEL001268, AAEL009419 | AGAP004863 | N |
| | Translocase of the inner mitochondrial membrane 44 | Timm44 | LLOJ009540 | FBgn0038683 | AAEL008412, AAEL012845 | AGAP000291 | N |
| | Tuberous sclerosis complex 1 | Tsc1 | LLOJ002399 | FBgn0026317 | AAEL002548 | AGAP003445 | Minor |
| | Tuberous sclerosis complex 2 | Tsc2 | LLOJ004899 | FBgn0005198 | AAEL007712 | AGAP011123 | Major |
| | Unc-45 | unc-45 | LLOJ005351 | FBgn0010812 | AAEL009168, AAEL010948 | AGAP003727 | N |
| | Upf1 regulator of nonsense transcripts | Upf1 | LLOJ009843 | FBgn0030354 | AAEL011817 | AGAP001133 | Minor |
| | Vacuolar H[+] ATPase subunit | Vha68-1 | LLOJ009754 | FBgn0265262 | AAEL008787 | AGAP003153 | N |
| | Vannin-like protein | Vnnl | LLOJ008292 | FBgn0040069 | AAEL002681, AAEL006034, AAEL006023, AAEL015162 | AGAP010733 | N |
| *P. papatasi* | Adenosine deaminase acting on RNA | adar | PPAI010104 | FBgn0026086 | AAEL002522 | AGAP000185 | Major |
| | Altered disjunction-like dual specificity protein kinase | ald | PPAI003159 | FBgn0000063 | AAEL011118 | AGAP002075 | N |
| | Archipelago | ago | PPAI002855 | FBgn0041171 | AAEL015522, AAEL011215 | AGAP005359 | Major |
| | Cyclic AMP-dependent activating transcription factor-2 | atf-2 | PPAI009946 | FBgn0265193 | AAEL013261 | AGAP008416 | Minor |
| | Cyclic nucleotide gated channel | Cng | PPAI008474 | FBgn0261612 | AAEL007345 | AGAP009050 | N |
| | Cyclin dependent kinase 9 | cdk9 | PPAI000097 | FBgn0019949 | AAEL013002 | AGAP008541 | N |

| | | | | | | |
|---|---|---|---|---|---|---|
| Cyclin-T like | CycT | PPAI001177 | FBgn0025455 | AAEL004839 | AGAP006678 | Major |
| DNA directed RNA polymerase II subunit 33 | RpII33 | PPAI009645 | FBgn0026373 | AAEL008755 | AGAP010321 | Major |
| DnaJ-like-1 | DnaJ-1 | PPAI005370 | FBgn0031322 | AAEL006899 | AGAP010239 | N |
| DnaJ-like-2 | DnaJ-2 | PPAI006146 | FBgn0038145 | AAEL005165 | AGAP005981, AGAP000008 | N |
| DnaJ-like-3 | DnaJ-3 | PPAI008141 | FBgn0031256 | AAEL009946, AAEL013020 | AGAP008327 | N |
| Dynamin | dyn | PPAI005546 | FBgn0000536 | AAEL009765 | AGAP000010 | N |
| eas-ethanolamine kinase | eas | PPAI009322 | FBgn0000536 | AAEL009765 | AGAP000010 | N |
| Ecdysone induced protein | Eip | PPAI001250 | FBgn0264490 | AAEL004578 | AGAP004494 | Minor |
| eIF2B-beta | eIF2B-beta | PPAI004100 | FBgn0024996 | AAEL005825 | AGAP007097 | N |
| eIF2B-gamma | eIF2B-gamma | PPAI009373 | FBgn0034029 | AAEL011442 | AGAP005210 | N |
| enoyl-CoA-hydratase | ech | PPAI008583 | FBgn0038049 | AAEL003202 | AGAP001715 | N |
| Exportin-1 | xpo1 | PPAI001648 | FBgn0020497 | AAEL001484 | AGAP009929 | N |
| Glutathione S transferase | GstD | PPAI010868 | FBgn0001149 | AAEL001078, AAEL001061 | AGAP004164 | N |
| Guanylyl Cyclase 88E | Gyc88E | PPAI004038 | FBgn0038295 | AAEL013328 | AGAP001985 | N |
| Guanylyl Cyclase 89D | Gyc89D | PPAI000706 | FBgn0038435, FBgn0038436 | AAEL013026, AAEL014569 | AGAP004564 | Minor |
| Hairy | h | PPAI006478 | FBgn0001168 | AAEL005480, AAEL011943 | AGAP006699 | N |
| Hangover | hang | PPAI009131 | FBgn0026575 | AAEL011847 | AGAP010021 | Major |
| Heat shock factor | Hsf | PPAI009263 | FBgn0001222 | AAEL010319 | AGAP011082 | N |
| Heat shock protein 23 | Hsp23 | PPAI005922 | FBgn0001224 | AAEL013350 | AGAP005548 | Major |
| Heat shock protein 26 | Hsp26 | PPAI008863 | FBgn0001225 | AAEL013350 | AGAP005548 | N |
| Heat shock protein 27 | Hsp27 | PPAI003244 | FBgn0001226 | AAEL013350, AAEL013345, AAEL013339 | AGAP005547, AGAP005548, AGAP007158, AGAP007159 | Minor |
| Heat shock protein 67B | Hsp67B | PPAI009632 | FBgn0001228 | AAEL005759, AAEL005884, AAEL005890 | AGAP013228 | N |
| Heat shock protein 83 | Hsp83 | PPAI005952 | FBgn0001233 | AAEL011708, AAEL014843, AAEL014845, AAEL011704 | AGAP006958 | Major |
| Heat shock protein cognate 70-2 | Hsc70-2 | PPAI010639 | FBgn0001217 | AAEL018062 | AGAP004944 | N |
| Heat shock protein cognate 70-3 | Hsc70-3 | PPAI008472 | FBgn0001218 | AAEL017349 | AGAP004192 | N |
| Heatshock protein 60 | Hsp60 | PPAI003141 | FBgn0015245 | AAEL011584 | AGAP004002 | N |
| Heatshock protein cognate 70-1 | Hsc70-1 | PPAI000496 | FBgn0001216 | AAEL018061 | AGAP004944 | N |
| Heterogeneous ribonucleic protein 87f | Hrb87F | PPAI003801 | FBgn0004237 | AAEL010467 | AGAP002374 | Major |
| Holocarboxylase synthetase | Hcs | PPAI001335 | FBgn0037332 | AAEL009340 | AGAP001481 | N |
| Hormone receptor 4 | Hr4 | PPAI009449 | FBgn0023546 | AAEL005850 | AGAP004693 | Major |

| | | | | | | |
|---|---|---|---|---|---|---|
| Hypoxia inducible factor-prolyl hydroxylase | Hph | PPAI009790 | FBgn0264785 | AAEL002798 | AGAP003523 | N |
| Inactive | iav | PPAI004901 | FBgn0086693 | AAEL009258 | AGAP000413 | N |
| Jun-N-terminal Kinase | jnk | PPAI005249 | FBgn0000229 | AAEL008634 | AGAP009461 | N |
| Leucine zipper EF-hand containing transmembrane protein 1-ortholog | Letm1 | PPAI003231 | FBgn0019886 | AAEL000485 | AGAP003296 | Minor |
| Locomotion defects | loco | PPAI009736 | FBgn0020278 | AAEL007358 | AGAP002411 | N |
| Menin | Mnn | PPAI001130 | FBgn0031885 | AAEL005686, AAEL005553 | AGAP008130 | N |
| Methusela | mth | PPAI008623 | FBgn0023000 | AAEL010982 | AGAP006215, AGAP006216 | Minor |
| Methyltransferase 2 | mt2 | PPAI010989 | FBgn0028707 | AAEL006166 | AGAP012836, AGAP004101 | N |
| Mitochondrial inner membrane import translocase | Tim9 | PPAI007284 | FBgn0027358 | AAEL002486 | AGAP007167 | Minor |
| Mitogen activated protein kinase kinase | Mekk | PPAI009824 | FBgn0024329 | AAEL010466 | AGAP002371 | Major |
| Mixed lineage kinase | Mlk | PPAI001629 | FBgn0030018 | AAEL012999 | AGAP002710 | Minor |
| Multidrug Resistance protein 1 | mrp1 | PPAI000774 | FBgn0032456 | AAEL004743 | AGAP009835 | N |
| Multidrug resistance protein 49 | Mdr49 | PPAI006063 | FBgn0004512 | AAEL010379 | AGAP005639 | Major |
| Myocardin-related transcription factor | Mrtf | PPAI007609 | FBgn0052296 | AAEL007304 | AGAP001472 | Major |
| p38 map kinase | p38 | PPAI008377 | FBgn0024846 | AAEL008379 | AGAP012148 | Minor |
| Painless | pain | PPAI003852 | FBgn0060296 | AAEL006835 | AGAP012995, AGAP013366, AGAP013463 | N |
| Para-like sodium channel | para | PPAI003387 | FBgn0264255 | AAEL008297 | AGAP004707 | N |
| Phosphoglycerate mutase family member | Pgam | PPAI001812 | FBgn0023517 | AAEL006915 | AGAP002535 | Minor |
| Phosphomannomutase | PMM | PPAI004644 | FBgn0036300 | AAEL009497 | AGAP011353 | Major |
| Poly (ADP-ribose) glycohydrolase | Parg | PPAI009486 | FBgn0023216 | AAEL001470 | AGAP000589 | Minor |
| Prohibitin | l(2)37Cc | PPAI008827 | FBgn0002031 | AAEL009345 | AGAP009323 | N |
| Protein tumerous imaginal disc | Ptid | PPAI003201 | FBgn0002174 | AAEL011055, AAEL007728 | AGAP007565 | Minor |
| Protein-Tyrosine phosphatase 61F | Ptp61F | PPAI006334 | FBgn0267487 | AAEL001919 | AGAP005352 | N |
| Ras which interacts with Calmodulin | Ric | PPAI009427 | FBgn0265605 | AAEL009180, AAEL015232 | AGAP005159 | Minor |
| RNA binding musashi-like protein | msi | PPAI009515 | FBgn0011666 | AAEL006057 | AGAP001993 | Major |
| Rutebaga | rut | PPAI001426 | FBgn0003301 | AAEL017420, AAEL009022 | AGAP000727 | Major |
| Sentrin specific protease | Ulp1 | PPAI004288 | FBgn0027603 | AAEL008952 | n/a | N |
| Similar-like hypoxia inducible factor | sima | PPAI002345 | FBgn0266411 | AAEL015383,AAEL001097 | AGAP002942 | Major |
| Sorting nexin 33 isoform X1 | SH3PX1 | PPAI002620 | FBgn0040475 | AAEL003416 | AGAP011751 | N |

| | | | | | | |
|---|---|---|---|---|---|---|
| Synapsin | syn | PPAI003266 | FBgn0004575 | AAEL004549 | AGAP003318 | Major |
| Tango | tgo | PPAI009623 | FBgn0264075 | AAEL010343 | AGAP009748 | N |
| Technical knockout | tko | PPAI000676 | FBgn0003714 | AAEL008169 | AGAP013102 | N |
| Temperature induced paralytic E sodium channel | tipE | PPAI002431 | FBgn0003710 | AAEL002127 | AGAP002820 | Major |
| THO complex subunit 1 | THO1 | PPAI006037 | FBgn0037382 | AAEL002563 | AGAP004070 | N |
| THO complex subunit 2 | tho2 | PPAI001341 | FBgn0031390 | AAEL001716 | AGAP008236 | N |
| Transient receptor potential A1 | TrpA1 | PPAI004036 | FBgn0035934 | AAEL001268, AAEL009419 | AGAP004863 | Major |
| Tuberous sclerosis complex 1 | Tsc1 | PPAI007877 | FBgn0026317 | AAEL002548 | AGAP003445 | Major |
| Tuberous sclerosis complex 2 | Tsc2 | PPAI010500 | FBgn0005198 | AAEL007712 | AGAP011123 | Major |
| Unc-45 | unc-45 | PPAI010371 | FBgn0010812 | AAEL009168, AAEL010948 | AGAP003727 | N |
| Up frameshift suppressor 1 | Upf1 | PPAI010838 | FBgn0030354 | AAEL011817 | AGAP001133 | N |
| Vacuolar H[+] ATPase subunit | Vha68-1 | PPAI004876 | FBgn0265262 | AAEL008787 | AGAP003153 | N |
| Vanin-like protein | Vnnl | PPAI008082 | FBgn0040069 | AAEL002681, AAEL006034, AAEL006023, AAEL015162 | AGAP010733 | N |

**S26 Table.** Cuticular protein gene family annotation.

| Species | Vectorbase Gene ID | Gene | Symbol | Family | Required Revisions | *D. melanogaster* | *A. aegypi* | *A. gambiae* |
|---|---|---|---|---|---|---|---|---|
| *Lu. Longipalpis* | LLOJ000008 | CG13643 | CG13643 | CPAP1 | Y | FBpp0302610 | AAEL006933 | AGAP028105 |
| | LLOJ000231 | CG13676 | CG13676 | CPAP1 | Y | FBpp0305000 | AAEL004279 | AGAP005489 |
| | LLOJ000690 | CG32036 | CG32036 | CPAP1 | Y | FBpp0113017 | AAEL005622 | AGAP001203 |
| | LLOJ000711 | CG14304 | CG14304 | CPAP1 | Y | FBpp0112989 | AAEL011586 | AGAP009479 |
| | LLOJ001937 | CG12009 | CG12009 | CPAP1 | Y | FBpp0300633 | AAEL012013 | AGAP005586 |
| | LLOJ002193 | CG14608 | CG14608 | CPAP1 | Y | FBpp0292311 | AAEL011586 | AGAP010302 |
| | LLOJ002468 | CG14880 | CG14880 | CPAP1 | Y | FBpp0082770 | AAEL006201 | AGAP003751 |
| | LLOJ006484 | CG14608 | CG14608 | CPAP1 | Y | FBpp0292311 | AAEL011586 | AGAP010302 |
| | LLOJ006974 | CG32036 | CG32036 | CPAP1 | Y | FBpp0113017 | AAEL005622 | AGAP001203 |
| | LLOJ007841 | CG6757 | CG6757 | CPAP1 | Y | FBpp0303036 | AAEL003496 | AGAP007089 |
| | LLOJ002676 | obstruct-A | obst | CPAP3 | Y | FBpp0297663 | AAEL012245 | AGAP000989 |
| | LLOJ005620 | CG10725 | CG10725 | CPAP3 | Y | FBpp0089010 | AAEL002625 | AGAP028110 |
| | LLOJ007615 | Gasp | Gasp | CPAP3 | N | FBpp0111835 | AAEL013766 | AGAP003308 |
| | LLOJ009444 | CG10725 | CG10725 | CPAP3 | Y | FBpp0089010 | AAEL002625 | AGAP028110 |
| | LLOJ001501 | Cuticular Protein | CG8543 | CPF | Y | FBpp0076573 | AAEL000890 | AGAP001664 |
| | LLOJ008723 | Cuticular Protein | CG8541 | CPF | Y | FBpp0076572 | AAEL011068 | AGAP004690 |
| | LLOJ004690 | CG13042 | CG13042 | CPLCA | Y | FBpp0075191 | AAEL001704 | AGAP006148 |
| | LLOJ004691 | CG13042 | CG13042 | CPLCA | Y | FBpp0075191 | AAEL001704 | AGAP006148 |
| | LLOJ006190 | CG34462 | CG34462 | CPR_Uncl | Y | FBpp0111680 | AAEL003269 | AGAP009162 |
| | LLOJ008765 | Cuticular Protein 67B | Cpr67B | CPR_Uncl | Y | FBpp0076183 | AAEL004588 | AGAP006497 |
| | LLOJ002517 | Cuticular Protein 62Bb | Cpr62Bb | RR-1 | Y | FBpp0305097 | AAEL011045 | AGAP006829 |
| | LLOJ002518 | Cuticular Protein 62Bb | Cpr62Bb | RR-1 | Y | FBpp0305097 | AAEL011043 | AGAP006830 |
| | LLOJ003218 | Cuticular Protein 65Av | Cpr65Av | RR-1 | Y | FBpp0076732 | AAEL013515 | AGAP005996 |
| | LLOJ003219 | Cuticular Protein 65Av | Cpr65Av | RR-1 | Y | FBpp0076732 | AAEL011444 | AGAP005996 |
| | LLOJ004217 | Cuticular Protein 62Bb | Cpr62Bb | RR-1 | Y | FBpp0305097 | AAEL011045 | AGAP006829 |
| | LLOJ004632 | Cuticular Protein 47Ea | Cpr47Ea | RR-1 | Y | FBpp0087286 | AAEL014419 | AGAP009879 |
| | LLOJ005050 | Cuticular Protein 57A | Cpr57A | RR-1 | Y | FBpp0289081 | AAEL000419 | AGAP002726 |
| | LLOJ005175 | diskette | dik | RR-1 | N | FBpp0074284 | AAEL004675 | AGAP010459 |
| | LLOJ005497 | Pupal Cuticular Protein | Lcp65Av | RR-1 | Y | FBpp0076732 | AAEL013515 | AGAP006001 |
| | LLOJ005501 | Cuticular Protein 78E | Cpr78E | RR-1 | Y | FBpp0078117 | AAEL002440 | AGAP006006 |

| | | | | | | | |
|---|---|---|---|---|---|---|---|
| LLOJ005502 | Cuticular Protein 12A | Cpr12A | RR-1 | Y | FBpp0073615 | AAEL002452 | AGAP006009 |
| LLOJ005503 | Cuticular Protein 47Eb | Cpr47Eb | RR-1 | Y | FBpp0033598 | AAEL015460 | AGAP009876 |
| LLOJ005504 | Larval Cuticular Protein | Lcp65Ac | RR-1 | Y | FBpp0076700 | AAEL011444 | AGAP006003 |
| LLOJ005505 | Cuticular Protein 65Az | Cpr65Az | RR-1 | Y | FBpp0076704 | AAEL001735 | AGAP006013 |
| LLOJ005696 | Cuticular Protein 65Ec | Cpr65Ec | RR-1 | Y | FBpp0035737 | AAEL008285 | AGAP005459 |
| LLOJ006053 | Cuticular Protein 50Cb | Cpr50Cb | RR-1 | Y | FBpp0086756 | AAEL008764 | AGAP012466 |
| LLOJ006193 | Cuticular Protein 49Ae | Cpr49Ae | RR-1 | Y | FBpp0087031 | AAEL003259 | AGAP009871 |
| LLOJ006195 | Cuticular Protein 49Ab | Cpr49Ab | RR-1 | Y | FBpp0087025 | AAEL003235 | AGAP009874 |
| LLOJ006197 | Cuticular Protein 49Aa | Cpr49Aa | RR-1 | Y | FBpp0112271 | AAEL003272 | AGAP009876 |
| LLOJ006199 | Cuticular Protein 47Ee | Cpr47Ee | RR-1 | Y | FBpp0087215 | AAEL014416 | AGAP012728 |
| LLOJ007167 | Cuticular Protein 72Eb | Cpr72Eb | RR-1 | Y | FBpp0075182 | AAEL011458 | AGAP006931 |
| LLOJ008046 | Cuticular Protein | Cpr | RR-1 | Y | N/A | AAEL009005 | AGAP006848 |
| LLOJ007438 | Cuticular Protein 72Eb | Cpr72Eb | RR-2 | Y | FBpp0075141 | AAEL013030 | AGAP006931 |
| LLOJ007439 | Cuticular Protein 92F | Cpr92F | RR-2 | Y | FBpp0083327 | AAEL015119 | AGAP000047 |
| LLOJ000190 | Cuticular Protein | Cpr | RR-2 | Y | N/A | AAEL012090 | AGAP000745 |
| LLOJ000400 | Cuticular Protein | Cpr | RR-2 | Y | N/A | AAEL008997 | AGAP006848 |
| LLOJ000603 | Cuticular Protein 97Ea | Cpr97Ea | RR-2 | Y | FBpp0084506 | AAEL003027 | AGAP000345 |
| LLOJ000829 | Cuticular Protein 50Ca | Cpr50Ca | RR-2 | Y | FBpp0086755 | AAEL008759 | AGAP010717 |
| LLOJ001386 | CG13670 | CG13670 | RR-2 | Y | FBpp0076431 | AAEL007739 | AGAP006321 |
| LLOJ002420 | Cuticular Protein 76Bb | Cpr76Bb | RR-2 | Y | FBpp0074702 | AAEL009005 | AGAP006866 |
| LLOJ002421 | Cuticular Protein 76Bc | Cpr76Bc | RR-2 | Y | FBpp0304993 | AAEL008970 | AGAP006867 |
| LLOJ002877 | Cuticular Protein 47Ef | Cpr47Ef | RR-2 | Y | FBpp0291859 | AAEL004951 | AGAP010887 |
| LLOJ004294 | Cuticular Protein | Cpr | RR-2 | Y | N/A | AAEL014983 | AGAP006865 |
| LLOJ005682 | Cuticular Protein 66D | Cpr66D | RR-2 | Y | FBpp0076294 | AAEL003112 | AGAP006261 |
| LLOJ005712 | Cuticular Protein 56F | Cpr56F | RR-2 | Y | FBpp0085613 | AAEL008765 | AGAP012462 |
| LLOJ005917 | resillin | resillin | RR-2 | N | FBpp0086175 | AAEL008754 | AGAP012487 |
| LLOJ005918 | resillin | resillin | RR-2 | N | FBpp0086175 | AAEL008754 | AGAP012487 |
| LLOJ006524 | CG43367 | CG43367 | RR-2 | Y | FBpp0298311 | AAEL002114 | AGAP003389 |
| LLOJ006525 | Cuticular Protein | Cpr | RR-2 | Y | N/A | AAEL002092 | AGAP003385 |
| LLOJ006537 | Cuticular Protein 84Ag | Ccp84Ag | RR-2 | Y | FBpp0081187 | AAEL009803 | AGAP008960 |
| LLOJ006538 | Cuticular Protein 64Ad | Cpr64Ad | RR-2 | Y | FBpp0073079 | AAEL009803 | AGAP008960 |
| LLOJ007166 | Cuticular Protein 72Ec | Cpr72Ec | RR-2 | Y | FBpp0075182 | AAEL013030 | AGAP006931 |

| | | | | | | | | |
|---|---|---|---|---|---|---|---|---|
| | LLOJ007350 | Pupal Cuticular Protein | Lcp65Ad | RR-2 | Y | FBpp0305758 | AAEL002449 | AGAP002613 |
| | LLOJ008045 | Cuticular Protein 76Bc | Cpr76Bc | RR-2 | Y | FBpp0304993 | AAEL008970 | AGAP006867 |
| | LLOJ009584 | Cuticular Protein | Cpr | RR-2 | Y | N/A | AAEL002092 | AGAP003385 |
| | LLOJ009609 | Cuticular Protein | Cpr | RR-2 | Y | N/A | AAEL014973 | AGAP006858 |
| | LLOJ009610 | Cuticular Protein | Cpr | RR-2 | Y | N/A | AAEL009005 | AGAP006856 |
| | LLOJ009612 | Cuticular Protein | Cpr | RR-2 | Y | N/A | AAEL008979 | AGAP006864 |
| | LLOJ009893 | Cuticular Protein 92A | Cpr92A | RR-2 | Y | FBpp0083221 | AAEL009803 | AGAP001668 |
| | LLOJ009933 | Cuticular Protein 73D | Cpr73D | RR-2 | Y | FBpp0112089 | AAEL006737 | AGAP006369 |
| | LLOJ000600 | Tweedle T | TwdlT | TWDL | Y | FBpp0084471 | AAEL017015 | AGAP000352 |
| | LLOJ004281 | Tweedle T | TwdlT | TWDL | Y | FBpp0084471 | AAEL017015 | AGAP000352 |
| | LLOJ006105 | Tweedle E | TwdlE | TWDL | Y | FBpp0079185 | AAEL017015 | AGAP000352 |
| | LLOJ009522 | Tweedle Beta | TwdlBeta | TWDL | Y | FBpp0087168 | AAEL017015 | AGAP000352 |
| *P. papatasi* | PPAI001796 | CG8192 | CG8192 | CPAP1 | Y | FBpp0086487 | AAEL001553 | AGAP007613 |
| | PPAI003916 | CG14607 | CG14607 | CPAP1 | Y | FBpp0292310 | AAEL014445 | AGAP010302 |
| | PPAI004179 | CG32036 | CG32036 | CPAP1 | Y | FBpp0113017 | AAEL005622 | AGAP001203 |
| | PPAI006757 | CG14304 | CG14304 | CPAP1 | Y | FBpp0112989 | AAEL011586 | AGAP009479 |
| | PPAI006890 | CG14301 | CG14301 | CPAP1 | Y | FBpp0288638 | AAEL013117 | AGAP009480 |
| | PPAI006920 | CG5767 | CG5756 | CPAP1 | Y | FBpp0303036 | AAEL003496 | AGAP007089 |
| | PPAI008794 | CG14608 | CG14608 | CPAP1 | Y | FBpp0292311 | AAEL011586 | AGAP010302 |
| | PPAI010490 | CG13643 | CG13643 | CPAP1 | Y | FBpp0302610 | AAEL007483 | AGAP002052 |
| | PPAI001263 | obstructor-E | obst-E | CPAP3 | Y | FBpp0078794 | AAEL011897 | AGAP009405 |
| | PPAI001604 | obstructor-B | obst-B | CPAP3 | Y | FBpp0079524 | AAEL013222 | AGAP009790 |
| | PPAI004716 | Gasp | Gasp | CPAP3 | Y | FBpp0078268 | AAEL013766 | AGAP003308 |
| | PPAI004741 | Peritrophin-A | Peritrophin-A | CPAP3 | Y | FBpp0077028 | AAEL009585 | AGAP000986 |
| | PPAI004749 | Peritrophin-A | Peritrophin-A | CPAP3 | Y | FBpp0077028 | AAEL009585 | AGAP000986 |
| | PPAI004750 | obstructor-A | obst-A | CPAP3 | Y | FBpp0297663 | AAEL012245 | AGAP000989 |
| | PPAI002321 | CG8541 | CG8541 | CPF | Y | FBpp0076572 | AAEL011068 | AGAP004690 |
| | PPAI005554 | CG8543 | CG8543 | CPF | Y | FBpp0076573 | AAEL000890 | AGAP010901 |
| | PPAI004733 | CG13044 | CG13044 | CPLCA | Y | FBpp0075225 | AAEL001704 | AGAP006145 |
| | PPAI003971 | CG7203 | CG7203 | CPLCG | Y | FBpp0312122 | AAEL002187 | AGAP008459 |
| | PPAI001139 | Cuticular Protein 50Cb | Cpr50Cb | CPR_Uncl | Y | FBpp0086175 | AAEL017142 | AGAP000177 |
| | PPAI003073 | Cuticular Protein 56F | Cpr56F | CPR_Uncl | Y | FBpp0085613 | AAEL008754 | AGAP012487 |

| | | | | | | | |
|---|---|---|---|---|---|---|---|
| PPAI005524 | Cuticular Protein 65Ec | Cpr65Ec | CPR_Uncl | Y | FBpp0076618 | AAEL008285 | AGAP005459 |
| PPAI005525 | Pupal Cuticular Protein | Pcp | CPR_Uncl | Y | FBpp0079016 | AAEL008289 | AGAP005459 |
| PPAI005527 | Septin 2 | Sep2-PA | CPR_Uncl | Y | FBpp0083352 | AAEL004668 | AGAP011532 |
| PPAI009711 | Cuticular Protein 56F | Cpr56F | CPR_Uncl | Y | FBpp0085613 | AAEL008765 | AGAP012462 |
| PPAI000923 | Cuticular Protein 76Bd | Cpr76Bd | RR-1 | Y | FBpp0304995 | AAEL011504 | AGAP006868 |
| PPAI000924 | Cuticular Protein 76Bc | Cpr76Bc | RR-1 | Y | FBpp0304993 | AAEL008970 | AGAP006867 |
| PPAI000925 | Cuticular Protein 76Bb | Cpr76Bb | RR-1 | Y | FBpp0074702 | AAEL009005 | AGAP006866 |
| PPAI000927 | Cuticular Protein 62Ba | Cpr62Ba | RR-1 | Y | FBpp0288836 | AAEL009002 | AGAP006838 |
| PPAI000944 | Cuticular Protein 73D | Cpr73D | RR-1 | Y | FBpp0112089 | AAEL006737 | AGAP006369 |
| PPAI001238 | Cuticular Protein 47Ea | Cpr47Ea | RR-1 | Y | FBpp0087286 | AAEL003711 | AGAP009879 |
| PPAI001239 | Cuticular Protein 47Ee | Cpr47Ee | RR-1 | Y | FBpp0087215 | AAEL014416 | AGAP009878 |
| PPAI001921 | Cuticular Protein 97Ea | Cpr97Ea | RR-1 | Y | FBpp0084506 | AAEL012088 | AGAP000744 |
| PPAI002038 | Cuticular Protein | Cpr | RR-1 | Y | N/A | AAEL014987 | AGAP006858 |
| PPAI002218 | Cuticular Protein 47Ea | Cpr47Ea | RR-1 | Y | FBpp0087286 | AAEL014419 | AGAP009879 |
| PPAI002332 | Cuticular Protein 49 | Cpr49 | RR-1 | Y | FBpp0087035 | AAEL003216 | AGAP009868 |
| PPAI002939 | Cuticular Protein 49Ab | Cpr49Ab | RR-1 | Y | FBpp0087025 | AAEL003232 | AGAP009874 |
| PPAI002940 | Pupal Cuticular Protein | Pcp | RR-1 | Y | FBpp0079016 | AAEL003272 | AGAP009875 |
| PPAI002941 | Cuticular Protein 49Ae | Cpr49Ae | RR-1 | Y | FBpp0087031 | AAEL003259 | AGAP009871 |
| PPAI002942 | Cuticular Protein 49Af | Cpr49Af | RR-1 | Y | FBpp0311713 | AAEL003221 | AGAP009870 |
| PPAI003072 | Cuticular Protein 97Eb | Cpr97Eb | RR-1 | Y | FBpp0084505 | AAEL003041 | AGAP000344 |
| PPAI003900 | Cuticular Protein 92F | Cpr92F | RR-1 | Y | FBpp0083327 | AAEL015119 | AGAP000047 |
| PPAI004050 | Cuticular Protein | Cpr | RR-1 | Y | N/A | AAEL009005 | AGAP006857 |
| PPAI005059 | bancal | bl | RR-1 | Y | FBpp0089364 | AAEL014419 | AGAP005015 |
| PPAI005526 | Pupal Cuticular Protein | Pcp | RR-1 | Y | FBpp0079016 | AAEL008289 | AGAP005459 |
| PPAI005622 | Cuticular Protein 65Av | Cpr65Av | RR-1 | Y | FBpp0076732 | AAEL013515 | AGAP006001 |
| PPAI005624 | Cuticular Protein 65Av | Cpr65Av | RR-1 | Y | FBpp0076732 | AAEL013515 | AGAP005996 |
| PPAI005625 | Cuticular Protein 65Av | Cpr65Av | RR-1 | Y | FBpp0076732 | AAEL013515 | AGAP005996 |
| PPAI005626 | Cuticular Protein 65Av | Cpr65Av | RR-1 | N | FBpp0076732 | AAEL012883 | AGAP005996 |
| PPAI005627 | lethal (3) malignant blood neoplasm | l(3)mbn | RR-1 | Y | FBpp0076770 | AAEL011451 | AGAP005995 |
| PPAI007059 | Cuticular Protein 49Aa | Cpr49Aa | RR-1 | Y | FBpp0112271 | AAEL003272 | AGAP009876 |
| PPAI007060 | Cuticular Protein 49Ab | Cpr49Ab | RR-1 | Y | FBpp0087025 | AAEL003235 | AGAP009874 |
| PPAI007283 | Larval Cuticular Protein 65Ad | Lcp65Ad | RR-1 | Y | FBpp0305758 | AAEL002449 | AGAP002613 |

| | | | | | | | |
|---|---|---|---|---|---|---|---|
| PPAI007450 | Cuticular Protein 47Ef | Cpr47Ef | RR-1 | Y | FBpp0291859 | AAEL014418 | AGAP009877 |
| PPAI007533 | Cuticular Protein 76Bb | Cpr76Bb | RR-1 | Y | FBpp0074702 | AAEL009807 | AGAP006837 |
| PPAI007636 | CG15515 | CG15515 | RR-1 | Y | FBpp0304702 | AAEL015460 | AGAP006002 |
| PPAI008220 | Adult Cuticular Protein | Cpr | RR-1 | Y | N/A | AAEL008969 | AGAP006859 |
| PPAI009060 | Cuticular Protein 65Aw | Cpr65Aw | RR-1 | Y | FBpp0297891 | AAEL015460 | AGAP006001 |
| PPAI009072 | Cuticular Protein 65Az | Cpr65Az | RR-1 | Y | FBpp0076704 | AAEL001735 | AGAP006012 |
| PPAI009742 | Cuticular Protein 62Bb | Cpr62Bb | RR-1 | Y | FBpp0305097 | AAEL011043 | AGAP006830 |
| PPAI009743 | Cuticular Protein 62Bc | Cpr62Bc | RR-1 | Y | FBpp0035281 | AAEL011045 | AGAP006829 |
| PPAI009744 | Cuticular Protein 66Cb | Cpr66Cb | RR-1 | Y | FBpp0076328 | AAEL013380 | AGAP006828 |
| PPAI010782 | Cuticular Protein 65Ec | Cpr65Ec | RR-1 | Y | FBpp0076618 | AAEL008285 | AGAP005459 |
| PPAI000410 | Cuticular Protein 11A | Cpr11A | RR-2 | Y | FBpp0073483 | AAEL013683 | AGAP000085 |
| PPAI002037 | Cuticular Protein | Cpr | RR-2 | Y | N/A | N/A | AGAP006859 |
| PPAI002038 | Cuticular Protein | Cpr | RR-2 | Y | N/A | AAEL014987 | AGAP006858 |
| PPAI002283 | Cuticular Protein 100A | Cpr100A | RR-2 | Y | FBpp0085072 | AAEL003049 | AGAP000820 |
| PPAI002426 | CG34461 | Cpr66Ca | RR-2 | Y | FBpp0305270 | AAEL011504 | AGAP006283 |
| PPAI002427 | CG34462 | CG34462 | RR-2 | Y | FBpp0111680 | AAEL011501 | AGAP013248 |
| PPAI002760 | Cuticular Protein 92F | Cpr92F | RR-2 | Y | FBpp0083327 | AAEL015119 | AGAP000047 |
| PPAI002808 | CG8927 | CG8927 | RR-2 | Y | FBpp0310843 | AAEL012090 | AGAP000745 |
| PPAI003325 | Cuticular Protein 50Cb | Cpr50Cb | RR-2 | Y | FBpp0086756 | AAEL008764 | AGAP012466 |
| PPAI003841 | Cuticular Protein | Cpr | RR-2 | Y | N/A | AAEL002092 | AGAP003385 |
| PPAI003842 | Cuticular Protein 64Ab | Cpr64Ab | RR-2 | Y | FBpp0073109 | AAEL002110 | AGAP003378 |
| PPAI003843 | Cuticular Protein 64Ad | Cpr64Ad | RR-2 | Y | FBpp0073079 | AAEL002110 | AGAP003390 |
| PPAI006635 | Cuticular Protein 56F | Cpr56F | RR-2 | Y | FBpp0085613 | AAEL008765 | AGAP012462 |
| PPAI006706 | CG10625 | CG10625 | RR-2 | Y | FBpp0271739 | AAEL018209 | AGAP011506 |
| PPAI007221 | Cuticular Protein | Cpr | RR-2 | Y | N/A | AAEL015163 | AGAP001668 |
| PPAI008738 | Cuticular Protein 47Ef | Cpr47Ef | RR-2 | Y | FBpp0291859 | AAEL004951 | AGAP010887 |
| PPAI009993 | Cuticular Protein | Cpr | RR-2 | Y | N/A | AAEL004776 | AGAP010128 |
| PPAI010171 | Cuticular Protein 50Cb | Cpr50Cb | RR-2 | Y | FBpp0086756 | AAEL008726 | AGAP010717 |
| PPAI002105 | Tweedle T | TwdlT | TWDL | Y | FBpp0084471 | AAEL013793 | AGAP000352 |
| PPAI002309 | Tweedle T | TwdlT | TWDL | Y | FBpp0084471 | AAEL017015 | AGAP000352 |
| PPAI004510 | Tweedle Beta | TWdlBeta | TWDL | Y | FBpp0087168 | AAEL014994 | AGAP004576 |
| PPAI006993 | Tweedle E | TwdlE | TWDL | Y | FBpp0079185 | AAEL009478 | AGAP000537 |

2082

**S27 Table.** Juvenile hormone family annotation.

| Species | Vectorbase Gene ID | Gene | Symbol | Required Revision | *D. melanogaster* | *A. aegypi* | *A. gambiae* |
|---|---|---|---|---|---|---|---|
| *Lu. Longipalpis* | LLOJ007479 | paralytic | para | N | FBpp0292720 | AAEL006019 | AGAP004707 |
| | LLOJ000799 | short gastrulation | sog | N | FBpp0304148 | AAEL005627 | AGAP000472 |
| | LLOJ004071 | Juvenile hormone epoxide hydrolase 1 | jheh1 | N | FBpp0085807 | AAEL011314 | AGAP008684 |
| | LLOJ004199 | Juvenile hormone epoxide hydrolase 2 | jheh2 | N | FBpp0311868 | AAEL011313 | AGAP008686 |
| | LLOJ005310 | Vacuolar H+ ATPase 44kD subunit | Vha44 | N | FBpp0288471 | AAEL005173 | AGAP005845 |
| | LLOJ005486 | Haemolymph juvenile hormone binding | CG3246 | N | FBpp0077261 | AAEL009927 | AGAP002341 |
| | LLOJ008548 | Juvenile hormone esterase | Jhe | N | FBpp0290965 | AAEL005200 | AGAP005834 |
| | LLOJ005938 | ultraspiracle | usp | N | FBpp0305937 | AAEL000395 | AGAP002095 |
| | LLOJ001255 | sine oculis | so | N | FBpp0089177 | AAEL009170 | AGAP011695 |
| | LLOJ004815 | Chd64 | Chd64 | N | FBpp0073126 | AAEL006922 | AGAP001313 |
| | LLOJ000605 | tailless | tll | N | FBpp0085071 | AAEL003020 | AGAP000819 |
| | LLOJ008362 | Haemolymph juvenile hormone binding | CG10264 | N | FBpp0082676 | AAEL013324 | AGAP001982 |
| | LLOJ000266 | ftz transcription factor 1 | ftz-f1 | N | FBpp0074853 | AAEL002062 | AGAP005661 |
| | LLOJ007734 | Jhl-21 | Jhl-21 | N | FBpp0087449 | AAEL010336 | AGAP009743 |
| | LLOJ001069 | glass | gl | N | FBpp0083005 | AAEL002390 | AGAP004331 |
| | LLOJ005381 | Haemolymph juvenile hormone binding | CG3776 | N | FBpp0310109 | AAEL007196 | AGAP009891 |
| | LLOJ000480 | apterous | ap | N | FBpp0085394 | AAEL008694 | AGAP008980 |
| | LLOJ008363 | Haemolymph juvenile hormone binding | CG2016 | N | FBpp0297105 | AAEL013323 | AGAP001983 |
| | LLOJ008361 | Haemolymph juvenile hormone binding | CG2016 | N | FBpp0307387 | AAEL014366 | AGAP001981 |
| | LLOJ008364 | Haemolymph juvenile hormone binding | CG1124 | N | FBpp0078482 | AAEL013327 | AGAP013533 |
| | LLOJ009149 | Haemolymph juvenile hormone binding | CG15497 | N | FBpp0083512 | AAEL013325 | AGAP001984 |
| | LLOJ005012 | Haemolymph juvenile hormone binding | CG33680 | N | FBpp0091002 | AAEL001323 | AGAP028142 |
| | LLOJ003576 | juvenile hormone acid methyltransferase | jhamt | N | FBpp0309208 | AAEL006280 | AGAP005256 |
| | LLOJ006987 | Haemolymph juvenile hormone binding | CG11854 | N | FBpp0308366 | AAEL011966 | AGAP012703 |
| | LLOJ008792 | takeout | to | N | FBpp0308365 | AAEL011966 | AGAP004262 |
| | LLOJ000874 | FK506-binding protein 1 | FK506-bp1 | N | FBpp0082574 | AAEL003303 | AGAP007473 |
| | LLOJ006030 | Haemolymph juvenile hormone binding | CG11852 | N | FBpp0084184 | AAEL011966 | AGAP012703 |
| | LLOJ007016 | Haemolymph juvenile hormone binding | CG13618 | N | FBpp0084027 | AAEL011966 | AGAP012703 |
| | LLOJ003590 | juvenile hormone-inducible protein, putative | | N | N/A | AAEL004023 | AGAP003220 |
| | LLOJ001973 | Haemolymph juvenile hormone binding | CG5867 | N | FBpp0080029 | AAEL004987 | AGAP008182 |
| | LLOJ005637 | giant | gt | N | FBpp0070444 | AAEL009376 | AGAP001880 |
| | LLOJ007068 | *More closely related to gce gene, Methoprene-tolerant | met | N | FBpp0301991 | AAEL001746 | AGAP006022 |
| | LLOJ003890 | Malate dehydrogenase 1 | Mdh1 | N | FBpp0311560 | N/A | N/A |
| | LLOJ008974 | Yolk-protein, vitellogenin *Closer related to vitellogenin | Yp | N | FBpp0071359 | AAEL006966 | AGAP009104 |
| | LLOJ006997 | Yolk-protein, vitellogenin *Closer related to vitellogenin | Yp | N | FBpp0308710 | AAEL012311 | AGAP005013 |
| | LLOJ008286 | Yolk-protein, vitellogenin *Closer related to vitellogenin | Yp | Y | FBpp0308710 | AAEL012311 | AGAP005013 |
| *P. papatasi* | PPAI001672 | short gastrulation | sog | N | FBpp0304148 | AAEL018236 | AGAP000472 |

| | | | | | | |
|---|---|---|---|---|---|---|
| PPAI002908 | Vacuolar H+ ATPase 44kD subunit | Vha44 | N | FBpp0086320 | AAEL015594 | AGAP009485 |
| PPAI000394 | paralytic | para | N | FBpp0292691 | N/A | AGAP004707 |
| PPAI009818 | Juvenile hormone epoxide hydrolase 1 | Jheh1 | N | FBpp0085807 | AAEL0011314 | AGAP008684 |
| PPAI004780 | Juvenile hormone epoxide hydrolase 2 | Jheh2 | N | FBpp0311868 | AAEL0011313 | AGAP008685 |
| PPAI004470 | Jhl-1 | Jhl-1 | N | FBpp0087449 | AAEL010336 | AGAP005109 |
| PPAI006829 | Juvenile hormone esterase | Jhe | N | FBpp0290965 | AAEL004323 | AGAP005834 |
| PPAI006912 | Jhedup | Jhedup | N | FBpp0086361 | AAEL005210 | AGAP005837 |
| PPAI009809 | Haemolymph juvenile hormone binding | CG3246 | N | FBpp0077261 | AAEL009927 | AGAP002341 |
| PPAI009068 | Chd64 | Chd64 | N | FBpp0073126 | AAEL006922 | AGAP000749 |
| PPAI001367 | Haemolymph juvenile hormone binding | CG15497 | N | FBpp0083512 | AAEL013325 | AGAP001984 |
| PPAI009891 | Haemolymph juvenile hormone binding | CG10175 | N | FBpp0311034 | AAEL007486 | AGAP002391 |
| PPAI000899 | juvenile hormone acid methyltransferase | jhamt | N | FBpp0309208 | AAEL006280 | AGAP005256 |
| PPAI010395 | ftz transcription factor 1 | ftz-f1 | N | FBpp0074853 | AAEL002062 | AGAP005661 |
| PPAI010173 | Haemolymph juvenile hormone binding | CG3776 | N | FBpp0310109 | AAEL007196 | AGAP009891 |
| PPAI001860 | Haemolymph juvenile hormone binding | CG10264 | N | FBpp0082676 | AAEL013324 | AGAP001982 |
| PPAI009697 | sine oculis | so | N | FBpp0089177 | AAEL009170 | AGAP011695 |
| PPAI001859 | Haemolymph juvenile hormone binding | CG1124 | N | FBpp0078482 | AAEL013327 | AGAP013533 |
| PPAI005409 | ultraspiracle | usp | N | FBpp0305937 | AAEL000395 | AGAP002095 |
| PPAI007441 | Haemolymph juvenile hormone binding | CG13618 | N | FBpp0084027 | AAEL011966 | AGAP011167 |
| PPAI012824 | takeout | to | N | FBpp0308365 | AAEL011966 | AGAP004262 |
| PPAI010560 | glass | gl | N | FBpp0083005 | N/A | AGAP004331 |
| PPAI001532 | FK506-binding protein 1 | FK506-bp1 | N | FBpp0082574 | AAEL003303 | AGAP007473 |
| PPAI002209 | tailless | tll | N | FBpp0085071 | AAEL003020 | AGAP000819 |
| PPAI001861 | Haemolymph juvenile hormone binding | CG2016 | N | FBpp0297106 | AAEL014366 | AGAP001981 |
| PPAI009982 | Haemolymph juvenile hormone binding | CG5867 | N | FBpp0080029 | AAEL004987 | AGAP008182 |
| PPAI000977 | giant | gt | N | FBpp0070444 | AAEL009376 | AGAP001880 |
| PPAI000938 | Haemolymph juvenile hormone binding | CG14457 | N | FBpp0100135 | AAEL008307 | AGAP000750 |
| PPAI000466 | Haemolymph juvenile hormone binding | CG33680 | N | FBpp0091002 | AAEL001323 | AGAP028142 |
| PPAI007045 | Haemolymph juvenile hormone binding | CG31189 | N | FBpp0290041 | AAEL011966 | AGAP004262 |
| PPAI006506 | Yolk-protein, vitellogenin *Closer related to vitellogenin | Yp | Y | FBpp0308710 | AAEL012311 | AGAP005103 |
| PPAI003061 | Yolk-protein, vitellogenin *Closer related to vitellogenin | Yp | Y | FBpp0073652 | AAEL000828 | AGAP005103 |
| PPAI001123 | Yolk-protein, vitellogenin *Closer related to vitellogenin | Yp | Y | FBpp0071359 | AAEL005815 | AGAP005103 |
| PPAI001922 | apterous | ap | Y | FBpp0289731 | AAEL008685 | AGAP008979 |
| PPAI008795 | Haemolymph juvenile hormone binding | CG5867 | N | FBpp0080029 | AAEL004987 | AGAP008182 |

2084

**S28 Table.** Insulin signaling pathway annotation.

| Species | Gene Name | Gene Symbol | Lutzomyia Vectorbase ID | *D. melanogaster* | *A. aegypti* | *A. gambiae* | Revisions required | Notes |
|---|---|---|---|---|---|---|---|---|
| *Lu. Longipalpis* | Adiponectin Receptor | AdipoR | LLOJ001737 | FBgn0038984 | AAEL014717 | AGAP004486 | Major | |
| | Chico-insulin receptor substrate 1 | chico | LLOJ003065 | FBgn0024248 | AAEL007819, AAEL007817 | AGAP008859 | Minor | |
| | Coenzyme Q biosynthesis protein 2 | Coq2 | LLOJ007686 | FBgn0037574 | AAEL011249 | AGAP004513 | Major | |
| | Cullin | Cul1 | LLOJ007109 | FBgn0015509 | AAEL013530 | AGAP008007 | Major | apparent duplication? Two full copies in scaffold |
| | Downstream of Kinase | Dok | LLOJ006565 | FBgn0029944 | N/A | AGAP000825 | Major | |
| | Dreadlocks-like | dock | LLOJ003901 | FBgn0010583 | AAEL013539 | AGAP010135 | N | |
| | Fibroblast growth factor receptor substrate | FGFRs | LLOJ006221 | FBgn0032042 | AAEL001475 | AGAP009930 | Minor | |
| | Forkhead box subgroup o | foxo | LLOJ008538 | FBgn0038197 | AAEL012847 | AGAP001577 | Major | |
| | Happy-hour-like MAP4K3 | MAP4K3 | LLOJ002088 | FBgn0263395 | AAEL017338, AAEL013859 | AGAP010837 | N | |
| | Insulin like peptide 1 | Ilp1 | LLOJ006186 | FBgn0044051 | AAEL000937, AAEL000973 | AGAP010603 | N | |
| | Insulin-like-Receptor | InR | LLOJ003142 | FBgn0013984 | AAEL002317 | AGAP012424 | Minor | |
| | Lnk-like signal transduction protein | Lnk | LLOJ004990 | FBgn0028717 | AAEL004125 | AGAP003863 | N | |
| | Melted-like protein | Melt | LLOJ009634 | FBgn0023001 | AAEL009213 | AGAP007472 | N | |
| | metabotropic GABA-B receptor subtype 2 | GABA-B-R2 | LLOJ000435 | FBgn0027575 | AAEL010408 | AGAP004595 | N | |
| | Nucleostemin ortholog1 | Ns1 | LLOJ004073 | FBgn0038473 | AAEL011255 | AGAP004514 | N | |
| | PDGF- and VEGF-receptor related | Pvr | LLOJ001405 | FBgn0032006 | AAEL003928 | AGAP008813 | N | |
| | Phosphatidylinositol 3-kinase catalytic subunit | Pi3K92E | LLOJ001313 | FBgn0015279 | AAEL002903 | AGAP000018 | N | |
| | Phosphoinositide-dependent kinase 1 | Pdk1 | LLOJ006185 | FBgn0020386 | AAEL006440, AAEL007284 | AGAP001471 | Major | |
| | Poly-like protein | poly | LLOJ005207 | FBgn0086371 | AAEL012624 | AGAP003075 | N | |
| | Protein-Tyrosine phosphatase 61F | Ptp61F | LLOJ008161 | FBgn0267487 | AAEL001919 | AGAP005352 | Minor | |
| | Pten-phosphatase | Pten | LLOJ001893 | FBgn0026379 | AAEL017224 | AGAP009628 | N | |
| | Pudgy-like AMP dependent CoA ligase | pdgy | LLOJ009886 | FBgn0027601 | AAEL000415 | AGAP002718 | N | |

| | | | | | | | |
|---|---|---|---|---|---|---|---|
| | Short neuropeptide F precursor | sNPF | LLOJ003236 | FBgn0032840 | AAEL012542 | N/A | Minor |
| | Similar-like Hypoxia inducible factor | sima | LLOJ000925 | FBgn0266411 | AAEL015383, AAEL001097 | AGAP002942 | N |
| | S-phase kinase-associated protein | skpA | LLOJ001899 | FBgn0025637 | AAEL010651, AAEL009160 | AGAP008719 | N |
| | Slmb-like F-box/WD repeat protein | slmb | LLOJ006764 | FBgn0023423 | AAEL003371 | AGAP001944 | Major |
| | Small wing-like PLC-gamma | sl | LLOJ005588 | FBgn0003416 | AAEL004431 | AGAP011151, AGAP011152 | Major |
| | Spargle-like peroxisome proliferator-activated receptor gamma coactivator 1 | srl | LLOJ008949 | FBgn0037248 | AAEL003768 | AGAP004361 | Major |
| | Talin | rhea | LLOJ008380 | FBgn0260442 | AAEL006222 | AGAP007474 | Major |
| | Tango protein | tgo | LLOJ003291 | FBgn0264075 | AAEL010343 | AGAP009748 | Major |
| | Tuberous sclerosis complex 2 | Tsc2 | LLOJ004899 | FBgn0005198 | AAEL007712 | AGAP011123 | Major |
| *P. papatasi* | Adiponectin Receptor | AdipoR | PPAI002915 | FBgn0038984 | AAEL014717 | AGAP004486 | N |
| | Chico-insulin receptor substrate 1 | chico | PPAI007519 | FBgn0024248 | AAEL007819, AAEL007817 | AGAP008859 | N |
| | Cullin | Cul1 | PPAI005081 | FBgn0015509 | AAEL013530 | AGAP008007 | N |
| | Dreadlocks-like | dock | PPAI008675 | FBgn0010583 | AAEL013539 | AGAP010135 | N |
| | Fibroblast growth factor receptor substrate | FGFRs | PPAI010918 | FBgn0032042 | AAEL001475 | AGAP009930 | Minor |
| | Forkhead box subgroup o | foxo | PPAI000724 | FBgn0038197 | AAEL012847 | AGAP001577 | Minor |
| | Happy-hour-like MAP4K3 | MAP4K3 | PPAI002155 | FBgn0263395 | AAEL017338, AAEL013859 | AGAP010837 | N |
| | Insulin like protein 7 | Ilp7 | PPAI000801 | FBgn0044046 | AAEL003000 | AGAP003927 | Minor |
| | Insulin-like-Receptor | InR | PPAI002386 | FBgn0013984 | AAEL002317 | AGAP012424 | Major |
| | Lnk-like signal transduction protein | Lnk | PPAI004216 | FBgn0028717 | AAEL004125 | AGAP003863 | N |
| | Melted-like protein | Melt | PPAI005856 | FBgn0023001 | AAEL009213 | AGAP007472 | N |
| | metabotropic GABA-B receptor subtype 2 | GABA-B-R2 | PPAI006769 | FBgn0027575 | AAEL010408 | AGAP004595 | Major |
| | PDGF- and VEGF-receptor related | Pvr | PPAI010744 | FBgn0032006 | AAEL003928 | AGAP008813 | Major |
| | Phosphatidylinositol 3-kinase catalytic subunit | Pi3K92E | PPAI005862 | FBgn0015279 | AAEL002903 | AGAP000018 | N |
| | Phosphatidylinositol 3-kinase regulatory subunit 21B | Pi3K21B | PPAI004335 | FBgn0020622 | AAEL013596 | AGAP005583 | Major |

| | | | | | | |
|---|---|---|---|---|---|---|
| Phosphoinositide-dependent kinase 1 | Pdk1 | PPAI005178 | FBgn0020386 | AAEL006440, AAEL007284 | AGAP001471 | Minor |
| Protein-Tyrosine phosphatase 61F | Ptp61F | PPAI006334 | FBgn0267487 | AAEL001919 | AGAP005352 | N |
| Pten-phosphatase | Pten | PPAI003273 | FBgn0026379 | AAEL017224 | AGAP009628 | N |
| Pudgy-like AMP dependent CoA ligase | pdgy | PPAI001635 | FBgn0027601 | AAEL000415 | AGAP002718 | Minor |
| Ribosomal protein L8 | RpL8 | PPAI008202 | FBgn0261602 | AAEL000987 | AGAP005802 | Minor |
| Similar-like Hypoxia inducible factor | sima | PPAI002345 | FBgn0266411 | AAEL015383,AAEL001097 | AGAP002942 | Major |
| S-phase kinase-associated protein | skpA | PPAI010409 | FBgn0025637 | AAEL010651, AAEL009160 | AGAP008719 | N |
| Slmb-like F-box/WD repeat protein | slmb | PPAI004879 | FBgn0023423 | AAEL003371 | AGAP001944 | Major |
| Small wing-like PLC-gamma | sl | PPAI004351 | FBgn0003416 | AAEL004431 | AGAP011151, AGAP011152 | N |
| Spargle-like peroxisome proliferator-activated receptor gamma coactivator 1 | srl | PPAI001168 | FBgn0037248 | AAEL003768 | AGAP004361 | Major |
| Sugarbabe-like zinc finger protein | sug | PPAI005712 | FBgn0033782 | AAEL005120 | AGAP006736 | Major |
| Talin | rhea | PPAI009170 | FBgn0260442 | AAEL006222 | AGAP007474 | Major |
| Tango | tgo | PPAI009623 | FBgn0264075 | AAEL010343 | AGAP009748 | N |
| Tuberous sclerosis complex 1 | Tsc1 | PPAI007877 | FBgn0026317 | AAEL002548 | AGAP003445 | Minor |
| Tuberous sclerosis complex 2 | Tsc2 | PPAI010500 | FBgn0005198 | AAEL007712 | AGAP011123 | Major |

**S29 Table.** Antioxidant family annotation.

| Species | Gene Name | Gene Symbol | Phlebotomus Vectorbase ID | *D. melanogaster* | *A. aegypti* | *A. gambiae* | Revisions required |
|---|---|---|---|---|---|---|---|
| *Lu. Longipalpis* | Acyl-coa dehydrogenase | acd | LLOJ008579 | FBgn0034432 | AAEL004778 | AGAP008769 | Minor |
| | Alphabet-like protein phosphatase | alph | LLOJ007706 | FBgn0086361 | AAEL003326 | AGAP008149 | N |
| | Autophagy specific protein 18 | Atg18 | LLOJ002094 | FBgn0032935 | AAEL013063 | AGAP007970 | N |
| | Autophagy specific protein 9 | Atg9 | LLOJ000395 | FBgn0034110 | AAEL009105 | AGAP001762 | Minor |
| | Cardinal-like-oxidasae/peroxidase | cd | LLOJ007503 | FBgn0263986 | AAEL012481, AAEL011941 | AGAP012561, AGAP003502 | N |
| | Catalase | Cat | LLOJ007605 | FBgn0000261 | AAEL013407 | AGAP004904 | N |
| | Chloride intracellular channel | Clic | LLOJ008028 | FBgn0030529 | AAEL007761 | AGAP000943 | N |
| | Chorion/haem-binding peroxidase | HPX1 | LLOJ006481 | FBgn0038511 | AAEL002354 | AGAP000051 | N |
| | Chorion/haem-binding peroxidase | HPX2 | LLOJ006480 | FBgn0038511 | AAEL002354 | AGAP000051 | N |
| | CREB-regulated transcription coactivator | Crtc | LLOJ007525 | FBgn0036746 | AAEL008842 | AGAP008677 | Minor |
| | CYLD-like ubiquitin carboxyl-terminal hydrolase | CYLD | LLOJ007012 | FBgn0032210 | AAEL013253 | AGAP008412 | Major |
| | Daxx-like protein | DLP | LLOJ009097 | FBgn0031820 | AAEL004489 | AGAP009432 | Minor |
| | DJ-1beta | DJ-1beta | LLOJ002791 | FBgn0039802 | AAEL004081 | AGAP000705 | N |
| | Dual oxidase | duox | LLOJ010494 | FBgn0031464 | AAEL007563 | AGAP009978 | N |
| | Dual specificity protein phosphatase | dsp | LLOJ005232 | FBgn0036844 | AAEL001145 | AGAP012237 | N |
| | Forkhead box subgroup o | foxo | LLOJ008538 | FBgn0038197 | AAEL012847 | AGAP001577 | Major |
| | Fumble- like panthotenate kinase | fbl | LLOJ005896 | FBgn0011205 | AAEL012409 | AGAP011715 | N |
| | GLaz-like apolipoprotein D | GLaz | LLOJ000084 | FBgn0033799 | AAEL011278 | AGAP005223 | N |
| | Glutathione s transferase | GstD | LLOJ004462 | FBgn0001149 | AAEL001078, AAEL001061 | AGAP004164 | N |
| | Glutathione s transferase omega 2 | GstO2 | LLOJ009136 | FBgn0035906 | AAEL017085 | AGAP005749 | Minor |
| | Glutathione-S-transferase S 1 | GstS1 | LLOJ009037 | FBgn0010226 | AAEL011741 | AGAP010404 | N |
| | Hangover | hang | LLOJ010026 | FBgn0026575 | AAEL011847 | AGAP010021 | Major |
| | Heat shock protein 22 | Hsp22 | LLOJ005427 | FBgn0001223 | AAEL013350 | AGAP005548 | Minor |
| | Immune induced catalase | Icat | LLOJ007355 | FBgn0038465 | AAEL013171 | AGAP009033 | N |
| | Inositol triphosphate-3 kianse 1 | IP3K1 | LLOJ004867 | FBgn0032147 | AAEL010766 | AGAP002194 | Minor |
| | Jun-N-Terminal Kinase | Jnk | LLOJ005677 | FBgn0000229 | AAEL008634 | AGAP009461 | N |
| | Kelch-like ECH associate protein | Keap | LLOJ005290 | FBgn0038475 | AAEL005424 | AGAP003645 | Minor |
| | Locomotion defects | loco | LLOJ001915 | FBgn0020278 | AAEL007358 | AGAP002411 | Major |
| | Lon Protease | Lon | LLOJ008442 | FBgn0036892 | AAEL006474 | AGAP010451 | N |
| | MEKK1 | MEKK1 | LLOJ007004 | FBgn0024329 | AAEL010466 | AGAP002371 | Major |
| | Menin | Mnn | LLOJ002837 | FBgn0031885 | AAEL005686 | AGAP008130 | N |
| | Methionine sulfoxide reductase/ecdysone induced protein | msr | LLOJ008087 | FBgn0000565 | AAEL000670, AAEL000660 | AGAP012394, AGAP012395 | N |
| | Methusela | mth | LLOJ002123 | FBgn0023000 | AAEL010982 | AGAP006215, AGAP006216 | N |
| | Methyltransferase 2 | Mt2 | LLOJ001788 | FBgn0028707 | AAEL006166 | AGAP012836, AGAP004101 | Major |

| | | | | | | | |
|---|---|---|---|---|---|---|---|
| | Moladietz-dual oxidation maturation factor | mol | LLOJ005817 | FBgn0086711 | AAEL007562 | AGAP009976 | N |
| | Nicotinamide amidase | naam | LLOJ004470 | FBgn0051216 | AAEL002995 | AGAP001435 | Minor |
| | Nlaz-like Apolipoprotein | Nlaz | LLOJ003195 | FBgn0053126 | AAEL009561 | AGAP009281 | Major |
| | p38 mapk | p38a | LLOJ007636 | FBgn0015765 | AAEL008379 | AGAP012148 | Major |
| | Parkin (E3 ubiquitin ligase) | park | LLOJ008599 | FBgn0041100 | AAEL004260, AAEL004267 | AGAP006580 | N |
| | Period | per | LLOJ000134 | FBgn0003068 | AAEL008141 | AGAP001856 | Major |
| | Peroxidase | Pxd | LLOJ007354 | FBgn0004577 | AAEL015354 | AGAP010734 | Minor |
| | Peroxidasin | Pxn | LLOJ010039 | FBgn0011828 | AAEL000376 | AGAP007237 | Major |
| | Peroxinectin/chorion peroxidase | pxt | LLOJ003300 | FBgn0261987 | AAEL004388 | AGAP004038 | N |
| | Peroxiredoxin | prx | LLOJ006984 | FBgn0033521 | AAEL002309 | AGAP011824 | N |
| | Peroxiredoxin 2540 | prx-2540 | LLOJ006986 | FBgn0033520 | AAEL002309 | AGAP011824 | N |
| | Peroxiredoxin-1 | Prx-1 | LLOJ008460 | FBgn0040309 | AAEL004112 | AGAP011054 | N |
| | Peroxiredoxin-3 like | Prx-3 | LLOJ010087 | FBgn0038519 | AAEL013528 | AGAP000396 | Major |
| | Peroxiredoxin-6 like | Prx-6 | LLOJ004050 | FBgn0031479 | AAEL009051 | AGAP007020 | N |
| | Phosphatidylinositol 3-kinase catalytic subunit | Pi3K92E | LLOJ001313 | FBgn0015279 | AAEL002903 | AGAP000018 | N |
| | Phospholipid hydroperoxide glutathione peroxidase | PHGPx | LLOJ009243 | FBgn0035438 | AAEL012069 | AGAP004247 | N |
| | Pink1-like serine/threonine protein kinase | Pink1 | LLOJ003075 | FBgn0029891 | AAEL011594 | AGAP004315 | Minor |
| | Rutebaga | rut | LLOJ004622 | FBgn0003301 | AAEL017420, AAEL009022 | AGAP000727 | Major |
| | Selenoprotein-like methionine-R-sulfoxide reductase | SelR | LLOJ009961 | FBgn0267376 | AAEL002620 | AGAP013286 | Minor |
| | Stress-sensitive B -like ADP/ATP translocase | sesB | LLOJ003254 | FBgn0003360 | AAEL010884 | AGAP002358 | N |
| | Succinate dehydrogenase C | SdhC | LLOJ002875 | FBgn0037873 | AAEL008871 | AGAP010672 | N |
| | Superoxide dismutase | sod | LLOJ008594 | FBgn0003462 | AAEL011498, AAEL000259, AAEL000274 | AGAP010347 | N |
| | Superoxide dismutase 2 | Sod2 | LLOJ003300 | FBgn0010213 | AAEL004823 | AGAP010517 | Major |
| | Superoxide dismutase 3 | Sod3 | LLOJ010547 | FBgn0033631 | AAEL006271 | AGAP005234 | Major |
| | Thioredoxin dependent peroxide reductase | Tdpr | LLOJ002275 | FBgn0040308 | AAEL014548 | AGAP007543 | N |
| | Thioredoxin reductase-1 | Trxr-1 | LLOJ007938 | FBgn0020653 | AAEL002886 | AGAP000565 | N |
| | Thor-like Eukaryotic translation initiation factor 4E-binding protein | eif4E-bp | LLOJ007311 | FBgn0261560 | AAEL001864 | AGAP007781 | N |
| | Transcription factor A, mitochondrial | TFAM | LLOJ007295 | FBgn0038805 | AAEL013643, AAEL014794 | AGAP008499 | N |
| | Trap1-mitochondrial heat shock protein | Trap1 | LLOJ003520 | FBgn0026761 | AAEL000301 | AGAP010691 | N |
| | unc51-like kinase/Atg1 | unc 51 | LLOJ007855 | FBgn0260945 | AAEL003819, AAEL016987 | AGAP011295 | Minor |
| | Withered-like carnitine o-acyltransferase | withered | LLOJ005074 | FBgn0261862 | AAEL005458 | AGAP010164 | N |
| *P. papatasi* | Alphabet-like protein phosphatase | alph | PPAI010455 | FBgn0086361 | AAEL003326 | AGAP008149 | N |
| | Autophagy specific protein 18 | Atg18 | PPAI000077 | FBgn0032935 | AAEL013063 | AGAP007970 | N |
| | Autophagy specific protein 9 | Atg9 | PPAI002648 | FBgn0034110 | AAEL009105 | AGAP001762 | N |

| | | | | | | |
|---|---|---|---|---|---|---|
| Cap-n-collar | cnc | PPAI009244 | FBgn0262975 | AAEL015467, AAEL005077 | AGAP005300 | Minor |
| Cardinal-like oxidase/peroxidase | cd | PPAI001636 | FBgn0263986 | AAEL012481, AAEL011941 | AGAP012561, AGAP003502 | N |
| Catalase | Cat | PPAI006351 | FBgn0000261 | AAEL013407 | AGAP004904 | N |
| Chloride intracellular channel | Clic | PPAI005128 | FBgn0030529 | AAEL007761 | AGAP000943 | N |
| Chorion peroxidase | Cpx | PPAI009865 | FBgn0038511 | AAEL002354 | AGAP000051 | Minor |
| cln3/Batten's disease protein | cln3 | PPAI000935 | FBgn0036756 | AAEL009909 | AGAP003934 | N |
| CYLD-like ubiquitin carboxyl-terminal hydrolase | CYLD | PPAI005913 | FBgn0032210 | AAEL013253 | AGAP008412 | N |
| Daxx-like protein | DLP | PPAI001351 | FBgn0031820 | AAEL004489 | AGAP009432 | Minor |
| DJ-1beta | DJ-1beta | PPAI010696 | FBgn0039802 | AAEL004081 | AGAP000705 | Minor |
| Dual oxidase | duox | PPAI003114 | FBgn0031464 | AAEL007563 | AGAP009978 | N |
| Forkhead box subgroup o | foxo | PPAI000724 | FBgn0038197 | AAEL012847 | AGAP001577 | Minor |
| Fumble- like panthotenate kinase | fbl | PPAI009427 | FBgn0011205 | AAEL012409 | AGAP011715 | Minor |
| GLaz-like apolipoprotein D | GLaz | PPAI000239 | FBgn0033799 | AAEL011278 | AGAP005223 | Major |
| Glutathione S transferase | GstD | PPAI010868 | FBgn0001149 | AAEL001078, AAEL001061 | AGAP004164 | N |
| Glutathione-S-transferase S 1 | GstS1 | PPAI002540 | FBgn0010226 | AAEL011741 | AGAP010404 | N |
| Haem-binding Peroxidase | HPX1 | PPAI009674 | FBgn0032685 | AAEL003933 | AGAP008350 | Minor |
| Haem-binding Peroxidase | HPX2 | PPAI005168 | FBgn0259233 | AAEL005416 | AGAP003714 | Minor |
| Hangover | hang | PPAI009131 | FBgn0026575 | AAEL011847 | AGAP010021 | Major |
| Inositol triphosphate-3 kianse 1 | IP3K1 | PPAI003293 | FBgn0032147 | AAEL010766 | AGAP002194 | Minor |
| Jun-N-terminal Kinase | jnk | PPAI005249 | FBgn0000229 | AAEL008634 | AGAP009461 | N |
| Kelch-like ECH associate protein | Keap | PPAI010175 | FBgn0038475 | AAEL005424 | AGAP003645 | N |
| Locomotion defects | loco | PPAI009736 | FBgn0020278 | AAEL007358 | AGAP002411 | N |
| Menin | Mnn | PPAI001130 | FBgn0031885 | AAEL005686, AAEL005553 | AGAP008130 | N |
| Methionine sulfoxide reductase/ecdysone induced protein | msr | PPAI000349 | FBgn0000565 | AAEL000670, AAEL000660 | AGAP012394, AGAP012395 | Minor |
| Methusela | mth | PPAI008623 | FBgn0023000 | AAEL010982 | AGAP006215, AGAP006216 | Minor |
| Methyltransferase 2 | mt2 | PPAI010989 | FBgn0028707 | AAEL006166 | AGAP012836, AGAP004101 | N |
| Mitogen activated protein kinase kinase | Mekk | PPAI009824 | FBgn0024329 | AAEL010466 | AGAP002371 | Major |
| Moladietz-dual oxidation maturation factor | mol | PPAI003112 | FBgn0086711 | AAEL007562 | AGAP009976 | N |
| Nicotinamide amidase | naam | PPAI002435 | FBgn0051216 | AAEL002995 | AGAP001435 | N |
| Nuclear protein 1 | Np1 | PPAI003218 | FBgn0032400 | AAEL003160 | n/a | N |
| p38 map kinase | p38 | PPAI008377 | FBgn0024846 | AAEL008379 | AGAP012148 | Minor |
| Parkin (E3 ubiquitin ligase) | park | PPAI004091 | FBgn0041100 | AAEL004260, AAEL004267 | AGAP006580 | N |
| Period | per | PPAI006484 | FBgn0003068 | AAEL008141 | AGAP001856 | Minor |
| Peroxidase | Pxd | PPAI009536 | FBgn0004577 | AAEL015354 | AGAP010734 | Major |
| Peroxinectin/chorion peroxidase | pxt | PPAI012814 | FBgn0261987 | AAEL004388 | AGAP004038 | N |

| | | | | | | |
|---|---|---|---|---|---|---|
| Peroxiredoxin 2540 | prx-2540 | PPAI002050 | FBgn0033520 | AAEL002309 | AGAP011824 | Minor |
| Peroxiredoxin-1 | Prx-1 | PPAI006220 | FBgn0040309 | AAEL004112 | AGAP011054 | N |
| Peroxiredoxin-3 like | Prx-3 | PPAI006237 | FBgn0038519 | AAEL013528 | AGAP000396 | N |
| Peroxiredoxin-5 | Prx-5 | PPAI003012 | FBgn0038570 | AAEL007135 | AGAP001325 | N |
| Phosphatidylinositol 3-kinase catalytic subunit | Pi3K92E | PPAI005862 | FBgn0015279 | AAEL002903 | AGAP000018 | N |
| Phospholipid hydroperoxide glutathione peroxidase | PHGPx | PPAI010139 | FBgn0035438 | AAEL012069 | AGAP004247 | Minor |
| Ras which interacts with Calmodulin | Ric | PPAI009427 | FBgn0265605 | AAEL009180, AAEL015232 | AGAP005159 | Minor |
| Rutebaga | rut | PPAI001426 | FBgn0003301 | AAEL017420, AAEL009022 | AGAP000727 | Major |
| Selenoprotein-like methionine-R-sulfoxide reductase | SelR | PPAI008065 | FBgn0267376 | AAEL002620 | AGAP013286 | N |
| Stress-sensitive B -like ADP/ATP translocase | sesB | PPAI001953 | FBgn0003360 | AAEL010884 | AGAP002358 | Major |
| Stunted-like ATP synthase subunit | sun | PPAI005180 | FBgn0014391 | AAEL004060 | AGAP005098, AGAP009564, AGAP000260 | N |
| Succinate dehydrogenase C | SdhC | PPAI008179 | FBgn0037873 | AAEL008871 | AGAP010672 | Minor |
| Superoxide dismutase | sod | PPAI005974 | FBgn0003462 | AAEL011498, AAEL000259, AAEL000274 | AGAP010347 | Minor |
| Superoxide dismutase 2 | Sod2 | PPAI006666 | FBgn0010213 | AAEL004823 | AGAP010517 | Minor |
| Superoxide dismutase 3 | Sod3 | PPAI005793 | FBgn0033631 | AAEL006271 | AGAP005234 | Major |
| Thioredoxin 2 | Trx-2 | PPAI003111 | FBgn0040070 | AAEL010777 | AGAP009584 | Major |
| Thioredoxin dependent peroxide reductase | Tdpr | PPAI005157 | FBgn0040308 | AAEL014548 | AGAP007543 | N |
| Thor-like Eukaryotic translation initiation factor 4E-binding protein | eif4E-bp | PPAI008513 | FBgn0261560 | AAEL001864 | AGAP007781 | N |
| Transcription factor A, mitochondrial | TFAM | PPAI004961 | FBgn0038805 | AAEL013643, AAEL014794 | AGAP008499 | Minor |
| Trap1-mitochondrial heat shock protein | Trap1 | PPAI002821 | FBgn0026761 | AAEL000301 | AGAP010691 | N |
| unc51-like kinase/Atg1 | unc 51 | PPAI003929 | FBgn0260945 | AAEL003819, AAEL016987 | AGAP011295 | N |
| Withered-like carnitine o-acyltransferase | whd | PPAI002758 | FBgn0261862 | AAEL005458 | AGAP010164 | minor |

**S30 Table.** Vitamin metabolism pathway annotation.

| Species | Vectorbase Gene ID | Gene | Symbol | Revision Required | *D. melanogaster* | *A. gambiae* | *A. aegypti* |
|---|---|---|---|---|---|---|---|
| *Lu. Longipalpis* | LLOJ003887 | CG12170 D. Mel. & 3-oxoacyl-[acyl-carrier-protein] synthase | | N | N/A | AGAP002809 | AAEL002113 |
| | LLOJ008292 | Vanin-like protein D.Mel. (Pantetheine activity) | Vanin-like | N | FBgn0040069 | AGAP010733 | AAEL002681 |
| | LLOJ009995 | CG10238 molybdenum cofactor synthesis 2 | Mocs2 | N | FBgn0039280 | AGAP013168 | AAEL015584 |
| | LLOJ008404 | Sepiapterin reductase (short chain dehyrogenase) | Sptr | N | FBgn0014032 | N/A | N/A |
| | LLOJ001196 | Dihydrofolate reductase | Dhfr | N | FBgn0004087 | AGAP010278 | AAEL007370 |
| | LLOJ008224 | CG2543 & CG31773 Folylpolyglutamate synthetase | | N | FBgn0032779 | AGAP004679 | AAEL012882 |
| | LLOJ002185 | Dihyropteridine reductase | Dhpr | N | FBgn0035964 | N/A | N/A |
| | LLOJ008769 | Lipoic Acid Synthase | Las | N | FBgn0029158 | AGAP010568 | AAEL009368 |
| | LLOJ001529 | Lipoic Acid Synthase | Las | N | FBgn0029158 | AGAP010568 | AAEL009368 |
| | LLOJ008489 | CG8446 lipoyltransferase 1 | | N | N/A | AGAP012186 | AAEL007881 |
| | LLOJ005375 | CG9804 lipoyltransferase 2 | | N | FBgn0037251 | AGAP000531 | AAEL009657 |
| | LLOJ007053 | CG16758 purine nucleoside phosphorylase | | N | FBgn0035348 | AGAP005945 | AAEL002269 |
| | LLOJ003858 | CG32549 cytosolic purine 5-nucleotidase | | N | FBgn0052549 | AGAP000380 | AAEL005833 |
| | LLOJ001680 | CG3362 Pyrimidine 5'-nucleotidase (cN-IIIB) | cN-IIIB | N | FBgn0034988 | AGAP010681 | AAEL000258 |
| | LLOJ005681 | CG42249 5'-Nucleotidase/apyrase | | N | FBgn0259101 | AGAP007140 | AAEL011346 |
| | LLOJ002740 | NAD-dependent methylenetetrahydrofolate | Nmdmc | N | FBgn0010222 | AGAP004677 | AAEL014871 |
| | LLOJ001722 | Thymidylate synthase | ts | N | FBgn0024920 | AGAP010457 | AAEL013305 |
| | LLOJ001723 | Thymidylate synthase | ts | N | FBgn0024920 | AGAP010457 | AAEL013305 |
| | LLOJ001723 | CG34424 5-formyltetrahydrofolate cyclo-ligase | | N | FBgn0085453 | AGAP006824 | AAEL010694 |
| | LLOJ004871 | CG8665 10-formyltetrahydrofolate dehydrogenase | | N | FBgn0032945 | AGAP009591 | AAEL010764 |
| | LLOJ000519 | CG6106 allantoinase | | N | FBgn0030914 | AGAP000239 | AAEL007653 |
| | LLOJ007452 | Suppressor of rudimentary/Dihydroorotate dehydrogenase | su(r | N | FBgn0086450 | AGAP001021 | AAEL014199 |
| | LLOJ000583 | Beta-ureidopropionase activity/beta-alanine synthase | pyd3 | N | FBgn0037513 | AGAP010299 | AAEL010284 |

| | | | | | | |
|---|---|---|---|---|---|---|
| LLOJ005275 | Phosphopantothenoylcysteine synthetase | Ppcs | N | FBgn0261285 | AGAP004311 | AAEL014176 |
| LLOJ005896 | CG5725 pantothenate kinase activity | fbl | N | FBgn0011205 | AGAP011715 | AAEL012409 |
| LLOJ003214 | CG5828 pantothenate kinase activity | | N | FBgn0031682 | AGAP010073 | AAEL006648 |
| LLOJ003067 | CG1673 Branched-chain amino acid aminotransferase | | N | FBgn0030482 | AGAP000011 | AAEL007909 |
| LLOJ001286 | CG11899 Phosphoserine aminotransferase | | N | FBgn0014427 | AGAP004598 | AAEL012578 |
| LLOJ001350 | CG31472 Pyridoxamine 5'-phosphate oxidase, sugarlethal | sgll | N | FBgn0051472 | AGAP002227 | AAEL002703 |
| LLOJ009373 | CG34455 pyridoxal kinase | Pdxk | N | FBgn0085484 | AGAP005929 | AAEL009601 |
| LLOJ007107 | CG6656 acid phosphatase activity | | N | FBgn0038912 | AGAP004591 | AAEL015151 |
| LLOJ007844 | CG9449 acid phosphatase activity | Acph-1 | N | FBgn0000032 | AGAP002387 | AAEL003903 |
| LLOJ006172 | CG10581 nucleoside-triphosphatase activity | | N | FBgn0037046 | AGAP005921 | AAEL011713 |
| LLOJ009206 | CG12264 probable cysteine desulfurase | | N | FBgn0032393 | AGAP009094 | AAEL010743 |
| LLOJ001100 | CG14721 Thiamin pyrophosphokinase | | N | FBgn0037942 | AGAP002968 | AAEL006587 |
| LLOJ004643 | CG11883, Calcineurin-like phosphoesterase domain | | N | FBgn0033538 | AGAP001361 | AAEL012154 |
| LLOJ005953 | Nicotinamide mononucleotide adenylyltransferase | Nmnat | N | FBgn0039254 | AGAP009544 | AAEL007908 |
| LLOJ007244 | Ecto-5'-nucleotidase 2 | NT5E-2 | N | FBgn0050104 | AGAP003629 | AAEL010986 |
| LLOJ009119 | Ecto-5'-nucleotidase 2 | NT5E-2 | N | FBgn0050104 | AGAP003629 | AAEL010986 |
| LLOJ009014 | CG33156, NAD+ kinase activity | | N | FBgn0053156 | AGAP011122 | AAEL000278 |
| LLOJ006368 | CG3714, Nicotinate phophoribosyltransferase activity | | N | FBgn0031589 | AGAP008516 | AAEL014217 |
| LLOJ001928 | Adenosine 3 | ade3 | Y | FBgn0000053 | AGAP009786 | AAEL010640 |
| LLOJ003755 | Pugilist, methylenetetrahydrofolate dehydrogenase | pug | Y | FBgn0020385 | AGAP002830 | AAEL006085 |
| LLOJ005316 | Bifunctional Phosphopantetheine adenylyltransferase | Ppat-Dpck | N | FBgn0035632 | N/A | N/A |
| LLOJ002277 | Ebony activating protein | eap | N | FBgn0052099 | AGAP007546 | AAEL013942 |
| LLOJ007040 | CG4407 Phosphoadenosine phophosulphate reductase | | N | FBgn0030431 | AGAP002740 | AAEL011099 |
| LLOJ003989 | Primo-1 | Primo-1 | N | FBgn0040077 | AGAP007400 | AAEL001423 |

| | | | | | | | |
|---|---|---|---|---|---|---|---|
| | LLOJ008982 | CG9451 acid phosphatase activity | | N | FBgn0036876 | AGAP007400 | AAEL001424 |
| | LLOJ004173 | CG5150 alkaline phosphatase | | N | FBgn0035620 | N/A | N/A |
| | LLOJ004452 | CG1809 alkaline phosphatase | | N | FBgn0033423 | N/A | N/A |
| | LLOJ008256 | CG10592 alkaline phosphatase | | N | FBgn0035619 | AGAP011302 | AAEL003298 |
| | LLOJ004231 | CG8147 alkaline phosphatase | | N | FBgn0043791 | AGAP007300 | AAEL009077 |
| | LLOJ005944 | CG5656 alkaline phosphatase | | N | FBgn0037083 | AGAP001684 | AAEL003905 |
| | LLOJ006460 | Punch, GTP cyclohydrolase I | Pu | N | FBgn0003162 | AGAP006441 | AAEL002261 |
| | LLOJ009172 | Molybdenum cofactor synthesis 1 | Mocs1 | N | FBgn0263241 | N/A | N/A |
| | LLOJ003286 | lethal (3) 72Dp, gamma-glutamyl hydrolase | l(3)72Dp | N | FBgn0263607 | AGAP006670 | AAEL000271 |
| | LLOJ001549 | CG8147 alkaline phosphatase | | N | FBgn0043791 | AGAP011302 | AAEL003298 |
| | LLOJ008255 | CG1809 alkaline phosphatase | | N | FBgn0033423 | AGAP011302 | AAEL003298 |
| *P. papatasi* | PPAI001335 | Holocarboxylase synthetase | Hcs | N | FBgn0037332 | AGAP001481 | AAEL009340 |
| | PPAI008082 | Pantetheine hydrolase activity (Vanin-like) | Vanin-like | N | FBgn0040069 | AGAP010733 | AAEL006023 |
| | PPAI007151 | Molybdopterin synthase 2 | Mocs2 | N | FBgn0039280 | AGAP013168 | AAEL015584 |
| | PPAI001477 | sepiapterin reductase/short-chain dehydrogenase | Sptr | N | FBgn0014032 | AGAP008000 | AAEL001845 |
| | PPAI003707 | Dihydrofolate reductase | Dhfr | N | FBgn0004087 | AGAP010278 | AAEL007370 |
| | PPAI005073 | Short-chain dehydrogenase/reductase SDR | Dhpr | N | FBgn0035964 | AGAP002534 | AAEL006836 |
| | PPAI006170 | GTP cyclohydrolase I/Punch | Pu | N | FBgn0003162 | AGAP006441 | AAEL002261 |
| | PPAI004782 | Lipoic acid synthase | Las | N | FBgn0029158 | AGAP010568 | AAEL009368 |
| | PPAI000354 | CG8446 lipoyltransferase 1 | | N | N/A | AGAP012186 | AAEL007881 |
| | PPAI005715 | CG11883 5'-nucleotidase activity | | N | FBgn0033538 | AGAP007730 | AAEL012154 |
| | PPAI004593 | Nicotinamide mononucleotide adenylyltransferase | Nmnat | N | FBgn0039254 | AGAP009544 | AAEL007908 |
| | PPAI009440 | CG16758 & CG18128 purine-nucleoside phosphorylase activity | | N | FBgn0035348 | AGAP005945 | AAEL002269 |
| | PPAI005876 | CG32549 Cytosolic purine 5'-nucleotidase | | N | FBgn0052549 | AGAP000380 | AAEL005833 |
| | PPAI010184 | CG33156 NAD+ kinase-like | | N | FBgn0053156 | AGAP011122 | AAEL000278 |
| | PPAI006109 | CG3362 Pyrimidine 5'-nucleotidase | cN-IIIB | N | FBgn0034988 | AGAP010681 | AAEL000258 |

| | | | | | | |
|---|---|---|---|---|---|---|
| PPAI004594 | CG3714 nicotinate phosphoribosyltransferase activity | | N | FBgn0031589 | AGAP008516 | AAEL014217 |
| PPAI006440 | CG42249 5'-Nucleotidase/apyrase | | N | FBgn0259101 | AGAP001601 | AAEL011341 |
| PPAI005747 | CG11089 Bifunctional purine biosynthesis protein | | N | FBgn0039241 | AGAP001423 | AAEL012825 |
| PPAI004362 | CG1750 methionyl-tRNA formyltransferase activity | | N | FBgn0039836 | AGAP000398 | AAEL005279 |
| PPAI002644 | CG4067 methylenetetrahydrofolate dehydrogenase activity | pug | N | FBgn0020385 | AGAP002830 | AAEL006085 |
| PPAI002644 | CG6415 aminomethyltransferase activity | | N | FBgn0032287 | AGAP001124 | AAEL010276 |
| PPAI003287 | CG8665 10-formyltetrahydrofolate dehydrogenase | | N | FBgn0032945 | AGAP009591 | AAEL010764 |
| PPAI005169 | Bifunctional phosphopantetheine adenylytransferase | Ppat-Dpck | N | FBgn0035632 | AGAP005460 | AAEL003770 |
| PPAI005772 | CG1673 Branched-chain amino acid aminotransferase | | N | FBgn0030482 | AGAP000011 | AAEL007909 |
| PPAI002266 | Suppressor of rudimentary/Dihydroorotate dehydrogenase | su(r | N | FBgn0086450 | AGAP001021 | AAEL014199 |
| PPAI001004 | Phosphopantothenoylcysteine decarboxylase | Ppcdc | N | FBgn0050290 | AGAP011994 | AAEL010356 |
| PPAI008974 | CG5725 pantothenate kinase activity | fbl | N | FBgn0011205 | AGAP011715 | AAEL012409 |
| PPAI001356 | CG5828 pantothenate kinase activity | | N | FBgn0031682 | AGAP010073 | AAEL006648 |
| PPAI008797 | CG31472 Pyridoxamine 5'-phosphate oxidase, sugarlethal | sgll | N | FBgn0051472 | AGAP002227 | AAEL002703 |
| PPAI004756 | CG34455 pyridoxal kinase activity | Pdxk | N | FBgn0085484 | AGAP005929 | AAEL009601 |
| PPAI003010 | CG2846 Riboflavin kinase domain | | N | FBgn0014930 | AGAP007006 | N/A |
| PPAI002201 | Prophenoloxidase 1 | PPO1 | N | FBgn0261362 | AGAP002825 | AAEL014544 |
| PPAI010450 | Prophenoloxidase 1 | PPO1 | N | FBgn0261362 | AGAP002825 | AAEL014544 |
| PPAI001760 | CG6656 lysosomal acid phosphatase | | N | FBgn0038912 | AGAP004591 | AAEL013276 |
| PPAI000994 | CG7899 acid phosphatase 1 | Acph-1 | N | FBgn0000032 | AGAP003903 | AAEL003903 |
| PPAI005909 | CG12264 Cysteine desulfurase | | N | FBgn0032393 | AGAP009094 | AAEL010743 |
| PPAI007390 | CG2846 Riboflavin Kinase | | N | FBgn0014930 | N/A | N/A |
| PPAI004011 | Purple, 6-pyruvoyl tetrahydropterin synthase | pr | N | FBgn0003141 | AGAP008730 | AAEL001642 |
| PPAI000861 | CG2543, Folylpolyglutamate synthetase | | N | FBgn0030407 | AGAP004679 | AAEL012882 |
| PPAI002461 | Molybdenum cofactor synthesis 1 | Mocs1 | N | FBgn0263241 | AGAP006072 | AAEL000126 |

| | | | | | | |
|---|---|---|---|---|---|---|
| PPAI008977 | Veil, 5'- Nucleotidase activity | veil | N | FBgn0034225 | AGAP003629 | AAEL010986 |
| PPAI008706 | NAD-dependent methylenetetrahydrofolate | Nmdmc | N | FBgn0010222 | AGAP004677 | AAEL010751 |
| PPAI002746 | CG34424, 5'-formyltetrahydrofolate cyclo-ligase | | N | FBgn0085453 | AGAP006824 | AAEL010694 |
| PPAI008349 | CG7560, methylenetetrahydrofolate reductase | | N | FBgn0036157 | N/A | N/A |
| PPAI004605 | Collapsin Response Mediator Protein | CRMP | N | FBgn0023023 | AGAP003124 | AAEL007633 |
| PPAI007924 | Ebony activating protein | eap | N | FBgn0052099 | N/A | N/A |
| PPAI002918 | Pyridoxal phosphate binding | | N | N/A | AGAP004598 | AAEL012578 |
| PPAI005422 | Pyd3 beta-ureidopropionase activity | pyd3 | N | FBgn0037513 | AGAP010229 | AAEL010284 |
| PPAI005423 | Pyd3 beta-ureidopropionase activity | pyd3 | N | FBgn0037513 | AGAP010229 | AAEL010284 |
| PPAI008224 | CG4407 molybdopterin binding protein | | N | FBgn0030431 | AGAP002740 | AAEL011099 |
| PPAI005573 | Primo-1, Protein-tyrosine phosphatase, low molecular weight | primo-1 | Y | FBgn0040077 | AGAP004079 | AAEL003165 |
| PPAI009339 | CG10581 Nucleoside-triphosphatase | | N | FBgn0037046 | AGAP005921 | AAEL011713 |
| PPAI008766 | CG12170 3-oxoacyl-[acyl-carrier-protein] synthase | | N | FBgn0037356 | AGAP002809 | AAEL002113 |
| PPAI008311 | CG10592, Alkaline Phosphatase | | N | FBgn0035619 | AGAP001684 | AAEL003905 |
| PPAI006502 | CG10592, Alkaline Phosphatase | | N | FBgn0035619 | AGAP011302 | AAEL003298 |
| PPAI006501 | CG10592, Alkaline Phosphatase | | Y | FBgn0035619 | AGAP011302 | AAEL003298 |
| PPAI009724 | CG1809, Alkaline Phosphatase | | N | FBgn0033423 | N/A | N/A |
| PPAI004079 | CG5656, Alkaline Phosphatase | | Y | FBgn0037083 | AGAP007300 | AAEL009077 |
| PPAI004603 | CG16771, Alkaline Phosphatase | | N | FBgn0032779 | AGAP010596 | AAEL000931 |

**S31 Table.** *Phlebotomus papatasi* population sequencing median coverage depth.

| Sample | Median Coverage |
|---|---|
| PRAP.PPAFG_12.12 | 8 |
| PRAP.PPNS_80.80 | 9 |
| PRAP.PPNS_82.82 | 10 |
| PRAP.PPTUN_12.12 | 10 |
| PRAP.PPAFG_5.5 | 11 |
| PRAP.PPAFG_6.6 | 11 |
| PRAP.PPTUN_1.1 | 11 |
| PRAP.PPTUN_7.7 | 11 |
| PRAP.PPNS_25.25 | 12 |
| PRAP.PPNS_59.59 | 12 |
| PRAP.PPTUN_2.2 | 12 |
| PRAP.PPTUN_6.6 | 12 |
| PRAP.PPAFG_2.2 | 13 |
| PRAP.PPNS_20.20 | 13 |
| PRAP.PPNS_54.54 | 13 |
| PRAP.PPTUN_5.5 | 13 |
| PRAP.PPAFG_3.3 | 16 |

**S32 Table.** *Phlebotomus papatasi* population variant summary statistics.

| SampleName | TotalSites | Ref | Het | Hom | Missing | Singleton | Transitions | Transversions | Transition:Transversion |
|---|---|---|---|---|---|---|---|---|---|
| **PBAB-PBRG_8-8** | 6390876 | 2734387 | 103671 | 298671 | 3254147 | 214870 | 252744 | 149598 | 1.68949 |
| **PBAB-PBRG_9-9** | 6390876 | 2699700 | 90947 | 329871 | 3270358 | 227305 | 264798 | 156020 | 1.69721 |
| **PDAB-PDMA_1-1** | 6390876 | 2057503 | 58682 | 264612 | 4010079 | 269071 | 213182 | 110112 | 1.93605 |
| **PRAP-PPAFG_2-2** | 6390876 | 3890610 | 218019 | 317221 | 1965026 | 204137 | 340355 | 194885 | 1.74644 |
| **PRAP-PPAFG_3-3** | 6390876 | 3884284 | 205531 | 312332 | 1988729 | 196696 | 329417 | 188446 | 1.74807 |
| **PRAP-PPAFG_5-5** | 6390876 | 3887691 | 216063 | 337801 | 1949321 | 213510 | 352945 | 200919 | 1.75665 |
| **PRAP-PPAFG_6-6** | 6390876 | 3885638 | 215420 | 341971 | 1947847 | 215678 | 355465 | 201926 | 1.76037 |
| **PRAP-PPAFG_12-12** | 6390876 | 3916552 | 247294 | 383541 | 1843489 | 273833 | 400484 | 230351 | 1.73858 |
| **PRAP-PPNS_20-20** | 6390876 | 3959617 | 262945 | 289507 | 1878807 | 213926 | 350488 | 201964 | 1.7354 |
| **PRAP-PPNS_25-25** | 6390876 | 3922368 | 266284 | 287874 | 1914350 | 218120 | 352830 | 201328 | 1.75251 |
| **PRAP-PPNS_54-54** | 6390876 | 3927297 | 257149 | 293407 | 1913023 | 209690 | 350155 | 200401 | 1.74727 |
| **PRAP-PPNS_59-59** | 6390876 | 3922517 | 265489 | 294932 | 1907938 | 218594 | 357111 | 203310 | 1.75649 |
| **PRAP-PPNS_80-80** | 6390876 | 3900677 | 279634 | 305769 | 1904796 | 239966 | 373070 | 212333 | 1.757 |
| **PRAP-PPNS_82-82** | 6390876 | 3906115 | 259682 | 307694 | 1917385 | 222012 | 361406 | 205970 | 1.75465 |
| **PRAP-PPTUN_1-1** | 6390876 | 3908349 | 264739 | 296964 | 1920824 | 229982 | 357620 | 204083 | 1.75233 |
| **PRAP-PPTUN_2-2** | 6390876 | 3973523 | 277196 | 278867 | 1861290 | 226283 | 352787 | 203276 | 1.73551 |
| **PRAP-PPTUN_5-5** | 6390876 | 3948865 | 266104 | 290875 | 1885032 | 226419 | 353656 | 203323 | 1.73938 |
| **PRAP-PPTUN_6-6** | 6390876 | 3914971 | 259500 | 298662 | 1917743 | 225482 | 354995 | 203167 | 1.74731 |
| **PRAP-PPTUN_7-7** | 6390876 | 3969023 | 282740 | 302274 | 1836839 | 245355 | 371325 | 213689 | 1.73769 |
| **PRAP-PPTUN_12-12** | 6390876 | 3993035 | 293666 | 326856 | 1777319 | 274944 | 392836 | 227686 | 1.72534 |

**S33 Table.** *Phlebotomus papatasi* $F_{ST}$-Tajima's D overlap (including 10 kb upstream and downstream).

| AFG_Selected_Compared_With_EGP | EGP_Selected_Compared_With_AFG | AFG_Selected_Compared_With_TUN | TUN_Selected_Compared_With_AFG |
|---|---|---|---|
| PPAI011586 | PPAI011586 | PPAI011586 | PPAI001563 |
| PPAI012581 | PPAI012581 | PPAI012581 | |
| PPAI012653 | PPAI012653 | PPAI012653 | |
| PPAI005524 | PPAI002377 | PPAI005152 | |
| PPAI005525 | PPAI006365 | PPAI005524 | |
| PPAI006386 | PPAI009162 | PPAI005525 | |
| PPAI006387 | PPAI010152 | PPAI005603 | |
| PPAI006388 | PPAI010181 | PPAI005659 | |
| PPAI008927 | PPAI010182 | PPAI005660 | |
| PPAI009162 | PPAI010183 | PPAI009162 | |
| PPAI009163 | | PPAI009163 | |
| PPAI009633 | | PPAI009899 | |
| PPAI009634 | | PPAI009900 | |
| PPAI009899 | | PPAI009902 | |
| PPAI009900 | | PPAI010182 | |
| PPAI009902 | | PPAI010183 | |
| PPAI010151 | | PPAI010184 | |
| PPAI010182 | | | |
| PPAI010183 | | | |
| PPAI010184 | | | |

TUN: Tunisia, EGP: Egypt, AFG: Afghanistan.

| Gene | Description (*P. papatasi*; *Ortholog Description*) | Biological Process | Molecular Function | Cellular Component |
|---|---|---|---|---|
| PPAI011586 | tRNA-Lys | No Protein | No Protein | No Protein |
| PPAI012581 | tRNA-Gly | No Protein | No Protein | No Protein |
| PPAI012653 | tRNA-Lys | No Protein | No Protein | No Protein |
| PPAI005524 | Cuticular Protein | None Predicted | Structural constituent of cuticle | None Predicted |
| PPAI005525 | Cuticular Protein | None Predicted | Structural constituent of cuticle | None Predicted |
| PPAI006386 | No Description | None Predicted | None Predicted | None Predicted |
| PPAI006387 | No Description | None Predicted | None Predicted | None Predicted |
| PPAI006388 | No Description | | | |
| PPAI008927 | No Description | None Predicted | None Predicted | Membrane |
| PPAI009162 | Longitudinals lacking protein-like Transcription Factor | None Predicted | Protein Binding | None Predicted |
| PPAI009163 | oligoribonulcease, mitochondrial | None Predicted | 3'-5' exoribonuclease activity; nucleic acid binding | None Predicted |

| | | | | |
|---|---|---|---|---|
| PPAI009633 | Cytochrome P450 | oxidation-reduction process | monooxygenase activity;iron ion binding;oxidoreductase activity, acting on paired donors, with incorporation or reduction of molecular oxygen;heme binding | None Predicted |
| PPAI009634 | Cytochrome P450 | oxidation-reduction process | iron ion binding;oxidoreductase activity, acting on paired donors, with incorporation or reduction of molecular oxygen;heme binding | None Predicted |
| PPAI009899 | TATA box-binding protein-associated factor RNA polymerase I subunit C-like | | | |
| PPAI009900 | U3 small nucleolar RNA-associated protein homolog, putative | rRNA processing | protein binding | nucleolus |
| PPAI009902 | ubiquitin-conjugating enzyme E2 q | None Predicted | None Predicted | None Predicted |
| PPAI010151 | alpha amylase | carbohydrate [illegible] | catalytic activity;cation binding | None Predicted |
| PPAI010182 | Lipase | lipid metabolic [illegible] | carboxylic ester hydrolase activity | None Predicted |
| PPAI010183 | No Description | | | |
| PPAI010184 | NAD kinase-like | NADP biosynthetic [illegible] | NAD+ kinase activity | None Predicted |
| PPAI002377 | lipase 1 precursor | Lipid metabolic | None Predicted | None Predicted |
| PPAI006365 | Protein aurora boreallis | None Predicted | None Predicted | None Predicted |
| PPAI010152 | Six/sine homebox transcription factors | None Predicted | None Predicted | None Predicted |
| PPAI010181 | Putative lysozyme | None Predicted | lysozyme activity | None Predicted |
| PPAI005152 | poly(U)-specific endoribonuclease | None Predicted | endoribonuclease activity | None Predicted |
| PPAI005603 | No Description | None Predicted | None Predicted | None Predicted |
| PPAI005659 | cytochrome P450 | oxidation-reduction process | iron ion binding;oxidoreductase activity, acting on paired donors, with incorporation or reduction of molecular oxygen;heme binding | None Predicted |
| PPAI005660 | kekkon5 | None Predicted | protein binding | None Predicted |
| PPAI001563 | Calcium-transporting ATPase | calcium ion transmembrane transport | nucleotide binding;calcium-transporting ATPase activity;ATP binding | membrane;integral component of membrane |

2097

**S34 Table.** *Lutzomyia longipalpis* population sequencing median coverage depth.

| Sample | Median Coverage |
|---|---|
| IDMB.INT1 | 14 |
| IDMB.INT3 | 16 |
| IDMB.JAC01 | 64 |
| IDMB.JAC04 | 63 |
| IDMB.JAC05 | 44 |
| IDMB.JAC06 | 44 |
| IDMB.JAC08 | 53 |
| IDMB.JAC09 | 53 |
| IDMB.JAC11 | 52 |
| IDMB.JAC12 | 65 |
| IDMB.JAC14 | 32 |
| IDMB.JAC15 | 50 |
| IDMB.JAC16 | 37 |
| IDMB.JAC18 | 27 |
| IDMB.JAC22 | 105 |
| IDMB.JAC23 | 47 |
| IDMB.LAP01 | 41 |
| IDMB.LAP05 | 47 |
| IDMB.LAP06 | 62 |
| IDMB.LAP07 | 53 |
| IDMB.LAP08 | 61 |
| IDMB.LAP09 | 40 |
| IDMB.LAP10 | 46 |
| IDMB.LAP11 | 34 |
| IDMB.LAP12 | 22 |
| IDMB.LAP15 | 64 |
| IDMB.LAP16 | 80 |
| IDMB.LAP18 | 39 |
| IDMB.LAP20 | 46 |
| IDMB.Marajo06 | 50 |
| IDMB.Marajo10 | 8 |
| IDMB.Marajo15 | 54 |
| IDMB.Marajo16 | 44 |
| IDMB.Marajo17 | 51 |
| IDMB.Marajo18 | 57 |
| IDMB.Marajo19 | 53 |
| IDMB.Marajo20 | 58 |
| IDMB.Marajo21 | 74 |
| IDMB.Marajo22 | 49 |
| IDMB.MIGc6 | 15 |
| IDMB.MIGc9 | 18 |
| IDMB.S1S01 | 58 |
| IDMB.S1S02 | 48 |
| IDMB.S1S03 | 50 |
| IDMB.S1S04 | 33 |
| IDMB.S1S05 | 41 |
| IDMB.S1S06 | 35 |
| IDMB.S1S07 | 37 |
| IDMB.S1S08 | 36 |
| IDMB.S1S09 | 63 |

| | |
|---|---|
| IDMB.S1S10 | 40 |
| IDMB.S1S12 | 42 |
| IDMB.S1S13 | 37 |
| IDMB.S1S14 | 36 |
| IDMB.S2S01 | 65 |
| IDMB.S2S02 | 61 |
| IDMB.S2S03 | 43 |
| IDMB.S2S04 | 42 |
| IDMB.S2S07 | 72 |
| IDMB.S2S08 | 40 |
| IDMB.S2S09 | 36 |
| IDMB.S2S10 | 39 |
| IDMB.S2S11 | 45 |
| IDMB.S2S12 | 45 |
| IDMB.S2S13 | 48 |
| IDMB.S2S14 | 46 |
| IDMB.S2S15 | 51 |
| IDMB.S2S16 | 51 |
| IDMB.S2S17 | 49 |
| IDMB.S2S19 | 52 |

2099

**S35 Table.** Parameter values of male copulatory songs from *Lutzomyia longipalpis* from Araci and Olindina.

| | IPI / IBI (ms) | TL (s) | NB | FREQ |
|---|---|---|---|---|
| **Araci** | 57.8 ±4.6 | 2.1 ±0.7 | 36.4 ±11.3 | 242.3 ±12.0 |
| **Olindina** | 283.4 ±45.2 | 3.1 ±0.7 | 11.5 ±1.9 | 272.8 ±10.2 |

N, number of samples; IPI, inter-pulse interval; IBI, inter-burst interval; TL, train length; NB, number of bursts per train; NP, number of pulses per train; Freq, carrier frequency. Mean (±SE) values are presented.

**S36 Table.** *Lutzomyia longipalpis* genes within differentiation islands.

| DIFFERENTIATION ISLAND | GENES | DESCRIPTION | *ANOPHELES GAMBIAE ORTHOLOGUE ID* | PANTHER PATHWAY | PANTHER PROTEIN CLASS |
|---|---|---|---|---|---|
| **SCAFFOLD103 6001-7000** | LLOJ000261 | No Description | AGAP029318 | No Pathway | |
| **SCAFFOLD1090 32001-33000** | No Gene | | | | |
| **SCAFFOLD116 31001-32000** | No Gene | | | | |
| **SCAFFOLD116 39001-40000** | No Gene | | | | |
| **SCAFFOLD123 5001-6000** | LLOJ001208 | Protein MAK16 | AGAP000514 | No Pathway | |
| **SCAFFOLD1230 24001-25000** | LLOJ001226 | No Description | | | |
| **SCAFFOLD124 79001-80000** | No Gene | | | | |
| **SCAFFOLD1246 12001-13000** | LLOJ001273 | No Description | | | |
| **SCAFFOLD1246 25001-26000** | No Gene | | | | |
| **SCAFFOLD1261 25001-26000** | LLOJ001342 | No Description | AGAP000996 | No Pathway | |
| **SCAFFOLD1261 33001-34000** | LLOJ001342 | | | | |
| **SCAFFOLD1261 38001-39000** | LLOJ001342 | | | | |
| **SCAFFOLD1261 48001-49000** | LLOJ001343 | No Description | AGAP000520 | No Pathway | |
| **SCAFFOLD129 79001-80000** | No Gene | | | | |
| **SCAFFOLD1299 13001-14000** | No Gene | | | | |
| **SCAFFOLD13 375001-376000** | No Gene | | | | |
| **SCAFFOLD1343 24001-25000** | No Gene | | | | |
| **SCAFFOLD135 120001-121000** | No Gene | | | | |
| **SCAFFOLD135 84001-85000** | LLOJ001645 | No Description | | | |
| **SCAFFOLD1393 14001-15000** | No Gene | | | | |
| **SCAFFOLD1435 5001-6000** | LLOJ001919 | No Description | AGAP004691 | No Pathway | |
| **SCAFFOLD145 88001-89000** | LLOJ001959 | No Description | AGAP004772 | PDGF Signaling | |
| **SCAFFOLD1528 5001-6000** | No Gene | | | | |
| **SCAFFOLD1602 5001-6000** | No Gene | | | | |
| **SCAFFOLD1655 21001-22000** | LLOJ002503 | No Description | | | |
| **SCAFFOLD1821 12001-13000** | LLOJ002897 | No Description | AGAP002567 | | Triacylglyerol metabolism |
| **SCAFFOLD1956 13001-14000** | LLOJ003168 | No Description | AGAP010621 | No Pathway | cation transporter |
| **SCAFFOLD1956 19001-20000** | LLOJ003168 | | | | |
| **SCAFFOLD1956 2001-3000** | No Gene | | | | |
| **SCAFFOLD197 118001-119000** | No Gene | | | | |
| **SCAFFOLD197 124001-125000** | LLOJ003204 | No Description | | | |

| | | | | | |
|---|---|---|---|---|---|
| **SCAFFOLD2 513001-514000** | LLOJ003298 | No Description | | | |
| **SCAFFOLD2014 1001-2000** | LLOJ003364 | No Description | AGAP001425 | No Pathway | |
| **SCAFFOLD21 94001-95000** | LLOJ003528 | No Description | AGAP011870 | Did not map | |
| **SCAFFOLD2144 2001-3000** | No Gene | | | | |
| **SCAFFOLD234 60001-61000** | LLOJ003942 | No Description | | | |
| **SCAFFOLD2467 16001-17000** | LLOJ004171 | No Description | AGAP002321 | No Pathway | |
| **SCAFFOLD2527 9001-10000** | LLOJ004261 | No Description | | | |
| **SCAFFOLD274 77001-78000** | No Gene | | | | |
| **SCAFFOLD282 96001-97000** | LLOJ004737 | No Description | | | |
| **SCAFFOLD283 3001-4000** | No Gene | | | | |
| **SCAFFOLD288 3001-4000** | No Gene | | | | |
| **SCAFFOLD320 65001-66000** | LLOJ005134 | No Description | AGAP000635 | No Pathway | |
| **SCAFFOLD320 77001-78000** | LLOJ005134 | | | | |
| **SCAFFOLD3248 1001-2000** | No Gene | | | | |
| **SCAFFOLD330 65001-66000** | No Gene | | | | |
| **SCAFFOLD3435 1001-2000** | No Gene | | | | |
| **SCAFFOLD3460 2001-3000** | No Gene | | | | |
| **SCAFFOLD448 9001-10000** | LLOJ006400 | No Description | AGAP007588 | No Pathway | |
| **SCAFFOLD5279 1001-2000** | No Gene | | | | |
| **SCAFFOLD537 37001-38000** | No Gene | | | | |
| **SCAFFOLD537 39001-40000** | No Gene | | | | |
| **SCAFFOLD537 74001-75000** | No Gene | | | | |
| **SCAFFOLD544 25001-26000** | LLOJ007206 | No Description | AGAP000998 | No Pathway | Receptor |
| **SCAFFOLD564 67001-68000** | No Gene | | | | |
| **SCAFFOLD564 88001-89000** | No Gene | | | | |
| **SCAFFOLD5940 1001-2000** | No Gene | | | | |
| **SCAFFOLD611 24001-25000** | LLOJ007782 | No Description | | | |
| **SCAFFOLD611 24001-25000** | LLOJ007783 | No Description | AGAP013509 | No Pathway | lipase/serine protease |
| **SCAFFOLD611 24001-25000** | LLOJ007784 | No Description | AGAP029597 | Did not map | |
| **SCAFFOLD611 44001-45000** | LLOJ007785 | No Description | AGAP028418 | Did not map | |
| **SCAFFOLD611 58001-59000** | LLOJ007786 | No Description | | | |
| **SCAFFOLD611 7001-8000** | LLOJ007780 | No Description | AGAP011717 | No Pathway | |
| **SCAFFOLD611 81001-82000** | LLOJ007790 | No Description | | | |
| **SCAFFOLD6532 8001-9000** | No Gene | | | | |
| **SCAFFOLD6593 1001-2000** | No Gene | | | | |
| **SCAFFOLD6738 1-1000** | No Gene | | | | |

| | | | | | |
|---|---|---|---|---|---|
| **SCAFFOLD71 101001-102000** | LLOJ008470 | No Description | AGAP008938 | Did not map | |
| **SCAFFOLD71 104001-105000** | LLOJ008470 | | | | |
| **SCAFFOLD7298 1-1000** | No Gene | | | | |
| **SCAFFOLD735 56001-57000** | LLOJ008643 | No Description | AGAP011344 | No Pathway | |
| **SCAFFOLD7663 1-1000** | No Gene | | | | |
| **SCAFFOLD778 6001-7000** | No Gene | | | | |
| **SCAFFOLD789 17001-18000** | No Gene | | | | |
| **SCAFFOLD7896 1-1000** | No Gene | | | | |
| **SCAFFOLD8362 1001-2000** | No Gene | | | | |
| **SCAFFOLD8384 1001-2000** | No Gene | | | | |
| **SCAFFOLD8385 1-1000** | No Gene | | | | |
| **SCAFFOLD85 8001-9000** | LLOJ009304 | No Description | AGAP028534 | Did not map | |
| **SCAFFOLD8639 1001-2000** | No Gene | | | | |
| **SCAFFOLD8700 1-1000** | No Gene | | | | |
| **SCAFFOLD872 21001-22000** | LLOJ009447 | rRNA adenine N(6)- | AGAP012129 | No Pathway | RNA methyltransferase |
| **SCAFFOLD879 52001-53000** | No Gene | | | | |
| **SCAFFOLD8881 3001-4000** | No Gene | | | | |
| **SCAFFOLD919 1001-2000** | No Gene | | | | |
| **SCAFFOLD919 19001-20000** | No Gene | | | | |
| **SCAFFOLD92 27001-28000** | LLOJ009732 | Lipase maturation | AGAP011922 | Did not map | |
| **SCAFFOLD92 27001-28000** | LLOJ009733 | No Description | AGAP001563 | No Pathway | carbohydrate kinase |
| **SCAFFOLD9363 1001-2000** | | | | | |
| **SCAFFOLD94 52001-53000** | LLOJ009833 | No Description | AGAP012342 | Notch Signaling | basic helix-loop-helix transcription factor |
| **SCAFFOLD942 19001-20000** | No Gene | | | | |
| **SCAFFOLD942 21001-22000** | No Gene | | | | |
| **SCAFFOLD9630 1-1000** | No Gene | | | | |
| **SCAFFOLD97 67001-68000** | No Gene | | | | |
| **SCAFFOLD977 22001-23000** | No Gene | | | | |

2106
